# Clustering of RNA co-expression network identifies novel long non-coding RNA biomarkers in squamous cell carcinoma

**DOI:** 10.1101/2023.12.20.571624

**Authors:** Liisa Nissinen, Josefiina Haalisto, Pilvi Riihilä, Minna Piipponen, Veli-Matti Kähäri

**Author notes:** **Corresponding author:** Professor Veli-Matti Kähäri, M.D., PhD, Department of Dermatology, University of Turku and Turku University Hospital, Hämeentie 11 TE6, FI-20520 Turku, Finland.

## Abstract

Long non-coding RNAs (lncRNAs) have been shown to play an important role in cancer progression. Cutaneous squamous cell carcinoma is the most common metastatic skin cancer with increasing incidence worldwide. The prognosis of the metastatic cSCC is poor, and currently there are no established biomarkers to predict metastatic risk nor specific therapeutic targets for advanced or metastatic cSCC. To elucidate the role of lncRNAs in cSCC, RNA sequencing of patient derived cSCC cell lines and normal human epidermal keratinocytes was performed. The correlation analysis of differentially expressed lncRNA and protein-coding genes revealed six distinct clusters. One of the upregulated clusters involved genes related to cell motility. Upregulation of the expression of lncRNAs involved in cSCC cell motility in cSCC and head and neck SCC (HNSCC) cells was confirmed by qRT-PCR. Upregulation of *HOTTIP* and *LINC00543* was also noted in SCC tumors *in vivo* and was associated with worse prognosis in HNSCC and lung SCC cohorts in the TCGA data, respectively. Altogether, these results reveal a novel set of lncRNAs involved in cSCC cell locomotion. These lncRNAs may serve as potential novel biomarkers or a biomarker panel and as putative therapeutic targets in locally advanced and metastatic cSCC.

## INTRODUCTION

Long non-protein coding RNAs (lncRNAs) have been identified as important cellular regulators of tissue homeostasis and pathological conditions including cancer initiation, growth and metastasis (1). LncRNAs with transcript length of 200 nucleotides or longer are a functionally diverse group of regulatory RNAs that may interact with DNA, proteins or other RNAs (2). They are strictly regulated in spatial and temporal manner making lncRNAs versatile regulators of cellular processes in all cellular compartments and also interesting therapeutic targets in pathological conditions (3).

Keratinocyte-derived cutaneous squamous cell carcinoma (cSCC) is the most common metastatic skin cancer and the incidence of this non-melanoma skin-cancer is increasing worldwide due to life-style changes and an ageing of the population (4). The metastasis rate of primary cSCC is estimated as 3–5 % and it is responsible for at least 20% of all skin cancer-related mortality (5,6). The most important risk factor for cSCC is cumulative lifetime exposure to UV radiation (7). This makes cSCC a cancer with a particularly high mutation rate (7). Tumor protein 53 (*TP53*) gene is mutated early during keratinocyte carcinogenesis resulting in a marked accumulation of additional UV-induced mutations in cSCC progression (8-10). Subsequent genomic alterations in oncogenes such as *NOTCH1, HRAS, CDKN2A* and *EGFR* further contribute to the disease progression (11,12). As epidermal keratinocytes in sun-exposed normal skin already harbor several mutations in the genes associated with cSCC progression, alterations in the tumor microenvironment are crucial for initiation and progression of cSCC (13,14).

LncRNAs have been studied in cSCC but their role in cSCC progression is largely unknown (15). We have previously characterized and named three lncRNAs that are upregulated in cSCC and regulate progression of cSCC via different mechanisms, namely *PICSAR* (p38 inhibited cutaneous squamous cell carcinoma-associated lincRNA) (16,17), *PRECSIT* (p53 regulated carcinoma-associated STAT3-activating long intergenic non-protein coding transcript) (18) and *SERLOC* (super enhancer and ERK1/2-regulated long Intergenic non-protein coding transcript overexpressed in carcinomas) (19). Additionally, *PVT1* was discovered as a lncRNA that promotes cSCC progression by suppressing cellular senescence by inhibiting *CDKN1A* expression and preventing cell cycle arrest (20,21).

In this study, to elucidate the role of lncRNAs in cSCC, RNA sequencing (RNA-seq) of patient derived cSCC cell lines and normal human epidermal keratinocytes (NHEK) was performed. The correlation analysis of differentially expressed lncRNA and protein-coding genes revealed six clusters. Cluster 1 was upregulated in cSCC cells and it involved genes related to cell motility. Further studies of the lncRNAs involved in this cSCC cell motility cluster revealed that they were also upregulated in patient derived head and neck SCC (HNSCC) cells. Interestingly, the expression of *HOTTIP* and *LINC00543* were associated with worse prognosis in HNSCC and lung SCC (LUSCC) cohorts in the TCGA data, respectively. Altogether, these results reveal a novel set of lncRNAs involved in cSCC cell locomotion. These lncRNAs may serve as novel biomarkers or in a biomarker panel and also as potential therapeutic targets in advanced cSCC.

## MATERIALS AND METHODS

### Ethical Issues

The use of tumor derived SCC cell lines and NHEKs and collection of cSCC tissues was approved by the Ethics Committee of the Hospital District of Southwest Finland. All participants gave their written informed consent, and the study was performed with the permission of Turku University Hospital, according to the Declaration of Helsinki.

### Cell Culture

NHEKs (n=4) were isolated from the skin of healthy individuals undergoing mammoplasty (16). NHEK-PC, NHEK (adult pooled), NHEK (adult single donor), HEK, NHEK were from PromoCell (Heidelberg, Germany), HEK and HOK (human oral keratinocytes) were from ScienCell Research Laboratories (Carlsbad, CA, USA). Primary non-metastatic (UT-SCC12A, UT-SCC91, UT-SCC105, UT-SCC111, and UT-SCC118) and metastatic (UT-SCC7, UT-SCC59A, and UT-SCC115) cSCC cell lines were established from surgically removed cSCCs in Turku University Hospital (22). The authenticity of cSCC cell lines was verified by short tandem repeat profiling as described previously (22). HNSCC cell lines (n=29) were established from surgically removed HNSCCs in Turku University Hospital (23,24). The immortalized non-tumorigenic human keratinocyte–derived cell line (HaCaT) and three Ha-*ras*-transformed tumorigenic HaCaT cell lines (A5, II-4, and RT3) were kindly provided by Dr Norbert Fusenig (Deutsche Krebsforschungszentrum, Heidelberg, Germany). A5 cells form benign, II-4 cells invasive and RT3 cells metastatic tumors in nude mice *in vivo* (24). Cells were cultured as previously described (18). For growth factor treatment cSCC cell lines (n = 2-8) were maintained in serum-free medium for 24 hours and then treated with transforming growth factor-β1 (5 ng/mL; Sigma Aldrich, St Louis, MO) for 24 hours.

### Tissue samples

Normal human skin samples (n = 6) were obtained from the upper arm of healthy volunteers or during mammoplasty operation in Turku University Hospital. Primary cSCC (n = 6) samples were collected from surgically removed tumors in Turku University Hospital (16).

### RNA Sequencing

RNA was isolated from cSCC cell lines (n = 8) and NHEKs (n = 4) using miRNeasy Mini Kit (Qiagen), and the RNA-seq analysis was performed using Illumina HiSeq2500 system using paired-end sequencing chemistry with 100bp read length (Illumina, San Diego, CA) at the Finnish Functional Genomics Centre, University of Turku, Åbo Akademi University and Biocenter Finland. The reads were aligned against the human GRCh38 (hg38) genome assembly using STAR alignerr, version 2.6.1 (26). Gene-level read counts were obtained simultaneously with the alignment process. All subsequent analysis steps were done using R (27). Read count data was filtered to exclude genes with very low read counts (less than ten reads in total in all samples). The obtained read counts were normalized using regularized log transformation function of DESeq2 R package, v. 1.20.0, which transforms the count data to the log2 scale in a way that minimizes differences between samples for rows with small counts and also normalizes the data with respect to library size. Differential expression (DE) analysis was performed using DESeq2, v. 1.20.0. P-values were adjusted using the Benjamini-Hochberg multiple testing adjustment method (28). Genes with absolute log2 fold change > 1 and adjusted p < 0.05 were considered as significantly differentially expressed. Differentially expressed genes were annotated with HGNC gene symbols, gene description and gene biotype using biomaRt, v. 2.36.1 (29). The results were split into differentially expressed protein-coding genes and lncRNAs based on the gene biotype annotations, using Ensembl biotype definitions as guidance.

### Correlation and clustering analysis

A combined dataset of significantly differentially expressed lncRNAs and protein coding genes was generated and pairwise Pearson’s correlations of the expression profiles of each gene were computed. Correlation distances (1 - correlation coefficient) were used as input for k-means clustering with 6 centres, generating clusters of co-expressed genes, i.e. genes with similar expression profiles across all samples. Correlation distance matrix was subjected to classical multidimensional scaling (MDS), or principal coordinates analysis, to be able to visualize the clustering result in a two-dimensional plot generated using ggplot2 (30). Correlation densities within each of the clusters were visualized as histograms using basic R functions and custom R scripts. Distributions of protein coding gene and lncRNA expression values in each cluster were examined using box plots generated using ggplot2 (30). Gene expression values of all six clusters were also visualized as a heatmap using R package pheatmap (31).

### Functional analysis

Enrichment of Gene ontology (GO) (32,33) terms in the genes in k-means clusters was studied using R package clusterProfiler, v.3.8.1 (34). The p-values of enrichment analysis were corrected for multiple testing using Benjamini-Hochberg multiple testing adjustment procedure (28). For the cluster genes, the enrichment analysis was conducted using both lncRNAs and mRNAs.

Ingenuity Pathway Analysis (IPA) (35) was performed separately for the k-means gene cluster including both lncRNAs and mRNAs. As before, genes with absolute log2 fold change > 1 and adjusted p < 0.05 were considered as significantly differentially expressed and were used in the analyses. P-values were corrected for multiple testing using Benjamini-Hochberg method (28). IPA also calculated for each pathway a z-score that indicated predicted pathway activation if z-score > 2 or inhibition if z-score < -2.

DAVID Bioinformatics Resources was used to study the genes in k-means clusters (36,37). It calculates the probability that particular GO annotations are overrepresented in a given gene list using a Fisher exact probability test. Molecular function and biological process GO terms with a *p* value <0.05 containing at least three genes were considered significant.

### Transcription factor annotation

To annotated potential transcription factors (TF) regulating the transcription of each gene experimentally validated TFs from Transcriptional Regulatory Element Database (TRED) (38) via RegNetwork (39), Encyclopedia of DNA Elements (ENCODE) and ChIP-X Enrichment Analysis (CHEA) database (40) were collected and combined to generate unique TF - TF target gene pairs. To visualize TFs in cluster 1 https://www.wordclouds.com was used.

### Real-Time Quantitative PCR

Total RNA was extracted from cultured NHEKs and cSCC cells using an RNeasy mini kit (Qiagen, Germantown, MD, USA), and 1 μg of total RNA was reverse transcribed into cDNA with random hexamer and M-MLV Reverse Transcriptase H Minus (both from Promega, Madison, WI, USA) for real-time quantitative reverse transcriptase-PCR (qRT-PCR) analysis. Primers and probe for *LINC01361* (Hs03084701_cn, Cat nro 4400291), *LINC01558* (Hs00205026_m1, Cat nro 4351372), *LINC01460* (Hs03405179_cn, Cat nro 4400291), *LINC00543* (Hs00905601_s1, Cat nro 4426961), *LINC00702* (Hs05184596_cn, Cat nro 4400291) and *HOTTIP* (Hs04965481_cn, Cat nro 4400291) were purchased from Thermo Fisher Scientific (Waltham, MA, USA). Primers and probes for *MMP13* and β-actin (*ACTB*) were designed as previously described (24). qRT-PCR reactions were performed utilizing the QuantStudio 12K Flex (Thermo Fisher Scientific) at the Finnish Functional Genomics Centre in Turku, Finland. qRT-PCR amplification was done using the following protocol: hold stage 2 min at 50 °C, 10 min at 95 °C, and PCR stage for 40 cycles 0.15 min at 95 °C and 1 min at 60 °C. Samples were analyzed using the standard curve method in 2-3 parallel reactions with threshold cycle values <5% of the mean threshold cycle.

### Gene expression profiling interactive analysis

The online Gene Expression Profiling Interactive Analysis (GEPIA; http://gepia.cancer-pku.cn/) analysis tool was utilized to analyze the expression of *HOTTIP* and *LINC00543* in SCCs involved in the database and the relationship between *HOTTIP* and *LINC00543* expression and prognosis of HNSCC and LUSCC in The Cancer Genome Atlas (TCGA) data (41,42).

## RESULTS

### The expression profile of lncRNAs and protein coding genes in cSCC cells and NHEKs

To determine differentially expressed lncRNAs and protein-coding genes, the RNA-seq of primary non-metastatic (n = 5) and metastatic (n = 3) cSCC cell lines and NHEKs was performed. Differentially expressed genes between cSCC and NHEKs were considered as significantly differentially expressed, if adjusted p-value (FDR) was less than 0.05 and absolute log2 fold change of expression level over 1. The genes belonging into two categories, either lncRNAs or protein-coding genes, were characterized further. The count of DE lncRNA genes was 723 and of DE protein-coding genes 1667. To reveal the distribution of differentially expressed lncRNAs (Supplement Figure S1A) and protein coding genes (Supplement Figure S1B) volcano blots were constructed.

The top 50 most significantly differentially expressed lncRNAs (Figure 1A) and protein coding genes (Figure 1B) are presented as a heatmap. As previously shown in cSCC, among the top downregulated lncRNAs there was *LINC00520* (Figure 1A) (43) and among top upregulated there were *PRECSIT* (*LINC00346*) (Figure 1A) (18)) and inflammasome component *AIM2* (Figure 1A) (44).

**Figure 1.**
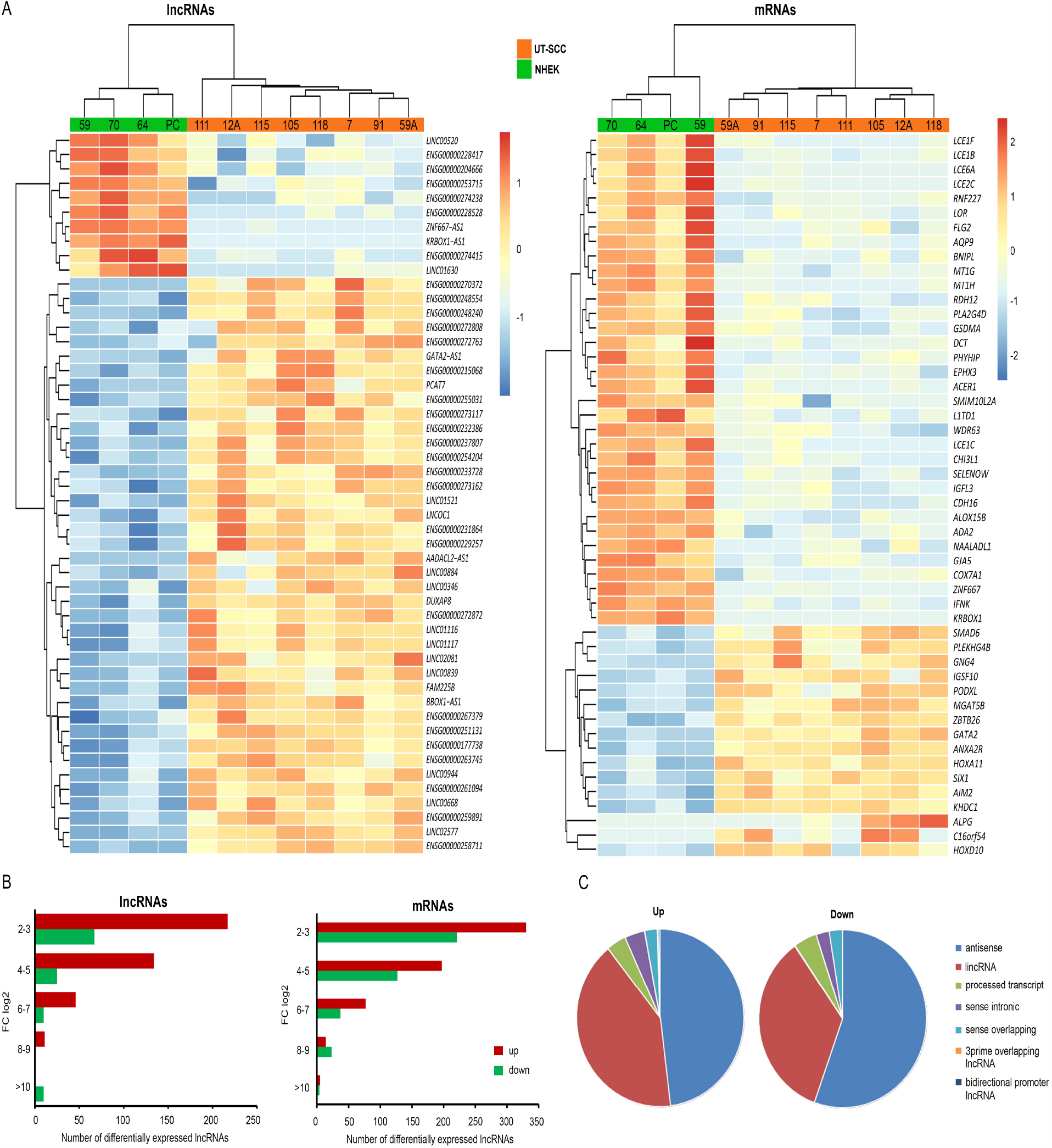
Profile of the differentially expressed lncRNAs and mRNAs in cSCC cells compared to NHEKs. RNA-seq of patient derived cSCC cell lines (n = 8) and normal human epidermal keratinocytes (NHEK, n = 4) was prepared. (**A**) Heatmaps and the hierarchical clustering of top differential expressed lncRNAs and mRNAs in cSCC cells compared to NHEKs. (**B**) The number of differentially expressed lncRNAs and mRNAs based on fold changes (FC log2). (**C**) Biotype classification of up and downregulated lncRNAs. All biotypes discovered in the data are shown.

Concerning the lncRNA expression with more than 8 fold change (log2), there were 11 upregulated genes, 5 of which were novel transcripts (Figure 1B, Supplement Table S1). In addition there was one >10 FC (log2) downregulated antisense lncRNA, *ZNF667-AS1* (ZNF667 antisense RNA 1) (Figure 1B, Supplement Table S1). *LINC01361* and *HOTTIP* and *LINC01558* were among upregulated lncRNAs ((FC (log2) >5 and >4, respectively) (Figure 1B, Supplement Table S1). The majority of the differentially expressed genes belonged to the 2-3 FC (log2) group e.g. *LINC00543, LINC00702* and *LINC01460* (Figure 1B, Supplement Table S1). Similarly to the differentially expressed lncRNAs, 5 upregulated protein coding genes (e.g. *MMP13*) and 4 downregulated ones were identified with >10 FC (log2), and the group with most differentially expressed protein coding genes was 2-3 FC (log2) (Figure 1B, Supplement Table S2).

### Classification of differentially expressed lncRNA biotypes

Altogether 568 lncRNAs were significantly upregulated (adjusted p-value < 0.05, FC (log2) 1) in cSCC cells and the majority of these were antisense or lincRNAs (48% and 41%, respectively) (Figure 1C). The number of significantly downregulated lncRNAs (adjusted p-value < 0.05, FC (log2) 1) in cSCC cells was 155 and the vast majority of these were antisense RNAs (55%) (Figure 1C). Additionally, 35% of significantly downregulated lncRNAs were lincRNAs (Figure 1C).

### Co-expression networks of the lncRNAs and protein coding genes

The differentially expressed protein-coding genes and lncRNAs were combined and pairwise Pearson’s correlation coefficients were computed for their expression values across the samples (Figure 2). k-means clustering using six centres was performed based on the correlation distances and the clustering of the genes is shown as MDS plot and the clusters are shown in different colors (Figure 2A). Pairwise Pearson’s correlations were calculated of the expression values across the samples within each cluster to determine the expression profile similarity among the cluster genes, and correlation distributions were visualized using histograms (Figure 2B). Average correlations of the clusters were also calculated (Supplement Table S3). The within-cluster correlations show that cluster 6 had the highest average correlation as well as the most narrow distribution of the correlation values (Supplement Table S3). Cluster 5, on the other hand, displayed bimodal distribution of the correlation values, and in MDS plot the genes in that cluster do not form a clearly defined group, suggesting that these genes had variable expression profiles that lacked clear correlation with the expression profiles of all the other genes (Figure 2A, B; Supplement Table S3).

**Figure 2.**
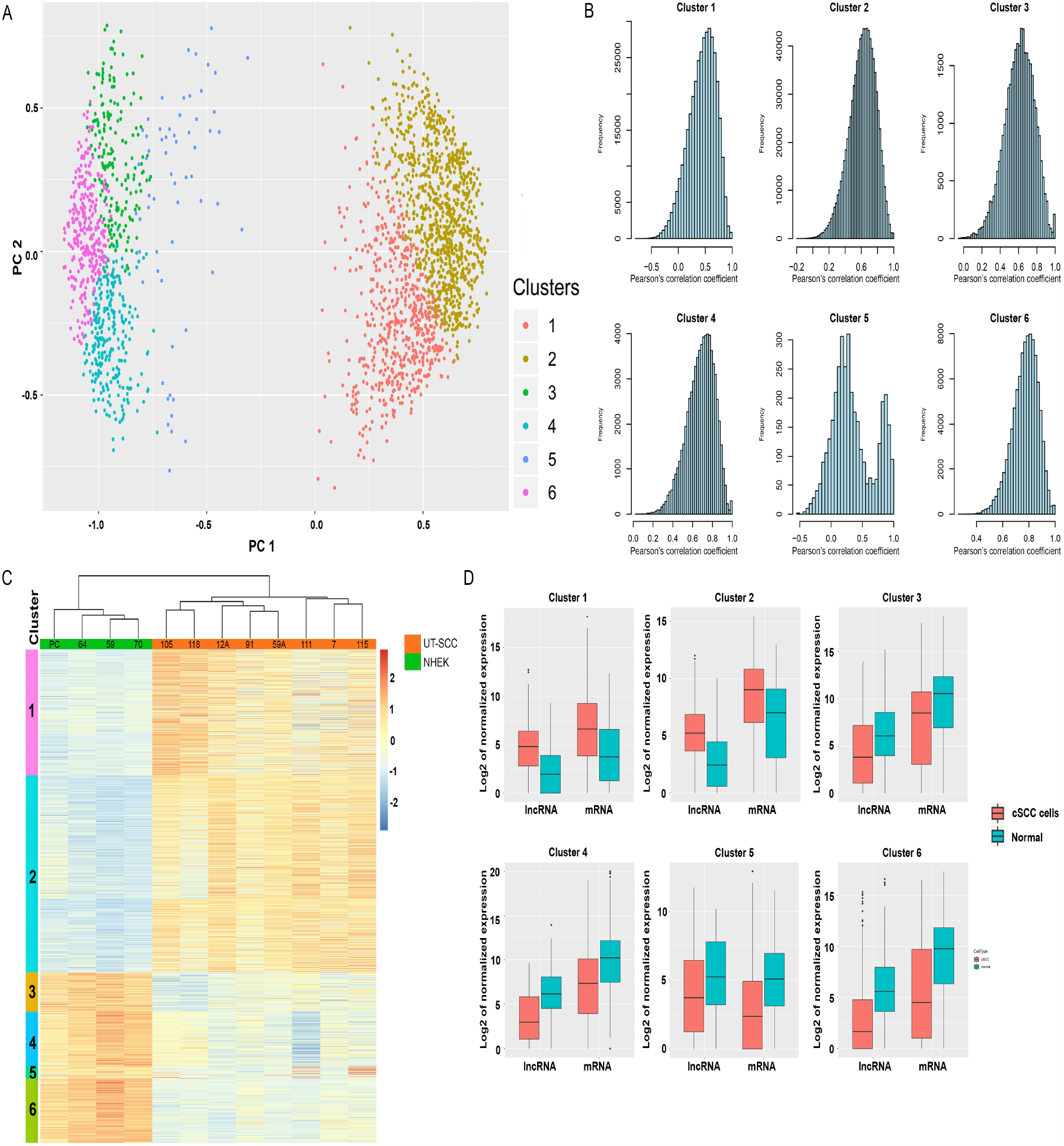
Construction of the co-expression network of lncRNAs and mRNAs. (**A**) Differentially expressed lncRNAs and mRNA in cSCC cells compared to NHEKs were joined and pairwise Pearson’s correlation coefficients were computed for their expression values. k-means clustering using six centres was performed based on the correlation distances. Clustering of the genes is shown as a multidimensional scaling (MDS) plot. In the plot, the principal coordinates of the pairwise correlation distances are shown on the axes, genes as dots and the clusters of genes in different colors. (**B**) Pairwise Pearson’s correlations were calculated of the expression values across the samples within each cluster to determine the expression profile similarity among the cluster genes, and correlation distributions were visualized using histograms. (**C**) Gene expression profiles of the clusters are visualized by a heatmap of the expression values of all clustered genes across the samples. (**D**) The box plot visualization shows the expression value distributions in cSCC cell lines (n = 8) and NHEKs (normal, n = 4) for lncRNAs and mRNAs belonging to one cluster.

### Cluster 1 and 2 genes are upregulated in cSCC cells

Gene expression profiles of the clusters were visualized by generating a heatmap of the expression values of all clustered genes across the samples (Figure 2C). The cluster 1 and 2 genes were upregulated, whereas the cluster 3, 4, 5 and 6 genes were downregulated in cSCC cells compared to NHEKs (Figure 2C). The expression values of the cluster genes were further inspected by generating box plot visualizations, showing expression value distributions in cSCC cells and NHEKs separately for the protein-coding genes and lncRNAs belonging to the same cluster (Figure 2D).

### Genes in Cluster1 are involved in cSCC cell motility

The upregulated Cluster 1 (Figure 3) was studied in more detail. Pathway analysis of the genes associated with cluster 1 revealed significant upregulation of the IPA biofunction categories *Cell movement, Migration of cells* and *Invasion of cells* (Figure 3A). This association was also supported by GO analysis. GO terms related to Biological process associated with cluster 1 genes included *Cell migration* and *Locomotion* (Figure 3B). GO terms related to Cellular component associated with cluster 1 genes included *Cell periphery* and *Plasma membrane* (Figure 3C). David analysis of cluster 1 genes showed the association of the protein coding genes with terms *Endopeptidase activity, Cell migration* and *Extracellular matrix disassembly* (Supplement Table S4). A network of the cluster 1 genes related to term *Extracellular space* (Supplement Figure S2) was formed by taking the protein–protein functional interaction data from STRING database (https://string-db.org/, August 16, 2023) (45). Interestingly, a previously known cSCC cell invasion related matrix metalloproteinase, MMP-13, was associated with this term (Supplement Figure S2) (46).

**Figure 3.**
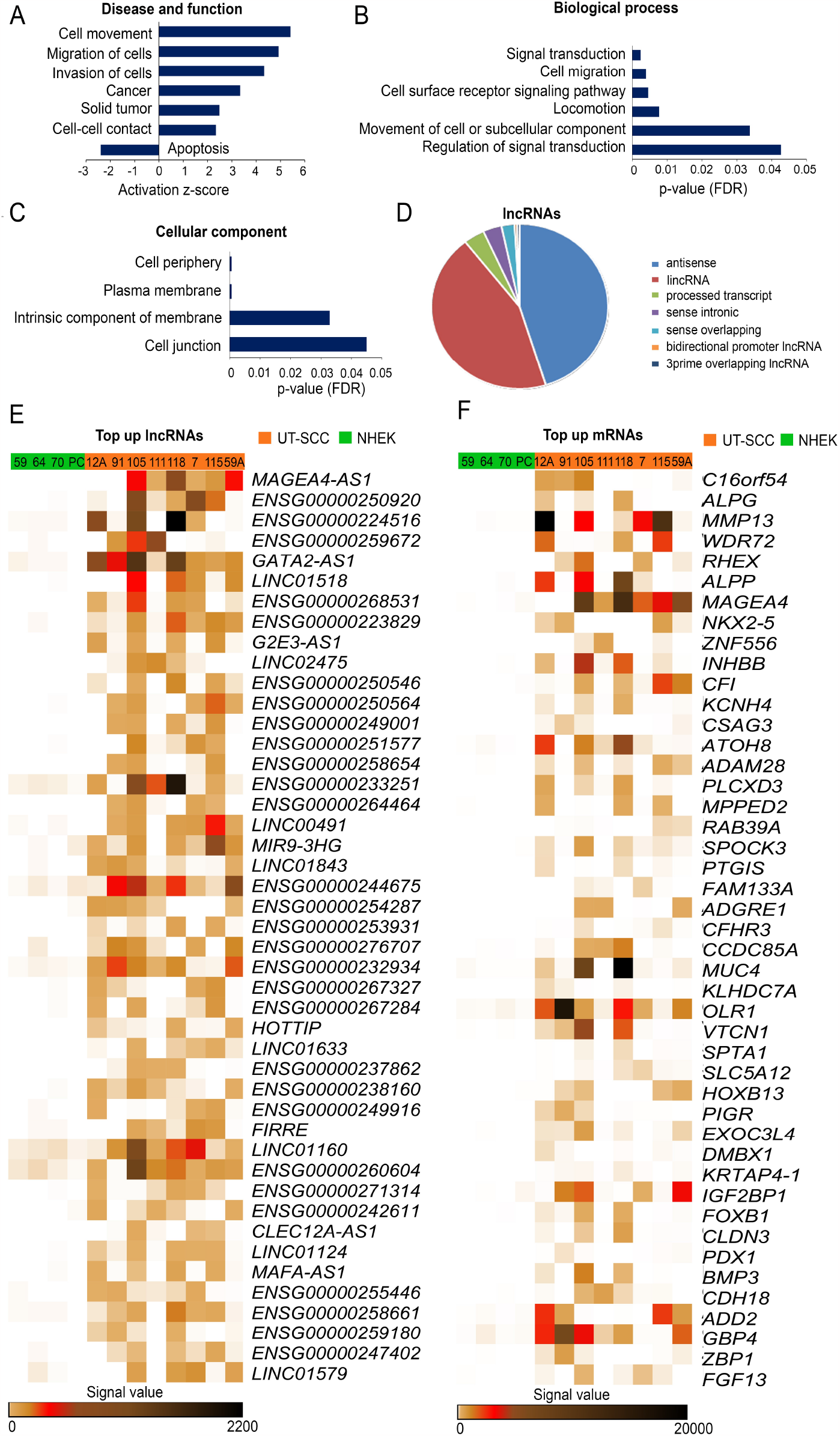
Cluster 1 genes are involved in cSCC cell motility. Summary of significantly regulated IPA biofunctions (**A**), the Gene ontology (GO) Biological process (**B**) and GO Cellular component (**C**) associated with genes involved in cluster 1. (**D**) Biotype classification of lncRNAs involved in cluster 1. All biotypes discovered in the data are shown. Heatmap visualization of top upregulated lncRNAs (**E**) and mRNAs (**F**) in cSCC cell lines (n = 8) compared to NHEKs (n = 4).

### Classification of differentially expressed lncRNA biotypes in cluster 1

LncRNA biotypes in cluster 1 were studied in more detail. In cluster 1 there were Altogether 208 lncRNAs, the majority of which were antisense or lincRNAs (45% and 44%, respectively) (Figure 3D). Among the top regulated lncRNAs in cluster 1 there were several novel transcripts and previously uncharacterized lincRNAs, and some previously characterized ones, like *HOTTIP* (47) and *LINC00491* (48) that have previously been shown to be involved in cancer cell invasion (Figure 3E) (Supplement Table S5). Among interesting lncRNAs in cluster 1 there were novel transcripts, such as *LINC01361, LINC01460* and *LINC01558* and RNA transcripts *LINC00543* and *LINC00702* that have not previously been identified in SCCs (Supplement Table S5). The expression of selected lncRNAs, both novel and the ones not characterized in cSCC, genes belonging to cluster 1 in cSCC cells and NHEKs was demonstrated as heatmap by RNA-seq (Supplement Figure S3). Interestingly, cSCC invasion related *MMP13* and complement factor I (*FI*) were among the top upregulated protein coding genes in cluster 1 (Figure 3F) (Supplement Table S5).

Cluster 1 genes were also annotated with potential upstream regulator transcription factors (TFs) (Supplement Figure S4, Supplement Table S6). Only experimentally validated TFs were included in the annotations. Myc was shown to be upstream regulator for 377 genes and EZH2 for 368 genes (Supplement Figure S4). These TFs were demonstrated to activate the transcription of genes coding for possible cancer progression related molecules, for example Myc regulating *FI* and EZH2 regulating *MMP13* and *FI* (Supplement Table S6).

### Validation of differentially expressed lncRNAs in cSCC

The expression of selected lncRNAs in cSCC cells was verified by qRT-PCR (Figure 4A). *HOTTIP, LINC00543, LINC00702, LINC01361, LINC01460*, and *LINC01558* were significantly upregulated in cSCC cells compared to NHEKs (Figure 4A). As a control, mRNA levels of *MMP13* were noted to be upregulated in cSCC cells (Figure 4A).

**Figure 4.**
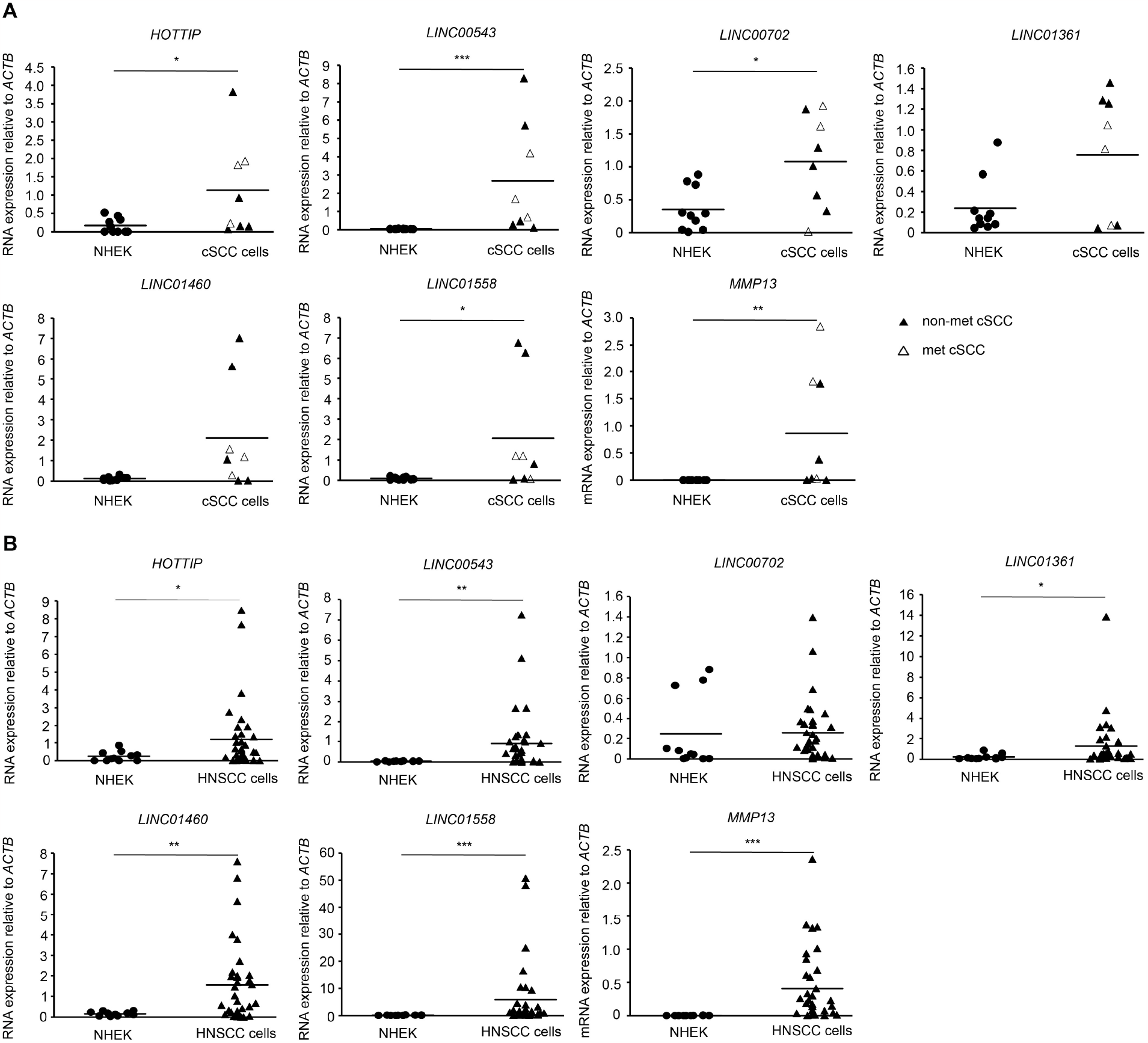
Validation of differentially expressed lncRNAs in cSCC and HNSCC cell lines. The RNA expression levels of *LINC00543, LINC00702, LINC01361, LINC01460, LINC01558, HOTTIP* and *MMP13* were determined by qRT-PCR in cSCC (**A**) and HNSCC (**B**) cells and NHEKs (A n = 10), B n = 11). β-actin mRNA (*ACTB*) levels were determined as a reference gene. *p<0.05; **p<0.01, ***p<0.001, Mann–Whitney U-test.

### Expression of lncRNAs in HNSCC cell lines

The expression of these selected lncRNAs was also investigated in patient derived HNSCC cells with qRT-PCR (Figure 4A). Significant overexpression of *LINC00543, LINC01361, LINC01460, LINC01558* and *HOTTIP* was detected in HNSCC cells compared to normal keratinocytes (Figure 4B). Additionally, *MMP13* was also significantly upregulated in HNSCC cells, as previously shown (23,24) (Figure 4B).

### Expression of *LINC00702, LINC01558* and *HOTTIP* in Ha-ras-transformed tumorigenic HaCaT cells

To further elucidate the role of the invasion cluster related lncRNAs during the progression of cSCC, the expression of *LINC00702, LINC01558, HOTTIP* and *MMP13* was determined in immortalized non-tumorigenic keratinocyte–derived cell line (HaCaT) lacking functional p53 and in three Ha-ras-transformed tumorigenic HaCaT-derived cell lines with different tumorigenicity *in vivo* (A5, II-4, and RT3, 24). Other selected lncRNAs were not expressed in this *in vitro* model of cSCC progression. *LINC00702* was expressed only in RT3 cells, the most aggressive ras-transformed tumorigenic HaCaT-derived cell line (Figure 5A). The expression of *LINC01558* was low in HaCaT and A5 cells, whereas markedly higher levels were noted in II-4 and RT3 cells (Figure 5A). The expression of *HOTTIP* and *MMP13* was increased in A5 compared to HaCaT cells and low in II-4 and RT3 cells (Figure 5A).

**Figure 5.**
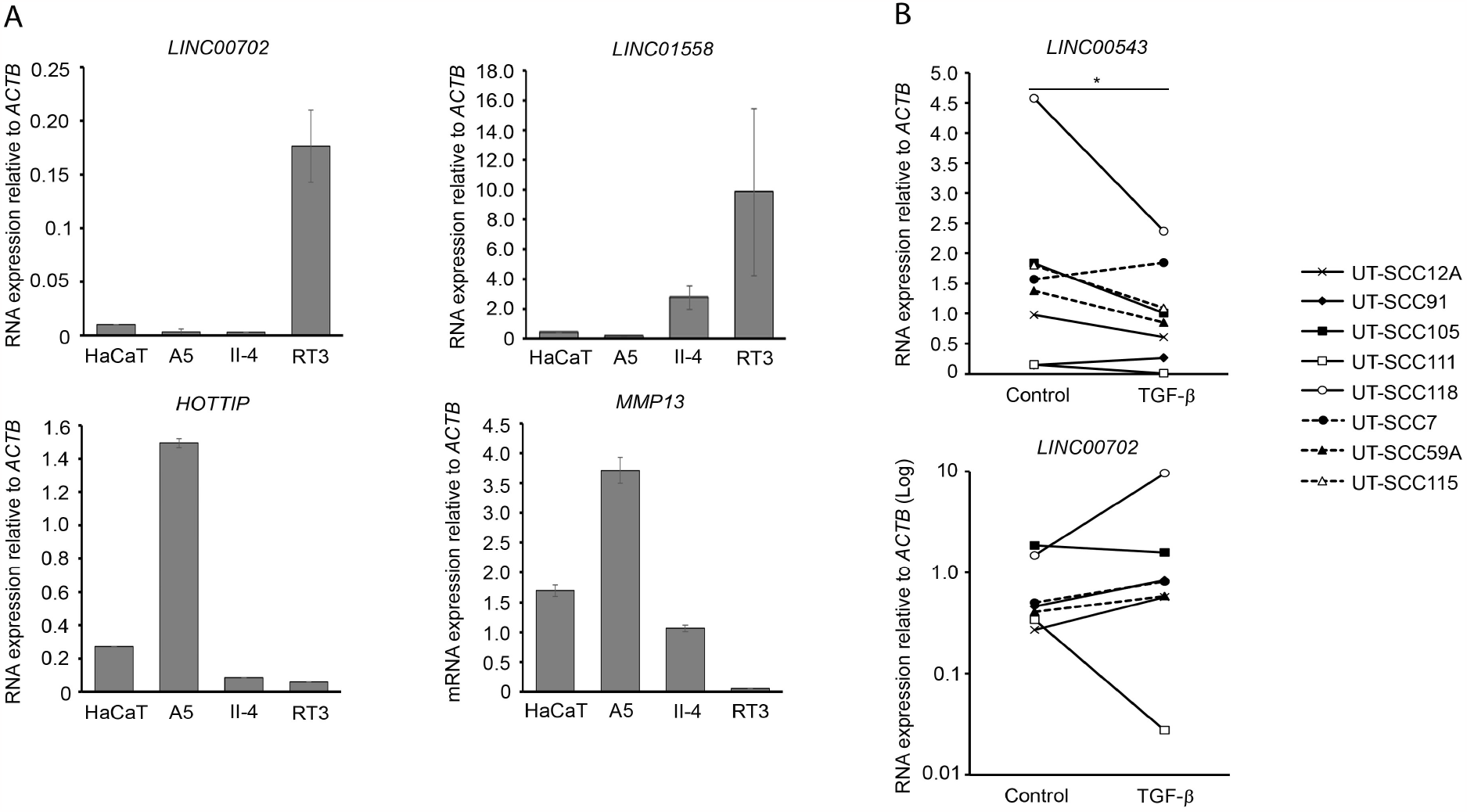
Regulation of lncRNA expression in Ha-***ras***-transformed HaCaT and cSCC cells. (**A**) *LINC00702* (n = 2-3), *LINC01558* (n = 3), *HOTTIP* (n = 2-3) and *MMP13* (n = 3) expression levels in HaCaT cells and tumorigenic Ha-ras-transformed HaCaT cell lines (A5, II-4, and RT3) were determined by qRT-PCR and corrected for the levels of β-actin (*ACTB*). (**B**) Cutaneous SCC cell lines (n = 7-8) were treated with transforming growth factor-β1 (TGF-β1; 5 ng/mL) for 24 hours. *LINC00543* and *LINC00702* levels were determined by qRT-PCR and corrected for the levels of β-actin (*ACTB*). *p<0.05, Mann–Whitney U-test.

### Regulation of lncRNAs *LINC00543* and *LINC00702* by TGF-β in cSCC cells

TGF-β signaling has been noted to function as a regulator of cSCC cell invasion (49-52). Thus the effect of TGF-β on the regulation of cell motility cluster lncRNAs was investigated. Cutaneous SCC cell lines were treated with TGF-β and the regulation on the lncRNAs was investigated by qRT-PCR. *LINC00702* was upregulated and *LINC00543* significantly downregulated in most SCC cell lines after treatment with TGF-β (Figure 5B). In contrast, *LINC01361* and *LINC01558* were downregulated and *LINC01460* was upregulated in cSCC cells after TGF-β treatment (Supplement Figure S5).

### Expression of lncRNAs in cSCC tumors *in vivo*

The overexpression of *LINC00543* was detected in cSCC tumor samples *in vivo* compared to normal skin by qRT-PCR (Figure 6A). Additionally, qRT-PCR analysis of the expression of protein coding genes revealed upregulation of *MMP13* in cSCC tumors *in vivo* (Figure 6B) (53).

**Figure 6.**
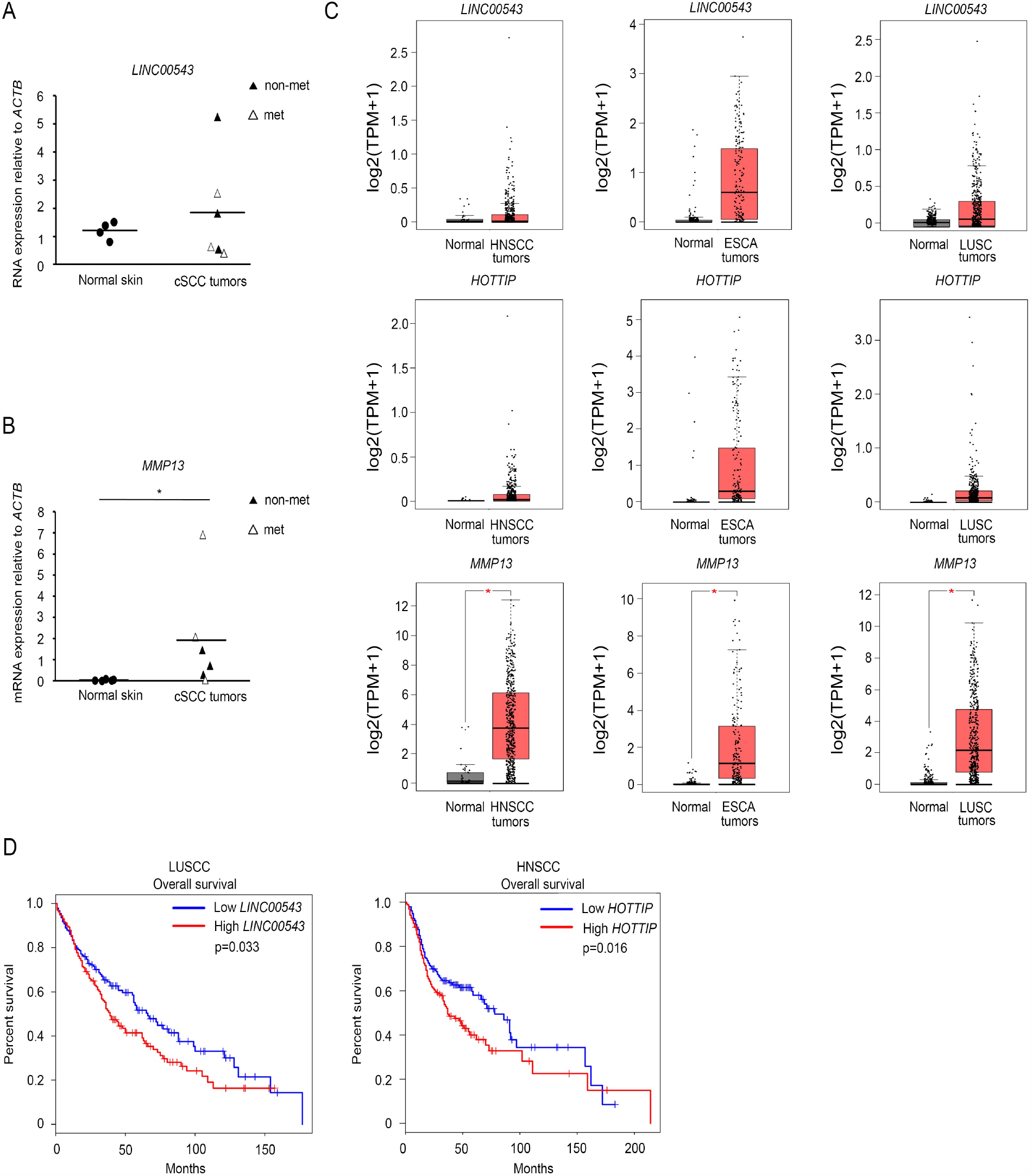
*LINC00543* and *HOTTIP* are associated with poor prognosis in SCC. (**A**,**B**) The expression of (**A**) *LINC00543* and (**B**) *MMP13* was determined in primary non-metastatic (non-met) and metastatic (met) cSCC tumor samples (n = 6) *in vivo* compared to normal skin (n = 4-5) by qRT-PCR. (**C**) The Cancer Genome Atlas (TCGA) data was utilized to investigate the expression of *LINC00543, HOTTIP* and *MMP13* in squamous cell carcinomas found in TCGA database (head and neck SCC (HNSCC), esophageal carcinoma (ESCA) and lung SCC (LUSC)). (**D**) The impact of *LINC00543* and *HOTTIP* on overall survival of patients with SCCs, was evaluated utilizing data available in the database of TCGA.

### Expression of *LINC00543* and *HOTTIP* in other SCCs is associated with poor prognosis

The Cancer Genome Atlas (TCGA) (41,42) data was utilized to investigate the expression of lncRNAs and *MMP13* in SCCs found in TCGA database (HNSCC, esophageal carcinoma (ESCA) and LUSC) (41,42). Cluster 1 related *LINC00543, HOTTIP* and *MMP13* were upregulated in SCCs investigated (Figure 6C). Next the impact of the expression of genes encoding lncRNAs on overall survival of patients with SCCs, was evaluated utilizing data available in the database of TCGA (41,42). Stronger expression of genes encoding *LINC00543* and *HOTTIP* were associated with poor prognosis, that is, with shorter overall survival in LUSCC and HNSCC, respectively (Figure 6D).

## DISCUSSION

The prognosis of the metastatic cSCC is poor, and there are no established biomarkers to predict metastatic risk nor specific therapeutic targets for advanced or metastatic cSCCs. Thus, characterization of cSCC progression at molecular level is important to be able to reveal dysregulated pathways and to identify key drivers of the disease. LncRNAs are under intensive investigation in cSCC but the role of lncRNAs in the progression of this cancer is still largely unknown (15). Recent studies have revealed several lncRNAs, such as *MALAT1, PVT1, LINC00319*, involved in the progression of cSCC (20,21,54,55). Additionally, our previous studies have revealed the role of lncRNAs *PICSAR, PRECSIT* and *SERLOC* in cSCC (16-19).

Here deep RNA-seq of patient derived cSCC cells and NHEKs was utilized to identify novel lincRNAs in cSCC progression. Among the top downregulated lncRNAs and mRNAs there were molecules previously linked to progression of cSCC. *LINC00520* was among the top downregulated lncRNAs and it has been shown to have inhibitory effect on cSCC progression (43). Among top upregulated lncRNAs noted *PRECSIT* (*LINC00346*) has previously been demonstrated to promote cSCC progression (18). A top protein coding gene *AIM2*, codes for inflammasome component AIM2, which has been shown to promote growth and invasion of cSCC (18,44). These observations confirm that the RNA-seq results are in accordance with previous analyses of cSCC cells.

To obtain additional insight into the progression of cSCC the deep RNA-seq data was utilized for the integration analysis of lncRNAs and protein coding genes to predict their potential functions in cSCC progression. Integration analysis is a common approach to predict the functions of lncRNAs in various biological processes (56). Previously a potential reference set of lncRNAs, mRNAs and circular RNAs in cSCC has been identified utilizing whole transcriptome sequencing (57). Additionally, microarray analysis of cSCC precursor actinic keratosis samples has revealed a lncRNA, which has a role in JAK-STAT3 signaling pathway in AK (58). However, to our knowledge no whole transcriptome sequencing based integration analysis of lncRNAs and protein coding genes has been prepared for genes regulated in cSCC samples earlier. The integration analysis revealed six clusters of genes two of which were upregulated. We further investigated one of the upregulated clusters, namely cluster 1. Bioinformatics analysis of enrichment of the genes revealed that the cluster 1 genes were significantly enriched in cell motility related terms and pathways. Thus the cluster 1 was named as cSCC motility cluster.

We focused on cSCC motility cluster related lncRNAs that was the second largest biotype of lncRNAs after antisense RNAs. Previous studies with other cancers have also noted that the two predominant categories of lncRNAs are antisense and lincRNAs (59). Among the top regulated lncRNAs in cluster 1 there were several novel transcripts and previously uncharacterized lincRNAs, but also some previously characterized cancer cell motility related lncRNAs, like *HOTTIP* (47) and *LINC00491* (48). *HOTTIP* has been shown to regulate invasion of prostate cancer cells (60). Additionally, *LINC00491* has been noted to regulate migration and invasion of colon adenocarcinoma cells (61). Also as previously known cSCC cell invasion related matrix metalloproteinase, MMP-13, and complement inhibitor serine protease FI were shown to be involved in this cluster of genes thus emphasizing the function of this cluster (46,62).

Myc and EZH2 were noted as the most frequent TFs in cell motility cluster. These findings are in accordance with previous studies showing that Myc can induce poor differentiation of cSCC and suggesting a role for it in the acquisition of a more aggressive tumor phenotype (63). EZH2 has been noted to promote progression of cSCC (64). Interestingly, EZH2 has also been shown to interact with Myc and coactivator (p300) via a cryptic transactivation domain (TAD), and this way induce gene activation and oncogenesis (65).

We investigated further the expression of selected lncRNAs and as a control one mRNA, *MMP13*, in patient derived cSCC and HNSCC cells. The upregulation of previously uncharacterized lncRNAs, such as *LINC01361, LINC01460* and *LINC01558* and already characterized *HOTTIP, LINC00543* and *LINC00702* was confirmed in cSCC cells compared to NHEKs by qRT-PCR. These lncRNAs except *LINC00702* were also upregulated in HNSCC cells. Additionally, the expression of *LINC00702, LINC01558* and *HOTTIP* in *in vitro* model of cSCC progression confirmed their upregulation during epidermal keratinocyte carcinogenesis. As mentioned above, *HOTTIP* has been shown to regulate cancer progression by promoting invasion of prostate cancer cells and *LINC00543* promotes metastasis of colorectal cancer (66). On the other hand, *LINC00702* has previously been shown to be downregulated in gastric and bladder cancer (67,68) but promote the pathogenesis of meningeoma (69).

TGF-β signaling pathway participates the invasion of SCC cells (49-51). Decrease of TGF-β signaling has been noted to be associated with increased invasion depth of cSCC (52). In this study we show, that *LINC00543* and *LINC00702* are regulated by TGF-β, and while TGF-β seems important to cell motility, our results support the idea that these lncRNAs belong to the cell motility cluster.

The expression of selected lncRNAs was also investigated in *in vivo* cSCC tumor samples by qRT-PCR. *LINC00543* was detected to be upregulated also in cSCC tumor samples *in vivo* compared to normal skin. In addition, qRT-PCR analysis revealed a significant upregulation of *MMP13* in cSCC compared to normal skin. Additionally TCGA data was utilized to investigate the expression of lncRNAs and *MMP13* in SCCs. *LINC00543, HOTTIP* and *MMP13* were demonstrated to be upregulated in HNSCC, esophageal carcinoma and LUSCC. Furthermore, stronger expression of *LINC00543* and *HOTTIP* were associated with poor prognosis, that is, with shorter overall survival in lung SCC and HNSCC, respectively. These results reveal that upregulation of *LINC00543* and *HOTTIP* may indicate poor prognosis of SCC patients. Further studies with larger cSCC sample set is required to confirm these results since cSCC expression data is not included in TCGA database.

In summary, we utilized the deep RNA-seq data for the integration analysis of lncRNAs and mRNAs to predict their potential functions in cSCC progression. Based on these results a set of lncRNAs was recognized to be involved in cSCC cell motility cluster. The lncRNAs investigated were shown to be upregulated in patient derived cSCC and HNSCC cell lines. Furthermore, *LINC00543* and *HOTTIP* were also overexpressed in tumor samples *in vivo* and stronger expression of *LINC00543* and *HOTTIP* was associated with poor prognosis of patient with lung SCC or HNSCC. The results of this study implicate cell motility cluster related lncRNAs *HOTTIP, LINC00543, LINC00702, LINC01361, LINC01460* and *LINC01558* as potential biomarkers and putative therapeutic target in advanced and metastatic cSCC.

## Supporting information

Nissinen_supplemental

lncRNA: long non-coding RNA;
linc: long intergenic non-coding;
cSCC: cutaneous squamous cell carcinoma;
qRT-PCR: quantitative real-time reverse transcriptase-PCR;
MMP: matrix metalloproteinase

## ACKNOWLEDGMENTS

We thank Johanna Markola for expert technical assistance; Dr. Reidar Grénman for cSCC and HNSCC cell lines and tumor samples; Dr. Norbert E. Fusenig (The German Cancer Research Center, Heidelberg, Germany) for HaCaT cells; GeneVia Technologies (Tampere, Finland) for bioinformatics analysis.

## DATA AVAILABILITY

The RNA-seq data of cSCC cell lines and NHEKs will be available in the GEO database.

## REFERENCES

1. Adnane, S., Marino, A., Leucci, E. (2022) LncRNAs in human cancers: signal from noise. Trends Cell. Biol., 32, 565–573.

2. Derrien, T., Johnson, R., Bussotti, G., Tanzer, A., Djebali, S., Tilgner, H., Guernec, G., Martin, D., Merkel, A., Knowles, D.G. et al. (2012) The GENCODE v7 catalog of human long noncoding RNAs: analysis of their gene structure, evolution, and expression. Genome Res., 22, 1775–1789.

3. Marchese, F.P., Raimondi, I., Huarte, M. (2017) The multidimensional mechanisms of long noncoding RNA function. Genome Biol., 18, 206.

4. Nehal, K.S. and Bichakjian, C.K. (2018) Update on keratinocyte carcinomas. N. Engl. J. Med., 379, 363–374.

5. Burton, K.A., Ashack, K.A., Khachemoune, A. (2016) Cutaneous squamous cell carcinoma: a review of high-risk and metastatic disease. Am. J. Clin. Dermatol., Oct;17, 491–508.

6. Knuutila, J.S., Riihilä, P., Kurki, S., Nissinen, L., Kähäri, V.M. (2020) Risk factors and prognosis for metastatic cutaneous squamous cell carcinoma: a cohort study. Acta Derm. Venereol., 100, adv00266.

7. Winge, M.C.G., Kellman, L.N., Guo, K., Tang, J.Y., Swetter, S.M., Aasi, S.Z., Sarin, K.Y., Chang, A.L.S., Khavari, P.A. (2023) Advances in cutaneous squamous cell carcinoma. Nat. Rev. Cancer, 23, 430–449.

8. Cho, R.J., Alexandrov, L.B., den Breems, N.Y., Atanasova, V.S., Farshchian, M., Purdom, E., Nguyen, T.N., Coarfa, C., Rajapakshe, K., Prisco, M. et al. (2018) APOBEC mutation drives early-onset squamous cell carcinomas in recessive dystrophic epidermolysis bullosa. Sci. Transl. Med., 10, eaas9668.

9. Piipponen, M., Riihilä, P., Nissinen, L., Kähäri, V.M. (2021) The Role of p53 in progression of cutaneous squamous cell carcinoma. Cancers (Basel), 13, 4507.

10. Hedberg, M.L., Berry, C.T., Moshiri, A.S., Xiang, Y., Yeh, C.J., Attilasoy, C., Capell, B.C., Seykora, J.T. (2022) Molecular Mechanisms of Cutaneous Squamous Cell Carcinoma. Int. J. Mol. Sci., 23, 3478.

11. Pickering, C.R., Zhou, J.H., Lee, J.J., Drummond, J.A., Peng, S.A., Saade, R.E., Tsai, K.Y., Curry, J.L., Tetzlaff, M.T., Lai, S.Y. et al. (2014) Mutational landscape of aggressive cutaneous squamous cell carcinoma. Clin. Cancer Res., 20, 6582–6592.

12. South, A.P., Purdie, K.J., Watt, S.A., Haldenby, S., den Breems, N., Dimon, M., Arron, S.T., Kluk, M.J., Aster, J.C., McHugh, A. et al. (2014) NOTCH1 mutations occur early during cutaneous squamous cell carcinogenesis. J. Invest. Dermatol., 134, 2630–2638.

13. Nissinen, L., Farshchian, M., Riihilä, P., Kähäri, V.M. (2016) New perspectives on role of tumor microenvironment in progression of cutaneous squamous cell carcinoma. Cell Tissue Res., 365, 691–702.

14. Riihilä, P., Nissinen, L., Kähäri, V.M. (2021) Matrix metalloproteinases in keratinocyte carcinomas. Exp. Dermatol., 30, 50–61.

15. Piipponen, M., Nissinen, L., Kähäri, V.M. (2020b) Long non-coding RNAs in cutaneous biology and keratinocyte carcinomas. Cell. Mol. Life Sci., 77, 4601–4614.

16. Piipponen, M., Nissinen, L., Farshchian, M., Riihilä, P., Kivisaari, A., Kallajoki, M., Peltonen, J., Peltonen, S., Kähäri, V.M. (2016) Long noncoding RNA PICSAR promotes growth of cutaneous squamous cell carcinoma by regulating ERK1/2 Activity. J. Invest. Dermatol., 136, 1701–1710.

17. Piipponen, M., Heino, J., Kähäri, V.M., Nissinen, L. (2018) Long non-coding RNA PICSAR decreases adhesion and promotes migration of squamous carcinoma cells by downregulating α2β1 and α5β1 integrin expression. Biol. Open, 7(11):bio037044.

18. Piipponen, M., Nissinen, L., Riihilä, P., Farshchian, M., Kallajoki, M., Peltonen, J., Peltonen, S., Kähäri, V.M. (2020a) p53-regulated long noncoding RNA PRECSIT Promotes progression of cutaneous squamous cell carcinoma via STAT3 Signaling. Am. J. Pathol., 190, 503–517.

19. Piipponen, M., Riihilä, P., Knuutila, J.S., Kallajoki, M., Kähäri, V.M., Nissinen, L. (2022) Super Enhancer-Regulated LINC00094 (SERLOC) Upregulates the expression of MMP-1 and MMP-13 and promotes invasion of cutaneous squamous cell carcinoma. Cancers (Basel), 14, 3980.

20. Li, C., Sun, C., Mahapatra, K.D., Riihilä, P., Knuutila, J., Nissinen, L., Lapins, J., Kähäri, V.M., Homey, B., Sonkoly, E., Pivarcsi, A. (2023) Long non-coding RNA PVT1 is overexpressed in cutaneous squamous cell carcinoma and exon 2 is critical for its oncogenicity. Br. J. Dermatol., Nov 1:ljad419.

21. Li, R., Huang, D., Ju, M., Chen, H.Y., Luan, C., Zhang, J.A., Chen, K. (2023) The long non-coding RNA PVT1 promotes tumorigenesis of cutaneous squamous cell carcinoma via interaction with 4EBP1. Cell Death Discov., 9, 101.

22. Farshchian, M., Nissinen, L., Grénman, R., Kähäri, V.M. (2017) Dasatinib promotes apoptosis of cutaneous squamous carcinoma cells by regulating activation of ERK1/2. Exp. Dermatol., 26, 89–92.

23. Johansson, N., Airola, K., Grénman, R., Kariniemi, A.L., Saarialho-Kere, U., Kähäri, V.M. (1997) Expression of collagenase-3 (matrix metalloproteinase-13) in squamous cell carcinomas of the head and neck. Am. J. Pathol., 151, 499–508.

24. Stokes, A., Joutsa, J., Ala-Aho, R., Pitchers, M., Pennington, C.J., Martin, C., Premachandra, D.J., Okada, Y., Peltonen, J., Grénman, R. et al. (2010) Expression profiles and clinical correlations of degradome components in the tumor microenvironment of head and neck squamous cell carcinoma. Clin. Cancer Res., 16, 2022–2035.

25. Mueller, M.M., Peter, W., Mappes, M., Huelsen, A., Steinbauer, H., Boukamp, P., Vaccariello, M., Garlick, J., Fusenig, N.E. (2001) Tumor progression of skin carcinoma cells in vivo promoted by clonal selection, mutagenesis, and autocrine growth regulation by granulocyte colony-stimulating factor and granulocyte-macrophage colony-stimulating factor. Am. J. Pathol., 159, 1567–1579.

26. Dobin, A., Davis, C.A., Schlesinger, F., Drenkow, J., Zaleski, C., Jha, S., Batut, P., Chaisson, M., Gingeras, TR. (2013) STAR: ultrafast universal RNA-seq aligner. Bioinformatics, 29, 15–21.

27. R Core Team. (2013) R: A language and environment for statistical computing. R Foundation for Statistical Computing, Vienna, Austria. URL: http://www.R-project.org/

28. Benjamini, Y. and Hochberg, Y. (1995) Controlling the false discovery rate: a practical and powerful approach to multiple testing. Journal of the Royal Statistical Society B, 57, 289–300.

29. Durinck, S., Spellman, P.T., Birney, E., Huber, W. (2009) Mapping identifiers for the integration of genomic datasets with the R/Bioconductor package biomaRt. Nat. Protoc., 4, 1184–1191.

30. Wickham, H. (2016) ggplot2: Elegant Graphics for Data Analysis. New York: Springer-Verlag

31. Kolde, R. (2018) pheatmap: Pretty Heatmaps. R package version 1.0.10. URL: https://CRAN.R-project.org/package=pheatmap

32. Ashburner, M., Ball, C.A., Blake, J.A., Botstein, D., Butler, H., Cherry, J.M., Davis, A.P., Dolinski, K., Dwight, S.S., Eppig, J.T. et al. (2000) Gene ontology: tool for the unification of biology. The Gene Ontology Consortium. Nat. Genet., 25, 25–29.

33. The Gene Ontology Consortium. (2017) Expansion of the Gene Ontology knowledgebase and resources. Nucleic Acids Res., 45(D1), D331–D338.

34. Yu, G., Wang, L.G., Han, Y., He, Q.Y. (2012) clusterProfiler: an R package for comparing biological themes among gene clusters. OMICS, 16, 284–287.

35. Krämer, A., Green, J., Pollard, J. Jr., Tugendreich, S. (2014) Causal analysis approaches in Ingenuity Pathway Analysis. Bioinformatics, 30, 523–530.

36. Huang da, W., Sherman, B.T., Lempicki, R.A. (2009a) Systematic and integrative analysis of large gene lists using DAVID bioinformatics resources. Nat. Protoc., 4, 44–57.

37. Huang da, W., Sherman, B.T., Lempicki, R.A. (2009b) Bioinformatics enrichment tools: paths toward the comprehensive functional analysis of large gene lists. Nucleic Acids Res., 37, 1–13.

38. Jiang, C., Xuan, Z., Zhao, F., Zhang, M.Q. (2007) TRED: a transcriptional regulatory element database, new entries and other development. Nucleic Acids Res., 35(Database issue), D137–140.

39. Liu, Z.P., Wu, C., Miao, H., Wu, H. (2015) RegNetwork: an integrated database of transcriptional and post-transcriptional regulatory networks in human and mouse. Database (Oxford), Sep 30;bav095.

40. Lachmann, A., Xu, H., Krishnan, J., Berger, S.I., Mazloom, A.R., Ma’ayan, A. (2010) ChEA: transcription factor regulation inferred from integrating genome-wide ChIP-X experiments. Bioinformatics, 26, 2438–2444.

41. Uhlen, M., Zhang, C., Lee, S., Sjöstedt, E., Fagerberg, L., Bidkhori, G., Benfeitas, R., Arif, M., Liu, Z., Edfors, F. et al. (2017) A pathology atlas of the human cancer transcriptome. Science, 357, (6352), eaan2507.

42. Cancer Genome Atlas Research Network, Weinstein, J.N., Collisson, E.A., Mills, G.B., Shaw, K.R., Ozenberger, B.A., Ellrott, K., Shmulevich, I., Sander, C., Stuart, J.M. (2013) The Cancer Genome Atlas pan-cancer analysis project. Nat. Genet., 45, 1113–1120.

43. Mei, X.L. and Zhong, S. (2019) Long noncoding RNA LINC00520 prevents the progression of cutaneous squamous cell carcinoma through the inactivation of the PI3K/Akt signaling pathway by downregulating EGFR. Chin. Med. J., 132, 454–465.

44. Farshchian, M., Nissinen, L., Siljamäki, E., Riihilä, P., Piipponen, M., Kivisaari, A., Kallajoki, M., Grénman, R., Peltonen, J., Peltonen, S. et al. (2017) Tumor cell-specific AIM2 regulates growth and invasion of cutaneous squamous cell carcinoma. Oncotarget, 8, 45825–45836.

45. Szklarczyk, D., Franceschini, A., Kuhn, M., Simonovic, M., Roth, A., Minguez, P., Doerks, T., Stark, M., Muller, J., Bork, P. et al. (2011) The STRING database in 2011: functional interaction networks of proteins, globally integrated and scored. Nucleic Acids Res., 39(Database issue), D561–568.

46. Ala-aho, R., Ahonen, M., George, S.J., Heikkilä, J., Grénman, R., Kallajoki, M., Kähäri, V.M. (2004) Targeted inhibition of human collagenase-3 (MMP-13) expression inhibits squamous cell carcinoma growth in vivo. Oncogene, 23, 5111–5123.

47. Feng, H., Zhao, F., Luo, J., Xu, S., Liang, Z., Xu, W., Bao, Y., Qin, G. (2023) Long non-coding RNA HOTTIP exerts an oncogenic function by regulating HOXA13 in nasopharyngeal carcinoma. Mol. Biol. Rep., 50, 6807–6818.

48. Wan, H., Lin, T., Shan, M., Lu, J., Guo, Z. (2022) LINC00491 Facilitates Tumor Progression of lung adenocarcinoma via Wnt/β-catenin-signaling pathway by regulating MTSS1 Ubiquitination. Cells, 11, 3737.

49. Leivonen, S.K., Ala-Aho, R., Koli, K., Grénman, R., Peltonen, J., Kähäri, V.M. (2006) Activation of Smad signaling enhances collagenase-3 (MMP-13) expression and invasion of head and neck squamous carcinoma cells. Oncogene, 25, 2588–2600.

50. Siljamäki, E., Rappu, P., Riihilä, P., Nissinen, L., Kähäri, V.M., Heino, J. (2020) H-Ras activation and fibroblast-induced TGF-β signaling promote laminin-332 accumulation and invasion in cutaneous squamous cell carcinoma. Matrix Biol., 87, 26–47.

51. Siljamäki, E., Riihilä, P., Suwal, U., Nissinen, L., Rappu, P., Kallajoki, M., Kähäri, V.M., Heino, J. (2023) Inhibition of TGF-β signaling, invasion, and growth of cutaneous squamous cell carcinoma by PLX8394. Oncogene, 42, 3633–3647.

52. Rose, A.M., Spender, L.C., Stephen, C., Mitchell, A., Rickaby, W., Bray, S., Evans, A.T., Dayal, J., Purdie, K.J., Harwood, C.A. et al. (2018) Reduced SMAD2/3 activation independently predicts increased depth of human cutaneous squamous cell carcinoma. Oncotarget, 9, 14552–14566.

53. Viiklepp, K., Nissinen, L., Ojalill, M., Riihilä, P., Kallajoki, M., Meri, S., Heino, J., Kähäri, V.M. (2022) C1r upregulates production of matrix metalloproteinase-13 and promotes invasion of cutaneous squamous cell carcinoma. J. Invest. Dermatol., 142, 1478–1488.

54. Zhang, C., Wang, J., Guo, L., Peng, M. (2021) Long non-coding RNA MALAT1 regulates cell proliferation, invasion and apoptosis by modulating the Wnt signaling pathway in squamous cell carcinoma. Am. J. Transl. Res., 13, 9233–9240.

55. Li, F., Liao, J., Duan, X., He, Y., Liao, Y. (2018) Upregulation of LINC00319 indicates a poor prognosis and promotes cell proliferation and invasion in cutaneous squamous cell carcinoma. J. Cell. Biochem., 119, :10393–10405.

56. Chen, H., Zhang, K., Lu, J., Wu, G., Yang, H., Chen, K. (2017) Comprehensive analysis of mRNA-lncRNA co-expression profile revealing crucial role of imprinted gene cluster DLK1-MEG3 in chordoma. Oncotarget, 8, 112623–112635.

57. Das Mahapatra, K., Pasquali, L., Søndergaard, J.N., Lapins, J., Nemeth, I.B., Baltás, E., Kemény, L., Homey, B., Moldovan, L.I., Kjems, J. et al. (2020) A comprehensive analysis of coding and non-coding transcriptomic changes in cutaneous squamous cell carcinoma. Sci. Rep., 10, 3637.

58. Luan, C., Jin, S., Hu, Y., Zhou, X., Liu, L., Li, R., Ju, M., Huang, D., Chen, K. (2022) Whole-genome identification and construction of the lncRNA-mRNA co-expression network in patients with actinic keratosis. Transl. Cancer Res., 11, 4070–4078.

59. Siena, Á.D.D., Plaça, J.R., Araújo, L.F., de Barros, I.I., Peronni, K., Molfetta, G., de Biagi, C.A.O. Jr., Espreafico, E.M., Sousa, J.F., Silva, W.A. Jr. (2019) Whole transcriptome analysis reveals correlation of long noncoding RNA ZEB1-AS1 with invasive profile in melanoma. Sci. Rep., 9, 11350.

60. Qian, Y., Wang, J., Wang, B., Wang, W., Li, P., Zhao, Z., Jiang, Y., Ren, H., Huang, D., Yang, Y. et al. (2023) Systematic fine-mapping and functional studies of prostate cancer risk variants. iScience, 26, 106497.

61. Wan, J., Deng, D., Wang, X., Wang, X., Jiang, S., Cui, R. (2019) LINC00491 as a new molecular marker can promote the proliferation, migration and invasion of colon adenocarcinoma cells. Onco Targets Ther., 12, 6471–6480.

62. Rahmati Nezhad, P., Riihilä, P., Piipponen, M., Kallajoki, M., Meri, S., Nissinen, L., Kähäri, V.M. (2021) Complement factor I upregulates expression of matrix metalloproteinase-13 and -2 and promotes invasion of cutaneous squamous carcinoma cells. Exp. Dermatol., 30, 1631–1641.

63. Toll, A., Salgado, R., Yébenes, M., Martín-Ezquerra, G., Gilaberte, M., Baró, T., Solé, F., Alameda, F., Espinet, B., Pujol, R.M. (2009) MYC gene numerical aberrations in actinic keratosis and cutaneous squamous cell carcinoma. Br. J. Dermatol., 161, 1112–1118.

64. Xie, Q., Wang, H., Heilman, E.R., Walsh, M.G., Haseeb, M.A., Gupta, R. (2014) Increased expression of enhancer of Zeste Homolog 2 (EZH2) differentiates squamous cell carcinoma from normal skin and actinic keratosis. Eur. J. Dermatol., 24, 41–45.

65. Wang, J., Yu, X., Gong, W., Liu, X., Park, K.S., Ma, A., Tsai, Y.H., Shen, Y., Onikubo, T., Pi, W.C. et al. (2022) EZH2 noncanonically binds cMyc and p300 through a cryptic transactivation domain to mediate gene activation and promote oncogenesis. Nat. Cell Biol., 24, 384–399.

66. Zheng, J., Dou, R., Zhang, X., Zhong, B., Fang, C., Xu, Q., Di, Z., Huang, S., Lin, Z., Song, J. et al. (2023) LINC00543 promotes colorectal cancer metastasis by driving EMT and inducing the M2 polarization of tumor associated macrophages. J. Transl. Med., 21, 153.

67. Ma, M., Li, J., Zeng, Z., Zheng, Z., Kang, W. (2023) Integrated analysis from multicentre studies identities m7G-related lncRNA-derived molecular subtypes and risk stratification systems for gastric cancer. Front. Immunol., 14, 1096488.

68. Pan, W., Han, J., Wei, N., Wu, H., Wang, Y., Sun, J. (2022) LINC00702-mediated DUSP1 transcription in the prevention of bladder cancer progression: Implications in cancer cell proliferation and tumor inflammatory microenvironment. Genomics, 114, 110428.

69. Ghafouri-Fard, S., Abak, A., Hussen, B.M., Taheri, M., Sharifi, G. (2021) The emerging role of non-coding rnas in pituitary gland tumors and meningioma. Cancers (Basel), 13, 5987.

