## Supplementary material for "Clustering of RNA co-expression network identifies novel long non-coding RNA biomarkers in squamous cell carcinoma": Nissinen_supplemental

Supplement figures (S1-S5) and tables (S1-S6)

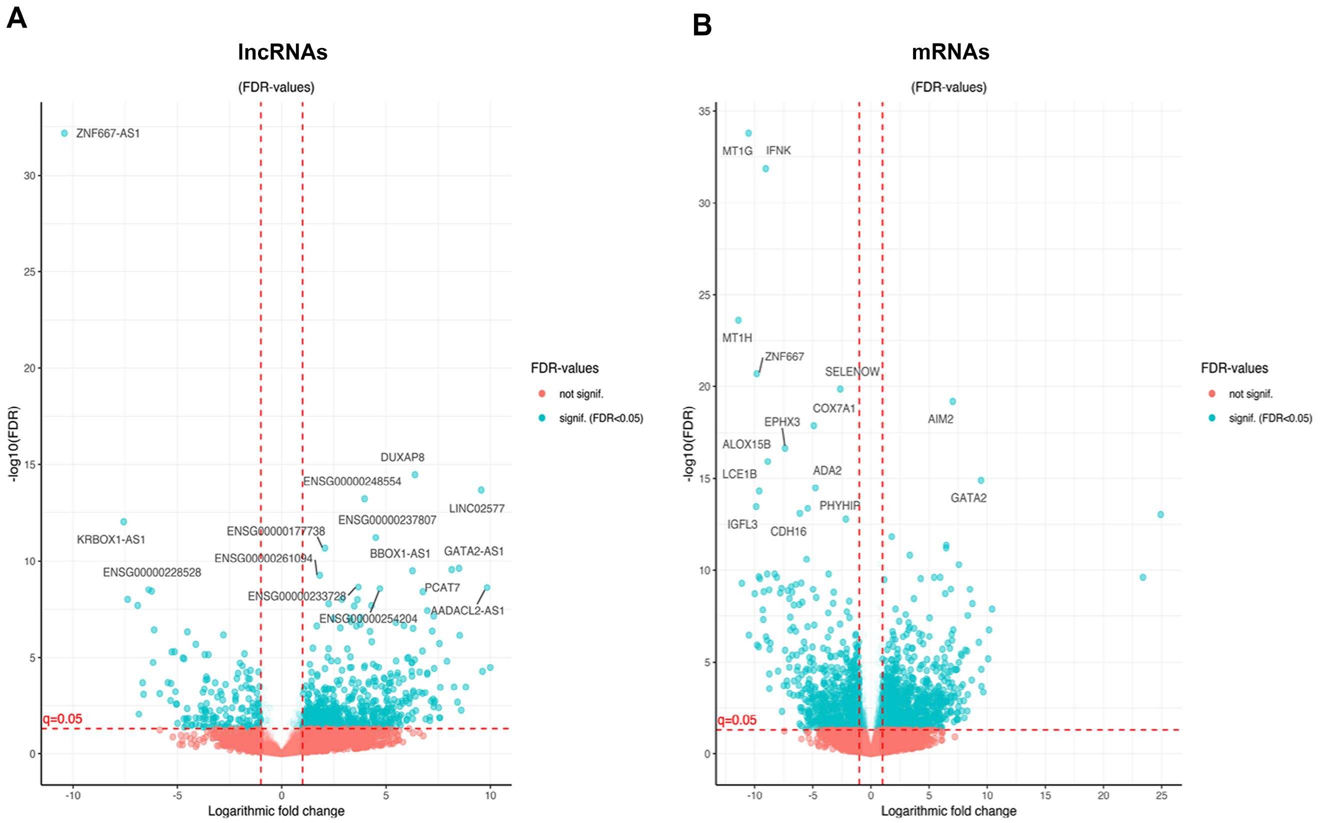


**Supplement figure S1.** The volcano plots show the distribution of differentially expressed long non-coding RNAs (lncRNA) (**A**) and protein coding genes (mRNA) (**B**) identified in cSCC cells (n=8) compared to normal epidermal keratinocytes (n=4) by RNA-seq.

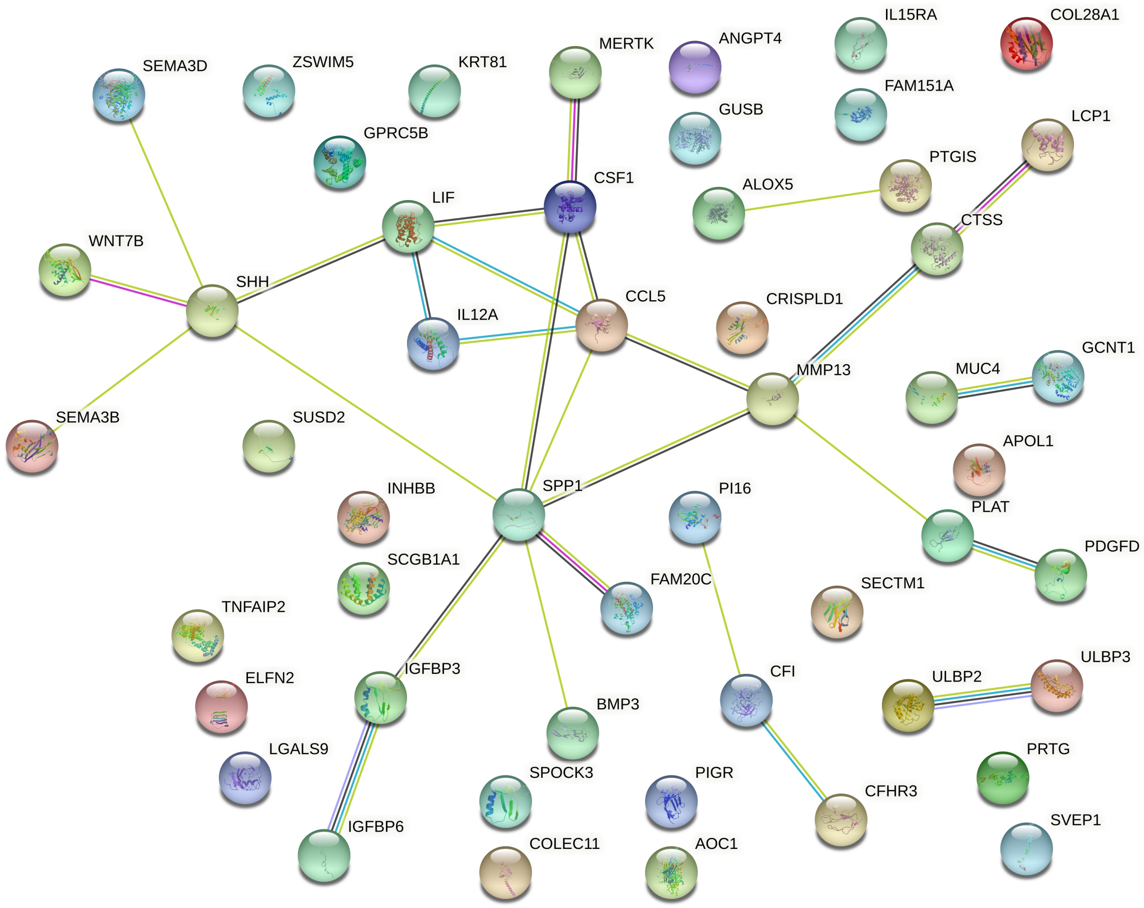


**Supplement figure S2.** A network of the cluster 1 genes related to term *extracellular space* was formed by taking the protein–protein functional interaction data from STRING database.

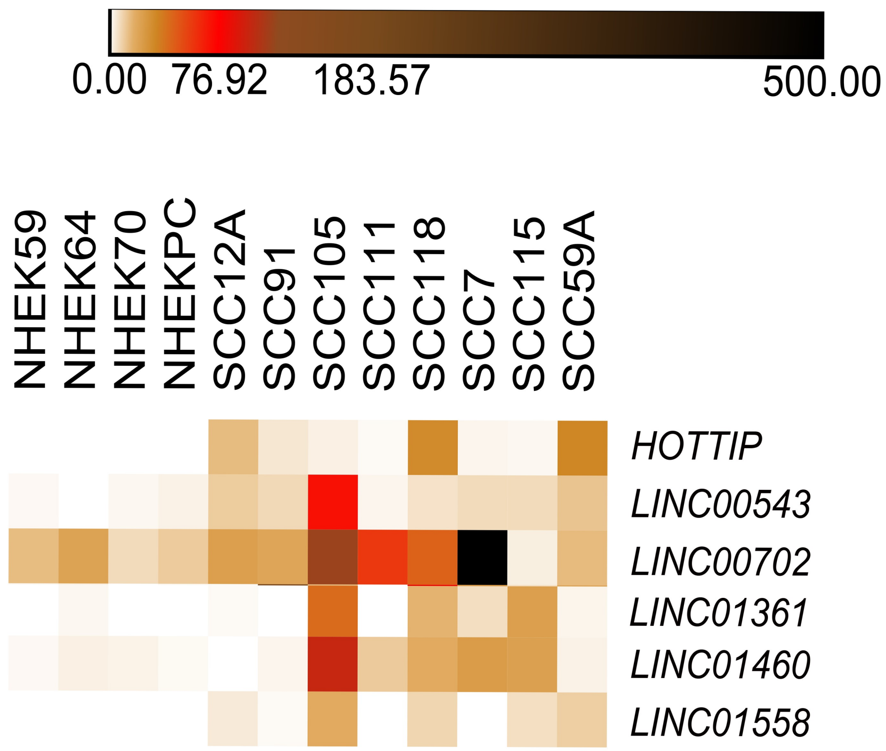


**Supplement figure S3.** The expression of selected lncRNA genes belonging to cluster 1 in cSCC cells and normal human epidermal keratinocytes (NHEK) analyzed by RNA-seq.

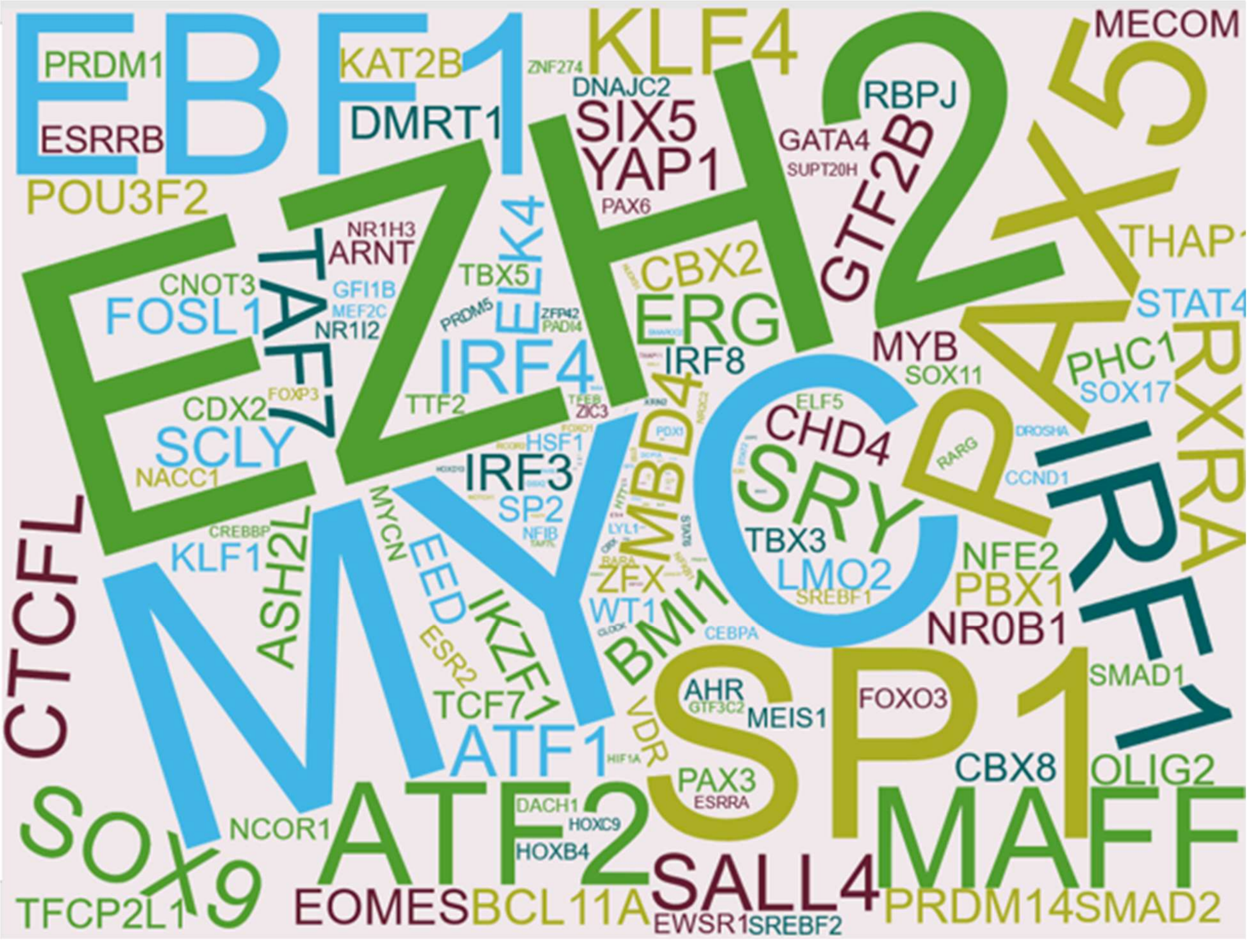


**Supplement figure S4.** Transcription factors (TF) are visualized by word cloud indicating the most frequent TFs regulating the expression of cluster 1 genes.

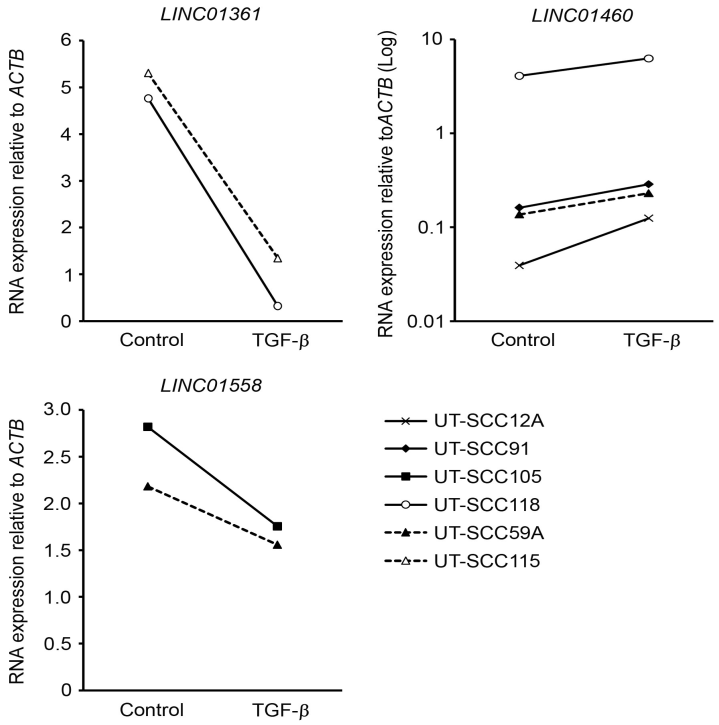


**Supplement figure S5.** Cutaneous SCC cell lines (n=2-4) were treated with transforming growth factor-β1 (TGF-β1; 5 ng $ml^{-1}$) for 24 hours. *LINC01361*, *LINC01460* and *LINC01558* levels were determined by qRT-PCR and corrected for the levels of β-actin (*ACTB*).

**Supplement table S1.** Significantly regulated lncRNAs in cSCC cells compared to NHEKs. Differentially expressed genes between cSCC and NHEKs were considered as significantly differentially expressed, if adjusted p-value (FDR) was less than 0.05 and absolute log2 fold change of expression level over 1.

| **Ensembl.ID** | **HGNC.symbol** | **Log2.foldchange** | **Adjusted.p.value** |
|---|---|---|---|
| *ENSG00000229967* | *MAGEA4-AS1* | 9,99 | 3,30675E-05 |
| *ENSG00000242908* | *AADACL2-AS1* | 9,84 | 2,40124E-09 |
| *ENSG00000250920* | | 9,62 | 5,33374E-05 |
| *ENSG00000228742* | *LINC02577* | 9,56 | 2,15558E-14 |
| *ENSG00000224516* | | 8,82 | 0,00034106 |
| *ENSG00000259672* | | 8,60 | 0,005378909 |
| *ENSG00000270372* | | 8,53 | 7,28526E-07 |
| *ENSG00000244300* | *GATA2-AS1* | 8,50 | 2,42271E-10 |
| *ENSG00000233515* | *LINC01518* | 8,42 | 0,002094643 |
| *ENSG00000268531* | | 8,26 | 0,00033776 |
| *ENSG00000231806* | *PCAT7* | 8,15 | 2,83068E-10 |
| *ENSG00000223829* | | 7,92 | 1,54737E-05 |

| | | | |
|---|---|---|---|
| *ENSG00000257636* | *G2E3-AS1* | 7,60 | 0,000353979 |
| *ENSG00000251350* | *LINC02475* | 7,58 | 0,013918649 |
| *ENSG00000268388* | *FENDRR* | 7,56 | 0,013262098 |
| *ENSG00000272763* | | 7,55 | 1,99041E-06 |
| *ENSG00000250546* | | 7,52 | 0,003965154 |
| *ENSG00000250564* | | 7,34 | 0,001824812 |
| *ENSG00000242207* | *HOXB-AS4* | 7,30 | 3,76278E-05 |
| *ENSG00000248370* | *LINC02434* | 7,30 | 0,00222641 |
| *ENSG00000249001* | | 7,29 | 0,00360993 |
| *ENSG00000267123* | *LINC02081* | 7,28 | 7,4267E-08 |
| *ENSG00000229970* | | 7,26 | 0,001461161 |
| *ENSG00000272872* | | 7,21 | 4,38469E-07 |
| *ENSG00000261706* | *LINC00165* | 7,16 | 0,001837674 |
| *ENSG00000251577* | | 7,10 | 0,012542655 |
| *ENSG00000258654* | | 7,10 | 0,019119191 |
| *ENSG00000258711* | | 6,98 | 3,79462E-08 |
| *ENSG00000233251* | | 6,98 | 6,36382E-05 |
| *ENSG00000244649* | *LINC02086* | 6,97 | 7,424E-05 |
| *ENSG00000264464* | | 6,90 | 0,016194043 |
| *ENSG00000267147* | *LINC01842* | 6,88 | 0,000666458 |
| *ENSG00000265933* | *LINC00668* | 6,76 | 4,0112E-09 |
| *ENSG00000250682* | *LINC00491* | 6,75 | 0,000746548 |
| *ENSG00000225684* | *FAM225B* | 6,74 | 4,69031E-06 |
| *ENSG00000257114* | *LINC02450* | 6,61 | 5,75626E-05 |
| *ENSG00000255571* | *MIR9-3HG* | 6,60 | 0,000621335 |
| *ENSG00000261730* | | 6,43 | 0,005647023 |
| *ENSG00000251169* | *LINC01843* | 6,41 | 0,001029448 |
| *ENSG00000206195* | *DUXAP8* | 6,38 | 3,36524E-15 |
| *ENSG00000244675* | | 6,37 | 1,29013E-05 |
| *ENSG00000204876* | | 6,31 | 0,000531918 |
| *ENSG00000260757* | | 6,30 | 0,000647528 |
| *ENSG00000256128* | *LINC00944* | 6,29 | 3,1886E-07 |
| *ENSG00000254287* | | 6,28 | 0,015877082 |
| *ENSG00000254560* | *BBOX1-AS1* | 6,27 | 3,23762E-10 |
| *ENSG00000253931* | | 6,26 | 0,007296581 |
| *ENSG00000282950* | | 6,24 | 9,19644E-05 |
| *ENSG00000245526* | *LINC00461* | 6,24 | 1,81092E-05 |
| *ENSG00000236347* | | 6,23 | 1,95266E-05 |
| *ENSG00000204792* | *LINC01291* | 6,23 | 5,2973E-05 |
| *ENSG00000232259* | | 6,15 | 0,001355437 |
| *ENSG00000277128* | | 6,12 | 0,001138149 |
| *ENSG00000232325* | | 6,10 | 0,00234096 |
| *ENSG00000248693* | *LINC02100* | 6,08 | 9,98806E-06 |
| *ENSG00000276707* | | 6,07 | 0,010327664 |
| *ENSG00000232934* | | 6,02 | 1,09373E-05 |

| | | | |
|---|---|---|---|
| *ENSG00000267327* | | 6,00 | 0,016598168 |
| *ENSG00000267284* | | 5,97 | 0,019187488 |
| *ENSG00000237181* | | 5,95 | 0,001248896 |
| *ENSG00000260944* | *FOXC2-AS1* | 5,95 | 0,00181694 |
| *ENSG00000260877* | | 5,95 | 0,000721283 |
| *ENSG00000263745* | | 5,86 | 2,26484E-07 |
| *ENSG00000278936* | | 5,83 | 0,00012844 |
| *ENSG00000243766* | *HOTTIP* | 5,80 | 0,002364294 |
| *ENSG00000277268* | *LHX1-DT* | 5,77 | 0,001829216 |
| *ENSG00000260976* | *LINC01633* | 5,74 | 0,00954983 |
| *ENSG00000237862* | | 5,70 | 0,032707799 |
| *ENSG00000238160* | | 5,66 | 0,003433127 |
| *ENSG00000272154* | | 5,66 | 0,011220294 |
| *ENSG00000249916* | | 5,62 | 0,023905976 |
| *ENSG00000234147* | | 5,61 | 0,000201417 |
| *ENSG00000213468* | *FIRRE* | 5,61 | 0,016506642 |
| *ENSG00000249894* | | 5,58 | 0,002029575 |
| *ENSG00000231346* | *LINC01160* | 5,56 | 6,37996E-05 |
| *ENSG00000260604* | | 5,54 | 8,87514E-05 |
| *ENSG00000271314* | | 5,52 | 0,008947344 |
| *ENSG00000242611* | | 5,49 | 0,01383483 |
| *ENSG00000259240* | *MIR4713HG* | 5,48 | 1,48009E-05 |
| *ENSG00000231560* | *CLEC12A-AS1* | 5,48 | 0,037752218 |
| *ENSG00000255874* | *LINC00346* | 5,47 | 1,59518E-07 |
| *ENSG00000222033* | *LINC01124* | 5,38 | 0,001559979 |
| *ENSG00000254338* | *MAFA-AS1* | 5,36 | 0,017008587 |
| *ENSG00000255446* | | 5,35 | 0,004120155 |
| *ENSG00000280026* | | 5,35 | 0,000445061 |
| *ENSG00000249772* | | 5,35 | 0,009346883 |
| *ENSG00000226053* | *LINC01776* | 5,34 | 0,014904445 |
| *ENSG00000229081* | *LINC01165* | 5,30 | 0,006567858 |
| *ENSG00000258661* | | 5,29 | 0,002592234 |
| *ENSG00000246095* | *LINC01096* | 5,26 | 0,004824501 |
| *ENSG00000205634* | *LINC00898* | 5,25 | 0,000100642 |
| *ENSG00000254349* | *MIR2052HG* | 5,25 | 0,023494137 |
| *ENSG00000251003* | *ZFPM2-AS1* | 5,24 | 0,000201045 |
| *ENSG00000259180* | | 5,24 | 0,026081279 |
| *ENSG00000247402* | | 5,21 | 0,016226471 |
| *ENSG00000189238* | *LINC00943* | 5,21 | 6,96908E-05 |
| *ENSG00000258754* | *LINC01579* | 5,21 | 0,042918871 |
| *ENSG00000249867* | | 5,19 | 0,016103362 |
| *ENSG00000237074* | | 5,18 | 0,022093684 |
| *ENSG00000233532* | *LINC00460* | 5,17 | 0,000217758 |
| *ENSG00000241202* | *ZIC4-AS1* | 5,16 | 0,041026579 |
| *ENSG00000226363* | *HAGLROS* | 5,15 | 0,000208642 |

| | | | |
|---|---|---|---|
| *ENSG00000243797* | | 5,12 | 0,000548692 |
| *ENSG00000237152* | *DLEU7-AS1* | 5,11 | 0,002621458 |
| *ENSG00000225087* | | 5,09 | 0,026081279 |
| *ENSG00000234695* | | 5,07 | 0,004812881 |
| *ENSG00000279375* | | 5,07 | 0,028995302 |
| *ENSG00000270607* | | 5,06 | 0,007036814 |
| *ENSG00000228412* | | 5,06 | 0,001376677 |
| *ENSG00000279047* | | 5,06 | 0,000545385 |
| *ENSG00000227619* | | 5,05 | 0,004330503 |
| *ENSG00000246526* | *LINC002481* | 5,04 | 0,006361627 |
| *ENSG00000261618* | | 5,02 | 0,001761797 |
| *ENSG00000236268* | *LINC01361* | 5,01 | 0,033451197 |
| *ENSG00000235277* | | 5,01 | 0,013485635 |
| *ENSG00000250328* | | 5,00 | 0,016812058 |
| *ENSG00000237978* | *KCNMB2-AS1* | 4,99 | 0,006018271 |
| *ENSG00000255983* | | 4,99 | 0,007941234 |
| *ENSG00000182912* | *TSPEAR-AS2* | 4,97 | 0,000367956 |
| *ENSG00000227279* | | 4,95 | 0,000707236 |
| *ENSG00000282851* | *BISPR* | 4,94 | 0,000128821 |
| *ENSG00000253307* | | 4,93 | 0,006011626 |
| *ENSG00000146521* | *LINC01558* | 4,93 | 0,033069753 |
| *ENSG00000233611* | | 4,90 | 0,000113651 |
| *ENSG00000203650* | *LINC01285* | 4,86 | 0,00820742 |
| *ENSG00000232949* | | 4,85 | 0,0330723 |
| *ENSG00000250994* | | 4,85 | 0,047653251 |
| *ENSG00000272372* | | 4,84 | 0,024919584 |
| *ENSG00000261798* | | 4,83 | 0,023992763 |
| *ENSG00000232759* | | 4,81 | 0,00674548 |
| *ENSG00000272328* | | 4,81 | 0,044954274 |
| *ENSG00000230061* | *TRPM2-AS* | 4,79 | 0,000307063 |
| *ENSG00000124915* | | 4,79 | 0,006354459 |
| *ENSG00000227076* | | 4,78 | 0,011141855 |
| *ENSG00000268894* | *PLCE1-AS1* | 4,77 | 0,018447949 |
| *ENSG00000278071* | | 4,71 | 0,03022934 |
| *ENSG00000254204* | | 4,70 | 2,7104E-09 |
| *ENSG00000229618* | | 4,65 | 3,26822E-05 |
| *ENSG00000231057* | | 4,64 | 0,004120155 |
| *ENSG00000236449* | | 4,63 | 0,007272109 |
| *ENSG00000254703* | *SENCR* | 4,59 | 0,031510101 |
| *ENSG00000283982* | | 4,57 | 0,004418177 |
| *ENSG00000259584* | | 4,57 | 0,004609141 |
| *ENSG00000250421* | | 4,57 | 0,03635629 |
| *ENSG00000228680* | | 4,57 | 0,013920676 |
| *ENSG00000177699* | | 4,53 | 2,86098E-05 |
| *ENSG00000234015* | *LINC02530* | 4,53 | 0,043296486 |

| | | | |
|---|---|---|---|
| *ENSG00000273301* | | 4,52 | 0,017775366 |
| *ENSG00000237807* | | 4,51 | 6,02013E-12 |
| *ENSG00000222017* | | 4,51 | 0,00178724 |
| *ENSG00000230836* | *LINC01293* | 4,50 | 0,000166291 |
| *ENSG00000237870* | | 4,49 | 0,000297859 |
| *ENSG00000273669* | | 4,48 | 0,009233547 |
| *ENSG00000224848* | | 4,46 | 0,023103455 |
| *ENSG00000257906* | *LINC02156* | 4,45 | 0,018447949 |
| *ENSG00000272180* | | 4,43 | 0,045746335 |
| *ENSG00000253508* | | 4,43 | 0,000114547 |
| *ENSG00000271776* | | 4,43 | 0,03839163 |
| *ENSG00000262772* | *LINC01977* | 4,42 | 0,007717307 |
| *ENSG00000271127* | | 4,41 | 0,001779954 |
| *ENSG00000268186* | *ZNF114-AS1* | 4,41 | 0,000166291 |
| *ENSG00000261305* | | 4,40 | 0,012556579 |
| *ENSG00000250320* | | 4,38 | 0,005891465 |
| *ENSG00000231528* | *FAM225A* | 4,38 | 2,19752E-05 |
| *ENSG00000257696* | | 4,37 | 0,011043759 |
| *ENSG00000258903* | | 4,36 | 0,002196351 |
| *ENSG00000248890* | *HHIP-AS1* | 4,36 | 0,010443389 |
| *ENSG00000272071* | | 4,35 | 0,012693044 |
| *ENSG00000283991* | | 4,33 | 0,018151967 |
| *ENSG00000206532* | | 4,32 | 0,04847901 |
| *ENSG00000272808* | | 4,31 | 1,59611E-06 |
| *ENSG00000163364* | *LINC01116* | 4,30 | 2,04982E-08 |
| *ENSG00000246662* | *LINC00535* | 4,30 | 0,006168 |
| *ENSG00000251365* | | 4,30 | 0,022822854 |
| *ENSG00000257732* | | 4,28 | 0,003260555 |
| *ENSG00000223764* | *LINC02593* | 4,25 | 0,00661947 |
| *ENSG00000273162* | | 4,23 | 4,55732E-07 |
| *ENSG00000260302* | | 4,21 | 0,049059113 |
| *ENSG00000260855* | | 4,17 | 0,000184645 |
| *ENSG00000231690* | *LINC00574* | 4,16 | 0,000233362 |
| *ENSG00000259354* | | 4,13 | 0,00097477 |
| *ENSG00000230729* | | 4,10 | 0,032541023 |
| *ENSG00000230333* | | 4,10 | 0,027542795 |
| *ENSG00000257515* | | 4,09 | 0,031171525 |
| *ENSG00000265349* | | 4,08 | 0,003352289 |
| *ENSG00000258842* | | 4,07 | 0,00323915 |
| *ENSG00000272769* | | 4,07 | 0,028265735 |
| *ENSG00000223485* | *LINC01615* | 4,06 | 0,00181694 |
| *ENSG00000249082* | *C5orf66-AS1* | 4,04 | 0,020612838 |
| *ENSG00000261379* | | 4,02 | 0,005891465 |
| *ENSG00000235741* | | 4,01 | 0,036761608 |
| *ENSG00000250240* | | 3,98 | 2,66136E-05 |

| | | | |
|---|---|---|---|
| *ENSG00000248554* | | 3,97 | 6,07816E-14 |
| *ENSG00000258922* | | 3,96 | 0,009819197 |
| *ENSG00000264569* | | 3,96 | 0,014144061 |
| *ENSG00000255306* | | 3,95 | 3,34984E-05 |
| *ENSG00000259306* | | 3,95 | 0,0005652 |
| *ENSG00000167912* | | 3,94 | 0,006862511 |
| *ENSG00000268686* | | 3,94 | 0,037964732 |
| *ENSG00000259701* | | 3,94 | 0,020448805 |
| *ENSG00000232110* | | 3,93 | 0,035542668 |
| *ENSG00000280029* | | 3,92 | 0,005925397 |
| *ENSG00000232555* | | 3,91 | 0,005946517 |
| *ENSG00000227039* | *ITGB2-AS1* | 3,88 | 0,000459878 |
| *ENSG00000261773* | | 3,88 | 0,032410709 |
| *ENSG00000272551* | | 3,87 | 0,015442575 |
| *ENSG00000272070* | | 3,87 | 0,003324759 |
| *ENSG00000273216* | | 3,86 | 0,018763689 |
| *ENSG00000224189* | *HAGLR* | 3,84 | 0,006769485 |
| *ENSG00000272463* | | 3,84 | 0,001969142 |
| *ENSG00000275450* | | 3,84 | 0,046063651 |
| *ENSG00000272622* | | 3,83 | 0,006210431 |
| *ENSG00000278276* | | 3,83 | 0,031130054 |
| *ENSG00000273473* | | 3,83 | 0,024778371 |
| *ENSG00000277152* | | 3,83 | 0,006617013 |
| *ENSG00000259376* | | 3,81 | 0,004189183 |
| *ENSG00000280744* | *LINC01173* | 3,81 | 0,048730581 |
| *ENSG00000185904* | *LINC00839* | 3,77 | 8,72893E-08 |
| *ENSG00000224577* | *LINC01117* | 3,76 | 1,93494E-07 |
| *ENSG00000227028* | *SLC8A1-AS1* | 3,76 | 0,003146186 |
| *ENSG00000163009* | *C2orf48* | 3,75 | 0,001346033 |
| *ENSG00000255118* | | 3,69 | 0,021075969 |
| *ENSG00000244968* | *LIFR-AS1* | 3,68 | 0,000201417 |
| *ENSG00000233728* | | 3,67 | 2,27048E-09 |
| *ENSG00000234883* | *MIR155HG* | 3,64 | 0,016570652 |
| *ENSG00000248240* | | 3,63 | 1,0269E-08 |
| *ENSG00000280890* | *ELDR* | 3,62 | 0,028745051 |
| *ENSG00000272068* | | 3,60 | 0,033055762 |
| *ENSG00000232386* | | 3,58 | 2,41816E-07 |
| *ENSG00000236643* | | 3,57 | 0,009385177 |
| *ENSG00000268289* | | 3,56 | 0,043709054 |
| *ENSG00000255082* | *GRM5-AS1* | 3,55 | 0,048437189 |
| *ENSG00000227953* | *LINC01341* | 3,55 | 0,000409342 |
| *ENSG00000281756* | *C2-AS1* | 3,53 | 0,038552204 |
| *ENSG00000270604* | *HCG17* | 3,53 | 0,021450299 |
| *ENSG00000259341* | | 3,51 | 0,031452357 |
| *ENSG00000229953* | | 3,50 | 0,013671407 |

| | | | |
|---|---|---|---|
| *ENSG00000254859* | | 3,50 | 0,000618441 |
| *ENSG00000273100* | | 3,48 | 0,035125572 |
| *ENSG00000231864* | | 3,47 | 2,14949E-08 |
| *ENSG00000227200* | | 3,46 | 0,043211334 |
| *ENSG00000266401* | | 3,43 | 0,005479872 |
| *ENSG00000251361* | | 3,41 | 0,016275283 |
| *ENSG00000232878* | *DPYD-AS1* | 3,41 | 0,000135307 |
| *ENSG00000224897* | *POT1-AS1* | 3,40 | 0,00094137 |
| *ENSG00000230798* | *FOXD3-AS1* | 3,40 | 0,018791287 |
| *ENSG00000205334* | *LINC01460* | 3,40 | 0,034276683 |
| *ENSG00000234913* | | 3,39 | 0,02964251 |
| *ENSG00000261758* | | 3,39 | 0,032159486 |
| *ENSG00000261693* | | 3,36 | 0,00141184 |
| *ENSG00000274270* | | 3,35 | 0,001211635 |
| *ENSG00000229214* | *LINC00242* | 3,35 | 0,000206183 |
| *ENSG00000244040* | *IL12A-AS1* | 3,34 | 0,020128883 |
| *ENSG00000240990* | *HOXA11-AS* | 3,34 | 3,23147E-05 |
| *ENSG00000255045* | | 3,32 | 0,014450177 |
| *ENSG00000229257* | | 3,31 | 1,41214E-07 |
| *ENSG00000277855* | | 3,30 | 0,01646457 |
| *ENSG00000261054* | | 3,29 | 9,21225E-06 |
| *ENSG00000250635* | *CXXC5-AS1* | 3,29 | 0,033105844 |
| *ENSG00000248538* | | 3,28 | 0,029976436 |
| *ENSG00000267379* | | 3,28 | 9,46261E-08 |
| *ENSG00000261586* | | 3,27 | 0,010715828 |
| *ENSG00000280734* | *LINC01232* | 3,27 | 3,34984E-05 |
| *ENSG00000260328* | | 3,27 | 0,00142262 |
| *ENSG00000272694* | | 3,25 | 0,013054112 |
| *ENSG00000227479* | | 3,25 | 0,040449719 |
| *ENSG00000259869* | | 3,24 | 0,022508836 |
| *ENSG00000272382* | | 3,22 | 0,009222757 |
| *ENSG00000267607* | | 3,21 | 0,001137434 |
| *ENSG00000272662* | | 3,21 | 0,018839808 |
| *ENSG00000266283* | | 3,19 | 0,001712198 |
| *ENSG00000268812* | | 3,17 | 0,012597354 |
| *ENSG00000260896* | *ARLNC1* | 3,17 | 0,036985543 |
| *ENSG00000235806* | | 3,17 | 0,004698533 |
| *ENSG00000243902* | | 3,16 | 0,023714745 |
| *ENSG00000266010* | *GATA6-AS1* | 3,16 | 0,007593681 |
| *ENSG00000233058* | *LINC00884* | 3,14 | 3,69234E-06 |
| *ENSG00000272405* | | 3,13 | 0,006251769 |
| *ENSG00000283554* | *LINC02341* | 3,06 | 0,043709054 |
| *ENSG00000261654* | | 3,02 | 0,004557692 |
| *ENSG00000237380* | *HOXD-AS2* | 3,01 | 0,017485042 |
| *ENSG00000165511* | *C10orf25* | 3,00 | 0,000481877 |

| | | | |
|---|---|---|---|
| *ENSG00000236031* | | 2,98 | 0,00320335 |
| *ENSG00000271930* | | 2,97 | 0,020938483 |
| *ENSG00000275322* | | 2,94 | 0,036985543 |
| *ENSG00000229688* | *ISPD-AS1* | 2,89 | 0,044233076 |
| *ENSG00000273443* | | 2,89 | 4,61408E-05 |
| *ENSG00000251131* | | 2,88 | 9,90261E-09 |
| *ENSG00000230648* | | 2,88 | 0,044210903 |
| *ENSG00000233117* | *LINC00702* | 2,88 | 0,039658335 |
| *ENSG00000260430* | | 2,85 | 0,025922411 |
| *ENSG00000254363* | | 2,85 | 0,023248889 |
| *ENSG00000236914* | *LINC01852* | 2,84 | 0,005860221 |
| *ENSG00000231187* | | 2,84 | 0,004245671 |
| *ENSG00000270823* | | 2,84 | 0,025422713 |
| *ENSG00000275223* | | 2,82 | 0,024952074 |
| *ENSG00000215068* | | 2,81 | 2,99315E-07 |
| *ENSG00000227308* | | 2,80 | 0,000149694 |
| *ENSG00000237187* | *NR2F1-AS1* | 2,80 | 0,02971427 |
| *ENSG00000214650* | | 2,79 | 0,01914151 |
| *ENSG00000275874* | *PICSAR* | 2,78 | 0,010421459 |
| *ENSG00000260190* | | 2,74 | 0,004698533 |
| *ENSG00000224764* | | 2,74 | 0,008669777 |
| *ENSG00000259345* | | 2,73 | 0,000245159 |
| *ENSG00000234076* | *TPRG1-AS1* | 2,72 | 0,033292188 |
| *ENSG00000233101* | *HOXB-AS3* | 2,71 | 0,019748046 |
| *ENSG00000225411* | | 2,71 | 0,014102704 |
| *ENSG00000226609* | | 2,71 | 0,033434709 |
| *ENSG00000259409* | *BMF-AS1* | 2,70 | 0,013485635 |
| *ENSG00000258733* | *LINC02328* | 2,70 | 0,031462979 |
| *ENSG00000283078* | | 2,69 | 0,005621627 |
| *ENSG00000260000* | | 2,68 | 1,48009E-05 |
| *ENSG00000272620* | *AFAP1-AS1* | 2,65 | 0,022317586 |
| *ENSG00000282386* | | 2,65 | 0,000164748 |
| *ENSG00000273026* | | 2,64 | 0,004012249 |
| *ENSG00000258168* | | 2,64 | 0,027397395 |
| *ENSG00000225489* | | 2,63 | 0,005860221 |
| *ENSG00000224905* | | 2,60 | 0,015945589 |
| *ENSG00000258101* | | 2,60 | 0,002243684 |
| *ENSG00000227963* | *RBM15-AS1* | 2,60 | 0,003606431 |
| *ENSG00000260633* | | 2,60 | 0,008333086 |
| *ENSG00000250697* | | 2,60 | 0,02029801 |
| *ENSG00000243176* | | 2,59 | 0,008009008 |
| *ENSG00000235204* | | 2,59 | 0,023103455 |
| *ENSG00000272825* | | 2,58 | 0,01836873 |
| *ENSG00000244327* | | 2,56 | 0,03053687 |
| *ENSG00000273210* | | 2,55 | 0,022057655 |

| | | | |
|---|---|---|---|
| *ENSG00000231948* | *HS1BP3-IT1* | 2,55 | 0,016066611 |
| *ENSG00000261211* | | 2,53 | 0,006724738 |
| *ENSG00000239332* | *LINC01119* | 2,53 | 0,020403494 |
| *ENSG00000249572* | | 2,50 | 0,009559785 |
| *ENSG00000260400* | | 2,50 | 0,004982759 |
| *ENSG00000260588* | | 2,49 | 0,000987029 |
| *ENSG00000225792* | | 2,49 | 0,00252136 |
| *ENSG00000236682* | | 2,49 | 0,003153604 |
| *ENSG00000283828* | | 2,48 | 0,021230933 |
| *ENSG00000213888* | *LINC01521* | 2,47 | 1,02172E-07 |
| *ENSG00000231764* | *DLX6-AS1* | 2,47 | 0,036761608 |
| *ENSG00000251602* | | 2,47 | 0,000450856 |
| *ENSG00000278709* | *NKILA* | 2,47 | 0,00187862 |
| *ENSG00000259347* | | 2,45 | 0,006478447 |
| *ENSG00000260293* | | 2,44 | 0,005405555 |
| *ENSG00000273998* | | 2,43 | 0,015326758 |
| *ENSG00000255966* | | 2,42 | 0,036544271 |
| *ENSG00000259062* | *ACTN1-AS1* | 2,41 | 0,025244771 |
| *ENSG00000272501* | | 2,40 | 0,000435113 |
| *ENSG00000259368* | | 2,40 | 0,023944162 |
| *ENSG00000277767* | | 2,40 | 0,00928415 |
| *ENSG00000276075* | | 2,40 | 0,016604674 |
| *ENSG00000259215* | | 2,39 | 0,044954274 |
| *ENSG00000260704* | *LINC00543* | 2,38 | 0,022553716 |
| *ENSG00000226659* | | 2,38 | 0,046119589 |
| *ENSG00000272183* | | 2,37 | 1,48497E-05 |
| *ENSG00000240996* | | 2,37 | 0,006044895 |
| *ENSG00000247317* | *LY6E-DT* | 2,35 | 0,031462979 |
| *ENSG00000276744* | | 2,34 | 0,004217418 |
| *ENSG00000256802* | | 2,34 | 0,009390061 |
| *ENSG00000263015* | | 2,31 | 0,044548287 |
| *ENSG00000230555* | | 2,30 | 0,001446742 |
| *ENSG00000272301* | | 2,29 | 0,01701061 |
| *ENSG00000227908* | | 2,28 | 0,004071419 |
| *ENSG00000257097* | *CLIP1-AS1* | 2,28 | 0,005660174 |
| *ENSG00000261762* | | 2,26 | 0,011881555 |
| *ENSG00000277619* | | 2,26 | 0,02630431 |
| *ENSG00000236242* | *MYO16-AS1* | 2,25 | 0,024952074 |
| *ENSG00000274220* | | 2,25 | 0,000411061 |
| *ENSG00000253741* | *LNCOC1* | 2,25 | 1,65704E-08 |
| *ENSG00000273117* | | 2,25 | 3,6217E-06 |
| *ENSG00000182648* | *LINC01006* | 2,24 | 0,006522036 |
| *ENSG00000273335* | | 2,22 | 0,036950598 |
| *ENSG00000203999* | *LINC01270* | 2,22 | 0,018258046 |
| *ENSG00000197536* | *C5orf56* | 2,21 | 0,010742245 |

| | | | |
|---|---|---|---|
| *ENSG00000273004* | | 2,21 | 0,009664853 |
| *ENSG00000272509* | | 2,21 | 0,005169538 |
| *ENSG00000258334* | | 2,21 | 0,000450856 |
| *ENSG00000238113* | *LINC01410* | 2,20 | 9,486E-06 |
| *ENSG00000277159* | | 2,20 | 0,032874637 |
| *ENSG00000235501* | | 2,19 | 0,008929311 |
| *ENSG00000267356* | | 2,17 | 0,008388715 |
| *ENSG00000248079* | *DPH6-DT* | 2,16 | 0,026193491 |
| *ENSG00000276931* | | 2,14 | 0,011913475 |
| *ENSG00000260495* | | 2,14 | 0,014400472 |
| *ENSG00000254389* | *RHPN1-AS1* | 2,13 | 0,04847901 |
| *ENSG00000183250* | *LINC01547* | 2,13 | 0,001856198 |
| *ENSG00000259953* | | 2,12 | 0,006043695 |
| *ENSG00000253552* | *HOXA-AS2* | 2,12 | 0,018201928 |
| *ENSG00000253395* | | 2,11 | 0,049731469 |
| *ENSG00000227542* | | 2,11 | 0,006554872 |
| *ENSG00000235823* | *OLMALINC* | 2,10 | 0,00155424 |
| *ENSG00000253636* | | 2,10 | 0,004327436 |
| *ENSG00000255692* | | 2,09 | 0,000691283 |
| *ENSG00000204054* | *LINC00963* | 2,09 | 0,00046068 |
| *ENSG00000238058* | | 2,08 | 0,000360507 |
| *ENSG00000224358* | | 2,07 | 3,82764E-05 |
| *ENSG00000177738* | | 2,07 | 2,12247E-11 |
| *ENSG00000225077* | *LINC00337* | 2,07 | 0,033171124 |
| *ENSG00000225032* | | 2,06 | 0,041951427 |
| *ENSG00000274849* | | 2,06 | 0,026245611 |
| *ENSG00000259884* | | 2,06 | 0,025023505 |
| *ENSG00000274528* | | 2,06 | 0,006654407 |
| *ENSG00000225361* | *PPP1R26-AS1* | 2,06 | 0,003765042 |
| *ENSG00000268996* | *MAN1B1-DT* | 2,05 | 0,010047915 |
| *ENSG00000245614* | *DDX11-AS1* | 2,03 | 0,008938272 |
| *ENSG00000274922* | | 2,03 | 0,04176694 |
| *ENSG00000259438* | *MAPK6-DT* | 2,03 | 0,030211245 |
| *ENSG00000279873* | *LINC01126* | 2,02 | 0,002480544 |
| *ENSG00000213057* | *C1orf220* | 2,01 | 0,001268659 |
| *ENSG00000250159* | | 2,01 | 0,000450856 |
| *ENSG00000227803* | | 1,99 | 0,049621281 |
| *ENSG00000237301* | | 1,97 | 0,016268992 |
| *ENSG00000271888* | | 1,96 | 0,005815059 |
| *ENSG00000239718* | *HLTF-AS1* | 1,96 | 0,029349841 |
| *ENSG00000251018* | *HMMR-AS1* | 1,94 | 0,019759248 |
| *ENSG00000269038* | | 1,94 | 0,021979424 |
| *ENSG00000276603* | | 1,93 | 0,029976436 |
| *ENSG00000224418* | *STK24-AS1* | 1,90 | 0,008110884 |
| *ENSG00000206344* | *HCG27* | 1,89 | 0,002441575 |

| | | | |
|---|---|---|---|
| *ENSG00000261266* | | 1,89 | 0,001255775 |
| *ENSG00000246334* | *PRR7-AS1* | 1,89 | 0,044930126 |
| *ENSG00000233654* | | 1,88 | 0,002530337 |
| *ENSG00000242147* | | 1,88 | 0,014706061 |
| *ENSG00000204706* | *MAMDC2-AS1* | 1,88 | 0,043209904 |
| *ENSG00000226180* | | 1,86 | 0,030714049 |
| *ENSG00000254812* | | 1,86 | 0,025736178 |
| *ENSG00000257596* | | 1,86 | 0,001996774 |
| *ENSG00000250220* | | 1,86 | 0,015451967 |
| *ENSG00000271821* | | 1,85 | 0,010172262 |
| *ENSG00000272115* | | 1,85 | 0,006286042 |
| *ENSG00000257038* | | 1,85 | 0,001837674 |
| *ENSG00000259209* | | 1,85 | 0,034909116 |
| *ENSG00000239523* | *MYLK-AS1* | 1,84 | 0,023738538 |
| *ENSG00000255498* | | 1,84 | 0,033226954 |
| *ENSG00000271966* | | 1,84 | 0,02546948 |
| *ENSG00000276728* | | 1,84 | 0,008202895 |
| *ENSG00000237753* | | 1,83 | 0,029847089 |
| *ENSG00000271781* | | 1,83 | 0,006554872 |
| *ENSG00000261094* | | 1,82 | 5,46504E-10 |
| *ENSG00000268364* | *SMC5-AS1* | 1,81 | 0,006091582 |
| *ENSG00000257607* | | 1,81 | 0,030162258 |
| *ENSG00000246523* | | 1,81 | 0,028676541 |
| *ENSG00000272764* | | 1,81 | 0,022046482 |
| *ENSG00000266495* | | 1,81 | 0,014783249 |
| *ENSG00000271646* | | 1,80 | 0,002013424 |
| *ENSG00000231113* | | 1,80 | 0,002611696 |
| *ENSG00000273203* | | 1,80 | 0,014481724 |
| *ENSG00000267201* | *LINC01775* | 1,78 | 0,010537137 |
| *ENSG00000276116* | *FUT8-AS1* | 1,77 | 0,013632135 |
| *ENSG00000267751* | | 1,77 | 0,035053651 |
| *ENSG00000273888* | *FRMD6-AS1* | 1,75 | 0,006478447 |
| *ENSG00000284052* | | 1,74 | 0,01454813 |
| *ENSG00000270006* | | 1,74 | 0,017938271 |
| *ENSG00000273702* | | 1,74 | 0,014102704 |
| *ENSG00000255135* | | 1,73 | 0,001299346 |
| *ENSG00000263731* | | 1,73 | 0,000791552 |
| *ENSG00000260063* | | 1,72 | 0,017973304 |
| *ENSG00000250899* | | 1,71 | 7,70249E-05 |
| *ENSG00000232600* | *TONSL-AS1* | 1,71 | 0,003430879 |
| *ENSG00000277142* | *LINC00235* | 1,71 | 0,005457116 |
| *ENSG00000257702* | *LBX2-AS1* | 1,70 | 0,003734684 |
| *ENSG00000270959* | *LPP-AS2* | 1,70 | 9,2689E-05 |
| *ENSG00000224046* | | 1,70 | 0,00365964 |
| *ENSG00000260742* | | 1,68 | 0,022436591 |

| | | | |
|---|---|---|---|
| *ENSG00000255031* | | 1,68 | 2,40112E-07 |
| *ENSG00000246777* | | 1,68 | 0,033664827 |
| *ENSG00000258534* | | 1,67 | 0,037228358 |
| *ENSG00000257167* | *TMPO-AS1* | 1,66 | 0,000606368 |
| *ENSG00000242474* | | 1,66 | 0,012134402 |
| *ENSG00000262001* | *DLGAP1-AS2* | 1,65 | 0,010855892 |
| *ENSG00000243849* | *CFAP44-AS1* | 1,65 | 0,002508101 |
| *ENSG00000278383* | | 1,64 | 0,047619638 |
| *ENSG00000258701* | *LINC00638* | 1,64 | 0,020345445 |
| *ENSG00000260874* | | 1,64 | 0,003765171 |
| *ENSG00000273355* | | 1,63 | 0,024598874 |
| *ENSG00000228544* | *CCDC183-AS1* | 1,63 | 0,006282026 |
| *ENSG00000253210* | | 1,62 | 0,026414041 |
| *ENSG00000177337* | *DLGAP1-AS1* | 1,61 | 0,021890083 |
| *ENSG00000236423* | *LINC01134* | 1,61 | 0,036383792 |
| *ENSG00000228919* | | 1,61 | 0,049503279 |
| *ENSG00000258725* | *PRC1-AS1* | 1,59 | 0,006396115 |
| *ENSG00000259826* | | 1,59 | 0,0145852 |
| *ENSG00000228434* | | 1,55 | 0,007396021 |
| *ENSG00000277369* | | 1,55 | 0,009092517 |
| *ENSG00000231770* | *TMEM44-AS1* | 1,55 | 0,008682681 |
| *ENSG00000276384* | | 1,54 | 0,010715971 |
| *ENSG00000259460* | | 1,54 | 0,021261932 |
| *ENSG00000231304* | *SGO1-AS1* | 1,53 | 0,017973304 |
| *ENSG00000272767* | *JMJD1C-AS1* | 1,53 | 0,038726511 |
| *ENSG00000280303* | *ERICD* | 1,53 | 0,019093544 |
| *ENSG00000266947* | | 1,53 | 0,017491886 |
| *ENSG00000229152* | *ANKRD10-IT1* | 1,53 | 0,012138719 |
| *ENSG00000272324* | | 1,53 | 0,019446652 |
| *ENSG00000225951* | *ODF2-AS1* | 1,53 | 0,016046477 |
| *ENSG00000161149* | *TUBA3FP* | 1,52 | 0,038321715 |
| *ENSG00000237491* | | 1,50 | 0,011326571 |
| *ENSG00000284428* | | 1,50 | 0,021979424 |
| *ENSG00000259891* | | 1,49 | 3,36318E-06 |
| *ENSG00000265393* | | 1,48 | 0,007776862 |
| *ENSG00000255886* | | 1,47 | 0,005782477 |
| *ENSG00000253806* | | 1,46 | 0,035271779 |
| *ENSG00000224635* | | 1,46 | 0,044354143 |
| *ENSG00000228109* | *MELTF-AS1* | 1,44 | 0,010848815 |
| *ENSG00000275437* | | 1,44 | 0,000370791 |
| *ENSG00000224020* | *MIR181A2HG* | 1,44 | 0,027025274 |
| *ENSG00000238186* | | 1,44 | 0,004698533 |
| *ENSG00000272721* | | 1,43 | 0,039364099 |
| *ENSG00000272768* | | 1,43 | 0,000162375 |
| *ENSG00000267321* | *LINC02001* | 1,43 | 5,90311E-05 |

| | | | |
|---|---|---|---|
| *ENSG00000260773* | | 1,43 | 0,000293013 |
| *ENSG00000179743* | | 1,40 | 0,012723096 |
| *ENSG00000273486* | | 1,40 | 0,004755762 |
| *ENSG00000239213* | *NCK1-DT* | 1,40 | 0,00553504 |
| *ENSG00000272870* | | 1,39 | 0,047898686 |
| *ENSG00000258789* | | 1,38 | 0,03967463 |
| *ENSG00000259877* | | 1,38 | 0,043709054 |
| *ENSG00000275557* | | 1,38 | 0,024621364 |
| *ENSG00000232586* | *KIAA1614-AS1* | 1,38 | 0,040671939 |
| *ENSG00000230185* | *C9orf147* | 1,35 | 0,045452924 |
| *ENSG00000176124* | *DLEU1* | 1,35 | 0,002699615 |
| *ENSG00000259523* | | 1,35 | 0,026414041 |
| *ENSG00000187951* | | 1,35 | 2,25708E-05 |
| *ENSG00000188242* | | 1,35 | 0,017440649 |
| *ENSG00000272994* | | 1,33 | 0,003779836 |
| *ENSG00000272195* | | 1,33 | 0,005745324 |
| *ENSG00000274265* | | 1,32 | 0,016709801 |
| *ENSG00000259426* | | 1,32 | 0,005080559 |
| *ENSG00000271122* | | 1,32 | 0,046114492 |
| *ENSG00000231305* | | 1,31 | 0,02866502 |
| *ENSG00000246323* | | 1,31 | 0,000858288 |
| *ENSG00000261759* | | 1,29 | 0,002238906 |
| *ENSG00000278206* | | 1,29 | 0,000790263 |
| *ENSG00000223478* | | 1,27 | 0,015041117 |
| *ENSG00000188693* | *CYP51A1-AS1* | 1,27 | 0,005530435 |
| *ENSG00000228801* | | 1,26 | 0,007722197 |
| *ENSG00000273329* | | 1,25 | 0,041543877 |
| *ENSG00000246889* | | 1,25 | 0,048940238 |
| *ENSG00000246851* | | 1,24 | 0,001930863 |
| *ENSG00000245149* | *RNF139-AS1* | 1,22 | 0,001267809 |
| *ENSG00000230149* | | 1,22 | 0,025511886 |
| *ENSG00000261188* | | 1,21 | 0,005457116 |
| *ENSG00000248092* | *NNT-AS1* | 1,20 | 0,004285221 |
| *ENSG00000254551* | | 1,19 | 0,042711381 |
| *ENSG00000235530* | | 1,19 | 0,014518978 |
| *ENSG00000263235* | | 1,19 | 0,012283661 |
| *ENSG00000196366* | *C9orf163* | 1,17 | 0,010811453 |
| *ENSG00000227278* | | 1,17 | 0,000293013 |
| *ENSG00000255153* | *TOLLIP-AS1* | 1,17 | 0,027328994 |
| *ENSG00000228395* | | 1,17 | 0,006385718 |
| *ENSG00000237188* | | 1,16 | 0,044366056 |
| *ENSG00000260924* | *LINC01311* | 1,15 | 0,010090118 |
| *ENSG00000247092* | *SNHG10* | 1,15 | 0,006102171 |
| *ENSG00000256712* | | 1,14 | 0,008496797 |
| *ENSG00000260552* | | 1,14 | 0,040671939 |

| | | | |
|---|---|---|---|
| *ENSG00000185065* | | 1,13 | 0,025384797 |
| *ENSG00000228106* | | 1,11 | 0,01108627 |
| *ENSG00000260774* | | 1,08 | 0,028534206 |
| *ENSG00000255629* | | 1,08 | 0,031910745 |
| *ENSG00000267100* | *ILF3-DT* | 1,08 | 0,034871363 |
| *ENSG00000276529* | | 1,06 | 0,007969557 |
| *ENSG00000236986* | | 1,06 | 0,00042716 |
| *ENSG00000264456* | | 1,06 | 0,040742326 |
| *ENSG00000261008* | *LINC01572* | 1,05 | 0,020685553 |
| *ENSG00000247934* | | 1,05 | 0,030494454 |
| *ENSG00000232671* | | 1,03 | 0,034712835 |
| *ENSG00000225791* | *TRAM2-AS1* | 1,03 | 0,041221349 |
| *ENSG00000253327* | *RAD21-AS1* | 1,03 | 0,000822845 |
| *ENSG00000247228* | | 1,02 | 0,029969944 |
| *ENSG00000196810* | *CTBP1-DT* | 1,02 | 0,034986518 |
| *ENSG00000274425* | | 1,01 | 0,046195505 |
| *ENSG00000280195* | | -1,01 | 0,019034308 |
| *ENSG00000234203* | | -1,01 | 0,016537541 |
| *ENSG00000222043* | | -1,05 | 0,007285927 |
| *ENSG00000232931* | *LINC00342* | -1,06 | 0,01251703 |
| *ENSG00000238184* | *CD81-AS1* | -1,06 | 0,000316583 |
| *ENSG00000269946* | | -1,08 | 0,043875023 |
| *ENSG00000238279* | | -1,09 | 0,034246317 |
| *ENSG00000259198* | | -1,11 | 0,00237506 |
| *ENSG00000272293* | | -1,11 | 0,006069055 |
| *ENSG00000224660* | *SH3BP5-AS1* | -1,11 | 5,68919E-05 |
| *ENSG00000229839* | | -1,12 | 0,003405938 |
| *ENSG00000250379* | | -1,12 | 0,001592135 |
| *ENSG00000258451* | | -1,13 | 0,034524016 |
| *ENSG00000237232* | *ZNF295-AS1* | -1,16 | 0,02605982 |
| *ENSG00000273419* | | -1,17 | 0,00267231 |
| *ENSG00000224032* | *EPB41L4A-AS1* | -1,19 | 0,000721283 |
| *ENSG00000267551* | | -1,19 | 0,026232836 |
| *ENSG00000257453* | | -1,22 | 0,0423337 |
| *ENSG00000270751* | *FBXW7-AS1* | -1,23 | 0,007997581 |
| *ENSG00000257181* | | -1,28 | 0,010855892 |
| *ENSG00000226438* | | -1,29 | 0,00427531 |
| *ENSG00000177410* | *ZFAS1* | -1,31 | 0,010153712 |
| *ENSG00000254860* | *TMEM9B-AS1* | -1,37 | 0,000450856 |
| *ENSG00000235831* | *BHLHE40-AS1* | -1,40 | 0,003763597 |
| *ENSG00000244676* | | -1,40 | 0,002360702 |
| *ENSG00000241288* | | -1,42 | 0,046628677 |
| *ENSG00000246763* | *RGMB-AS1* | -1,45 | 0,000399126 |
| *ENSG00000264215* | | -1,47 | 0,036176802 |
| *ENSG00000226715* | *LINC01709* | -1,49 | 4,7014E-05 |

| | | | |
|---|---|---|---|
| *ENSG00000164621* | *SMAD5-AS1* | -1,53 | 0,005758882 |
| *ENSG00000267251* | | -1,53 | 0,00012844 |
| *ENSG00000276952* | | -1,53 | 0,019288314 |
| *ENSG00000266402* | *SNHG25* | -1,56 | 0,032583474 |
| *ENSG00000237781* | | -1,57 | 0,049454437 |
| *ENSG00000175061* | *LRRC75A-AS1* | -1,58 | 0,001406425 |
| *ENSG00000265496* | | -1,60 | 0,000400867 |
| *ENSG00000203804* | *ADAMTSL4-AS1* | -1,60 | 0,038188577 |
| *ENSG00000273149* | | -1,61 | 0,00012229 |
| *ENSG00000261668* | | -1,68 | 0,019775641 |
| *ENSG00000271730* | | -1,68 | 0,006097028 |
| *ENSG00000203875* | *SNHG5* | -1,69 | 0,044542544 |
| *ENSG00000234741* | *GAS5* | -1,69 | 5,94822E-05 |
| *ENSG00000270084* | *GAS5-AS1* | -1,71 | 0,000274599 |
| *ENSG00000247903* | | -1,78 | 6,47671E-06 |
| *ENSG00000236404* | *VLDLR-AS1* | -1,80 | 0,027046054 |
| *ENSG00000257681* | | -1,84 | 0,040671939 |
| *ENSG00000272112* | | -1,84 | 0,038703358 |
| *ENSG00000203709* | *MIR29B2CHG* | -1,85 | 0,013573577 |
| *ENSG00000234546* | *LNCTAM34A* | -1,88 | 0,004120155 |
| *ENSG00000262884* | | -1,89 | 2,70398E-05 |
| *ENSG00000251095* | | -1,93 | 0,000373681 |
| *ENSG00000237499* | | -1,94 | 0,004609141 |
| *ENSG00000269893* | *SNHG8* | -2,00 | 1,94112E-05 |
| *ENSG00000254936* | | -2,00 | 0,01872092 |
| *ENSG00000236234* | | -2,00 | 0,038873082 |
| *ENSG00000248787* | | -2,08 | 0,017375669 |
| *ENSG00000229881* | | -2,09 | 6,66691E-05 |
| *ENSG00000245261* | | -2,15 | 0,00012215 |
| *ENSG00000260948* | | -2,19 | 0,001017135 |
| *ENSG00000249379* | | -2,22 | 0,001757593 |
| *ENSG00000259964* | *THSD4-AS1* | -2,22 | 0,04630369 |
| *ENSG00000249252* | | -2,23 | 0,001366327 |
| *ENSG00000274904* | | -2,25 | 0,000603769 |
| *ENSG00000255241* | | -2,37 | 0,004043197 |
| *ENSG00000223701* | *RAET1E-AS1* | -2,40 | 0,04854038 |
| *ENSG00000230417* | *LINC00595* | -2,44 | 0,015870147 |
| *ENSG00000268119* | | -2,48 | 0,014904445 |
| *ENSG00000257495* | *KRT73-AS1* | -2,55 | 0,02732771 |
| *ENSG00000275620* | | -2,59 | 0,039001647 |
| *ENSG00000224251* | | -2,61 | 0,029227962 |
| *ENSG00000229891* | *LINC01315* | -2,67 | 0,010443389 |
| *ENSG00000274341* | | -2,69 | 0,000247618 |
| *ENSG00000255470* | | -2,69 | 0,004629837 |
| *ENSG00000224387* | | -2,71 | 0,041551831 |

| | | | |
|---|---|---|---|
| *ENSG00000269349* | | -2,71 | 0,03957766 |
| *ENSG00000228417* | | -2,80 | 6,86383E-07 |
| *ENSG00000269989* | | -2,83 | 0,001636935 |
| *ENSG00000231768* | *LINC01354* | -2,88 | 0,027589975 |
| *ENSG00000231840* | | -2,89 | 0,000821502 |
| *ENSG00000246100* | *LINC00900* | -2,90 | 0,041921569 |
| *ENSG00000261996* | | -2,94 | 0,000563785 |
| *ENSG00000229393* | | -2,98 | 0,005457116 |
| *ENSG00000281162* | *LINC01127* | -3,07 | 0,031781324 |
| *ENSG00000226599* | | -3,08 | 0,001137434 |
| *ENSG00000248663* | *LINC00992* | -3,14 | 0,013230726 |
| *ENSG00000251320* | | -3,17 | 0,038318868 |
| *ENSG00000257599* | *OVCH1-AS1* | -3,19 | 0,000202978 |
| *ENSG00000278668* | | -3,20 | 0,012872244 |
| *ENSG00000237975* | *FLG-AS1* | -3,26 | 0,014230824 |
| *ENSG00000224822* | *THRB-IT1* | -3,28 | 0,022057655 |
| *ENSG00000246740* | *PLA2G4E-AS1* | -3,28 | 0,017991657 |
| *ENSG00000259704* | | -3,29 | 0,010559047 |
| *ENSG00000248587* | *GDNF-AS1* | -3,35 | 0,037419725 |
| *ENSG00000251191* | *LINC00589* | -3,38 | 0,000628848 |
| *ENSG00000253125* | | -3,43 | 0,005839955 |
| *ENSG00000236915* | | -3,48 | 0,041235211 |
| *ENSG00000249790* | | -3,51 | 7,17442E-06 |
| *ENSG00000181800* | *CELF2-AS1* | -3,52 | 0,016546924 |
| *ENSG00000231683* | | -3,56 | 0,000134427 |
| *ENSG00000230113* | | -3,60 | 0,002440598 |
| *ENSG00000251364* | | -3,61 | 0,000101881 |
| *ENSG00000230990* | | -3,63 | 0,006874039 |
| *ENSG00000235481* | *UBE2R2-AS1* | -3,64 | 0,000145442 |
| *ENSG00000265743* | | -3,65 | 0,039311332 |
| *ENSG00000215386* | *MIR99AHG* | -3,66 | 0,006346378 |
| *ENSG00000234756* | | -3,67 | 0,014668756 |
| *ENSG00000267077* | | -3,67 | 0,005162 |
| *ENSG00000276542* | | -3,69 | 7,0726E-06 |
| *ENSG00000236427* | | -3,69 | 0,005033886 |
| *ENSG00000261268* | | -3,71 | 0,000429449 |
| *ENSG00000270547* | *LINC01235* | -3,72 | 0,011746924 |
| *ENSG00000232855* | | -3,73 | 0,030594651 |
| *ENSG00000272865* | | -3,76 | 0,042594191 |
| *ENSG00000262619* | *LINC00621* | -3,78 | 0,00064089 |
| *ENSG00000225611* | *LINC02158* | -3,79 | 0,014631329 |
| *ENSG00000267462* | | -3,83 | 0,005395658 |
| *ENSG00000224307* | | -3,88 | 0,00323915 |
| *ENSG00000257500* | | -3,95 | 0,035314672 |
| *ENSG00000272234* | | -3,96 | 0,014199272 |

| | | | |
|---|---|---|---|
| *ENSG00000269985* | | -4,12 | 0,013418328 |
| *ENSG00000204666* | | -4,12 | 2,1708E-06 |
| *ENSG00000264727* | | -4,12 | 0,028787136 |
| *ENSG00000257797* | | -4,30 | 0,031781324 |
| *ENSG00000276241* | | -4,33 | 0,034212136 |
| *ENSG00000259541* | | -4,47 | 0,014281835 |
| *ENSG00000274238* | | -4,52 | 4,77203E-07 |
| *ENSG00000251636* | *LINC01218* | -4,53 | 0,004421267 |
| *ENSG00000237923* | *LINC02570* | -4,54 | 0,018264564 |
| *ENSG00000205106* | | -4,58 | 0,014706061 |
| *ENSG00000255091* | | -4,59 | 0,033280351 |
| *ENSG00000223863* | *LINC01805* | -4,66 | 0,042146459 |
| *ENSG00000265888* | *DSCAS* | -4,68 | 1,13423E-05 |
| *ENSG00000256969* | | -4,69 | 0,015750222 |
| *ENSG00000253766* | | -4,74 | 1,04484E-05 |
| *ENSG00000225243* | | -4,99 | 0,026837147 |
| *ENSG00000224957* | *LINC01266* | -5,04 | 0,002599007 |
| *ENSG00000261026* | | -5,06 | 0,002622767 |
| *ENSG00000269745* | | -5,09 | 0,000590808 |
| *ENSG00000267905* | | -5,14 | 5,1016E-06 |
| *ENSG00000237290* | *LINC01343* | -5,27 | 5,18268E-06 |
| *ENSG00000230587* | *LINC02580* | -5,31 | 0,000217758 |
| *ENSG00000234477* | | -5,33 | 0,00173949 |
| *ENSG00000263698* | | -5,46 | 0,000197999 |
| *ENSG00000249923* | | -5,85 | 0,00076408 |
| *ENSG00000227115* | *LINC01630* | -6,12 | 3,78382E-07 |
| *ENSG00000257700* | | -6,17 | 1,81433E-05 |
| *ENSG00000258791* | *LINC00520* | -6,24 | 3,7696E-09 |
| *ENSG00000228528* | | -6,38 | 3,09954E-09 |
| *ENSG00000260469* | *INSYN1-AS1* | -6,62 | 0,000829706 |
| *ENSG00000269729* | | -6,67 | 0,000201417 |
| *ENSG00000254101* | *LINC02055* | -6,85 | 0,008804124 |
| *ENSG00000253715* | | -6,90 | 2,02913E-08 |
| *ENSG00000274415* | | -7,39 | 9,90261E-09 |
| *ENSG00000206552* | *KRBOX1-AS1* | -7,58 | 9,32341E-13 |
| *ENSG00000166770* | *ZNF667-AS1* | -10,42 | 6,80342E-33 |

**Supplement table S2.** Significantly regulated mRNAs in cSCC cells compared to NHEKs. Differentially expressed genes between cSCC and NHEKs were considered as significantly differentially expressed, if adjusted p-value (FDR) was less than 0.05 and absolute log2 fold change of expression level over 1.

| **Ensembl.ID** | **HGNC.symbol** | **Log2.foldchange** | **Adjusted.p.value** |
|---|---|---|---|
| *ENSG00000185905* | *C16orf54* | 24,93 | 9,62004E-14 |
| *ENSG00000163286* | *ALPG* | 23,40 | 2,42271E-10 |

| | | | |
|---|---|---|---|
| *ENSG00000137745* | *MMP13* | 10,42 | 1,30228E-08 |
| *ENSG00000183778* | *B3GALT5* | 10,16 | 1,80236E-07 |
| *ENSG00000166415* | *WDR72* | 10,08 | 6,51162E-06 |
| *ENSG00000263961* | *RHEX* | 9,67 | 0,00043091 |
| *ENSG00000131203* | *IDO1* | 9,51 | 0,000207112 |
| *ENSG00000163283* | *ALPP* | 9,49 | 2,50805E-05 |
| *ENSG00000179348* | *GATA2* | 9,47 | 1,26855E-15 |
| *ENSG00000147381* | *MAGEA4* | 9,28 | 8,77192E-07 |
| *ENSG00000130957* | *FBP2* | 8,73 | 6,57625E-09 |
| *ENSG00000128714* | *HOXD13* | 8,67 | 9,47186E-05 |
| *ENSG00000128710* | *HOXD10* | 8,52 | 1,09911E-09 |
| *ENSG00000164362* | *TERT* | 8,33 | 2,98938E-08 |
| *ENSG00000143355* | *LHX9* | 8,33 | 0,000606368 |
| *ENSG00000183072* | *NKX2-5* | 8,29 | 0,004602127 |
| *ENSG00000150625* | *GPM6A* | 8,26 | 2,55086E-06 |
| *ENSG00000128713* | *HOXD11* | 8,12 | 2,25545E-06 |
| *ENSG00000172000* | *ZNF556* | 8,03 | 0,011086479 |
| *ENSG00000142408* | *CACNG8* | 7,99 | 0,001283835 |
| *ENSG00000163083* | *INHBB* | 7,90 | 0,000166291 |
| *ENSG00000163825* | *RTP3* | 7,84 | 0,004554641 |
| *ENSG00000205403* | *CFI* | 7,82 | 4,42131E-06 |
| *ENSG00000165323* | *FAT3* | 7,74 | 3,69234E-06 |
| *ENSG00000089558* | *KCNH4* | 7,73 | 0,000234383 |
| *ENSG00000268916* | *CSAG3* | 7,62 | 0,001352421 |
| *ENSG00000153404* | *PLEKHG4B* | 7,56 | 4,95471E-11 |
| *ENSG00000168874* | *ATOH8* | 7,56 | 1,03813E-05 |
| *ENSG00000042980* | *ADAM28* | 7,52 | 0,000518531 |
| *ENSG00000182836* | *PLCXD3* | 7,51 | 0,00267446 |
| *ENSG00000066382* | *MPPED2* | 7,51 | 0,013661494 |
| *ENSG00000179331* | *RAB39A* | 7,44 | 0,00565188 |
| *ENSG00000196104* | *SPOCK3* | 7,40 | 2,47636E-05 |
| *ENSG00000124212* | *PTGIS* | 7,40 | 0,013632135 |
| *ENSG00000179083* | *FAM133A* | 7,39 | 0,014100424 |
| *ENSG00000174837* | *ADGRE1* | 7,38 | 0,006043695 |
| *ENSG00000132744* | *ACY3* | 7,32 | 2,69594E-05 |
| *ENSG00000176692* | *FOXC2* | 7,22 | 2,4955E-05 |
| *ENSG00000116785* | *CFHR3* | 7,18 | 0,000873943 |
| *ENSG00000055813* | *CCDC85A* | 7,18 | 0,000748575 |
| *ENSG00000145113* | *MUC4* | 7,17 | 0,000242877 |
| *ENSG00000179023* | *KLHDC7A* | 7,14 | 0,018242846 |
| *ENSG00000173391* | *OLR1* | 7,08 | 1,79942E-07 |
| *ENSG00000134258* | *VTCN1* | 7,07 | 0,000482924 |
| *ENSG00000163568* | *AIM2* | 7,04 | 6,28825E-20 |
| *ENSG00000170689* | *HOXB9* | 6,98 | 6,7335E-07 |
| *ENSG00000151025* | *GPR158* | 6,96 | 0,000133598 |

| | | | |
|---|---|---|---|
| *ENSG00000172927* | *MYEOV* | 6,94 | 3,10671E-07 |
| *ENSG00000163554* | *SPTA1* | 6,92 | 0,001472311 |
| *ENSG00000148942* | *SLC5A12* | 6,90 | 0,000745927 |
| *ENSG00000156466* | *GDF6* | 6,88 | 0,004987464 |
| *ENSG00000168243* | *GNG4* | 6,87 | 1,06825E-09 |
| *ENSG00000197249* | *SERPINA1* | 6,86 | 1,62887E-05 |
| *ENSG00000159184* | *HOXB13* | 6,85 | 0,004372167 |
| *ENSG00000162896* | *PIGR* | 6,83 | 0,001750014 |
| *ENSG00000205436* | *EXOC3L4* | 6,83 | 3,3301E-05 |
| *ENSG00000197587* | *DMBX1* | 6,83 | 0,006111561 |
| *ENSG00000163623* | *NKX6-1* | 6,82 | 0,000228458 |
| *ENSG00000106031* | *HOXA13* | 6,79 | 1,79892E-06 |
| *ENSG00000198443* | *KRTAP4-1* | 6,69 | 0,000134514 |
| *ENSG00000120328* | *PCDHB12* | 6,66 | 3,30165E-05 |
| *ENSG00000137142* | *IGFBPL1* | 6,64 | 0,000900016 |
| *ENSG00000197479* | *PCDHB11* | 6,61 | 9,77354E-07 |
| *ENSG00000204335* | *SP5* | 6,58 | 7,44075E-05 |
| *ENSG00000159217* | *IGF2BP1* | 6,57 | 0,000641436 |
| *ENSG00000171956* | *FOXB1* | 6,55 | 0,00481982 |
| *ENSG00000165215* | *CLDN3* | 6,53 | 0,001421351 |
| *ENSG00000139515* | *PDX1* | 6,53 | 0,007091247 |
| *ENSG00000138650* | *PCDH10* | 6,52 | 0,0014053 |
| *ENSG00000126778* | *SIX1* | 6,48 | 4,33447E-12 |
| *ENSG00000152580* | *IGSF10* | 6,47 | 6,02013E-12 |
| *ENSG00000152785* | *BMP3* | 6,46 | 0,013160302 |
| *ENSG00000145526* | *CDH18* | 6,45 | 0,00140907 |
| *ENSG00000075340* | *ADD2* | 6,42 | 0,001463929 |
| *ENSG00000137834* | *SMAD6* | 6,39 | 1,92952E-09 |
| *ENSG00000162654* | *GBP4* | 6,37 | 1,10873E-05 |
| *ENSG00000165685* | *TMEM52B* | 6,33 | 5,51492E-06 |
| *ENSG00000275793* | *RIMBP3* | 6,31 | 0,000101276 |
| *ENSG00000124256* | *ZBP1* | 6,29 | 0,009819197 |
| *ENSG00000129682* | *FGF13* | 6,27 | 0,00268043 |
| *ENSG00000120322* | *PCDHB8* | 6,26 | 0,000326526 |
| *ENSG00000187689* | *AMTN* | 6,25 | 4,61408E-05 |
| *ENSG00000118785* | *SPP1* | 6,23 | 0,001392486 |
| *ENSG00000132182* | *NUP210* | 6,23 | 1,97969E-05 |
| *ENSG00000115468* | *EFHD1* | 6,21 | 0,001783709 |
| *ENSG00000128709* | *HOXD9* | 6,16 | 2,04631E-06 |
| *ENSG00000095596* | *CYP26A1* | 6,13 | 0,012393593 |
| *ENSG00000184845* | *DRD1* | 6,12 | 0,03613826 |
| *ENSG00000105894* | *PTN* | 6,11 | 0,000559626 |
| *ENSG00000257138* | *TAS2R38* | 6,10 | 0,000823404 |
| *ENSG00000176165* | *FOXG1* | 6,06 | 0,00277233 |
| *ENSG00000162711* | *NLRP3* | 6,06 | 6,39878E-05 |

| | | | |
|---|---|---|---|
| *ENSG00000196335* | *STK31* | 6,04 | 0,00077714 |
| *ENSG00000128645* | *HOXD1* | 6,04 | 0,001047776 |
| *ENSG00000172322* | *CLEC12A* | 6,03 | 0,038220635 |
| *ENSG00000182901* | *RGS7* | 6,00 | 0,000649022 |
| *ENSG00000185686* | *PRAME* | 6,00 | 0,012445414 |
| *ENSG00000185758* | *CLDN24* | 5,99 | 0,001837674 |
| *ENSG00000111700* | *SLCO1B3* | 5,98 | 0,040472039 |
| *ENSG00000043355* | *ZIC2* | 5,98 | 0,002434295 |
| *ENSG00000136267* | *DGKB* | 5,97 | 0,014635534 |
| *ENSG00000273706* | *LHX1* | 5,97 | 0,00123908 |
| *ENSG00000119547* | *ONECUT2* | 5,96 | 2,04836E-05 |
| *ENSG00000128567* | *PODXL* | 5,95 | 2,42271E-10 |
| *ENSG00000163499* | *CRYBA2* | 5,95 | 0,008133178 |
| *ENSG00000182732* | *RGS6* | 5,94 | 0,006097028 |
| *ENSG00000145451* | *GLRA3* | 5,93 | 0,021607033 |
| *ENSG00000144229* | *THSD7B* | 5,92 | 0,000542516 |
| *ENSG00000206531* | *CD200R1L* | 5,90 | 0,037985481 |
| *ENSG00000198959* | *TGM2* | 5,88 | 0,001780371 |
| *ENSG00000213512* | *GBP7* | 5,87 | 0,00092882 |
| *ENSG00000170577* | *SIX2* | 5,86 | 0,003177973 |
| *ENSG00000139800* | *ZIC5* | 5,86 | 0,013485635 |
| *ENSG00000152804* | *HHEX* | 5,84 | 0,000196842 |
| *ENSG00000164749* | *HNF4G* | 5,83 | 0,000400867 |
| *ENSG00000184012* | *TMPRSS2* | 5,83 | 0,005981511 |
| *ENSG00000205359* | *SLCO6A1* | 5,81 | 0,01464648 |
| *ENSG00000064692* | *SNCAIP* | 5,79 | 0,000921546 |
| *ENSG00000175877* | *TMEM270* | 5,77 | 0,000228337 |
| *ENSG00000186732* | *MPPED1* | 5,76 | 0,005815059 |
| *ENSG00000205517* | *RGL3* | 5,72 | 0,001376677 |
| *ENSG00000109625* | *CPZ* | 5,72 | 0,001531099 |
| *ENSG00000160183* | *TMPRSS3* | 5,71 | 0,000911334 |
| *ENSG00000148965* | *SAA4* | 5,71 | 0,013920676 |
| *ENSG00000153253* | *SCN3A* | 5,70 | 0,000445061 |
| *ENSG00000204287* | *HLA-DRA* | 5,69 | 0,00222641 |
| *ENSG00000189052* | *CGB5* | 5,67 | 0,018839808 |
| *ENSG00000197142* | *ACSL5* | 5,66 | 6,51162E-06 |
| *ENSG00000146955* | *RAB19* | 5,65 | 0,004031257 |
| *ENSG00000169994* | *MYO7B* | 5,63 | 0,010494527 |
| *ENSG00000198626* | *RYR2* | 5,61 | 1,48009E-05 |
| *ENSG00000261787* | *TCF24* | 5,61 | 3,06289E-05 |
| *ENSG00000019505* | *SYT13* | 5,59 | 0,026806251 |
| *ENSG00000011677* | *GABRA3* | 5,59 | 0,004876664 |
| *ENSG00000170558* | *CDH2* | 5,55 | 0,00086085 |
| *ENSG00000145536* | *ADAMTS16* | 5,53 | 0,000370381 |
| *ENSG00000110848* | *CD69* | 5,52 | 0,010450912 |

| | | | |
|---|---|---|---|
| *ENSG00000120645* | *IQSEC3* | 5,49 | 0,010559047 |
| *ENSG00000213030* | *CGB8* | 5,48 | 0,01134914 |
| *ENSG00000012779* | *ALOX5* | 5,48 | 0,007593681 |
| *ENSG00000189056* | *RELN* | 5,46 | 1,32799E-05 |
| *ENSG00000126895* | *AVPR2* | 5,46 | 0,006866653 |
| *ENSG00000155849* | *ELMO1* | 5,45 | 0,00820742 |
| *ENSG00000116833* | *NR5A2* | 5,44 | 0,000450856 |
| *ENSG00000167889* | *MGAT5B* | 5,44 | 2,42271E-10 |
| *ENSG00000158528* | *PPP1R9A* | 5,41 | 0,004812881 |
| *ENSG00000178568* | *ERBB4* | 5,37 | 0,002548684 |
| *ENSG00000173253* | *DMRT2* | 5,36 | 0,020710296 |
| *ENSG00000179776* | *CDH5* | 5,36 | 0,03451553 |
| *ENSG00000149926* | *FAM57B* | 5,33 | 0,001764771 |
| *ENSG00000177839* | *PCDHB9* | 5,33 | 1,8613E-05 |
| *ENSG00000128266* | *GNAZ* | 5,29 | 0,000761608 |
| *ENSG00000079841* | *RIMS1* | 5,29 | 0,017498711 |
| *ENSG00000130303* | *BST2* | 5,29 | 0,00153476 |
| *ENSG00000165071* | *TMEM71* | 5,29 | 0,0005652 |
| *ENSG00000182759* | *MAFA* | 5,28 | 0,002642565 |
| *ENSG00000023171* | *GRAMD1B* | 5,27 | 0,0093958 |
| *ENSG00000124657* | *OR2B6* | 5,24 | 0,00181694 |
| *ENSG00000164690* | *SHH* | 5,24 | 0,010848815 |
| *ENSG00000146674* | *IGFBP3* | 5,23 | 0,002352012 |
| *ENSG00000204296* | *C6orf10* | 5,23 | 0,03983201 |
| *ENSG00000272674* | *PCDHB16* | 5,20 | 0,000244652 |
| *ENSG00000180777* | *ANKRD30B* | 5,17 | 0,044923153 |
| *ENSG00000180660* | *MAB21L1* | 5,14 | 0,031131113 |
| *ENSG00000103241* | *FOXF1* | 5,09 | 0,031523158 |
| *ENSG00000112333* | *NR2E1* | 5,08 | 0,006478447 |
| *ENSG00000125462* | *C1orf61* | 5,08 | 0,024790046 |
| *ENSG00000176485* | *PLA2G16* | 5,08 | 0,000733917 |
| *ENSG00000205809* | *KLRC2* | 5,05 | 0,014157914 |
| *ENSG00000005249* | *PRKAR2B* | 5,04 | 4,07051E-08 |
| *ENSG00000175294* | *CATSPER1* | 5,03 | 0,000736183 |
| *ENSG00000181143* | *MUC16* | 5,03 | 0,014423347 |
| *ENSG00000254221* | *PCDHGB1* | 5,02 | 0,000145216 |
| *ENSG00000058335* | *RASGRF1* | 5,02 | 4,18017E-07 |
| *ENSG00000145248* | *SLC10A4* | 4,99 | 0,017413456 |
| *ENSG00000146678* | *IGFBP1* | 4,97 | 0,010290132 |
| *ENSG00000065609* | *SNAP91* | 4,93 | 0,049995155 |
| *ENSG00000172461* | *FUT9* | 4,93 | 0,044548287 |
| *ENSG00000099994* | *SUSD2* | 4,91 | 0,000921546 |
| *ENSG00000198944* | *SOWAHA* | 4,91 | 0,011770314 |
| *ENSG00000187912* | *CLEC17A* | 4,91 | 0,03220846 |
| *ENSG00000145198* | *VWA5B2* | 4,91 | 0,001270795 |

| | | | |
|---|---|---|---|
| *ENSG00000121797* | *CCRL2* | 4,89 | 0,004985654 |
| *ENSG00000120324* | *PCDHB10* | 4,88 | 9,92192E-05 |
| *ENSG00000164176* | *EDIL3* | 4,87 | 0,002772108 |
| *ENSG00000162687* | *KCNT2* | 4,86 | 0,043179377 |
| *ENSG00000038945* | *MSR1* | 4,83 | 0,029719045 |
| *ENSG00000132026* | *RTBDN* | 4,83 | 0,029980956 |
| *ENSG00000049249* | *TNFRSF9* | 4,82 | 7,17442E-06 |
| *ENSG00000189108* | *IL1RAPL2* | 4,82 | 0,013071654 |
| *ENSG00000176381* | *PRR18* | 4,81 | 0,047683733 |
| *ENSG00000106852* | *LHX6* | 4,81 | 0,000107746 |
| *ENSG00000174827* | *PDZK1* | 4,81 | 0,006710513 |
| *ENSG00000174963* | *ZIC4* | 4,80 | 0,001660035 |
| *ENSG00000162722* | *TRIM58* | 4,80 | 0,000336133 |
| *ENSG00000171631* | *P2RY6* | 4,79 | 2,92819E-05 |
| *ENSG00000187372* | *PCDHB13* | 4,78 | 8,17489E-05 |
| *ENSG00000198125* | *MB* | 4,78 | 0,034167547 |
| *ENSG00000134339* | *SAA2* | 4,78 | 0,01674103 |
| *ENSG00000172915* | *NBEA* | 4,78 | 0,000144481 |
| *ENSG00000152977* | *ZIC1* | 4,78 | 0,004629837 |
| *ENSG00000165238* | *WNK2* | 4,77 | 0,012759946 |
| *ENSG00000258881* | | 4,76 | 0,017991657 |
| *ENSG00000156206* | *CFAP161* | 4,74 | 0,035810112 |
| *ENSG00000149021* | *SCGB1A1* | 4,73 | 0,049218739 |
| *ENSG00000148704* | *VAX1* | 4,73 | 0,00012229 |
| *ENSG00000137273* | *FOXF2* | 4,73 | 0,000865732 |
| *ENSG00000096996* | *IL12RB1* | 4,72 | 0,001533009 |
| *ENSG00000172159* | *FRMD3* | 4,72 | 0,00170765 |
| *ENSG00000139865* | *TTC6* | 4,72 | 0,001692221 |
| *ENSG00000053918* | *KCNQ1* | 4,71 | 0,016459791 |
| *ENSG00000177462* | *OR2T8* | 4,69 | 0,046628677 |
| *ENSG00000144227* | *NXPH2* | 4,68 | 0,026530022 |
| *ENSG00000152430* | *BOLL* | 4,68 | 0,00114902 |
| *ENSG00000112852* | *PCDHB2* | 4,67 | 0,001220675 |
| *ENSG00000198846* | *TOX* | 4,67 | 0,000223669 |
| *ENSG00000196092* | *PAX5* | 4,65 | 0,006235064 |
| *ENSG00000168685* | *IL7R* | 4,64 | 0,000623977 |
| *ENSG00000143512* | *HHIPL2* | 4,64 | 0,00476491 |
| *ENSG00000124191* | *TOX2* | 4,62 | 0,00054942 |
| *ENSG00000168754* | *FAM178B* | 4,59 | 0,001000722 |
| *ENSG00000169126* | *ARMC4* | 4,58 | 0,014957584 |
| *ENSG00000129514* | *FOXA1* | 4,57 | 0,00326163 |
| *ENSG00000080224* | *EPHA6* | 4,56 | 0,016277722 |
| *ENSG00000186047* | *DLEU7* | 4,55 | 0,000530614 |
| *ENSG00000121075* | *TBX4* | 4,54 | 0,007841301 |
| *ENSG00000169418* | *NPR1* | 4,54 | 0,004876664 |

| | | | |
|---|---|---|---|
| *ENSG00000186998* | *EMID1* | 4,52 | 0,023923289 |
| *ENSG00000178965* | *ERICH3* | 4,52 | 0,038209528 |
| *ENSG00000077092* | *RARB* | 4,51 | 0,014904445 |
| *ENSG00000164778* | *EN2* | 4,49 | 0,000109316 |
| *ENSG00000142185* | *TRPM2* | 4,49 | 1,04922E-06 |
| *ENSG00000120251* | *GRIA2* | 4,48 | 0,024844178 |
| *ENSG00000164744* | *SUN3* | 4,48 | 0,001201695 |
| *ENSG00000115194* | *SLC30A3* | 4,48 | 0,033914468 |
| *ENSG00000205045* | *SLFN12L* | 4,47 | 1,31537E-05 |
| *ENSG00000105246* | *EBI3* | 4,47 | 0,015373376 |
| *ENSG00000182177* | *ASB18* | 4,46 | 0,031171525 |
| *ENSG00000166450* | *PRTG* | 4,46 | 0,044366056 |
| *ENSG00000256980* | *KHDC1L* | 4,45 | 0,002516774 |
| *ENSG00000170801* | *HTRA3* | 4,45 | 0,017386537 |
| *ENSG00000168843* | *FSTL5* | 4,44 | 0,03870815 |
| *ENSG00000118004* | *COLEC11* | 4,42 | 0,048594321 |
| *ENSG00000236320* | *SLFN14* | 4,42 | 0,022990397 |
| *ENSG00000075213* | *SEMA3A* | 4,41 | 3,38688E-07 |
| *ENSG00000152192* | *POU4F1* | 4,40 | 0,003141529 |
| *ENSG00000029534* | *ANK1* | 4,39 | 0,00077714 |
| *ENSG00000166448* | *TMEM130* | 4,39 | 0,016598168 |
| *ENSG00000156886* | *ITGAD* | 4,37 | 0,032767078 |
| *ENSG00000170166* | *HOXD4* | 4,36 | 0,001234014 |
| *ENSG00000125355* | *TMEM255A* | 4,36 | 0,004071419 |
| *ENSG00000156966* | *B3GNT7* | 4,34 | 0,00076408 |
| *ENSG00000002726* | *AOC1* | 4,34 | 0,016598168 |
| *ENSG00000130829* | *DUSP9* | 4,34 | 0,004378582 |
| *ENSG00000180730* | *SHISA2* | 4,34 | 0,001138149 |
| *ENSG00000157303* | *SUSD3* | 4,33 | 0,001107597 |
| *ENSG00000163121* | *NEURL3* | 4,32 | 0,001370503 |
| *ENSG00000134198* | *TSPAN2* | 4,30 | 0,001267809 |
| *ENSG00000274267* | *HIST1H3B* | 4,30 | 0,009387139 |
| *ENSG00000253910* | *PCDHGB2* | 4,30 | 1,51019E-05 |
| *ENSG00000146938* | *NLGN4X* | 4,29 | 0,032844012 |
| *ENSG00000124507* | *PACSIN1* | 4,29 | 0,021510428 |
| *ENSG00000005073* | *HOXA11* | 4,28 | 2,83068E-10 |
| *ENSG00000137261* | *KIAA0319* | 4,28 | 0,004221593 |
| *ENSG00000121005* | *CRISPLD1* | 4,26 | 0,038188577 |
| *ENSG00000271503* | *CCL5* | 4,26 | 0,014964508 |
| *ENSG00000175164* | *ABO* | 4,23 | 0,023027472 |
| *ENSG00000100055* | *CYTH4* | 4,23 | 0,005405555 |
| *ENSG00000184985* | *SORCS2* | 4,22 | 0,001061946 |
| *ENSG00000170214* | *ADRA1B* | 4,22 | 0,000292819 |
| *ENSG00000178401* | *DNAJC22* | 4,21 | 7,28526E-07 |
| *ENSG00000113212* | *PCDHB7* | 4,21 | 0,025922411 |

| | | | |
|---|---|---|---|
| *ENSG00000176244* | *ACBD7* | 4,19 | 2,33696E-05 |
| *ENSG00000139055* | *ERP27* | 4,18 | 0,042141111 |
| *ENSG00000182103* | *FAM181B* | 4,15 | 0,033813187 |
| *ENSG00000007372* | *PAX6* | 4,14 | 2,87926E-07 |
| *ENSG00000023445* | *BIRC3* | 4,14 | 0,000355892 |
| *ENSG00000128284* | *APOL3* | 4,14 | 0,002196585 |
| *ENSG00000139973* | *SYT16* | 4,12 | 0,000762304 |
| *ENSG00000136167* | *LCP1* | 4,11 | 0,044296742 |
| *ENSG00000148053* | *NTRK2* | 4,11 | 0,01836873 |
| *ENSG00000120068* | *HOXB8* | 4,09 | 0,013365525 |
| *ENSG00000185215* | *TNFAIP2* | 4,08 | 0,001057491 |
| *ENSG00000238243* | *OR2W3* | 4,08 | 0,003331431 |
| *ENSG00000197977* | *ELOVL2* | 4,06 | 0,036625673 |
| *ENSG00000164530* | *PI16* | 4,05 | 0,047717215 |
| *ENSG00000004468* | *CD38* | 4,04 | 0,004953444 |
| *ENSG00000105464* | *GRIN2D* | 4,04 | 0,001573307 |
| *ENSG00000170962* | *PDGFD* | 4,04 | 0,032874637 |
| *ENSG00000131094* | *C1QL1* | 4,04 | 0,001531099 |
| *ENSG00000159263* | *SIM2* | 4,04 | 0,000324564 |
| *ENSG00000115896* | *PLCL1* | 4,03 | 0,0053174 |
| *ENSG00000205221* | *VIT* | 4,01 | 0,029184801 |
| *ENSG00000167680* | *SEMA6B* | 3,99 | 0,002564273 |
| *ENSG00000151702* | *FLI1* | 3,98 | 0,015786369 |
| *ENSG00000006534* | *ALDH3B1* | 3,97 | 0,000921546 |
| *ENSG00000135454* | *B4GALNT1* | 3,97 | 0,002635361 |
| *ENSG00000226690* | | 3,94 | 0,029349841 |
| *ENSG00000197181* | *PIWIL2* | 3,93 | 0,017119935 |
| *ENSG00000148677* | *ANKRD1* | 3,93 | 0,016066611 |
| *ENSG00000105967* | *TFEC* | 3,93 | 0,048412155 |
| *ENSG00000205426* | *KRT81* | 3,92 | 0,017991657 |
| *ENSG00000147573* | *TRIM55* | 3,91 | 0,036048308 |
| *ENSG00000134780* | *DAGLA* | 3,90 | 0,001901514 |
| *ENSG00000144230* | *GPR17* | 3,89 | 0,033802021 |
| *ENSG00000133321* | *RARRES3* | 3,86 | 0,006102171 |
| *ENSG00000083454* | *P2RX5* | 3,86 | 0,007325256 |
| *ENSG00000090339* | *ICAM1* | 3,86 | 0,000331144 |
| *ENSG00000175556* | *LONRF3* | 3,85 | 0,021696882 |
| *ENSG00000182621* | *PLCB1* | 3,84 | 0,002410169 |
| *ENSG00000143195* | *ILDR2* | 3,84 | 0,019441679 |
| *ENSG00000168961* | *LGALS9* | 3,84 | 0,002238906 |
| *ENSG00000180914* | *OXTR* | 3,84 | 0,006105785 |
| *ENSG00000183833* | *MAATS1* | 3,83 | 0,032005099 |
| *ENSG00000138075* | *ABCG5* | 3,83 | 0,035954201 |
| *ENSG00000183729* | *NPBWR1* | 3,82 | 0,007593681 |
| *ENSG00000165124* | *SVEP1* | 3,82 | 0,000881353 |

| | | | |
|---|---|---|---|
| *ENSG00000019186* | *CYP24A1* | 3,82 | 0,029976436 |
| *ENSG00000119125* | *GDA* | 3,81 | 0,024138581 |
| *ENSG00000168959* | *GRM5* | 3,80 | 0,019423321 |
| *ENSG00000198771* | *RCSD1* | 3,79 | 0,019258062 |
| *ENSG00000121380* | *BCL2L14* | 3,77 | 0,008705254 |
| *ENSG00000164308* | *ERAP2* | 3,75 | 0,000216724 |
| *ENSG00000179869* | *ABCA13* | 3,74 | 0,005891465 |
| *ENSG00000080031* | *PTPRH* | 3,74 | 0,005457116 |
| *ENSG00000120327* | *PCDHB14* | 3,73 | 5,08434E-06 |
| *ENSG00000178150* | *ZNF114* | 3,71 | 1,4987E-07 |
| *ENSG00000185090* | *MANEAL* | 3,70 | 3,5755E-09 |
| *ENSG00000102271* | *KLHL4* | 3,69 | 0,035049712 |
| *ENSG00000019582* | *CD74* | 3,68 | 0,037457049 |
| *ENSG00000164292* | *RHOBTB3* | 3,66 | 8,24118E-05 |
| *ENSG00000044524* | *EPHA3* | 3,66 | 0,033132992 |
| *ENSG00000164161* | *HHIP* | 3,65 | 0,047521805 |
| *ENSG00000104368* | *PLAT* | 3,65 | 0,007617481 |
| *ENSG00000106025* | *TSPAN12* | 3,62 | 0,003165071 |
| *ENSG00000168702* | *LRP1B* | 3,61 | 0,036943016 |
| *ENSG00000215018* | *COL28A1* | 3,61 | 0,030702287 |
| *ENSG00000133216* | *EPHB2* | 3,60 | 7,28828E-05 |
| *ENSG00000204176* | *SYT15* | 3,60 | 0,000150122 |
| *ENSG00000187867* | *PALM3* | 3,60 | 0,020322371 |
| *ENSG00000278463* | *HIST1H2AB* | 3,60 | 0,047030563 |
| *ENSG00000182253* | *SYNM* | 3,58 | 1,80541E-06 |
| *ENSG00000100628* | *ASB2* | 3,57 | 0,021230933 |
| *ENSG00000167191* | *GPRC5B* | 3,57 | 0,009319186 |
| *ENSG00000130675* | *MNX1* | 3,56 | 0,030162258 |
| *ENSG00000149564* | *ESAM* | 3,55 | 0,007056394 |
| *ENSG00000152078* | *TMEM56* | 3,52 | 1,80541E-06 |
| *ENSG00000156509* | *FBXO43* | 3,50 | 6,66691E-05 |
| *ENSG00000146555* | *SDK1* | 3,50 | 0,000660749 |
| *ENSG00000133488* | *SEC14L4* | 3,49 | 0,023287998 |
| *ENSG00000277758* | | 3,48 | 0,000473979 |
| *ENSG00000242265* | *PEG10* | 3,46 | 0,000234383 |
| *ENSG00000234602* | *MCIDAS* | 3,46 | 8,19962E-06 |
| *ENSG00000163449* | *TMEM169* | 3,46 | 0,017302395 |
| *ENSG00000136327* | *NKX2-8* | 3,45 | 0,021680644 |
| *ENSG00000135502* | *SLC26A10* | 3,44 | 0,005553022 |
| *ENSG00000250305* | *TRMT9B* | 3,43 | 0,046951421 |
| *ENSG00000177519* | *RPRM* | 3,42 | 0,041628765 |
| *ENSG00000128833* | *MYO5C* | 3,42 | 0,009198387 |
| *ENSG00000043143* | *JADE2* | 3,41 | 2,64865E-05 |
| *ENSG00000063015* | *SEZ6* | 3,41 | 0,01431288 |
| *ENSG00000127990* | *SGCE* | 3,40 | 0,000220894 |

| | | | |
|---|---|---|---|
| *ENSG00000130045* | *NXNL2* | 3,40 | 0,004226828 |
| *ENSG00000262576* | *PCDHGA4* | 3,39 | 0,000295192 |
| *ENSG00000221890* | *NPTXR* | 3,39 | 0,004698533 |
| *ENSG00000102452* | *NALCN* | 3,39 | 0,015915523 |
| *ENSG00000151023* | *ENKUR* | 3,37 | 0,000487487 |
| *ENSG00000168505* | *GBX2* | 3,37 | 0,025688045 |
| *ENSG00000137869* | *CYP19A1* | 3,35 | 0,000433442 |
| *ENSG00000177721* | *ANXA2R* | 3,34 | 1,4757E-11 |
| *ENSG00000141574* | *SECTM1* | 3,34 | 0,000295536 |
| *ENSG00000123146* | *ADGRE5* | 3,33 | 1,3383E-07 |
| *ENSG00000091656* | *ZFHX4* | 3,30 | 0,001166981 |
| *ENSG00000039600* | *SOX30* | 3,30 | 0,004378108 |
| *ENSG00000168913* | *ENHO* | 3,30 | 0,034029106 |
| *ENSG00000153993* | *SEMA3D* | 3,29 | 0,018894833 |
| *ENSG00000188404* | *SELL* | 3,29 | 0,040156349 |
| *ENSG00000058866* | *DGKG* | 3,29 | 0,030642628 |
| *ENSG00000102048* | *ASB9* | 3,28 | 0,004031257 |
| *ENSG00000112761* | *WISP3* | 3,28 | 0,002340307 |
| *ENSG00000267060* | *PTGES3L* | 3,28 | 5,75626E-05 |
| *ENSG00000154146* | *NRGN* | 3,27 | 0,008380258 |
| *ENSG00000184058* | *TBX1* | 3,26 | 0,003136739 |
| *ENSG00000186193* | *SAPCD2* | 3,24 | 7,40971E-08 |
| *ENSG00000167034* | *NKX3-1* | 3,24 | 1,77015E-07 |
| *ENSG00000135119* | *RNFT2* | 3,22 | 5,69819E-06 |
| *ENSG00000157927* | *RADIL* | 3,21 | 0,036134966 |
| *ENSG00000198246* | *SLC29A3* | 3,21 | 0,042323343 |
| *ENSG00000101306* | *MYLK2* | 3,21 | 0,000141878 |
| *ENSG00000130720* | *FIBCD1* | 3,21 | 0,041079294 |
| *ENSG00000174015* | *SPERT* | 3,20 | 0,025681808 |
| *ENSG00000121895* | *TMEM156* | 3,20 | 0,044210903 |
| *ENSG00000176595* | *KBTBD11* | 3,19 | 0,00125808 |
| *ENSG00000162068* | *NTN3* | 3,19 | 0,016747977 |
| *ENSG00000186603* | *HPDL* | 3,18 | 0,004120155 |
| *ENSG00000152779* | *SLC16A12* | 3,18 | 0,026523291 |
| *ENSG00000188763* | *FZD9* | 3,18 | 0,009037944 |
| *ENSG00000100342* | *APOL1* | 3,18 | 0,011415918 |
| *ENSG00000128342* | *LIF* | 3,18 | 0,015689012 |
| *ENSG00000124839* | *RAB17* | 3,18 | 0,018418646 |
| *ENSG00000092929* | *UNC13D* | 3,18 | 0,001709821 |
| *ENSG00000160255* | *ITGB2* | 3,18 | 0,012436536 |
| *ENSG00000164430* | *CGAS* | 3,17 | 1,55474E-06 |
| *ENSG00000166823* | *MESP1* | 3,17 | 0,013710563 |
| *ENSG00000082074* | *FYB1* | 3,15 | 0,02753681 |
| *ENSG00000141448* | *GATA6* | 3,14 | 0,000427047 |
| *ENSG00000260027* | *HOXB7* | 3,14 | 0,000409342 |

| | | | |
|---|---|---|---|
| *ENSG00000150551* | *LYPD1* | 3,14 | 0,000796218 |
| *ENSG00000072163* | *LIMS2* | 3,14 | 0,020207646 |
| *ENSG00000039139* | *DNAH5* | 3,13 | 0,001531099 |
| *ENSG00000204956* | *PCDHGA1* | 3,12 | 0,014787255 |
| *ENSG00000188641* | *DPYD* | 3,11 | 0,000331144 |
| *ENSG00000168454* | *TXNDC2* | 3,11 | 0,038015111 |
| *ENSG00000104848* | *KCNA7* | 3,10 | 0,00142262 |
| *ENSG00000183023* | *SLC8A1* | 3,09 | 0,008684368 |
| *ENSG00000063438* | *AHRR* | 3,09 | 0,018958922 |
| *ENSG00000160161* | *CILP2* | 3,08 | 0,008843639 |
| *ENSG00000012171* | *SEMA3B* | 3,08 | 0,040449719 |
| *ENSG00000124772* | *CPNE5* | 3,07 | 0,031075523 |
| *ENSG00000123080* | *CDKN2C* | 3,06 | 3,0877E-07 |
| *ENSG00000065325* | *GLP2R* | 3,05 | 0,043873653 |
| *ENSG00000074370* | *ATP2A3* | 3,04 | 0,030141923 |
| *ENSG00000274641* | *HIST1H2BO* | 3,03 | 0,021230933 |
| *ENSG00000115919* | *KYNU* | 3,03 | 0,035829762 |
| *ENSG00000130948* | *HSD17B3* | 3,02 | 0,005822314 |
| *ENSG00000138336* | *TET1* | 3,02 | 0,00014725 |
| *ENSG00000173890* | *GPR160* | 3,01 | 2,63985E-06 |
| *ENSG00000130038* | *CRACR2A* | 3,00 | 0,000847907 |
| *ENSG00000135083* | *CCNJL* | 2,99 | 0,00078261 |
| *ENSG00000088367* | *EPB41L1* | 2,99 | 0,024328955 |
| *ENSG00000136205* | *TNS3* | 2,99 | 2,63754E-08 |
| *ENSG00000138411* | *HECW2* | 2,98 | 0,048984364 |
| *ENSG00000173227* | *SYT12* | 2,98 | 0,012572684 |
| *ENSG00000100867* | *DHRS2* | 2,98 | 0,002617814 |
| *ENSG00000183638* | *RP1L1* | 2,98 | 0,016630957 |
| *ENSG00000198829* | *SUCNR1* | 2,98 | 0,041190265 |
| *ENSG00000138347* | *MYPN* | 2,98 | 0,043533225 |
| *ENSG00000163888* | *CAMK2N2* | 2,97 | 0,001691659 |
| *ENSG00000108511* | *HOXB6* | 2,97 | 0,017828898 |
| *ENSG00000261308* | *FIGNL2* | 2,96 | 0,026253191 |
| *ENSG00000166897* | *ELFN2* | 2,96 | 0,015820988 |
| *ENSG00000183873* | *SCN5A* | 2,94 | 0,033153026 |
| *ENSG00000145861* | *C1QTNF2* | 2,93 | 0,005630051 |
| *ENSG00000163131* | *CTSS* | 2,92 | 0,048030845 |
| *ENSG00000183688* | *RFLNB* | 2,91 | 0,014711815 |
| *ENSG00000101986* | *ABCD1* | 2,91 | 0,000390753 |
| *ENSG00000140323* | *DISP2* | 2,90 | 0,000324565 |
| *ENSG00000136999* | *NOV* | 2,90 | 0,025244771 |
| *ENSG00000157833* | *GAREM2* | 2,89 | 0,015317383 |
| *ENSG00000165730* | *STOX1* | 2,88 | 0,000459878 |
| *ENSG00000006377* | *DLX6* | 2,88 | 0,026316076 |
| *ENSG00000106484* | *MEST* | 2,88 | 0,015800139 |

| | | | |
|---|---|---|---|
| *ENSG00000273983* | *HIST1H3G* | 2,87 | 0,001947213 |
| *ENSG00000137825* | *ITPKA* | 2,87 | 7,36423E-05 |
| *ENSG00000149177* | *PTPRJ* | 2,86 | 0,000190436 |
| *ENSG00000100276* | *RASL10A* | 2,85 | 0,037188882 |
| *ENSG00000008277* | *ADAM22* | 2,85 | 0,025724901 |
| *ENSG00000139438* | *FAM222A* | 2,83 | 0,000884179 |
| *ENSG00000175352* | *NRIP3* | 2,83 | 0,00766004 |
| *ENSG00000175920* | *DOK7* | 2,82 | 0,043108098 |
| *ENSG00000165716* | *FAM69B* | 2,82 | 0,012755177 |
| *ENSG00000253485* | *PCDHGA5* | 2,82 | 3,3721E-05 |
| *ENSG00000152377* | *SPOCK1* | 2,81 | 0,001116385 |
| *ENSG00000123143* | *PKN1* | 2,80 | 0,023613811 |
| *ENSG00000158683* | *PKD1L1* | 2,80 | 0,003763597 |
| *ENSG00000162591* | *MEGF6* | 2,80 | 0,048814386 |
| *ENSG00000110446* | *SLC15A3* | 2,79 | 0,031455558 |
| *ENSG00000151491* | *EPS8* | 2,77 | 0,014518978 |
| *ENSG00000162639* | *HENMT1* | 2,77 | 4,3112E-07 |
| *ENSG00000182168* | *UNC5C* | 2,76 | 0,017491886 |
| *ENSG00000176678* | *FOXL1* | 2,75 | 0,000885545 |
| *ENSG00000111110* | *PPM1H* | 2,75 | 0,044210903 |
| *ENSG00000013588* | *GPRC5A* | 2,75 | 0,029247581 |
| *ENSG00000182791* | *CCDC87* | 2,74 | 0,00178724 |
| *ENSG00000182578* | *CSF1R* | 2,73 | 0,033274006 |
| *ENSG00000169946* | *ZFPM2* | 2,72 | 0,000753097 |
| *ENSG00000105929* | *ATP6V0A4* | 2,71 | 0,030747882 |
| *ENSG00000189366* | *ALG1L* | 2,71 | 0,005667388 |
| *ENSG00000234776* | *C11orf94* | 2,71 | 0,019288314 |
| *ENSG00000136514* | *RTP4* | 2,70 | 0,046284288 |
| *ENSG00000168280* | *KIF5C* | 2,70 | 0,027542795 |
| *ENSG00000167780* | *SOAT2* | 2,70 | 0,007563295 |
| *ENSG00000233198* | *RNF224* | 2,69 | 0,009817505 |
| *ENSG00000167619* | *TMEM145* | 2,68 | 0,026557631 |
| *ENSG00000112984* | *KIF20A* | 2,68 | 2,06346E-05 |
| *ENSG00000116039* | *ATP6V1B1* | 2,67 | 0,033169749 |
| *ENSG00000177706* | *FAM20C* | 2,66 | 0,007809823 |
| *ENSG00000138193* | *PLCE1* | 2,65 | 0,011867634 |
| *ENSG00000168993* | *CPLX1* | 2,64 | 0,047653064 |
| *ENSG00000099282* | *TSPAN15* | 2,64 | 0,022700939 |
| *ENSG00000179841* | *AKAP5* | 2,64 | 0,00096282 |
| *ENSG00000177675* | *CD163L1* | 2,64 | 0,015050201 |
| *ENSG00000153904* | *DDAH1* | 2,63 | 0,000730274 |
| *ENSG00000154479* | *CCDC173* | 2,62 | 0,010826066 |
| *ENSG00000125347* | *IRF1* | 2,62 | 0,001481436 |
| *ENSG00000162415* | *ZSWIM5* | 2,61 | 0,038654124 |
| *ENSG00000113368* | *LMNB1* | 2,61 | 0,000205147 |

| | | | |
|---|---|---|---|
| *ENSG00000168811* | *IL12A* | 2,61 | 0,049365563 |
| *ENSG00000175463* | *TBC1D10C* | 2,61 | 0,000580902 |
| *ENSG00000165507* | *DEPP1* | 2,59 | 0,004581423 |
| *ENSG00000174514* | *MFSD4A* | 2,59 | 0,032214426 |
| *ENSG00000198807* | *PAX9* | 2,58 | 0,015647653 |
| *ENSG00000183840* | *GPR39* | 2,56 | 0,002455963 |
| *ENSG00000051523* | *CYBA* | 2,54 | 0,000349081 |
| *ENSG00000113319* | *RASGRF2* | 2,53 | 0,02540368 |
| *ENSG00000198929* | *NOS1AP* | 2,53 | 0,003181824 |
| *ENSG00000115738* | *ID2* | 2,52 | 0,038690757 |
| *ENSG00000104783* | *KCNN4* | 2,52 | 0,033292188 |
| *ENSG00000183150* | *GPR19* | 2,50 | 0,008632834 |
| *ENSG00000175697* | *GPR156* | 2,49 | 0,017398238 |
| *ENSG00000111348* | *ARHGDIB* | 2,48 | 0,028061375 |
| *ENSG00000239887* | *C1orf226* | 2,48 | 0,007141885 |
| *ENSG00000180998* | *GPR137C* | 2,47 | 0,000935495 |
| *ENSG00000198569* | *SLC34A3* | 2,45 | 0,02880328 |
| *ENSG00000130377* | *ACSBG2* | 2,45 | 0,005813603 |
| *ENSG00000196159* | *FAT4* | 2,45 | 0,026416833 |
| *ENSG00000184371* | *CSF1* | 2,44 | 0,024455574 |
| *ENSG00000113594* | *LIFR* | 2,44 | 0,005606486 |
| *ENSG00000173621* | *LRFN4* | 2,43 | 6,21245E-08 |
| *ENSG00000108813* | *DLX4* | 2,42 | 0,020700222 |
| *ENSG00000153064* | *BANK1* | 2,42 | 0,032707799 |
| *ENSG00000165181* | *C9orf84* | 2,42 | 3,24431E-05 |
| *ENSG00000117020* | *AKT3* | 2,41 | 0,006480356 |
| *ENSG00000169684* | *CHRNA5* | 2,40 | 0,005530435 |
| *ENSG00000004777* | *ARHGAP33* | 2,39 | 8,78453E-05 |
| *ENSG00000140993* | *TIGD7* | 2,39 | 0,0002603 |
| *ENSG00000107736* | *CDH23* | 2,38 | 0,018776478 |
| *ENSG00000107551* | *RASSF4* | 2,38 | 0,012990312 |
| *ENSG00000075218* | *GTSE1* | 2,38 | 5,63542E-06 |
| *ENSG00000166596* | *CFAP52* | 2,37 | 0,001421351 |
| *ENSG00000177842* | *ZNF620* | 2,37 | 0,001268456 |
| *ENSG00000162676* | *GFI1* | 2,36 | 0,031426421 |
| *ENSG00000128408* | *RIBC2* | 2,36 | 0,02732415 |
| *ENSG00000181418* | *DDN* | 2,35 | 0,047516117 |
| *ENSG00000119888* | *EPCAM* | 2,35 | 0,041221349 |
| *ENSG00000163053* | *SLC16A14* | 2,35 | 0,01650506 |
| *ENSG00000101412* | *E2F1* | 2,34 | 0,004102186 |
| *ENSG00000110492* | *MDK* | 2,34 | 0,037197044 |
| *ENSG00000158104* | *HPD* | 2,33 | 0,02937684 |
| *ENSG00000105613* | *MAST1* | 2,33 | 0,048676078 |
| *ENSG00000117791* | *MARC2* | 2,33 | 0,004289784 |
| *ENSG00000073464* | *CLCN4* | 2,33 | 0,008574466 |

| | | | |
|---|---|---|---|
| *ENSG00000104081* | *BMF* | 2,33 | 0,02546948 |
| *ENSG00000080986* | *NDC80* | 2,32 | 0,000450856 |
| *ENSG00000007968* | *E2F2* | 2,31 | 0,002195333 |
| *ENSG00000144959* | *NCEH1* | 2,31 | 0,000367084 |
| *ENSG00000249209* | | 2,30 | 2,08659E-06 |
| *ENSG00000117724* | *CENPF* | 2,29 | 8,00057E-05 |
| *ENSG00000127928* | *GNGT1* | 2,28 | 0,000923944 |
| *ENSG00000113248* | *PCDHB15* | 2,27 | 0,016906688 |
| *ENSG00000111859* | *NEDD9* | 2,27 | 0,029980956 |
| *ENSG00000088325* | *TPX2* | 2,26 | 3,53808E-06 |
| *ENSG00000165072* | *MAMDC2* | 2,26 | 0,022037495 |
| *ENSG00000153233* | *PTPRR* | 2,26 | 0,018227275 |
| *ENSG00000100206* | *DMC1* | 2,26 | 0,017263346 |
| *ENSG00000126562* | *WNK4* | 2,25 | 0,031079971 |
| *ENSG00000163171* | *CDC42EP3* | 2,25 | 0,044708858 |
| *ENSG00000116299* | *KIAA1324* | 2,25 | 0,029025336 |
| *ENSG00000142945* | *KIF2C* | 2,25 | 0,000110377 |
| *ENSG00000127329* | *PTPRB* | 2,25 | 0,019998739 |
| *ENSG00000105011* | *ASF1B* | 2,24 | 0,00116322 |
| *ENSG00000164304* | *CAGE1* | 2,23 | 0,004560355 |
| *ENSG00000275713* | *HIST1H2BH* | 2,23 | 0,000744711 |
| *ENSG00000164403* | *SHROOM1* | 2,23 | 2,34782E-05 |
| *ENSG00000188707* | *ZBED6CL* | 2,23 | 0,022907763 |
| *ENSG00000166278* | *C2* | 2,23 | 7,00886E-06 |
| *ENSG00000196074* | *SYCP2* | 2,22 | 0,022383596 |
| *ENSG00000164674* | *SYTL3* | 2,22 | 0,034878145 |
| *ENSG00000087586* | *AURKA* | 2,21 | 9,08656E-05 |
| *ENSG00000121653* | *MAPK8IP1* | 2,21 | 0,00427531 |
| *ENSG00000185697* | *MYBL1* | 2,20 | 0,020322371 |
| *ENSG00000146197* | *SCUBE3* | 2,20 | 0,004174767 |
| *ENSG00000111057* | *KRT18* | 2,20 | 0,008204637 |
| *ENSG00000117650* | *NEK2* | 2,20 | 0,000140671 |
| *ENSG00000124116* | *WFDC3* | 2,20 | 0,013286923 |
| *ENSG00000162946* | *DISC1* | 2,19 | 0,047766939 |
| *ENSG00000117318* | *ID3* | 2,19 | 0,015776524 |
| *ENSG00000111206* | *FOXM1* | 2,19 | 1,2464E-05 |
| *ENSG00000240065* | *PSMB9* | 2,17 | 0,036985543 |
| *ENSG00000138678* | *GPAT3* | 2,17 | 0,010998802 |
| *ENSG00000278828* | *HIST1H3H* | 2,15 | 0,049454437 |
| *ENSG00000188064* | *WNT7B* | 2,15 | 0,017627622 |
| *ENSG00000121621* | *KIF18A* | 2,15 | 1,26413E-05 |
| *ENSG00000148773* | *MKI67* | 2,15 | 2,98238E-05 |
| *ENSG00000114473* | *IQCG* | 2,14 | 0,042345599 |
| *ENSG00000136866* | *ZFP37* | 2,14 | 0,002102816 |
| *ENSG00000213626* | *LBH* | 2,13 | 0,006210431 |

| | | | |
|---|---|---|---|
| *ENSG00000107821* | *KAZALD1* | 2,13 | 0,000531964 |
| *ENSG00000090889* | *KIF4A* | 2,12 | 1,95266E-05 |
| *ENSG00000101057* | *MYBL2* | 2,12 | 0,026550015 |
| *ENSG00000134508* | *CABLES1* | 2,12 | 0,014100424 |
| *ENSG00000196787* | *HIST1H2AG* | 2,11 | 0,00845239 |
| *ENSG00000135451* | *TROAP* | 2,11 | 1,29096E-08 |
| *ENSG00000123358* | *NR4A1* | 2,11 | 0,008632834 |
| *ENSG00000166578* | *IQCD* | 2,11 | 0,001629709 |
| *ENSG00000138587* | *MNS1* | 2,10 | 0,019577407 |
| *ENSG00000105866* | *SP4* | 2,10 | 0,000355141 |
| *ENSG00000153208* | *MERTK* | 2,09 | 0,046485764 |
| *ENSG00000160886* | *LY6K* | 2,09 | 6,11868E-07 |
| *ENSG00000108106* | *UBE2S* | 2,09 | 0,003863065 |
| *ENSG00000076382* | *SPAG5* | 2,07 | 0,004421267 |
| *ENSG00000167779* | *IGFBP6* | 2,07 | 0,019132777 |
| *ENSG00000186185* | *KIF18B* | 2,06 | 0,000426715 |
| *ENSG00000082497* | *SERTAD4* | 2,06 | 0,001270248 |
| *ENSG00000164849* | *GPR146* | 2,06 | 0,037953463 |
| *ENSG00000122966* | *CIT* | 2,06 | 0,000174618 |
| *ENSG00000100526* | *CDKN3* | 2,06 | 0,000386339 |
| *ENSG00000143228* | *NUF2* | 2,06 | 0,005309758 |
| *ENSG00000166582* | *CENPV* | 2,05 | 0,000436471 |
| *ENSG00000126561* | *STAT5A* | 2,05 | 0,020207646 |
| *ENSG00000107105* | *ELAVL2* | 2,04 | 0,04325325 |
| *ENSG00000152253* | *SPC25* | 2,04 | 0,004221593 |
| *ENSG00000253731* | *PCDHGA6* | 2,04 | 0,005402252 |
| *ENSG00000117399* | *CDC20* | 2,04 | 0,000675713 |
| *ENSG00000065308* | *TRAM2* | 2,03 | 0,021168691 |
| *ENSG00000177465* | *ACOT4* | 2,03 | 0,004415292 |
| *ENSG00000101280* | *ANGPT4* | 2,03 | 0,048030845 |
| *ENSG00000111674* | *ENO2* | 2,03 | 0,034969226 |
| *ENSG00000105339* | *DENND3* | 2,02 | 0,000150749 |
| *ENSG00000121716* | *PILRB* | 2,02 | 0,034203538 |
| *ENSG00000086300* | *SNX10* | 2,02 | 0,014480296 |
| *ENSG00000163507* | *CIP2A* | 2,01 | 0,001559979 |
| *ENSG00000263513* | *FAM72C* | 2,01 | 0,030868968 |
| *ENSG00000169105* | *CHST14* | 2,00 | 0,000531918 |
| *ENSG00000182957* | *SPATA13* | 1,99 | 0,042821827 |
| *ENSG00000106976* | *DNM1* | 1,99 | 0,029189981 |
| *ENSG00000069011* | *PITX1* | 1,99 | 0,002081575 |
| *ENSG00000277203* | *F8A1* | 1,98 | 0,000631017 |
| *ENSG00000178878* | *APOLD1* | 1,98 | 0,004254812 |
| *ENSG00000221944* | *TIGD1* | 1,98 | 2,64236E-05 |
| *ENSG00000101447* | *FAM83D* | 1,97 | 0,007715004 |
| *ENSG00000008300* | *CELSR3* | 1,97 | 0,010016599 |

| | | | |
|---|---|---|---|
| *ENSG00000134690* | *CDCA8* | 1,97 | 0,000641436 |
| *ENSG00000099953* | *MMP11* | 1,96 | 0,034029042 |
| *ENSG00000215912* | *TTC34* | 1,96 | 0,005524227 |
| *ENSG00000072571* | *HMMR* | 1,96 | 0,021088823 |
| *ENSG00000158402* | *CDC25C* | 1,96 | 0,005815059 |
| *ENSG00000143653* | *SCCPDH* | 1,95 | 0,005665411 |
| *ENSG00000175063* | *UBE2C* | 1,95 | 0,0031072 |
| *ENSG00000073111* | *MCM2* | 1,95 | 0,014518978 |
| *ENSG00000162390* | *ACOT11* | 1,95 | 0,046182226 |
| *ENSG00000119397* | *CNTRL* | 1,95 | 2,86098E-05 |
| *ENSG00000179240* | *GVQW3* | 1,94 | 0,000818649 |
| *ENSG00000156802* | *ATAD2* | 1,94 | 0,001918723 |
| *ENSG00000137804* | *NUSAP1* | 1,94 | 0,000339653 |
| *ENSG00000255508* | | 1,93 | 0,000883546 |
| *ENSG00000105877* | *DNAH11* | 1,93 | 3,33096E-07 |
| *ENSG00000139354* | *GAS2L3* | 1,93 | 9,96712E-05 |
| *ENSG00000187210* | *GCNT1* | 1,93 | 0,032219488 |
| *ENSG00000111665* | *CDCA3* | 1,92 | 0,000502229 |
| *ENSG00000170525* | *PFKFB3* | 1,92 | 0,008632834 |
| *ENSG00000134222* | *PSRC1* | 1,92 | 0,00064473 |
| *ENSG00000110057* | *UNC93B1* | 1,92 | 0,002375868 |
| *ENSG00000172731* | *LRRC20* | 1,91 | 0,004039845 |
| *ENSG00000123485* | *HJURP* | 1,91 | 0,003774222 |
| *ENSG00000135476* | *ESPL1* | 1,91 | 0,000559626 |
| *ENSG00000138160* | *KIF11* | 1,90 | 0,000167998 |
| *ENSG00000136052* | *SLC41A2* | 1,90 | 0,025724901 |
| *ENSG00000102226* | *USP11* | 1,90 | 1,46214E-06 |
| *ENSG00000011426* | *ANLN* | 1,90 | 0,000312207 |
| *ENSG00000119403* | *PHF19* | 1,90 | 0,000900016 |
| *ENSG00000234616* | *JRK* | 1,89 | 0,010009174 |
| *ENSG00000197256* | *KANK2* | 1,89 | 0,008974987 |
| *ENSG00000138180* | *CEP55* | 1,89 | 0,001212512 |
| *ENSG00000100583* | *SAMD15* | 1,89 | 0,014397929 |
| *ENSG00000197457* | *STMN3* | 1,89 | 0,040866112 |
| *ENSG00000184227* | *ACOT1* | 1,88 | 0,013665513 |
| *ENSG00000196550* | *FAM72A* | 1,88 | 0,003267866 |
| *ENSG00000283154* | *IQCJ-SCHIP1* | 1,88 | 6,06723E-07 |
| *ENSG00000092853* | *CLSPN* | 1,87 | 0,049551057 |
| *ENSG00000179598* | *PLD6* | 1,87 | 0,000110892 |
| *ENSG00000122367* | *LDB3* | 1,87 | 0,023420371 |
| *ENSG00000185304* | *RGPD2* | 1,87 | 0,019205691 |
| *ENSG00000179528* | *LBX2* | 1,87 | 1,24754E-05 |
| *ENSG00000113810* | *SMC4* | 1,86 | 4,43478E-05 |
| *ENSG00000146670* | *CDCA5* | 1,86 | 0,012088905 |
| *ENSG00000198830* | *HMGN2* | 1,86 | 2,22749E-07 |

| | | | |
|---|---|---|---|
| *ENSG00000262209* | *PCDHGB3* | 1,85 | 0,024884979 |
| *ENSG00000131941* | *RHPN2* | 1,85 | 0,00416921 |
| *ENSG00000089685* | *BIRC5* | 1,85 | 0,000939189 |
| *ENSG00000105997* | *HOXA3* | 1,85 | 2,42424E-05 |
| *ENSG00000134940* | *ACRV1* | 1,85 | 0,045507295 |
| *ENSG00000165244* | *ZNF367* | 1,84 | 0,00014725 |
| *ENSG00000071794* | *HLTF* | 1,83 | 0,01836873 |
| *ENSG00000166851* | *PLK1* | 1,83 | 0,001531099 |
| *ENSG00000058056* | *USP13* | 1,83 | 0,00323915 |
| *ENSG00000076351* | *SLC46A1* | 1,83 | 0,038160828 |
| *ENSG00000131747* | *TOP2A* | 1,83 | 0,001069123 |
| *ENSG00000116771* | *AGMAT* | 1,83 | 0,019288314 |
| *ENSG00000139192* | *TAPBPL* | 1,83 | 0,007919018 |
| *ENSG00000163808* | *KIF15* | 1,82 | 0,01383483 |
| *ENSG00000104518* | *GSDMD* | 1,82 | 7,6797E-06 |
| *ENSG00000157456* | *CCNB2* | 1,81 | 0,000173455 |
| *ENSG00000225805* | *DEFB131B* | 1,81 | 0,016906688 |
| *ENSG00000066279* | *ASPM* | 1,80 | 0,000502229 |
| *ENSG00000127399* | *LRRC61* | 1,80 | 0,038372673 |
| *ENSG00000166359* | *WDR88* | 1,80 | 0,000542711 |
| *ENSG00000163072* | *NOSTRIN* | 1,80 | 0,001212512 |
| *ENSG00000206140* | *TMEM191C* | 1,79 | 0,044354143 |
| *ENSG00000162391* | *FAM151A* | 1,79 | 0,044296742 |
| *ENSG00000166813* | *KIF7* | 1,79 | 0,000215478 |
| *ENSG00000178919* | *FOXE1* | 1,79 | 0,001384456 |
| *ENSG00000135314* | *KHDC1* | 1,79 | 1,4567E-12 |
| *ENSG00000134470* | *IL15RA* | 1,78 | 0,032988605 |
| *ENSG00000215784* | *FAM72D* | 1,77 | 0,037953463 |
| *ENSG00000237649* | *KIFC1* | 1,76 | 0,001521275 |
| *ENSG00000164649* | *CDCA7L* | 1,76 | 5,48823E-05 |
| *ENSG00000110237* | *ARHGEF17* | 1,76 | 0,000822715 |
| *ENSG00000123609* | *NMI* | 1,76 | 7,12962E-06 |
| *ENSG00000106868* | *SUSD1* | 1,75 | 0,001421351 |
| *ENSG00000100162* | *CENPM* | 1,75 | 0,009212296 |
| *ENSG00000112029* | *FBXO5* | 1,75 | 0,000215478 |
| *ENSG00000004139* | *SARM1* | 1,74 | 0,022748291 |
| *ENSG00000205220* | *PSMB10* | 1,74 | 0,041221349 |
| *ENSG00000111291* | *GPRC5D* | 1,74 | 0,026143554 |
| *ENSG00000112742* | *TTK* | 1,74 | 0,002417134 |
| *ENSG00000130193* | *THEM6* | 1,73 | 0,017546363 |
| *ENSG00000040275* | *SPDL1* | 1,73 | 0,048588919 |
| *ENSG00000140451* | *PIF1* | 1,73 | 6,77538E-05 |
| *ENSG00000137807* | *KIF23* | 1,72 | 0,00121472 |
| *ENSG00000213186* | *TRIM59* | 1,72 | 6,64494E-05 |
| *ENSG00000126787* | *DLGAP5* | 1,72 | 0,003048792 |

| | | | |
|---|---|---|---|
| *ENSG00000165568* | *AKR1E2* | 1,72 | 0,001421351 |
| *ENSG00000136824* | *SMC2* | 1,72 | 0,000290153 |
| *ENSG00000164985* | *PSIP1* | 1,71 | 0,017303464 |
| *ENSG00000121152* | *NCAPH* | 1,71 | 0,037953463 |
| *ENSG00000111247* | *RAD51AP1* | 1,71 | 0,023519857 |
| *ENSG00000169679* | *BUB1* | 1,70 | 0,009954176 |
| *ENSG00000276043* | *UHRF1* | 1,70 | 0,019198061 |
| *ENSG00000129195* | *PIMREG* | 1,70 | 0,016215899 |
| *ENSG00000147119* | *CHST7* | 1,70 | 0,049503279 |
| *ENSG00000138778* | *CENPE* | 1,70 | 0,010811453 |
| *ENSG00000206535* | *LNP1* | 1,69 | 0,00094348 |
| *ENSG00000175305* | *CCNE2* | 1,69 | 0,003854191 |
| *ENSG00000166803* | *PCLAF* | 1,69 | 0,013071654 |
| *ENSG00000178999* | *AURKB* | 1,69 | 0,032277165 |
| *ENSG00000010292* | *NCAPD2* | 1,68 | 0,00012629 |
| *ENSG00000188610* | *FAM72B* | 1,68 | 0,012415292 |
| *ENSG00000163923* | *RPL39L* | 1,68 | 0,03573181 |
| *ENSG00000024526* | *DEPDC1* | 1,68 | 0,00748688 |
| *ENSG00000122952* | *ZWINT* | 1,68 | 0,010037507 |
| *ENSG00000158470* | *B4GALT5* | 1,68 | 0,036968856 |
| *ENSG00000185404* | *SP140L* | 1,67 | 0,000110303 |
| *ENSG00000134398* | *ERN2* | 1,67 | 0,002267948 |
| *ENSG00000164061* | *BSN* | 1,67 | 0,009940079 |
| *ENSG00000175322* | *ZNF519* | 1,67 | 0,005030516 |
| *ENSG00000118193* | *KIF14* | 1,66 | 0,029247581 |
| *ENSG00000189057* | *FAM111B* | 1,66 | 0,029137051 |
| *ENSG00000204366* | *ZBTB12* | 1,66 | 0,01077822 |
| *ENSG00000226174* | *TEX22* | 1,65 | 0,004953319 |
| *ENSG00000101003* | *GINS1* | 1,65 | 0,023103455 |
| *ENSG00000148120* | *C9orf3* | 1,65 | 0,039737626 |
| *ENSG00000196526* | *AFAP1* | 1,65 | 0,032287881 |
| *ENSG00000198901* | *PRC1* | 1,63 | 0,00300696 |
| *ENSG00000140534* | *TICRR* | 1,63 | 0,007855542 |
| *ENSG00000164104* | *HMGB2* | 1,63 | 0,000721283 |
| *ENSG00000167900* | *TK1* | 1,63 | 0,026193491 |
| *ENSG00000196584* | *XRCC2* | 1,62 | 0,048170885 |
| *ENSG00000165661* | *QSOX2* | 1,62 | 0,002867818 |
| *ENSG00000120334* | *CENPL* | 1,62 | 0,001296711 |
| *ENSG00000127124* | *HIVEP3* | 1,61 | 0,0167851 |
| *ENSG00000162063* | *CCNF* | 1,61 | 0,003136739 |
| *ENSG00000112208* | *BAG2* | 1,61 | 0,046754307 |
| *ENSG00000106077* | *ABHD11* | 1,60 | 0,004593779 |
| *ENSG00000115163* | *CENPA* | 1,60 | 0,002168175 |
| *ENSG00000180448* | *ARHGAP45* | 1,60 | 0,029247581 |
| *ENSG00000149089* | *APIP* | 1,58 | 0,040472039 |

| | | | |
|---|---|---|---|
| *ENSG00000169255* | *B3GALNT1* | 1,58 | 0,018234704 |
| *ENSG00000175564* | *UCP3* | 1,57 | 0,017838645 |
| *ENSG00000135052* | *GOLM1* | 1,56 | 0,000336165 |
| *ENSG00000157064* | *NMNAT2* | 1,56 | 0,040601987 |
| *ENSG00000131351* | *HAUS8* | 1,56 | 0,005672128 |
| *ENSG00000114346* | *ECT2* | 1,56 | 0,001912358 |
| *ENSG00000123473* | *STIL* | 1,55 | 0,017796565 |
| *ENSG00000185920* | *PTCH1* | 1,55 | 0,04773314 |
| *ENSG00000078269* | *SYNJ2* | 1,55 | 0,002660798 |
| *ENSG00000119333* | *WDR34* | 1,55 | 1,14785E-06 |
| *ENSG00000033170* | *FUT8* | 1,55 | 0,03052963 |
| *ENSG00000100350* | *FOXRED2* | 1,54 | 0,015929836 |
| *ENSG00000254469* | | 1,53 | 0,000611594 |
| *ENSG00000187189* | *TSPYL4* | 1,53 | 0,000530614 |
| *ENSG00000120729* | *MYOT* | 1,53 | 0,001138149 |
| *ENSG00000104147* | *OIP5* | 1,53 | 0,003254269 |
| *ENSG00000172167* | *MTBP* | 1,53 | 0,033236533 |
| *ENSG00000075702* | *WDR62* | 1,52 | 0,033745386 |
| *ENSG00000110721* | *CHKA* | 1,52 | 0,000721283 |
| *ENSG00000095637* | *SORBS1* | 1,52 | 0,049517784 |
| *ENSG00000112186* | *CAP2* | 1,52 | 0,034029106 |
| *ENSG00000164758* | *MED30* | 1,52 | 0,000461613 |
| *ENSG00000160298* | *C21orf58* | 1,51 | 0,000228088 |
| *ENSG00000109674* | *NEIL3* | 1,51 | 0,041221349 |
| *ENSG00000160345* | *C9orf116* | 1,50 | 0,04114348 |
| *ENSG00000134057* | *CCNB1* | 1,50 | 0,016195737 |
| *ENSG00000186470* | *BTN3A2* | 1,50 | 0,049517784 |
| *ENSG00000113569* | *NUP155* | 1,49 | 0,016840334 |
| *ENSG00000198826* | *ARHGAP11A* | 1,48 | 0,007094538 |
| *ENSG00000131015* | *ULBP2* | 1,48 | 0,020165862 |
| *ENSG00000261884* | | 1,48 | 0,006011311 |
| *ENSG00000186564* | *FOXD2* | 1,47 | 0,00820742 |
| *ENSG00000156970* | *BUB1B* | 1,47 | 0,047670412 |
| *ENSG00000149636* | *DSN1* | 1,47 | 0,023283179 |
| *ENSG00000025423* | *HSD17B6* | 1,47 | 0,004554641 |
| *ENSG00000132773* | *TOE1* | 1,47 | 0,000742542 |
| *ENSG00000100625* | *SIX4* | 1,46 | 0,000190436 |
| *ENSG00000284512* | | 1,46 | 0,012854302 |
| *ENSG00000141934* | *PLPP2* | 1,46 | 0,023472554 |
| *ENSG00000168005* | *SPINDOC* | 1,46 | 0,004221593 |
| *ENSG00000137404* | *NRM* | 1,46 | 0,013545788 |
| *ENSG00000197576* | *HOXA4* | 1,45 | 0,019208952 |
| *ENSG00000143153* | *ATP1B1* | 1,45 | 0,035840629 |
| *ENSG00000171877* | *FRMD5* | 1,45 | 0,041693246 |
| *ENSG00000106524* | *ANKMY2* | 1,44 | 0,001183316 |

| | | | |
|---|---|---|---|
| *ENSG00000135835* | *KIAA1614* | 1,44 | 0,010137488 |
| *ENSG00000178202* | *KDELC2* | 1,44 | 0,030238019 |
| *ENSG00000186197* | *EDARADD* | 1,44 | 2,36842E-05 |
| *ENSG00000214827* | *MTCP1* | 1,44 | 8,24618E-05 |
| *ENSG00000013573* | *DDX11* | 1,43 | 5,30735E-05 |
| *ENSG00000077152* | *UBE2T* | 1,43 | 0,044908576 |
| *ENSG00000172508* | *CARNS1* | 1,43 | 0,038187292 |
| *ENSG00000146535* | *GNA12* | 1,43 | 0,015626744 |
| *ENSG00000174804* | *FZD4* | 1,42 | 0,016762196 |
| *ENSG00000060140* | *STYK1* | 1,42 | 0,04725477 |
| *ENSG00000104524* | *PYCR3* | 1,41 | 0,004876664 |
| *ENSG00000180626* | *ZNF594* | 1,41 | 0,007990566 |
| *ENSG00000160447* | *PKN3* | 1,41 | 0,000400281 |
| *ENSG00000152291* | *TGOLN2* | 1,41 | 2,04836E-05 |
| *ENSG00000111203* | *ITFG2* | 1,40 | 2,15567E-06 |
| *ENSG00000117632* | *STMN1* | 1,40 | 0,019170555 |
| *ENSG00000205730* | *ITPRIPL2* | 1,40 | 0,000629021 |
| *ENSG00000213397* | *HAUS7* | 1,40 | 0,036547613 |
| *ENSG00000143498* | *TAF1A* | 1,40 | 0,023332833 |
| *ENSG00000105968* | *H2AFV* | 1,40 | 0,000595699 |
| *ENSG00000166669* | *ATF7IP2* | 1,39 | 0,01209456 |
| *ENSG00000197980* | *LEKR1* | 1,39 | 0,012066691 |
| *ENSG00000042286* | *AIFM2* | 1,39 | 0,000891977 |
| *ENSG00000109881* | *CCDC34* | 1,39 | 0,01872092 |
| *ENSG00000221886* | *ZBED8* | 1,38 | 0,028441501 |
| *ENSG00000145386* | *CCNA2* | 1,38 | 0,032363212 |
| *ENSG00000078795* | *PKD2L2* | 1,38 | 6,35798E-05 |
| *ENSG00000104738* | *MCM4* | 1,38 | 0,040922918 |
| *ENSG00000120802* | *TMPO* | 1,37 | 0,005291631 |
| *ENSG00000089505* | *CMTM1* | 1,37 | 0,00353113 |
| *ENSG00000149503* | *INCENP* | 1,37 | 0,002621458 |
| *ENSG00000204815* | *TTC25* | 1,36 | 0,039760832 |
| *ENSG00000146918* | *NCAPG2* | 1,36 | 0,014594276 |
| *ENSG00000136811* | *ODF2* | 1,36 | 0,022001264 |
| *ENSG00000160957* | *RECQL4* | 1,36 | 0,022185829 |
| *ENSG00000151967* | *SCHIP1* | 1,36 | 0,009212303 |
| *ENSG00000176619* | *LMNB2* | 1,36 | 0,00412188 |
| *ENSG00000001036* | *FUCA2* | 1,34 | 0,000120232 |
| *ENSG00000110719* | *TCIRG1* | 1,34 | 0,026362878 |
| *ENSG00000189362* | *NEMP2* | 1,34 | 0,000876944 |
| *ENSG00000136122* | *BORA* | 1,32 | 0,002336961 |
| *ENSG00000114107* | *CEP70* | 1,31 | 0,014481724 |
| *ENSG00000146143* | *PRIM2* | 1,31 | 0,026990741 |
| *ENSG00000031003* | *FAM13B* | 1,31 | 0,000195191 |
| *ENSG00000155850* | *SLC26A2* | 1,31 | 0,036547613 |

| | | | |
|---|---|---|---|
| *ENSG00000160949* | *TONSL* | 1,30 | 0,034609275 |
| *ENSG00000013810* | *TACC3* | 1,30 | 0,018850304 |
| *ENSG00000166938* | *DIS3L* | 1,30 | 7,5372E-05 |
| *ENSG00000010270* | *STARD3NL* | 1,30 | 0,002630274 |
| *ENSG00000106266* | *SNX8* | 1,29 | 0,023905976 |
| *ENSG00000106012* | *IQCE* | 1,29 | 0,00037689 |
| *ENSG00000124795* | *DEK* | 1,29 | 0,000250701 |
| *ENSG00000137812* | *KNL1* | 1,29 | 0,006186577 |
| *ENSG00000131238* | *PPT1* | 1,29 | 0,038603703 |
| *ENSG00000025772* | *TOMM34* | 1,28 | 0,010144137 |
| *ENSG00000051825* | *MPHOSPH9* | 1,27 | 0,029003274 |
| *ENSG00000112977* | *DAP* | 1,27 | 0,038220635 |
| *ENSG00000136603* | *SKIL* | 1,27 | 0,014100424 |
| *ENSG00000166246* | *C16orf71* | 1,27 | 0,041628688 |
| *ENSG00000114859* | *CLCN2* | 1,26 | 0,000537228 |
| *ENSG00000118707* | *TGIF2* | 1,26 | 0,000900016 |
| *ENSG00000112773* | *TENT5A* | 1,26 | 0,006115321 |
| *ENSG00000146757* | *ZNF92* | 1,25 | 0,035810112 |
| *ENSG00000164002* | *EXO5* | 1,25 | 0,000335561 |
| *ENSG00000161847* | *RAVER1* | 1,25 | 0,00175949 |
| *ENSG00000197818* | *SLC9A8* | 1,25 | 8,37537E-06 |
| *ENSG00000100105* | *PATZ1* | 1,25 | 6,64494E-05 |
| *ENSG00000130584* | *ZBTB46* | 1,24 | 0,014292734 |
| *ENSG00000217555* | *CKLF* | 1,24 | 0,005891465 |
| *ENSG00000110583* | *NAA40* | 1,24 | 0,002267565 |
| *ENSG00000259330* | *INAFM2* | 1,23 | 0,022645967 |
| *ENSG00000071243* | *ING3* | 1,23 | 4,66678E-05 |
| *ENSG00000183647* | *ZNF530* | 1,23 | 0,016065816 |
| *ENSG00000121864* | *ZNF639* | 1,22 | 0,001973432 |
| *ENSG00000185361* | *TNFAIP8L1* | 1,22 | 0,028683617 |
| *ENSG00000197905* | *TEAD4* | 1,22 | 0,003149915 |
| *ENSG00000106009* | *BRAT1* | 1,22 | 0,002426087 |
| *ENSG00000101871* | *MID1* | 1,22 | 0,009392942 |
| *ENSG00000151725* | *CENPU* | 1,21 | 0,013918649 |
| *ENSG00000119599* | *DCAF4* | 1,21 | 0,016282599 |
| *ENSG00000163913* | *IFT122* | 1,21 | 0,04271703 |
| *ENSG00000157653* | *C9orf43* | 1,21 | 0,007424371 |
| *ENSG00000166508* | *MCM7* | 1,20 | 0,024425552 |
| *ENSG00000168778* | *TCTN2* | 1,20 | 0,036691433 |
| *ENSG00000183856* | *IQGAP3* | 1,20 | 0,027058332 |
| *ENSG00000143401* | *ANP32E* | 1,20 | 0,015456001 |
| *ENSG00000088448* | *ANKRD10* | 1,19 | 0,012587542 |
| *ENSG00000149639* | *SOGA1* | 1,19 | 0,001913414 |
| *ENSG00000132780* | *NASP* | 1,18 | 0,025688045 |
| *ENSG00000165752* | *STK32C* | 1,18 | 0,029989739 |

| | | | |
|---|---|---|---|
| *ENSG00000197724* | *PHF2* | 1,18 | 0,002324008 |
| *ENSG00000127191* | *TRAF2* | 1,17 | 0,000646986 |
| *ENSG00000145945* | *FAM50B* | 1,17 | 0,04114348 |
| *ENSG00000128944* | *KNSTRN* | 1,17 | 0,005925397 |
| *ENSG00000121775* | *TMEM39B* | 1,17 | 0,000196842 |
| *ENSG00000148200* | *NR6A1* | 1,17 | 0,04069547 |
| *ENSG00000106399* | *RPA3* | 1,16 | 0,020938483 |
| *ENSG00000198919* | *DZIP3* | 1,16 | 0,037210423 |
| *ENSG00000163535* | *SGO2* | 1,16 | 0,023248889 |
| *ENSG00000160563* | *MED27* | 1,16 | 0,000335561 |
| *ENSG00000117519* | *CNN3* | 1,16 | 0,032994873 |
| *ENSG00000139278* | *GLIPR1* | 1,16 | 0,043533225 |
| *ENSG00000180611* | *MB21D2* | 1,15 | 0,015478836 |
| *ENSG00000113360* | *DROSHA* | 1,15 | 0,011932616 |
| *ENSG00000178966* | *RMI1* | 1,15 | 0,009989146 |
| *ENSG00000188321* | *ZNF559* | 1,15 | 0,006362817 |
| *ENSG00000215883* | *CYB5RL* | 1,15 | 0,0449536 |
| *ENSG00000214654* | *B3GNT10* | 1,15 | 0,027697042 |
| *ENSG00000106144* | *CASP2* | 1,15 | 0,005511643 |
| *ENSG00000096092* | *TMEM14A* | 1,15 | 0,003008485 |
| *ENSG00000126883* | *NUP214* | 1,15 | 5,85766E-05 |
| *ENSG00000168014* | *C2CD3* | 1,15 | 0,003982302 |
| *ENSG00000163577* | *EIF5A2* | 1,14 | 0,000966161 |
| *ENSG00000171448* | *ZBTB26* | 1,14 | 3,23762E-10 |
| *ENSG00000119673* | *ACOT2* | 1,14 | 0,025521186 |
| *ENSG00000148362* | *PAXX* | 1,14 | 0,006153099 |
| *ENSG00000162607* | *USP1* | 1,14 | 0,007455698 |
| *ENSG00000177182* | *CLVS1* | 1,14 | 0,037188882 |
| *ENSG00000136206* | *SPDYE1* | 1,14 | 0,022466095 |
| *ENSG00000214078* | *CPNE1* | 1,13 | 0,047898686 |
| *ENSG00000185347* | *TEDC1* | 1,13 | 0,026967056 |
| *ENSG00000148229* | *POLE3* | 1,13 | 0,010731262 |
| *ENSG00000143799* | *PARP1* | 1,13 | 0,001157179 |
| *ENSG00000161800* | *RACGAP1* | 1,13 | 0,010447415 |
| *ENSG00000196597* | *ZNF782* | 1,13 | 0,003075389 |
| *ENSG00000183395* | *PMCH* | 1,12 | 0,04284181 |
| *ENSG00000130309* | *COLGALT1* | 1,12 | 0,018769164 |
| *ENSG00000198887* | *SMC5* | 1,12 | 0,004044917 |
| *ENSG00000184786* | *TCTE3* | 1,12 | 0,012016631 |
| *ENSG00000253293* | *HOXA10* | 1,11 | 0,00573323 |
| *ENSG00000171823* | *FBXL14* | 1,11 | 0,001338626 |
| *ENSG00000125520* | *SLC2A4RG* | 1,11 | 0,011129366 |
| *ENSG00000196776* | *CD47* | 1,10 | 0,048730581 |
| *ENSG00000119392* | *GLE1* | 1,10 | 0,00591707 |
| *ENSG00000163781* | *TOPBP1* | 1,10 | 0,010855892 |

| | | | |
|---|---|---|---|
| *ENSG00000138182* | *KIF20B* | 1,10 | 0,013573577 |
| *ENSG00000101413* | *RPRD1B* | 1,10 | 0,000964478 |
| *ENSG00000196935* | *SRGAP1* | 1,09 | 0,032175005 |
| *ENSG00000167525* | *PROCA1* | 1,09 | 0,005360497 |
| *ENSG00000124279* | *FASTKD3* | 1,09 | 0,004750492 |
| *ENSG00000156500* | *FAM122C* | 1,09 | 0,004698533 |
| *ENSG00000132434* | *LANCL2* | 1,09 | 0,009722439 |
| *ENSG00000188807* | *TMEM201* | 1,08 | 0,043595302 |
| *ENSG00000105643* | *ARRDC2* | 1,08 | 0,048018792 |
| *ENSG00000256537* | *SMIM10L1* | 1,08 | 0,000596315 |
| *ENSG00000158169* | *FANCC* | 1,08 | 0,004876664 |
| *ENSG00000226479* | *TMEM185B* | 1,07 | 0,04917418 |
| *ENSG00000130713* | *EXOSC2* | 1,07 | 0,028533305 |
| *ENSG00000138190* | *EXOC6* | 1,06 | 0,041716708 |
| *ENSG00000012660* | *ELOVL5* | 1,06 | 0,007809823 |
| *ENSG00000176142* | *TMEM39A* | 1,06 | 0,029247581 |
| *ENSG00000175787* | *ZNF169* | 1,06 | 0,000123933 |
| *ENSG00000137770* | *CTDSPL2* | 1,06 | 0,042304411 |
| *ENSG00000176058* | *TPRN* | 1,06 | 0,008561192 |
| *ENSG00000077044* | *DGKD* | 1,06 | 0,013671407 |
| *ENSG00000133111* | *RFXAP* | 1,05 | 0,013231343 |
| *ENSG00000139190* | *VAMP1* | 1,05 | 0,011086252 |
| *ENSG00000178764* | *ZHX2* | 1,04 | 0,003905908 |
| *ENSG00000112877* | *CEP72* | 1,04 | 0,04481492 |
| *ENSG00000164754* | *RAD21* | 1,03 | 0,000631017 |
| *ENSG00000119661* | *DNAL1* | 1,03 | 0,007306291 |
| *ENSG00000064012* | *CASP8* | 1,03 | 0,015373376 |
| *ENSG00000170469* | *SPATA24* | 1,03 | 0,041202686 |
| *ENSG00000204822* | *MRPL53* | 1,03 | 0,037401915 |
| *ENSG00000090857* | *PDPR* | 1,02 | 0,021524463 |
| *ENSG00000169683* | *LRRC45* | 1,02 | 0,00048446 |
| *ENSG00000086475* | *SEPHS1* | 1,02 | 0,027731889 |
| *ENSG00000158122* | *PRXL2C* | 1,02 | 0,029734546 |
| *ENSG00000164916* | *FOXK1* | 1,02 | 0,014440197 |
| *ENSG00000135436* | *FAM186B* | 1,02 | 0,00153476 |
| *ENSG00000169925* | *BRD3* | 1,01 | 0,000220661 |
| *ENSG00000111196* | *MAGOHB* | 1,01 | 0,036938773 |
| *ENSG00000131019* | *ULBP3* | 1,01 | 0,047731978 |
| *ENSG00000163935* | *SFMBT1* | 1,01 | 0,029976436 |
| *ENSG00000238227* | *TMEM250* | 1,01 | 0,003222111 |
| *ENSG00000183337* | *BCOR* | 1,01 | 6,77733E-06 |
| *ENSG00000158636* | *EMSY* | 1,01 | 0,000195278 |
| *ENSG00000221843* | *C2orf16* | 1,00 | 0,003182631 |
| *ENSG00000101452* | *DHX35* | 1,00 | 5,74708E-05 |
| *ENSG00000106346* | *USP42* | 1,00 | 0,003188101 |

| | | | |
|---|---|---|---|
| *ENSG00000176248* | *ANAPC2* | 1,00 | 0,000881353 |
| *ENSG00000166451* | *CENPN* | 1,00 | 0,041690256 |
| *ENSG00000143190* | *POU2F1* | 1,00 | 0,004875103 |
| *ENSG00000187634* | *SAMD11* | 1,00 | 0,002681509 |
| *ENSG00000169919* | *GUSB* | 1,00 | 0,010026494 |
| *ENSG00000198728* | *LDB1* | -1,00 | 0,006038004 |
| *ENSG00000100129* | *EIF3L* | -1,00 | 8,53669E-05 |
| *ENSG00000185650* | *ZFP36L1* | -1,01 | 0,018791287 |
| *ENSG00000160691* | *SHC1* | -1,01 | 0,043870836 |
| *ENSG00000099246* | *RAB18* | -1,01 | 0,016598168 |
| *ENSG00000137806* | *NDUFAF1* | -1,01 | 0,012860507 |
| *ENSG00000185189* | *NRBP2* | -1,01 | 0,008289516 |
| *ENSG00000153130* | *SCOC* | -1,01 | 0,000308033 |
| *ENSG00000063177* | *RPL18* | -1,01 | 7,83382E-05 |
| *ENSG00000047849* | *MAP4* | -1,01 | 0,044354143 |
| *ENSG00000110628* | *SLC22A18* | -1,02 | 0,031510101 |
| *ENSG00000177042* | *TMEM80* | -1,02 | 0,000151898 |
| *ENSG00000275183* | *LENG9* | -1,02 | 0,000557179 |
| *ENSG00000135698* | *MPHOSPH6* | -1,02 | 0,000123306 |
| *ENSG00000166441* | *RPL27A* | -1,02 | 0,021404066 |
| *ENSG00000011638* | *TMEM159* | -1,02 | 0,003727867 |
| *ENSG00000166886* | *NAB2* | -1,02 | 0,001750014 |
| *ENSG00000156508* | *EEF1A1* | -1,02 | 0,00270494 |
| *ENSG00000197093* | *GAL3ST4* | -1,02 | 0,026143554 |
| *ENSG00000084733* | *RAB10* | -1,03 | 0,003267551 |
| *ENSG00000027001* | *MIPEP* | -1,03 | 0,021075969 |
| *ENSG00000136997* | *MYC* | -1,03 | 0,049567863 |
| *ENSG00000063046* | *EIF4B* | -1,03 | 8,21149E-07 |
| *ENSG00000197958* | *RPL12* | -1,03 | 0,011656862 |
| *ENSG00000151552* | *QDPR* | -1,03 | 0,027877061 |
| *ENSG00000128590* | *DNAJB9* | -1,03 | 0,004065416 |
| *ENSG00000168575* | *SLC20A2* | -1,03 | 0,029247581 |
| *ENSG00000174748* | *RPL15* | -1,04 | 0,006909585 |
| *ENSG00000213906* | *LTB4R2* | -1,04 | 0,02810648 |
| *ENSG00000121897* | *LIAS* | -1,04 | 0,003073238 |
| *ENSG00000182774* | *RPS17* | -1,04 | 0,013181967 |
| *ENSG00000164253* | *WDR41* | -1,04 | 3,69987E-05 |
| *ENSG00000163682* | *RPL9* | -1,04 | 0,011428809 |
| *ENSG00000161533* | *ACOX1* | -1,05 | 0,010752506 |
| *ENSG00000087008* | *ACOX3* | -1,05 | 0,010277071 |
| *ENSG00000144824* | *PHLDB2* | -1,05 | 0,026193491 |
| *ENSG00000186184* | *POLR1D* | -1,05 | 0,003332752 |
| *ENSG00000110651* | *CD81* | -1,05 | 0,000854827 |
| *ENSG00000154025* | *SLC5A10* | -1,05 | 0,007090874 |
| *ENSG00000119537* | *KDSR* | -1,06 | 0,007520851 |

| | | | |
|---|---|---|---|
| *ENSG00000148985* | *PGAP2* | -1,06 | 0,027881677 |
| *ENSG00000072778* | *ACADVL* | -1,06 | 0,001962079 |
| *ENSG00000168028* | *RPSA* | -1,06 | 0,044708858 |
| *ENSG00000153879* | *CEBPG* | -1,06 | 1,56119E-05 |
| *ENSG00000141198* | *TOM1L1* | -1,07 | 0,004698533 |
| *ENSG00000006757* | *PNPLA4* | -1,07 | 0,038982998 |
| *ENSG00000110700* | *RPS13* | -1,07 | 0,044419062 |
| *ENSG00000197535* | *MYO5A* | -1,08 | 0,002802879 |
| *ENSG00000186577* | *SMIM29* | -1,08 | 0,010104162 |
| *ENSG00000145425* | *RPS3A* | -1,09 | 0,001708401 |
| *ENSG00000139684* | *ESD* | -1,09 | 5,09459E-06 |
| *ENSG00000198715* | *GLMP* | -1,09 | 0,048049145 |
| *ENSG00000214274* | *ANG* | -1,09 | 0,026665871 |
| *ENSG00000116044* | *NFE2L2* | -1,09 | 0,002411307 |
| *ENSG00000198839* | *ZNF277* | -1,09 | 0,002174983 |
| *ENSG00000169228* | *RAB24* | -1,10 | 0,016180603 |
| *ENSG00000109270* | *LAMTOR3* | -1,10 | 0,008632834 |
| *ENSG00000147364* | *FBXO25* | -1,10 | 0,000559626 |
| *ENSG00000188522* | *FAM83G* | -1,10 | 0,005022182 |
| *ENSG00000171469* | *ZNF561* | -1,10 | 0,022854606 |
| *ENSG00000166598* | *HSP90B1* | -1,11 | 0,012733627 |
| *ENSG00000147403* | *RPL10* | -1,11 | 0,007119877 |
| *ENSG00000168010* | *ATG16L2* | -1,11 | 0,037450658 |
| *ENSG00000196072* | *BLOC1S2* | -1,12 | 0,001011709 |
| *ENSG00000182208* | *MOB2* | -1,12 | 0,000183319 |
| *ENSG00000137802* | *MAPKBP1* | -1,12 | 0,00178724 |
| *ENSG00000175390* | *EIF3F* | -1,13 | 0,00094348 |
| *ENSG00000170855* | *TRIAP1* | -1,13 | 1,48224E-06 |
| *ENSG00000145050* | *MANF* | -1,13 | 0,021680644 |
| *ENSG00000134419* | *RPS15A* | -1,13 | 0,000145442 |
| *ENSG00000165169* | *DYNLT3* | -1,14 | 0,000284224 |
| *ENSG00000118181* | *RPS25* | -1,15 | 1,80541E-06 |
| *ENSG00000198743* | *SLC5A3* | -1,15 | 0,023554442 |
| *ENSG00000198933* | *TBKBP1* | -1,15 | 0,006068419 |
| *ENSG00000135596* | *MICAL1* | -1,15 | 0,012322398 |
| *ENSG00000170889* | *RPS9* | -1,16 | 1,95266E-05 |
| *ENSG00000204977* | *TRIM13* | -1,16 | 0,008351849 |
| *ENSG00000125691* | *RPL23* | -1,16 | 0,000605264 |
| *ENSG00000112874* | *NUDT12* | -1,16 | 0,002173295 |
| *ENSG00000141034* | *GID4* | -1,17 | 0,007593681 |
| *ENSG00000134108* | *ARL8B* | -1,17 | 0,002498938 |
| *ENSG00000167658* | *EEF2* | -1,17 | 0,006626244 |
| *ENSG00000169018* | *FEM1B* | -1,18 | 0,001829216 |
| *ENSG00000109458* | *GAB1* | -1,18 | 0,011888421 |
| *ENSG00000105372* | *RPS19* | -1,18 | 6,30854E-05 |

| | | | |
|---|---|---|---|
| *ENSG00000151665* | *PIGF* | -1,18 | 0,016519289 |
| *ENSG00000060558* | *GNA15* | -1,18 | 0,015379254 |
| *ENSG00000161970* | *RPL26* | -1,18 | 0,0029113 |
| *ENSG00000100979* | *PLTP* | -1,19 | 0,045049501 |
| *ENSG00000182606* | *TRAK1* | -1,19 | 0,016802552 |
| *ENSG00000138756* | *BMP2K* | -1,20 | 0,016275283 |
| *ENSG00000163191* | *S100A11* | -1,20 | 0,000348288 |
| *ENSG00000131370* | *SH3BP5* | -1,21 | 0,000247404 |
| *ENSG00000181029* | *TRAPPC5* | -1,21 | 0,013920676 |
| *ENSG00000154359* | *LONRF1* | -1,21 | 0,017991657 |
| *ENSG00000231500* | *RPS18* | -1,21 | 0,004178542 |
| *ENSG00000119950* | *MXI1* | -1,21 | 0,010559047 |
| *ENSG00000198755* | *RPL10A* | -1,22 | 1,72548E-06 |
| *ENSG00000172831* | *CES2* | -1,22 | 0,047404511 |
| *ENSG00000133678* | *TMEM254* | -1,23 | 0,006522036 |
| *ENSG00000154957* | *ZNF18* | -1,23 | 0,012138719 |
| *ENSG00000233927* | *RPS28* | -1,24 | 0,000507829 |
| *ENSG00000085465* | *OVGP1* | -1,24 | 0,041933019 |
| *ENSG00000031698* | *SARS* | -1,24 | 4,16271E-06 |
| *ENSG00000120129* | *DUSP1* | -1,25 | 0,011400456 |
| *ENSG00000144713* | *RPL32* | -1,25 | 0,014102704 |
| *ENSG00000198034* | *RPS4X* | -1,25 | 0,000323337 |
| *ENSG00000203546* | | -1,25 | 0,040137569 |
| *ENSG00000145390* | *USP53* | -1,25 | 0,001903218 |
| *ENSG00000129116* | *PALLD* | -1,25 | 0,038714694 |
| *ENSG00000120925* | *RNF170* | -1,25 | 1,85972E-05 |
| *ENSG00000180190* | *TDRP* | -1,25 | 0,000669821 |
| *ENSG00000171425* | *ZNF581* | -1,25 | 7,62721E-05 |
| *ENSG00000164096* | *C4orf3* | -1,26 | 0,009819197 |
| *ENSG00000078237* | *TIGAR* | -1,26 | 0,00815662 |
| *ENSG00000109670* | *FBXW7* | -1,26 | 0,003074841 |
| *ENSG00000118960* | *HS1BP3* | -1,26 | 0,001750014 |
| *ENSG00000135679* | *MDM2* | -1,27 | 0,012415292 |
| *ENSG00000165475* | *CRYL1* | -1,27 | 0,031510101 |
| *ENSG00000172493* | *AFF1* | -1,27 | 0,000507829 |
| *ENSG00000123191* | *ATP7B* | -1,27 | 0,020545316 |
| *ENSG00000100139* | *MICALL1* | -1,28 | 0,005502569 |
| *ENSG00000157077* | *ZFYVE9* | -1,28 | 4,06321E-06 |
| *ENSG00000099875* | *MKNK2* | -1,29 | 0,031371751 |
| *ENSG00000174444* | *RPL4* | -1,29 | 0,002723035 |
| *ENSG00000175197* | *DDIT3* | -1,29 | 3,11039E-07 |
| *ENSG00000068903* | *SIRT2* | -1,30 | 0,000202978 |
| *ENSG00000180964* | *TCEAL8* | -1,30 | 0,036675077 |
| *ENSG00000213903* | *LTB4R* | -1,31 | 0,012415292 |
| *ENSG00000148926* | *ADM* | -1,31 | 0,008843639 |

| | | | |
|---|---|---|---|
| *ENSG00000122026* | *RPL21* | -1,31 | 1,19386E-06 |
| *ENSG00000123600* | *METTL8* | -1,31 | 0,00286798 |
| *ENSG00000141337* | *ARSG* | -1,31 | 0,009176026 |
| *ENSG00000188846* | *RPL14* | -1,32 | 0,007985429 |
| *ENSG00000110315* | *RNF141* | -1,32 | 0,032319885 |
| *ENSG00000147416* | *ATP6V1B2* | -1,32 | 0,01415725 |
| *ENSG00000109475* | *RPL34* | -1,34 | 3,66074E-07 |
| *ENSG00000107679* | *PLEKHA1* | -1,34 | 0,005437636 |
| *ENSG00000162980* | *ARL5A* | -1,34 | 0,017386537 |
| *ENSG00000145439* | *CBR4* | -1,34 | 2,3466E-05 |
| *ENSG00000162458* | *FBLIM1* | -1,35 | 4,89143E-05 |
| *ENSG00000197756* | *RPL37A* | -1,36 | 9,81286E-05 |
| *ENSG00000050405* | *LIMA1* | -1,36 | 0,000269336 |
| *ENSG00000142669* | *SH3BGRL3* | -1,36 | 0,005860221 |
| *ENSG00000204947* | *ZNF425* | -1,36 | 0,037188882 |
| *ENSG00000178184* | *PARD6G* | -1,36 | 0,019483173 |
| *ENSG00000127366* | *TAS2R5* | -1,37 | 0,010177841 |
| *ENSG00000100364* | *KIAA0930* | -1,37 | 3,75872E-05 |
| *ENSG00000114993* | *RTKN* | -1,37 | 0,006038004 |
| *ENSG00000154642* | *C21orf91* | -1,37 | 0,003157521 |
| *ENSG00000115884* | *SDC1* | -1,38 | 0,029227962 |
| *ENSG00000181061* | *HIGD1A* | -1,38 | 0,00222641 |
| *ENSG00000171443* | *ZNF524* | -1,39 | 0,000521736 |
| *ENSG00000185761* | *ADAMTSL5* | -1,40 | 0,046796774 |
| *ENSG00000197329* | *PELI1* | -1,40 | 0,005946517 |
| *ENSG00000130766* | *SESN2* | -1,40 | 0,000208642 |
| *ENSG00000104361* | *NIPAL2* | -1,41 | 0,007589364 |
| *ENSG00000080546* | *SESN1* | -1,42 | 0,011929878 |
| *ENSG00000167065* | *DUSP18* | -1,42 | 0,000509651 |
| *ENSG00000143753* | *DEGS1* | -1,44 | 0,01261715 |
| *ENSG00000142541* | *RPL13A* | -1,45 | 8,66827E-06 |
| *ENSG00000170175* | *CHRNB1* | -1,45 | 0,031291281 |
| *ENSG00000162244* | *RPL29* | -1,45 | 0,000458672 |
| *ENSG00000100316* | *RPL3* | -1,45 | 1,01359E-05 |
| *ENSG00000125744* | *RTN2* | -1,45 | 8,0029E-06 |
| *ENSG00000243989* | *ACY1* | -1,47 | 0,030377957 |
| *ENSG00000136144* | *RCBTB1* | -1,47 | 0,002502543 |
| *ENSG00000108828* | *VAT1* | -1,48 | 0,003272009 |
| *ENSG00000163686* | *ABHD6* | -1,49 | 6,5062E-05 |
| *ENSG00000135926* | *TMBIM1* | -1,49 | 0,006458321 |
| *ENSG00000163393* | *SLC22A15* | -1,50 | 0,000631017 |
| *ENSG00000078902* | *TOLLIP* | -1,51 | 2,61335E-05 |
| *ENSG00000105514* | *RAB3D* | -1,52 | 0,047730076 |
| *ENSG00000103269* | *RHBDL1* | -1,53 | 0,036938773 |
| *ENSG00000179094* | *PER1* | -1,54 | 0,027473825 |

| | | | |
|---|---|---|---|
| *ENSG00000141542* | *RAB40B* | -1,54 | 0,040087719 |
| *ENSG00000134864* | *GGACT* | -1,54 | 0,010376197 |
| *ENSG00000164251* | *F2RL1* | -1,54 | 0,007141885 |
| *ENSG00000070540* | *WIPI1* | -1,55 | 0,003053527 |
| *ENSG00000051108* | *HERPUD1* | -1,55 | 0,008477166 |
| *ENSG00000052795* | *FNIP2* | -1,55 | 0,00259705 |
| *ENSG00000181350* | *LRRC75A* | -1,56 | 0,002310904 |
| *ENSG00000090097* | *PCBP4* | -1,56 | 0,024563017 |
| *ENSG00000090776* | *EFNB1* | -1,57 | 0,041202686 |
| *ENSG00000185088* | *RPS27L* | -1,57 | 5,48823E-05 |
| *ENSG00000164070* | *HSPA4L* | -1,57 | 0,019762485 |
| *ENSG00000122378* | *PRXL2A* | -1,57 | 0,011005315 |
| *ENSG00000119729* | *RHOQ* | -1,57 | 0,006478447 |
| *ENSG00000116717* | *GADD45A* | -1,57 | 0,002654617 |
| *ENSG00000215788* | *TNFRSF25* | -1,59 | 0,000864294 |
| *ENSG00000164086* | *DUSP7* | -1,59 | 0,007196031 |
| *ENSG00000160685* | *ZBTB7B* | -1,60 | 1,09113E-05 |
| *ENSG00000114738* | *MAPKAPK3* | -1,60 | 0,000477791 |
| *ENSG00000175984* | *DENND2C* | -1,60 | 0,031130054 |
| *ENSG00000164181* | *ELOVL7* | -1,61 | 0,016783018 |
| *ENSG00000221887* | *HMSD* | -1,61 | 0,026196364 |
| *ENSG00000140450* | *ARRDC4* | -1,61 | 0,007424371 |
| *ENSG00000181392* | *SYNE4* | -1,61 | 0,00152376 |
| *ENSG00000147697* | *GSDMC* | -1,62 | 0,004012249 |
| *ENSG00000113231* | *PDE8B* | -1,62 | 0,010855892 |
| *ENSG00000175416* | *CLTB* | -1,63 | 1,97969E-05 |
| *ENSG00000138772* | *ANXA3* | -1,63 | 0,003379812 |
| *ENSG00000169991* | *IFFO2* | -1,64 | 0,001267809 |
| *ENSG00000181804* | *SLC9A9* | -1,64 | 0,023332833 |
| *ENSG00000134107* | *BHLHE40* | -1,64 | 0,000201417 |
| *ENSG00000232859* | *LYRM9* | -1,65 | 0,037126975 |
| *ENSG00000163472* | *TMEM79* | -1,66 | 0,007520851 |
| *ENSG00000143669* | *LYST* | -1,66 | 0,002805231 |
| *ENSG00000133112* | *TPT1* | -1,66 | 6,88268E-05 |
| *ENSG00000104892* | *KLC3* | -1,66 | 0,005354701 |
| *ENSG00000167920* | *TMEM99* | -1,66 | 0,003031422 |
| *ENSG00000156299* | *TIAM1* | -1,68 | 0,025729975 |
| *ENSG00000070404* | *FSTL3* | -1,69 | 0,023309639 |
| *ENSG00000105755* | *ETHE1* | -1,69 | 0,001043936 |
| *ENSG00000171680* | *PLEKHG5* | -1,69 | 0,000885545 |
| *ENSG00000163803* | *PLB1* | -1,69 | 0,009586726 |
| *ENSG00000123892* | *RAB38* | -1,73 | 0,005682131 |
| *ENSG00000128346* | *C22orf23* | -1,74 | 2,07166E-05 |
| *ENSG00000176472* | *ZNF575* | -1,74 | 0,000289724 |
| *ENSG00000197956* | *S100A6* | -1,74 | 2,84487E-05 |

| | | | |
|---|---|---|---|
| *ENSG00000130751* | *NPAS1* | -1,77 | 0,019525551 |
| *ENSG00000172940* | *SLC22A13* | -1,79 | 0,001721416 |
| *ENSG00000142046* | *TMEM91* | -1,80 | 0,000195469 |
| *ENSG00000117407* | *ARTN* | -1,80 | 0,008964345 |
| *ENSG00000064115* | *TM7SF3* | -1,80 | 1,6311E-05 |
| *ENSG00000145284* | *SCD5* | -1,81 | 0,034029106 |
| *ENSG00000100994* | *PYGB* | -1,81 | 0,000186296 |
| *ENSG00000221869* | *CEBPD* | -1,82 | 0,000299624 |
| *ENSG00000160796* | *NBEAL2* | -1,82 | 0,001123192 |
| *ENSG00000067082* | *KLF6* | -1,83 | 8,53669E-05 |
| *ENSG00000179630* | *LACC1* | -1,83 | 0,005651574 |
| *ENSG00000112659* | *CUL9* | -1,83 | 0,002061069 |
| *ENSG00000157992* | *KRTCAP3* | -1,84 | 0,042770995 |
| *ENSG00000159348* | *CYB5R1* | -1,84 | 0,03069984 |
| *ENSG00000167676* | *PLIN4* | -1,84 | 0,002069801 |
| *ENSG00000157379* | *DHRS1* | -1,85 | 3,93501E-08 |
| *ENSG00000091622* | *PITPNM3* | -1,85 | 0,036025078 |
| *ENSG00000144671* | *SLC22A14* | -1,85 | 0,002333978 |
| *ENSG00000188643* | *S100A16* | -1,87 | 4,86471E-09 |
| *ENSG00000136011* | *STAB2* | -1,87 | 0,0449536 |
| *ENSG00000107957* | *SH3PXD2A* | -1,87 | 9,09001E-06 |
| *ENSG00000184545* | *DUSP8* | -1,88 | 0,00064473 |
| *ENSG00000063180* | *CA11* | -1,88 | 0,037139828 |
| *ENSG00000130066* | *SAT1* | -1,89 | 0,000133289 |
| *ENSG00000243056* | *EIF4EBP3* | -1,90 | 0,000306215 |
| *ENSG00000241106* | *HLA-DOB* | -1,93 | 0,021340852 |
| *ENSG00000196743* | *GM2A* | -1,93 | 0,009319186 |
| *ENSG00000141404* | *GNAL* | -1,95 | 0,009961785 |
| *ENSG00000120738* | *EGR1* | -1,96 | 3,69257E-05 |
| *ENSG00000087086* | *FTL* | -1,96 | 8,47873E-05 |
| *ENSG00000143382* | *ADAMTSL4* | -1,96 | 0,029227962 |
| *ENSG00000144712* | *CAND2* | -1,96 | 0,000753097 |
| *ENSG00000182580* | *EPHB3* | -1,97 | 0,03613826 |
| *ENSG00000099337* | *KCNK6* | -1,98 | 0,025688045 |
| *ENSG00000196139* | *AKR1C3* | -1,98 | 0,02546948 |
| *ENSG00000088726* | *TMEM40* | -1,98 | 0,045018106 |
| *ENSG00000101255* | *TRIB3* | -1,99 | 0,002053371 |
| *ENSG00000171649* | *ZIK1* | -2,00 | 0,007841301 |
| *ENSG00000106211* | *HSPB1* | -2,00 | 0,00291072 |
| *ENSG00000165807* | *PPP1R36* | -2,01 | 0,010972028 |
| *ENSG00000101230* | *ISM1* | -2,01 | 0,04697081 |
| *ENSG00000124762* | *CDKN1A* | -2,03 | 0,001421351 |
| *ENSG00000143590* | *EFNA3* | -2,03 | 0,002173295 |
| *ENSG00000117394* | *SLC2A1* | -2,03 | 0,000815553 |
| *ENSG00000187554* | *TLR5* | -2,04 | 0,025121523 |

| | | | |
|---|---|---|---|
| *ENSG00000118402* | *ELOVL4* | -2,04 | 0,036698139 |
| *ENSG00000008056* | *SYN1* | -2,05 | 0,000302175 |
| *ENSG00000159423* | *ALDH4A1* | -2,06 | 1,02099E-05 |
| *ENSG00000185909* | *KLHDC8B* | -2,06 | 0,00317631 |
| *ENSG00000253368* | *TRNP1* | -2,07 | 0,000889266 |
| *ENSG00000154217* | *PITPNC1* | -2,08 | 1,85157E-08 |
| *ENSG00000100889* | *PCK2* | -2,08 | 1,99041E-06 |
| *ENSG00000114771* | *AADAC* | -2,09 | 0,046519836 |
| *ENSG00000125148* | *MT2A* | -2,09 | 0,002330059 |
| *ENSG00000089486* | *CDIP1* | -2,09 | 6,08261E-06 |
| *ENSG00000187017* | *ESPN* | -2,11 | 0,005758882 |
| *ENSG00000151914* | *DST* | -2,12 | 3,28963E-06 |
| *ENSG00000166924* | *NYAP1* | -2,13 | 0,026035243 |
| *ENSG00000119862* | *LGALSL* | -2,13 | 0,000796218 |
| *ENSG00000179909* | *ZNF154* | -2,13 | 6,72719E-06 |
| *ENSG00000214711* | *CAPN14* | -2,14 | 0,038428679 |
| *ENSG00000214140* | *PRCD* | -2,14 | 0,045592497 |
| *ENSG00000198948* | *MFAP3L* | -2,15 | 0,035431058 |
| *ENSG00000119986* | *AVPI1* | -2,16 | 0,000308249 |
| *ENSG00000179859* | *RNF227* | -2,16 | 1,67958E-13 |
| *ENSG00000150782* | *IL18* | -2,17 | 3,59145E-05 |
| *ENSG00000166033* | *HTRA1* | -2,17 | 0,003188167 |
| *ENSG00000140807* | *NKD1* | -2,18 | 0,03069984 |
| *ENSG00000147465* | *STAR* | -2,18 | 0,023276982 |
| *ENSG00000175793* | *SFN* | -2,19 | 0,001922124 |
| *ENSG00000180113* | *TDRD6* | -2,20 | 0,000884179 |
| *ENSG00000189283* | *FHIT* | -2,20 | 0,005304609 |
| *ENSG00000205084* | *TMEM231* | -2,21 | 0,002046617 |
| *ENSG00000129038* | *LOXL1* | -2,22 | 0,000485836 |
| *ENSG00000150764* | *DIXDC1* | -2,22 | 0,001406425 |
| *ENSG00000153714* | *LURAP1L* | -2,23 | 0,000377535 |
| *ENSG00000104808* | *DHDH* | -2,23 | 0,035217679 |
| *ENSG00000170899* | *GSTA4* | -2,24 | 0,038406784 |
| *ENSG00000234465* | *PINLYP* | -2,25 | 0,004286545 |
| *ENSG00000166546* | *BEAN1* | -2,26 | 0,019809157 |
| *ENSG00000135549* | *PKIB* | -2,29 | 0,046587808 |
| *ENSG00000187908* | *DMBT1* | -2,29 | 0,041543877 |
| *ENSG00000078018* | *MAP2* | -2,30 | 0,019583932 |
| *ENSG00000138606* | *SHF* | -2,30 | 0,030481506 |
| *ENSG00000197632* | *SERPINB2* | -2,31 | 0,009569912 |
| *ENSG00000137449* | *CPEB2* | -2,33 | 0,001256601 |
| *ENSG00000101049* | *SGK2* | -2,33 | 0,043342591 |
| *ENSG00000001084* | *GCLC* | -2,34 | 0,000377535 |
| *ENSG00000151704* | *KCNJ1* | -2,36 | 0,036412913 |
| *ENSG00000103316* | *CRYM* | -2,37 | 0,001195715 |

| | | | |
|---|---|---|---|
| *ENSG00000267710* | *EDDM13* | -2,37 | 0,000123504 |
| *ENSG00000108352* | *RAPGEFL1* | -2,40 | 0,024235255 |
| *ENSG00000167992* | *VWCE* | -2,40 | 8,7949E-07 |
| *ENSG00000092421* | *SEMA6A* | -2,40 | 2,40042E-05 |
| *ENSG00000197953* | *AADACL2* | -2,41 | 0,039289343 |
| *ENSG00000253873* | *PCDHGA11* | -2,42 | 0,000201417 |
| *ENSG00000105523* | *FAM83E* | -2,45 | 0,043184046 |
| *ENSG00000103034* | *NDRG4* | -2,46 | 0,02940756 |
| *ENSG00000143369* | *ECM1* | -2,47 | 0,000202978 |
| *ENSG00000189410* | *SH2D5* | -2,50 | 0,003920447 |
| *ENSG00000163584* | *RPL22L1* | -2,51 | 3,74343E-06 |
| *ENSG00000114656* | *KIAA1257* | -2,51 | 0,000626894 |
| *ENSG00000101695* | *RNF125* | -2,52 | 0,021168691 |
| *ENSG00000164520* | *RAET1E* | -2,52 | 0,03069984 |
| *ENSG00000108309* | *RUNDC3A* | -2,53 | 0,014386182 |
| *ENSG00000173809* | *TDRD12* | -2,53 | 0,044548287 |
| *ENSG00000161911* | *TREML1* | -2,54 | 0,033781079 |
| *ENSG00000196754* | *S100A2* | -2,54 | 0,023967139 |
| *ENSG00000258405* | *ZNF578* | -2,57 | 0,001690927 |
| *ENSG00000213366* | *GSTM2* | -2,57 | 0,03434188 |
| *ENSG00000186648* | *CARMIL3* | -2,59 | 0,009388098 |
| *ENSG00000144285* | *SCN1A* | -2,59 | 0,049291972 |
| *ENSG00000085831* | *TTC39A* | -2,60 | 0,008566125 |
| *ENSG00000206075* | *SERPINB5* | -2,60 | 0,029184801 |
| *ENSG00000283632* | *EXOC3L2* | -2,61 | 0,002724789 |
| *ENSG00000096060* | *FKBP5* | -2,63 | 0,008710713 |
| *ENSG00000183943* | *PRKX* | -2,64 | 3,38427E-06 |
| *ENSG00000178980* | *SELENOW* | -2,64 | 1,39095E-20 |
| *ENSG00000170385* | *SLC30A1* | -2,66 | 8,78051E-08 |
| *ENSG00000198417* | *MT1F* | -2,66 | 0,020207646 |
| *ENSG00000111405* | *ENDOU* | -2,69 | 0,010422625 |
| *ENSG00000139433* | *GLTP* | -2,69 | 4,48572E-05 |
| *ENSG00000128918* | *ALDH1A2* | -2,72 | 0,000656412 |
| *ENSG00000130598* | *TNNI2* | -2,72 | 0,012621728 |
| *ENSG00000069812* | *HES2* | -2,74 | 0,027248297 |
| *ENSG00000117707* | *PROX1* | -2,74 | 0,000242876 |
| *ENSG00000166963* | *MAP1A* | -2,75 | 0,008682681 |
| *ENSG00000178573* | *MAF* | -2,76 | 0,013314303 |
| *ENSG00000064886* | *CHI3L2* | -2,76 | 7,00175E-05 |
| *ENSG00000187944* | *C2orf66* | -2,76 | 0,001842455 |
| *ENSG00000111863* | *ADTRP* | -2,77 | 0,000370365 |
| *ENSG00000187091* | *PLCD1* | -2,78 | 0,020605539 |
| *ENSG00000100285* | *NEFH* | -2,79 | 0,002498938 |
| *ENSG00000133256* | *PDE6B* | -2,80 | 0,004717083 |
| *ENSG00000189334* | *S100A14* | -2,81 | 0,035829762 |

| | | | |
|---|---|---|---|
| *ENSG00000122711* | *SPINK4* | -2,82 | 0,040180469 |
| *ENSG00000203722* | *RAET1G* | -2,83 | 0,010789936 |
| *ENSG00000186081* | *KRT5* | -2,83 | 0,035638902 |
| *ENSG00000178826* | *TMEM139* | -2,84 | 0,001489001 |
| *ENSG00000129455* | *KLK8* | -2,85 | 0,046732735 |
| *ENSG00000161544* | *CYGB* | -2,87 | 0,0330723 |
| *ENSG00000175600* | *SUGCT* | -2,88 | 0,000596315 |
| *ENSG00000120215* | *MLANA* | -2,89 | 0,00424666 |
| *ENSG00000108242* | *CYP2C18* | -2,89 | 0,038891894 |
| *ENSG00000140986* | *RPL3L* | -2,90 | 6,12766E-05 |
| *ENSG00000135919* | *SERPINE2* | -2,90 | 0,022431694 |
| *ENSG00000270885* | *RASL10B* | -2,90 | 1,95266E-05 |
| *ENSG00000131126* | *TEX101* | -2,93 | 0,005946517 |
| *ENSG00000253159* | *PCDHGA12* | -2,94 | 6,80426E-05 |
| *ENSG00000110195* | *FOLR1* | -2,94 | 0,031801545 |
| *ENSG00000021826* | *CPS1* | -2,98 | 0,029006841 |
| *ENSG00000172350* | *ABCG4* | -3,00 | 6,36194E-05 |
| *ENSG00000188828* | *GLRA4* | -3,01 | 0,000884179 |
| *ENSG00000007174* | *DNAH9* | -3,01 | 0,0330723 |
| *ENSG00000166183* | *ASPG* | -3,01 | 0,010241189 |
| *ENSG00000069667* | *RORA* | -3,01 | 0,003504406 |
| *ENSG00000135218* | *CD36* | -3,02 | 0,012445414 |
| *ENSG00000188783* | *PRELP* | -3,02 | 0,017877747 |
| *ENSG00000187479* | *C11orf96* | -3,02 | 0,015248836 |
| *ENSG00000186806* | *VSIG10L* | -3,03 | 0,009819197 |
| *ENSG00000163071* | *SPATA18* | -3,04 | 4,76618E-06 |
| *ENSG00000128422* | *KRT17* | -3,06 | 0,002046964 |
| *ENSG00000167617* | *CDC42EP5* | -3,08 | 0,035780904 |
| *ENSG00000182489* | *XKRX* | -3,08 | 0,010953764 |
| *ENSG00000144452* | *ABCA12* | -3,08 | 0,03343369 |
| *ENSG00000105880* | *DLX5* | -3,08 | 0,002580904 |
| *ENSG00000161249* | *DMKN* | -3,08 | 3,98553E-05 |
| *ENSG00000146070* | *PLA2G7* | -3,09 | 0,035711514 |
| *ENSG00000186115* | *CYP4F2* | -3,11 | 0,000916825 |
| *ENSG00000204420* | *MPIG6B* | -3,12 | 0,019037222 |
| *ENSG00000186847* | *KRT14* | -3,13 | 0,028214724 |
| *ENSG00000136010* | *ALDH1L2* | -3,13 | 0,000202978 |
| *ENSG00000166268* | *MYRFL* | -3,15 | 0,028607286 |
| *ENSG00000243708* | *PLA2G4B* | -3,15 | 0,001047776 |
| *ENSG00000132677* | *RHBG* | -3,16 | 0,027117171 |
| *ENSG00000181458* | *TMEM45A* | -3,17 | 0,000150122 |
| *ENSG00000184502* | *GAST* | -3,18 | 0,000545385 |
| *ENSG00000168060* | *NAALADL1* | -3,20 | 1,61875E-09 |
| *ENSG00000166396* | *SERPINB7* | -3,21 | 0,002238659 |
| *ENSG00000167751* | *KLK2* | -3,21 | 0,031201722 |

| | | | |
|---|---|---|---|
| *ENSG00000163207* | *IVL* | -3,22 | 0,033745386 |
| *ENSG00000169860* | *P2RY1* | -3,24 | 0,014904445 |
| *ENSG00000182040* | *USH1G* | -3,24 | 0,003396247 |
| *ENSG00000173947* | *PIFO* | -3,25 | 6,41205E-09 |
| *ENSG00000164893* | *SLC7A13* | -3,25 | 0,043010564 |
| *ENSG00000185052* | *SLC24A3* | -3,27 | 0,003073238 |
| *ENSG00000198483* | *ANKRD35* | -3,31 | 0,041498183 |
| *ENSG00000170044* | *ZPLD1* | -3,31 | 0,00022135 |
| *ENSG00000109321* | *AREG* | -3,32 | 0,002364294 |
| *ENSG00000173338* | *KCNK7* | -3,32 | 3,19993E-05 |
| *ENSG00000167641* | *PPP1R14A* | -3,33 | 0,029263002 |
| *ENSG00000175356* | *SCUBE2* | -3,34 | 9,2689E-05 |
| *ENSG00000133135* | *RNF128* | -3,34 | 0,006623608 |
| *ENSG00000129151* | *BBOX1* | -3,34 | 0,00173949 |
| *ENSG00000198028* | *ZNF560* | -3,34 | 0,005333151 |
| *ENSG00000168824* | *NSG1* | -3,35 | 0,000719749 |
| *ENSG00000158813* | *EDA* | -3,35 | 0,000430261 |
| *ENSG00000181333* | *HEPHL1* | -3,40 | 0,018519508 |
| *ENSG00000184060* | *ADAP2* | -3,41 | 1,8544E-05 |
| *ENSG00000137507* | *LRRC32* | -3,43 | 0,00277233 |
| *ENSG00000188100* | *FAM25A* | -3,45 | 0,022699826 |
| *ENSG00000135929* | *CYP27A1* | -3,46 | 1,84759E-07 |
| *ENSG00000186832* | *KRT16* | -3,46 | 0,020118764 |
| *ENSG00000102032* | *RENBP* | -3,46 | 5,26512E-06 |
| *ENSG00000205445* | *KRTAP10-2* | -3,48 | 0,03774343 |
| *ENSG00000167723* | *TRPV3* | -3,48 | 4,66276E-06 |
| *ENSG00000165474* | *GJB2* | -3,50 | 0,010585688 |
| *ENSG00000026751* | *SLAMF7* | -3,52 | 0,005667388 |
| *ENSG00000178882* | *RFLNA* | -3,52 | 0,011775751 |
| *ENSG00000275004* | *ZNF280B* | -3,52 | 0,027877061 |
| *ENSG00000152463* | *OLAH* | -3,52 | 0,011881555 |
| *ENSG00000128285* | *MCHR1* | -3,52 | 0,032277165 |
| *ENSG00000141579* | *ZNF750* | -3,52 | 0,016738667 |
| *ENSG00000143891* | *GALM* | -3,55 | 0,000731596 |
| *ENSG00000125571* | *IL37* | -3,55 | 0,001837674 |
| *ENSG00000010319* | *SEMA3G* | -3,58 | 0,00013033 |
| *ENSG00000157368* | *IL34* | -3,58 | 0,002879526 |
| *ENSG00000166118* | *SPATA19* | -3,59 | 0,032214426 |
| *ENSG00000149090* | *PAMR1* | -3,59 | 0,012597354 |
| *ENSG00000136697* | *IL1F10* | -3,59 | 0,043823488 |
| *ENSG00000145192* | *AHSG* | -3,63 | 0,000314701 |
| *ENSG00000131080* | *EDA2R* | -3,64 | 0,013632135 |
| *ENSG00000100368* | *CSF2RB* | -3,64 | 0,016459791 |
| *ENSG00000162643* | *WDR63* | -3,64 | 1,6057E-10 |
| *ENSG00000267795* | *SMIM22* | -3,64 | 0,004065416 |

| | | | |
|---|---|---|---|
| *ENSG00000109846* | *CRYAB* | -3,66 | 0,04069547 |
| *ENSG00000196542* | *SPTSSB* | -3,66 | 0,024342159 |
| *ENSG00000186395* | *KRT10* | -3,67 | 2,14949E-08 |
| *ENSG00000152049* | *KCNE4* | -3,68 | 0,003722871 |
| *ENSG00000163328* | *GPR155* | -3,70 | 4,55732E-07 |
| *ENSG00000074410* | *CA12* | -3,70 | 0,013485635 |
| *ENSG00000133665* | *DYDC2* | -3,70 | 0,033973199 |
| *ENSG00000187193* | *MT1X* | -3,71 | 2,25545E-06 |
| *ENSG00000156298* | *TSPAN7* | -3,75 | 0,002310904 |
| *ENSG00000171476* | *HOPX* | -3,76 | 0,013671407 |
| *ENSG00000183801* | *OLFML1* | -3,78 | 0,017534866 |
| *ENSG00000095970* | *TREM2* | -3,81 | 0,030152929 |
| *ENSG00000179057* | *IGSF22* | -3,81 | 0,001709821 |
| *ENSG00000270168* | | -3,82 | 4,89282E-05 |
| *ENSG00000204538* | *PSORS1C2* | -3,85 | 0,000889266 |
| *ENSG00000090512* | *FETUB* | -3,86 | 0,000394805 |
| *ENSG00000155966* | *AFF2* | -3,87 | 0,012723096 |
| *ENSG00000283227* | *SPRR5* | -3,87 | 0,04974496 |
| *ENSG00000088386* | *SLC15A1* | -3,88 | 0,034950107 |
| *ENSG00000103056* | *SMPD3* | -3,89 | 0,000187247 |
| *ENSG00000137265* | *IRF4* | -3,90 | 0,012572684 |
| *ENSG00000167748* | *KLK1* | -3,90 | 0,007119877 |
| *ENSG00000189090* | *FAM25G* | -3,90 | 0,008423753 |
| *ENSG00000111218* | *PRMT8* | -3,90 | 0,024622718 |
| *ENSG00000092295* | *TGM1* | -3,91 | 2,68646E-05 |
| *ENSG00000115138* | *POMC* | -3,91 | 0,001669316 |
| *ENSG00000135312* | *HTR1B* | -3,92 | 0,029155198 |
| *ENSG00000143387* | *CTSK* | -3,92 | 1,61496E-05 |
| *ENSG00000136695* | *IL36RN* | -3,93 | 0,000759887 |
| *ENSG00000112964* | *GHR* | -3,93 | 0,016502732 |
| *ENSG00000123360* | *PDE1B* | -3,95 | 0,005860221 |
| *ENSG00000109956* | *B3GAT1* | -3,97 | 0,001234014 |
| *ENSG00000250722* | *SELENOP* | -3,98 | 0,017546363 |
| *ENSG00000010438* | *PRSS3* | -3,98 | 0,000884179 |
| *ENSG00000123689* | *G0S2* | -4,00 | 0,032468623 |
| *ENSG00000205108* | *FAM205A* | -4,02 | 0,020118764 |
| *ENSG00000115590* | *IL1R2* | -4,03 | 0,000134427 |
| *ENSG00000177098* | *SCN4B* | -4,03 | 0,004609141 |
| *ENSG00000183439* | *TRIM61* | -4,03 | 0,026438883 |
| *ENSG00000211445* | *GPX3* | -4,04 | 3,23147E-05 |
| *ENSG00000163331* | *DAPL1* | -4,05 | 0,025814334 |
| *ENSG00000189320* | *FAM180A* | -4,05 | 0,000106017 |
| *ENSG00000168918* | *INPP5D* | -4,06 | 0,020403494 |
| *ENSG00000133710* | *SPINK5* | -4,07 | 0,005112791 |
| *ENSG00000121552* | *CSTA* | -4,08 | 0,000149694 |

| | | | |
|---|---|---|---|
| *ENSG00000177301* | *KCNA2* | -4,09 | 0,031371751 |
| *ENSG00000172828* | *CES3* | -4,11 | 1,80541E-06 |
| *ENSG00000142549* | *IGLON5* | -4,11 | 0,015647653 |
| *ENSG00000100146* | *SOX10* | -4,12 | 0,000550095 |
| *ENSG00000112981* | *NME5* | -4,13 | 0,002621458 |
| *ENSG00000198865* | *CCDC152* | -4,13 | 0,027369689 |
| *ENSG00000151117* | *TMEM86A* | -4,14 | 0,000544163 |
| *ENSG00000123243* | *ITIH5* | -4,15 | 0,025626204 |
| *ENSG00000130203* | *APOE* | -4,15 | 0,001040725 |
| *ENSG00000167757* | *KLK11* | -4,17 | 0,004012249 |
| *ENSG00000204632* | *HLA-G* | -4,19 | 2,3644E-05 |
| *ENSG00000149131* | *SERPING1* | -4,19 | 0,008632834 |
| *ENSG00000204021* | *LIPK* | -4,19 | 0,003396247 |
| *ENSG00000172782* | *FADS6* | -4,22 | 0,015488274 |
| *ENSG00000198092* | *TMPRSS11F* | -4,22 | 0,002163669 |
| *ENSG00000204866* | *IGFL2* | -4,23 | 0,047976509 |
| *ENSG00000204542* | *C6orf15* | -4,24 | 0,03957766 |
| *ENSG00000105549* | *THEG* | -4,25 | 0,002196128 |
| *ENSG00000159398* | *CES5A* | -4,25 | 0,021979424 |
| *ENSG00000205364* | *MT1M* | -4,29 | 0,015478836 |
| *ENSG00000136689* | *IL1RN* | -4,30 | 0,000631017 |
| *ENSG00000110203* | *FOLR3* | -4,30 | 0,007103988 |
| *ENSG00000198854* | *C1orf68* | -4,32 | 0,025514429 |
| *ENSG00000124019* | *FAM124B* | -4,36 | 0,001712326 |
| *ENSG00000109610* | *SOD3* | -4,36 | 0,010376197 |
| *ENSG00000102409* | *BEX4* | -4,37 | 0,016755351 |
| *ENSG00000269113* | *TRABD2B* | -4,38 | 0,002540454 |
| *ENSG00000179388* | *EGR3* | -4,40 | 0,000865732 |
| *ENSG00000106819* | *ASPN* | -4,42 | 2,21515E-05 |
| *ENSG00000089250* | *NOS1* | -4,43 | 0,033657488 |
| *ENSG00000168621* | *GDNF* | -4,43 | 0,012872244 |
| *ENSG00000175264* | *CHST1* | -4,44 | 0,004047523 |
| *ENSG00000134827* | *TCN1* | -4,45 | 0,008176989 |
| *ENSG00000177238* | *TRIM72* | -4,46 | 0,00118362 |
| *ENSG00000175121* | *WFDC5* | -4,46 | 0,000545385 |
| *ENSG00000185615* | *PDIA2* | -4,49 | 0,000629158 |
| *ENSG00000188000* | *OR7D2* | -4,55 | 0,045161654 |
| *ENSG00000094796* | *KRT31* | -4,57 | 0,018151967 |
| *ENSG00000126733* | *DACH2* | -4,59 | 0,005131538 |
| *ENSG00000172551* | *MUCL1* | -4,70 | 0,001918193 |
| *ENSG00000185633* | *NDUFA4L2* | -4,71 | 3,09954E-09 |
| *ENSG00000149968* | *MMP3* | -4,73 | 0,013021193 |
| *ENSG00000169035* | *KLK7* | -4,73 | 0,003931691 |
| *ENSG00000108018* | *SORCS1* | -4,75 | 0,017536164 |
| *ENSG00000126233* | *SLURP1* | -4,77 | 0,002680651 |

| | | | |
|---|---|---|---|
| *ENSG00000093072* | *ADA2* | -4,78 | 3,36524E-15 |
| *ENSG00000180316* | *PNPLA1* | -4,78 | 0,000223669 |
| *ENSG00000143631* | *FLG* | -4,82 | 0,000681694 |
| *ENSG00000168447* | *SCNN1B* | -4,83 | 0,017128771 |
| *ENSG00000174145* | *NWD2* | -4,83 | 0,000293572 |
| *ENSG00000167768* | *KRT1* | -4,85 | 0,018264564 |
| *ENSG00000166535* | *A2ML1* | -4,88 | 0,002046617 |
| *ENSG00000087076* | *HSD17B14* | -4,88 | 9,9204E-08 |
| *ENSG00000185640* | *KRT79* | -4,90 | 0,009792692 |
| *ENSG00000268104* | *SLC6A14* | -4,91 | 0,000814569 |
| *ENSG00000161281* | *COX7A1* | -4,91 | 1,30763E-18 |
| *ENSG00000132746* | *ALDH3B2* | -4,92 | 1,30392E-05 |
| *ENSG00000182983* | *ZNF662* | -4,93 | 0,023494137 |
| *ENSG00000204252* | *HLA-DOA* | -4,93 | 2,86098E-05 |
| *ENSG00000138152* | *BTBD16* | -4,94 | 0,005092331 |
| *ENSG00000178947* | *SMIM10L2A* | -4,98 | 1,09911E-09 |
| *ENSG00000276430* | *FAM25C* | -4,98 | 0,002619955 |
| *ENSG00000283703* | *VSIG10L2* | -4,99 | 0,001071367 |
| *ENSG00000214652* | *ZNF727* | -5,01 | 0,000784841 |
| *ENSG00000184459* | *BPIFC* | -5,02 | 0,034029042 |
| *ENSG00000204540* | *PSORS1C1* | -5,02 | 6,47671E-06 |
| *ENSG00000168907* | *PLA2G4F* | -5,03 | 1,55474E-06 |
| *ENSG00000088002* | *SULT2B1* | -5,06 | 0,000966196 |
| *ENSG00000174697* | *LEP* | -5,06 | 0,020403494 |
| *ENSG00000173239* | *LIPM* | -5,09 | 0,000198626 |
| *ENSG00000164825* | *DEFB1* | -5,12 | 0,000372025 |
| *ENSG00000152207* | *CYSLTR2* | -5,17 | 0,018201928 |
| *ENSG00000204020* | *LIPN* | -5,20 | 0,000942692 |
| *ENSG00000141485* | *SLC13A5* | -5,21 | 1,0118E-05 |
| *ENSG00000167759* | *KLK13* | -5,22 | 0,027038348 |
| *ENSG00000158022* | *TRIM63* | -5,23 | 0,00647837 |
| *ENSG00000188293* | *IGFL1* | -5,25 | 0,000489864 |
| *ENSG00000164756* | *SLC30A8* | -5,27 | 0,000333604 |
| *ENSG00000188803* | *SHISA6* | -5,27 | 0,008632834 |
| *ENSG00000145283* | *SLC10A6* | -5,28 | 0,008005526 |
| *ENSG00000100078* | *PLA2G3* | -5,28 | 5,58351E-06 |
| *ENSG00000178187* | *ZNF454* | -5,30 | 2,6034E-05 |
| *ENSG00000183273* | *CCDC60* | -5,30 | 0,001538112 |
| *ENSG00000263429* | *TMEM238L* | -5,30 | 8,65049E-05 |
| *ENSG00000170465* | *KRT6C* | -5,30 | 0,006280437 |
| *ENSG00000189001* | *SBSN* | -5,30 | 0,004783016 |
| *ENSG00000174502* | *SLC26A9* | -5,30 | 0,00015596 |
| *ENSG00000188508* | *KRTDAP* | -5,32 | 0,006808151 |
| *ENSG00000182261* | *NLRP10* | -5,35 | 0,005479872 |
| *ENSG00000078725* | *BRINP1* | -5,37 | 0,012572684 |

| | | | |
|---|---|---|---|
| *ENSG00000162040* | *HS3ST6* | -5,40 | 0,004012249 |
| *ENSG00000172867* | *KRT2* | -5,41 | 0,005245584 |
| *ENSG00000204421* | *LY6G6C* | -5,41 | 0,000144964 |
| *ENSG00000121898* | *CPXM2* | -5,42 | 0,015314857 |
| *ENSG00000168490* | *PHYHIP* | -5,44 | 4,40691E-14 |
| *ENSG00000143627* | *PKLR* | -5,45 | 1,2154E-07 |
| *ENSG00000078596* | *ITM2A* | -5,47 | 0,032159486 |
| *ENSG00000108244* | *KRT23* | -5,48 | 0,00140907 |
| *ENSG00000122176* | *FMOD* | -5,51 | 0,010254315 |
| *ENSG00000163141* | *BNIPL* | -5,53 | 2,52045E-11 |
| *ENSG00000134765* | *DSC1* | -5,54 | 2,4277E-05 |
| *ENSG00000174807* | *CD248* | -5,59 | 0,000274696 |
| *ENSG00000160539* | *PLPP7* | -5,61 | 0,048984364 |
| *ENSG00000144649* | *FAM198A* | -5,64 | 3,14981E-05 |
| *ENSG00000204539* | *CDSN* | -5,65 | 1,80541E-06 |
| *ENSG00000105141* | *CASP14* | -5,67 | 0,00107813 |
| *ENSG00000203837* | *PNLIPRP3* | -5,73 | 0,000307738 |
| *ENSG00000205076* | *LGALS7* | -5,73 | 0,005119937 |
| *ENSG00000137033* | *IL33* | -5,76 | 8,60812E-07 |
| *ENSG00000205592* | *MUC19* | -5,90 | 5,49575E-09 |
| *ENSG00000107165* | *TYRP1* | -5,94 | 4,89282E-05 |
| *ENSG00000170426* | *SDR9C7* | -5,94 | 0,000499325 |
| *ENSG00000186226* | *LCE1E* | -5,94 | 0,003048792 |
| *ENSG00000089472* | *HEPH* | -5,97 | 2,54262E-06 |
| *ENSG00000084674* | *APOB* | -5,98 | 0,004560355 |
| *ENSG00000159337* | *PLA2G4D* | -5,99 | 4,87927E-10 |
| *ENSG00000145423* | *SFRP2* | -6,00 | 0,030090572 |
| *ENSG00000189182* | *KRT77* | -6,05 | 0,000669044 |
| *ENSG00000178934* | *LGALS7B* | -6,08 | 0,002989591 |
| *ENSG00000166589* | *CDH16* | -6,12 | 8,18712E-14 |
| *ENSG00000109906* | *ZBTB16* | -6,13 | 0,023815802 |
| *ENSG00000176046* | *NUPR1* | -6,16 | 7,44985E-09 |
| *ENSG00000109182* | *CWH43* | -6,19 | 0,000180268 |
| *ENSG00000205363* | *INSYN1* | -6,23 | 7,71865E-05 |
| *ENSG00000158014* | *SLC30A2* | -6,27 | 0,000657552 |
| *ENSG00000163464* | *CXCR1* | -6,30 | 1,75114E-05 |
| *ENSG00000204764* | *RANBP17* | -6,36 | 2,89112E-05 |
| *ENSG00000075673* | *ATP12A* | -6,38 | 4,16271E-06 |
| *ENSG00000139988* | *RDH12* | -6,46 | 6,01051E-10 |
| *ENSG00000143365* | *RORC* | -6,47 | 2,11051E-07 |
| *ENSG00000197614* | *MFAP5* | -6,62 | 0,000105318 |
| *ENSG00000079393* | *DUSP13* | -6,63 | 7,81159E-09 |
| *ENSG00000116132* | *PRRX1* | -6,67 | 0,000657145 |
| *ENSG00000118520* | *ARG1* | -6,69 | 1,32661E-07 |
| *ENSG00000113805* | *CNTN3* | -6,77 | 4,15187E-06 |

| | | | |
|---|---|---|---|
| *ENSG00000240747* | *KRBOX1* | -6,78 | 1,61875E-09 |
| *ENSG00000133048* | *CHI3L1* | -6,86 | 7,81196E-10 |
| *ENSG00000182916* | *TCEAL7* | -6,93 | 1,84759E-07 |
| *ENSG00000167656* | *LY6D* | -6,94 | 3,7536E-08 |
| *ENSG00000094963* | *FMO2* | -7,00 | 0,000515978 |
| *ENSG00000100170* | *SLC5A1* | -7,01 | 3,5755E-09 |
| *ENSG00000121440* | *PDZRN3* | -7,02 | 1,91614E-05 |
| *ENSG00000095627* | *TDRD1* | -7,06 | 6,62363E-06 |
| *ENSG00000240563* | *L1TD1* | -7,16 | 1,31287E-09 |
| *ENSG00000158786* | *PLA2G2F* | -7,19 | 0,000184928 |
| *ENSG00000171954* | *CYP4F22* | -7,23 | 0,00010595 |
| *ENSG00000255501* | *CARD18* | -7,38 | 2,31343E-06 |
| *ENSG00000105131* | *EPHX3* | -7,39 | 2,27744E-17 |
| *ENSG00000188393* | *CLEC2A* | -7,50 | 0,000186794 |
| *ENSG00000185962* | *LCE3A* | -7,65 | 0,00481982 |
| *ENSG00000265107* | *GJA5* | -7,69 | 9,55837E-10 |
| *ENSG00000167914* | *GSDMA* | -7,80 | 1,7793E-09 |
| *ENSG00000169509* | *CRCT1* | -7,99 | 3,1306E-05 |
| *ENSG00000203782* | *LOR* | -8,10 | 2,42271E-10 |
| *ENSG00000143520* | *FLG2* | -8,28 | 1,6057E-10 |
| *ENSG00000185966* | *LCE3E* | -8,65 | 9,77354E-07 |
| *ENSG00000163202* | *LCE3D* | -8,71 | 2,04631E-06 |
| *ENSG00000169900* | *PYDC1* | -8,74 | 0,000280117 |
| *ENSG00000160307* | *S100B* | -8,74 | 4,45457E-05 |
| *ENSG00000103569* | *AQP9* | -8,75 | 1,17879E-09 |
| *ENSG00000179593* | *ALOX15B* | -8,87 | 1,21193E-16 |
| *ENSG00000080166* | *DCT* | -8,92 | 1,6057E-10 |
| *ENSG00000077498* | *TYR* | -8,94 | 6,7335E-07 |
| *ENSG00000187223* | *LCE2D* | -9,02 | 4,3112E-07 |
| *ENSG00000147896* | *IFNK* | -9,04 | 1,33301E-32 |
| *ENSG00000167769* | *ACER1* | -9,22 | 1,50651E-09 |
| *ENSG00000159455* | *LCE2B* | -9,26 | 5,0284E-08 |
| *ENSG00000203786* | *KPRP* | -9,30 | 1,45163E-08 |
| *ENSG00000240386* | *LCE1F* | -9,55 | 2,86369E-10 |
| *ENSG00000196734* | *LCE1B* | -9,61 | 4,9824E-15 |
| *ENSG00000197084* | *LCE1C* | -9,64 | 2,24659E-10 |
| *ENSG00000187173* | *LCE2A* | -9,66 | 1,51359E-06 |
| *ENSG00000186844* | *LCE1A* | -9,79 | 1,23279E-06 |
| *ENSG00000198046* | *ZNF667* | -9,82 | 2,00379E-21 |
| *ENSG00000188624* | *IGFL3* | -9,87 | 3,45322E-14 |
| *ENSG00000187180* | *LCE2C* | -9,99 | 1,94594E-09 |
| *ENSG00000172155* | *LCE1D* | -10,49 | 3,48886E-07 |
| *ENSG00000125144* | *MT1G* | -10,52 | 1,63949E-34 |
| *ENSG00000235942* | *LCE6A* | -11,10 | 5,04661E-10 |
| *ENSG00000205358* | *MT1H* | -11,40 | 2,40309E-24 |

**Supplement table S3.** Average correlations of the clusters.

| Cluster | No of genes | No of protein coding genes | No of lncRNAs | Average correlation |
|---|---|---|---|---|
| 1 | 611 | 403 | 208 | 0,439 |
| 2 | 954 | 594 | 360 | 0,612 |
| 3 | 190 | 151 | 39 | 0,609 |
| 4 | 265 | 225 | 40 | 0,681 |
| 5 | 61 | 48 | 13 | 0,351 |
| 6 | 309 | 246 | 63 | 0,771 |

**Supplement table S4.** Summary of significantly regulated functional annotations overrepresented among cluster 1 genes. Cluster 1 genes were used for gene ontology (GO) analysis with DAVID.

| Category | Term | PValue |
|---|---|---|
| UP_SEQ_FEATURE | TOPO_DOM:Extracellular | 6.85E-7 |
| UP_KW_CELLULAR_COMPONENT | KW-1003~Cell membrane | 1.86E-6 |
| GOTERM_BP_DIRECT | GO:0007169~transmembrane receptor protein tyrosine kinase signaling pathway | 6.05E-4 |
| GOTERM_BP_DIRECT | GO:2000249~regulation of actin cytoskeleton reorganization | 7.93E-4 |
| GOTERM_BP_DIRECT | GO:0061098~positive regulation of protein tyrosine kinase activity | 0.001 |
| GOTERM_BP_DIRECT | GO:0033674~positive regulation of kinase activity | 0.002 |
| UP_KW_BIOLOGICAL_PROCESS | KW-0130~Cell adhesion | 0.004 |
| GOTERM_CC_DIRECT | GO:0005615~extracellular space | 0.004 |
| GOTERM_MF_DIRECT | GO:0004175~endopeptidase activity | 0.006 |
| GOTERM_CC_DIRECT | GO:0005576~extracellular region | 0.006 |
| GOTERM_BP_DIRECT | GO:0016477~cell migration | 0.009 |
| GOTERM_BP_DIRECT | GO:0022617~extracellular matrix disassembly | 0.009 |
| GOTERM_BP_DIRECT | GO:0030335~positive regulation of cell migration | 0.017 |
| GOTERM_BP_DIRECT | GO:0044344~cellular response to fibroblast growth factor stimulus | 0.021 |
| GOTERM_MF_DIRECT | GO:0004714~transmembrane receptor protein tyrosine kinase activity | 0.022 |
| UP_KW_MOLECULAR_FUNCTION | KW-0829~Tyrosine-protein kinase | 0.022 |

| | | |
|---|---|---|
| GOTERM_BP_DIRECT | GO:0043410~positive regulation of MAPK cascade | 0.025 |
| GOTERM_BP_DIRECT | GO:0072659~protein localization to plasma membrane | 0.025 |

**Supplement table S5.** LncRNA and mRNA genes involved in cluster 1.

| Ensembl.ID | HGNC.symbol | Average.expression | Log2.foldchange | Adjusted.p.value |
|---|---|---|---|---|
| *ENSG00000185905* | *C16orf54* | 190,2 | 24,9 | 9,62004E-14 |
| *ENSG00000163286* | *ALPG* | 62,0 | 23,4 | 2,42271E-10 |
| *ENSG00000137745* | *MMP13* | 3401,1 | 10,4 | 1,30228E-08 |
| *ENSG00000173391* | *OLR1* | 1978,3 | 7,1 | 1,79942E-07 |
| *ENSG00000147381* | *MAGEA4* | 2700,5 | 9,3 | 8,77192E-07 |
| *ENSG00000142185* | *TRPM2* | 384,3 | 4,5 | 1,04922E-06 |
| *ENSG00000205403* | *CFI* | 351,8 | 7,8 | 4,42131E-06 |
| *ENSG00000166415* | *WDR72* | 374,5 | 10,1 | 6,51162E-06 |
| *ENSG00000197142* | *ACSL5* | 598,2 | 5,7 | 6,51162E-06 |
| *ENSG00000168874* | *ATOH8* | 721,6 | 7,6 | 1,03813E-05 |
| *ENSG00000162654* | *GBP4* | 1293,6 | 6,4 | 1,10873E-05 |
| *ENSG00000132182* | *NUP210* | 1344,2 | 6,2 | 1,97969E-05 |
| *ENSG00000196104* | *SPOCK3* | 90,8 | 7,4 | 2,47636E-05 |
| *ENSG00000163283* | *ALPP* | 1076,0 | 9,5 | 2,50805E-05 |
| *ENSG00000205436* | *EXOC3L4* | 89,4 | 6,8 | 3,3301E-05 |
| *ENSG00000187689* | *AMTN* | 133,7 | 6,3 | 4,61408E-05 |
| *ENSG00000164292* | *RHOBTB3* | 4824,1 | 3,7 | 8,24118E-05 |
| *ENSG00000198443* | *KRTAP4-1* | 14,2 | 6,7 | 0,000134514 |
| *ENSG00000172915* | *NBEA* | 211,6 | 4,8 | 0,000144481 |
| *ENSG00000254221* | *PCDHGB1* | 67,0 | 5,0 | 0,000145216 |
| *ENSG00000204176* | *SYT15* | 130,8 | 3,6 | 0,000150122 |
| *ENSG00000163083* | *INHBB* | 190,0 | 7,9 | 0,000166291 |
| *ENSG00000089558* | *KCNH4* | 46,5 | 7,7 | 0,000234383 |
| *ENSG00000145113* | *MUC4* | 667,0 | 7,2 | 0,000242877 |
| *ENSG00000141574* | *SECTM1* | 696,3 | 3,3 | 0,000295536 |
| *ENSG00000140323* | *DISP2* | 361,1 | 2,9 | 0,000324565 |
| *ENSG00000023445* | *BIRC3* | 1462,2 | 4,1 | 0,000355892 |
| *ENSG00000106012* | *IQCE* | 2048,5 | 1,3 | 0,00037689 |
| *ENSG00000101986* | *ABCD1* | 731,8 | 2,9 | 0,000390753 |
| *ENSG00000260027* | *HOXB7* | 476,4 | 3,1 | 0,000409342 |
| *ENSG00000141448* | *GATA6* | 230,6 | 3,1 | 0,000427047 |
| *ENSG00000263961* | *RHEX* | 177,1 | 9,7 | 0,00043091 |
| *ENSG00000116833* | *NR5A2* | 38,4 | 5,4 | 0,000450856 |
| *ENSG00000134258* | *VTCN1* | 194,9 | 7,1 | 0,000482924 |
| *ENSG00000042980* | *ADAM28* | 79,6 | 7,5 | 0,000518531 |
| *ENSG00000144229* | *THSD7B* | 216,5 | 5,9 | 0,000542516 |

| | | | | |
|---|---|---|---|---|
| *ENSG00000165071* | *TMEM71* | 44,1 | 5,3 | 0,0005652 |
| *ENSG00000175463* | *TBC1D10C* | 22,5 | 2,6 | 0,000580902 |
| *ENSG00000159217* | *IGF2BP1* | 578,1 | 6,6 | 0,000641436 |
| *ENSG00000176485* | *PLA2G16* | 1626,5 | 5,1 | 0,000733917 |
| *ENSG00000175294* | *CATSPER1* | 48,6 | 5,0 | 0,000736183 |
| *ENSG00000275713* | *HIST1H2BH* | 121,3 | 2,2 | 0,000744711 |
| *ENSG00000148942* | *SLC5A12* | 28,2 | 6,9 | 0,000745927 |
| *ENSG00000055813* | *CCDC85A* | 173,6 | 7,2 | 0,000748575 |
| *ENSG00000156966* | *B3GNT7* | 134,0 | 4,3 | 0,00076408 |
| *ENSG00000029534* | *ANK1* | 940,5 | 4,4 | 0,00077714 |
| *ENSG00000257138* | *TAS2R38* | 16,2 | 6,1 | 0,000823404 |
| *ENSG00000116785* | *CFHR3* | 19,9 | 7,2 | 0,000873943 |
| *ENSG00000165124* | *SVEP1* | 277,1 | 3,8 | 0,000881353 |
| *ENSG00000160183* | *TMPRSS3* | 72,8 | 5,7 | 0,000911334 |
| *ENSG00000099994* | *SUSD2* | 200,6 | 4,9 | 0,000921546 |
| *ENSG00000064692* | *SNCAIP* | 200,0 | 5,8 | 0,000921546 |
| *ENSG00000006534* | *ALDH3B1* | 986,9 | 4,0 | 0,000921546 |
| *ENSG00000213512* | *GBP7* | 12,9 | 5,9 | 0,00092882 |
| *ENSG00000168754* | *FAM178B* | 278,0 | 4,6 | 0,001000722 |
| *ENSG00000185215* | *TNFAIP2* | 6436,6 | 4,1 | 0,001057491 |
| *ENSG00000184985* | *SORCS2* | 666,7 | 4,2 | 0,001061946 |
| *ENSG00000157303* | *SUSD3* | 90,1 | 4,3 | 0,001107597 |
| *ENSG00000152430* | *BOLL* | 13,7 | 4,7 | 0,00114902 |
| *ENSG00000273706* | *LHX1* | 253,6 | 6,0 | 0,00123908 |
| *ENSG00000134198* | *TSPAN2* | 305,3 | 4,3 | 0,001267809 |
| *ENSG00000268916* | *CSAG3* | 27,0 | 7,6 | 0,001352421 |
| *ENSG00000163121* | *NEURL3* | 89,3 | 4,3 | 0,001370503 |
| *ENSG00000205517* | *RGL3* | 132,3 | 5,7 | 0,001376677 |
| *ENSG00000118785* | *SPP1* | 691,4 | 6,2 | 0,001392486 |
| *ENSG00000145526* | *CDH18* | 88,7 | 6,4 | 0,00140907 |
| *ENSG00000165215* | *CLDN3* | 21,9 | 6,5 | 0,001421351 |
| *ENSG00000106868* | *SUSD1* | 355,7 | 1,8 | 0,001421351 |
| *ENSG00000075340* | *ADD2* | 512,7 | 6,4 | 0,001463929 |
| *ENSG00000163554* | *SPTA1* | 16,6 | 6,9 | 0,001472311 |
| *ENSG00000125347* | *IRF1* | 2253,7 | 2,6 | 0,001481436 |
| *ENSG00000096996* | *IL12RB1* | 81,4 | 4,7 | 0,001533009 |
| *ENSG00000105464* | *GRIN2D* | 265,5 | 4,0 | 0,001573307 |
| *ENSG00000139865* | *TTC6* | 21,2 | 4,7 | 0,001692221 |
| *ENSG00000172159* | *FRMD3* | 368,6 | 4,7 | 0,00170765 |
| *ENSG00000092929* | *UNC13D* | 977,7 | 3,2 | 0,001709821 |
| *ENSG00000162896* | *PIGR* | 50,8 | 6,8 | 0,001750014 |
| *ENSG00000115468* | *EFHD1* | 17,6 | 6,2 | 0,001783709 |
| *ENSG00000182791* | *CCDC87* | 47,5 | 2,7 | 0,00178724 |
| *ENSG00000124657* | *OR2B6* | 11,9 | 5,2 | 0,00181694 |
| *ENSG00000134780* | *DAGLA* | 210,9 | 3,9 | 0,001901514 |

| | | | | |
|---|---|---|---|---|
| *ENSG00000273983* | *HIST1H3G* | 61,6 | 2,9 | 0,001947213 |
| *ENSG00000128284* | *APOL3* | 786,5 | 4,1 | 0,002196585 |
| *ENSG00000168961* | *LGALS9* | 213,1 | 3,8 | 0,002238906 |
| *ENSG00000197724* | *PHF2* | 2545,0 | 1,2 | 0,002324008 |
| *ENSG00000112761* | *WISP3* | 392,8 | 3,3 | 0,002340307 |
| *ENSG00000146674* | *IGFBP3* | 48241,2 | 5,2 | 0,002352012 |
| *ENSG00000110057* | *UNC93B1* | 1497,0 | 1,9 | 0,002375868 |
| *ENSG00000182621* | *PLCB1* | 276,4 | 3,8 | 0,002410169 |
| *ENSG00000043355* | *ZIC2* | 48,9 | 6,0 | 0,002434295 |
| *ENSG00000178568* | *ERBB4* | 20,1 | 5,4 | 0,002548684 |
| *ENSG00000182759* | *MAFA* | 17,9 | 5,3 | 0,002642565 |
| *ENSG00000182836* | *PLCXD3* | 103,0 | 7,5 | 0,00267446 |
| *ENSG00000129682* | *FGF13* | 52,5 | 6,3 | 0,00268043 |
| *ENSG00000164176* | *EDIL3* | 767,1 | 4,9 | 0,002772108 |
| *ENSG00000196597* | *ZNF782* | 181,9 | 1,1 | 0,003075389 |
| *ENSG00000184058* | *TBX1* | 353,1 | 3,3 | 0,003136739 |
| *ENSG00000106025* | *TSPAN12* | 121,9 | 3,6 | 0,003165071 |
| *ENSG00000170577* | *SIX2* | 86,4 | 5,9 | 0,003177973 |
| *ENSG00000198929* | *NOS1AP* | 106,3 | 2,5 | 0,003181824 |
| *ENSG00000221843* | *C2orf16* | 141,9 | 1,0 | 0,003182631 |
| *ENSG00000129514* | *FOXA1* | 256,4 | 4,6 | 0,00326163 |
| *ENSG00000178764* | *ZHX2* | 782,9 | 1,0 | 0,003905908 |
| *ENSG00000146955* | *RAB19* | 22,8 | 5,7 | 0,004031257 |
| *ENSG00000125355* | *TMEM255A* | 318,1 | 4,4 | 0,004071419 |
| *ENSG00000186603* | *HPDL* | 163,9 | 3,2 | 0,004120155 |
| *ENSG00000131941* | *RHPN2* | 775,8 | 1,9 | 0,00416921 |
| *ENSG00000137261* | *KIAA0319* | 38,0 | 4,3 | 0,004221593 |
| *ENSG00000130045* | *NXNL2* | 74,8 | 3,4 | 0,004226828 |
| *ENSG00000178878* | *APOLD1* | 255,3 | 2,0 | 0,004254812 |
| *ENSG00000121653* | *MAPK8IP1* | 306,6 | 2,2 | 0,00427531 |
| *ENSG00000117791* | *MARC2* | 281,5 | 2,3 | 0,004289784 |
| *ENSG00000159184* | *HOXB13* | 78,9 | 6,8 | 0,004372167 |
| *ENSG00000130829* | *DUSP9* | 513,1 | 4,3 | 0,004378582 |
| *ENSG00000177465* | *ACOT4* | 120,9 | 2,0 | 0,004415292 |
| *ENSG00000164304* | *CAGE1* | 10,4 | 2,2 | 0,004560355 |
| *ENSG00000165507* | *DEPP1* | 294,7 | 2,6 | 0,004581423 |
| *ENSG00000106077* | *ABHD11* | 733,7 | 1,6 | 0,004593779 |
| *ENSG00000183072* | *NKX2-5* | 91,7 | 8,3 | 0,004602127 |
| *ENSG00000221890* | *NPTXR* | 289,4 | 3,4 | 0,004698533 |
| *ENSG00000143512* | *HHIPL2* | 70,9 | 4,6 | 0,00476491 |
| *ENSG00000171956* | *FOXB1* | 67,5 | 6,5 | 0,00481982 |
| *ENSG00000011677* | *GABRA3* | 214,2 | 5,6 | 0,004876664 |
| *ENSG00000004468* | *CD38* | 34,3 | 4,0 | 0,004953444 |
| *ENSG00000121797* | *CCRL2* | 13,7 | 4,9 | 0,004985654 |
| *ENSG00000115896* | *PLCL1* | 28,4 | 4,0 | 0,0053174 |

| | | | | |
|---|---|---|---|---|
| *ENSG00000100055* | *CYTH4* | 38,9 | 4,2 | 0,005405555 |
| *ENSG00000080031* | *PTPRH* | 89,6 | 3,7 | 0,005457116 |
| *ENSG00000106144* | *CASP2* | 1800,6 | 1,1 | 0,005511643 |
| *ENSG00000179331* | *RAB39A* | 23,8 | 7,4 | 0,00565188 |
| *ENSG00000189366* | *ALG1L* | 282,4 | 2,7 | 0,005667388 |
| *ENSG00000253293* | *HOXA10* | 655,5 | 1,1 | 0,00573323 |
| *ENSG00000186732* | *MPPED1* | 54,8 | 5,8 | 0,005815059 |
| *ENSG00000130948* | *HSD17B3* | 26,1 | 3,0 | 0,005822314 |
| *ENSG00000184012* | *TMPRSS2* | 128,7 | 5,8 | 0,005981511 |
| *ENSG00000174837* | *ADGRE1* | 93,8 | 7,4 | 0,006043695 |
| *ENSG00000182732* | *RGS6* | 8,5 | 5,9 | 0,006097028 |
| *ENSG00000133321* | *RARRES3* | 848,0 | 3,9 | 0,006102171 |
| *ENSG00000180914* | *OXTR* | 334,4 | 3,8 | 0,006105785 |
| *ENSG00000197587* | *DMBX1* | 15,6 | 6,8 | 0,006111561 |
| *ENSG00000196092* | *PAX5* | 28,5 | 4,7 | 0,006235064 |
| *ENSG00000174827* | *PDZK1* | 258,6 | 4,8 | 0,006710513 |
| *ENSG00000149564* | *ESAM* | 13,5 | 3,5 | 0,007056394 |
| *ENSG00000139515* | *PDX1* | 12,6 | 6,5 | 0,007091247 |
| *ENSG00000239887* | *C1orf226* | 397,8 | 2,5 | 0,007141885 |
| *ENSG00000083454* | *P2RX5* | 61,7 | 3,9 | 0,007325256 |
| *ENSG00000167780* | *SOAT2* | 11,0 | 2,7 | 0,007563295 |
| *ENSG00000012779* | *ALOX5* | 98,7 | 5,5 | 0,007593681 |
| *ENSG00000183729* | *NPBWR1* | 301,4 | 3,8 | 0,007593681 |
| *ENSG00000104368* | *PLAT* | 2311,7 | 3,7 | 0,007617481 |
| *ENSG00000177706* | *FAM20C* | 2532,6 | 2,7 | 0,007809823 |
| *ENSG00000121075* | *TBX4* | 34,4 | 4,5 | 0,007841301 |
| *ENSG00000139192* | *TAPBPL* | 765,3 | 1,8 | 0,007919018 |
| *ENSG00000155849* | *ELMO1* | 74,3 | 5,5 | 0,00820742 |
| *ENSG00000154146* | *NRGN* | 139,7 | 3,3 | 0,008380258 |
| *ENSG00000196787* | *HIST1H2AG* | 160,3 | 2,1 | 0,00845239 |
| *ENSG00000176058* | *TPRN* | 626,5 | 1,1 | 0,008561192 |
| *ENSG00000170525* | *PFKFB3* | 3035,2 | 1,9 | 0,008632834 |
| *ENSG00000123358* | *NR4A1* | 149,1 | 2,1 | 0,008632834 |
| *ENSG00000121380* | *BCL2L14* | 30,5 | 3,8 | 0,008705254 |
| *ENSG00000128833* | *MYO5C* | 467,7 | 3,4 | 0,009198387 |
| *ENSG00000151967* | *SCHIP1* | 81,9 | 1,4 | 0,009212303 |
| *ENSG00000167191* | *GPRC5B* | 1181,2 | 3,6 | 0,009319186 |
| *ENSG00000132434* | *LANCL2* | 751,8 | 1,1 | 0,009722439 |
| *ENSG00000233198* | *RNF224* | 58,2 | 2,7 | 0,009817505 |
| *ENSG00000124256* | *ZBP1* | 71,1 | 6,3 | 0,009819197 |
| *ENSG00000234616* | *JRK* | 2509,0 | 1,9 | 0,010009174 |
| *ENSG00000008300* | *CELSR3* | 537,7 | 2,0 | 0,010016599 |
| *ENSG00000169919* | *GUSB* | 1315,0 | 1,0 | 0,010026494 |
| *ENSG00000110848* | *CD69* | 10,1 | 5,5 | 0,010450912 |
| *ENSG00000120645* | *IQSEC3* | 100,9 | 5,5 | 0,010559047 |

| | | | | |
|---|---|---|---|---|
| *ENSG00000164690* | *SHH* | 8,3 | 5,2 | 0,010848815 |
| *ENSG00000172000* | *ZNF556* | 56,4 | 8,0 | 0,011086479 |
| *ENSG00000100342* | *APOL1* | 2967,2 | 3,2 | 0,011415918 |
| *ENSG00000198944* | *SOWAHA* | 13,8 | 4,9 | 0,011770314 |
| *ENSG00000138193* | *PLCE1* | 299,8 | 2,7 | 0,011867634 |
| *ENSG00000184786* | *TCTE3* | 98,2 | 1,1 | 0,012016631 |
| *ENSG00000197980* | *LEKR1* | 36,3 | 1,4 | 0,012066691 |
| *ENSG00000166669* | *ATF7IP2* | 299,1 | 1,4 | 0,01209456 |
| *ENSG00000095596* | *CYP26A1* | 9,6 | 6,1 | 0,012393593 |
| *ENSG00000185686* | *PRAME* | 51,0 | 6,0 | 0,012445414 |
| *ENSG00000173227* | *SYT12* | 620,4 | 3,0 | 0,012572684 |
| *ENSG00000088448* | *ANKRD10* | 1919,0 | 1,2 | 0,012587542 |
| *ENSG00000165716* | *FAM69B* | 249,9 | 2,8 | 0,012755177 |
| *ENSG00000165238* | *WNK2* | 387,4 | 4,8 | 0,012759946 |
| *ENSG00000107551* | *RASSF4* | 141,0 | 2,4 | 0,012990312 |
| *ENSG00000189108* | *IL1RAPL2* | 16,1 | 4,8 | 0,013071654 |
| *ENSG00000152785* | *BMP3* | 30,5 | 6,5 | 0,013160302 |
| *ENSG00000120068* | *HOXB8* | 20,3 | 4,1 | 0,013365525 |
| *ENSG00000139800* | *ZIC5* | 7,9 | 5,9 | 0,013485635 |
| *ENSG00000124212* | *PTGIS* | 23,2 | 7,4 | 0,013632135 |
| *ENSG00000066382* | *MPPED2* | 82,5 | 7,5 | 0,013661494 |
| *ENSG00000184227* | *ACOT1* | 183,9 | 1,9 | 0,013665513 |
| *ENSG00000179083* | *FAM133A* | 22,9 | 7,4 | 0,014100424 |
| *ENSG00000134508* | *CABLES1* | 318,0 | 2,1 | 0,014100424 |
| *ENSG00000205809* | *KLRC2* | 8,0 | 5,1 | 0,014157914 |
| *ENSG00000100583* | *SAMD15* | 78,7 | 1,9 | 0,014397929 |
| *ENSG00000181143* | *MUC16* | 1374,3 | 5,0 | 0,014423347 |
| *ENSG00000164916* | *FOXK1* | 4151,0 | 1,0 | 0,014440197 |
| *ENSG00000114107* | *CEP70* | 809,8 | 1,3 | 0,014481724 |
| *ENSG00000151491* | *EPS8* | 1829,5 | 2,8 | 0,014518978 |
| *ENSG00000136267* | *DGKB* | 8,6 | 6,0 | 0,014635534 |
| *ENSG00000205359* | *SLCO6A1* | 13,5 | 5,8 | 0,01464648 |
| *ENSG00000183688* | *RFLNB* | 1792,8 | 2,9 | 0,014711815 |
| *ENSG00000077092* | *RARB* | 235,1 | 4,5 | 0,014904445 |
| *ENSG00000271503* | *CCL5* | 281,0 | 4,3 | 0,014964508 |
| *ENSG00000157833* | *GAREM2* | 207,4 | 2,9 | 0,015317383 |
| *ENSG00000198807* | *PAX9* | 236,5 | 2,6 | 0,015647653 |
| *ENSG00000128342* | *LIF* | 2028,0 | 3,2 | 0,015689012 |
| *ENSG00000117318* | *ID3* | 2691,4 | 2,2 | 0,015776524 |
| *ENSG00000106484* | *MEST* | 1568,1 | 2,9 | 0,015800139 |
| *ENSG00000166897* | *ELFN2* | 949,0 | 3,0 | 0,015820988 |
| *ENSG00000102452* | *NALCN* | 78,0 | 3,4 | 0,015915523 |
| *ENSG00000148677* | *ANKRD1* | 71,4 | 3,9 | 0,016066611 |
| *ENSG00000080224* | *EPHA6* | 45,8 | 4,6 | 0,016277722 |
| *ENSG00000053918* | *KCNQ1* | 40,0 | 4,7 | 0,016459791 |

| | | | | |
|---|---|---|---|---|
| *ENSG00000163053* | *SLC16A14* | 139,7 | 2,3 | 0,01650506 |
| *ENSG00000002726* | *AOC1* | 40,9 | 4,3 | 0,016598168 |
| *ENSG00000166448* | *TMEM130* | 29,9 | 4,4 | 0,016598168 |
| *ENSG00000183638* | *RP1L1* | 61,1 | 3,0 | 0,016630957 |
| *ENSG00000134339* | *SAA2* | 300,1 | 4,8 | 0,01674103 |
| *ENSG00000197181* | *PIWIL2* | 54,2 | 3,9 | 0,017119935 |
| *ENSG00000163449* | *TMEM169* | 21,6 | 3,5 | 0,017302395 |
| *ENSG00000170801* | *HTRA3* | 391,8 | 4,4 | 0,017386537 |
| *ENSG00000182168* | *UNC5C* | 17,0 | 2,8 | 0,017491886 |
| *ENSG00000130193* | *THEM6* | 1186,9 | 1,7 | 0,017546363 |
| *ENSG00000188064* | *WNT7B* | 1966,2 | 2,2 | 0,017627622 |
| *ENSG00000108511* | *HOXB6* | 221,0 | 3,0 | 0,017828898 |
| *ENSG00000175564* | *UCP3* | 41,4 | 1,6 | 0,017838645 |
| *ENSG00000205426* | *KRT81* | 194,6 | 3,9 | 0,017991657 |
| *ENSG00000258881* | | 9,5 | 4,8 | 0,017991657 |
| *ENSG00000179023* | *KLHDC7A* | 19,3 | 7,1 | 0,018242846 |
| *ENSG00000148053* | *NTRK2* | 136,9 | 4,1 | 0,01836873 |
| *ENSG00000124839* | *RAB17* | 294,1 | 3,2 | 0,018418646 |
| *ENSG00000107736* | *CDH23* | 48,5 | 2,4 | 0,018776478 |
| *ENSG00000189052* | *CGB5* | 7,0 | 5,7 | 0,018839808 |
| *ENSG00000153993* | *SEMA3D* | 1119,1 | 3,3 | 0,018894833 |
| *ENSG00000167779* | *IGFBP6* | 5079,8 | 2,1 | 0,019132777 |
| *ENSG00000185304* | *RGPD2* | 42,9 | 1,9 | 0,019205691 |
| *ENSG00000197576* | *HOXA4* | 39,9 | 1,5 | 0,019208952 |
| *ENSG00000198771* | *RCSD1* | 95,6 | 3,8 | 0,019258062 |
| *ENSG00000131015* | *ULBP2* | 588,0 | 1,5 | 0,020165862 |
| *ENSG00000072163* | *LIMS2* | 86,4 | 3,1 | 0,020207646 |
| *ENSG00000126561* | *STAT5A* | 371,6 | 2,0 | 0,020207646 |
| *ENSG00000187867* | *PALM3* | 13,6 | 3,6 | 0,020322371 |
| *ENSG00000108813* | *DLX4* | 69,2 | 2,4 | 0,020700222 |
| *ENSG00000173253* | *DMRT2* | 40,5 | 5,4 | 0,020710296 |
| *ENSG00000100628* | *ASB2* | 86,0 | 3,6 | 0,021230933 |
| *ENSG00000274641* | *HIST1H2BO* | 13,3 | 3,0 | 0,021230933 |
| *ENSG00000124507* | *PACSIN1* | 27,5 | 4,3 | 0,021510428 |
| *ENSG00000145451* | *GLRA3* | 14,7 | 5,9 | 0,021607033 |
| *ENSG00000136327* | *NKX2-8* | 70,6 | 3,4 | 0,021680644 |
| *ENSG00000175556* | *LONRF3* | 51,3 | 3,9 | 0,021696882 |
| *ENSG00000165072* | *MAMDC2* | 478,8 | 2,3 | 0,022037495 |
| *ENSG00000136206* | *SPDYE1* | 53,7 | 1,1 | 0,022466095 |
| *ENSG00000259330* | *INAFM2* | 941,6 | 1,2 | 0,022645967 |
| *ENSG00000099282* | *TSPAN15* | 610,1 | 2,6 | 0,022700939 |
| *ENSG00000004139* | *SARM1* | 1070,8 | 1,7 | 0,022748291 |
| *ENSG00000188707* | *ZBED6CL* | 565,9 | 2,2 | 0,022907763 |
| *ENSG00000236320* | *SLFN14* | 2,9 | 4,4 | 0,022990397 |
| *ENSG00000175164* | *ABO* | 123,0 | 4,2 | 0,023027472 |

| | | | | |
|---|---|---|---|---|
| *ENSG00000133488* | *SEC14L4* | 48,2 | 3,5 | 0,023287998 |
| *ENSG00000122367* | *LDB3* | 24,2 | 1,9 | 0,023420371 |
| *ENSG00000123143* | *PKN1* | 2075,0 | 2,8 | 0,023613811 |
| *ENSG00000186998* | *EMID1* | 25,7 | 4,5 | 0,023923289 |
| *ENSG00000119125* | *GDA* | 1469,1 | 3,8 | 0,024138581 |
| *ENSG00000088367* | *EPB41L1* | 5092,6 | 3,0 | 0,024328955 |
| *ENSG00000184371* | *CSF1* | 1318,7 | 2,4 | 0,024455574 |
| *ENSG00000125462* | *C1orf61* | 39,3 | 5,1 | 0,024790046 |
| *ENSG00000120251* | *GRIA2* | 3,1 | 4,5 | 0,024844178 |
| *ENSG00000104081* | *BMF* | 284,2 | 2,3 | 0,02546948 |
| *ENSG00000119673* | *ACOT2* | 557,8 | 1,1 | 0,025521186 |
| *ENSG00000174015* | *SPERT* | 14,5 | 3,2 | 0,025681808 |
| *ENSG00000008277* | *ADAM22* | 81,1 | 2,8 | 0,025724901 |
| *ENSG00000136052* | *SLC41A2* | 269,5 | 1,9 | 0,025724901 |
| *ENSG00000110719* | *TCIRG1* | 2145,8 | 1,3 | 0,026362878 |
| *ENSG00000196159* | *FAT4* | 713,1 | 2,5 | 0,026416833 |
| *ENSG00000152779* | *SLC16A12* | 92,0 | 3,2 | 0,026523291 |
| *ENSG00000167619* | *TMEM145* | 9,0 | 2,7 | 0,026557631 |
| *ENSG00000019505* | *SYT13* | 14,5 | 5,6 | 0,026806251 |
| *ENSG00000082074* | *FYB1* | 1787,5 | 3,2 | 0,02753681 |
| *ENSG00000168280* | *KIF5C* | 24,7 | 2,7 | 0,027542795 |
| *ENSG00000111348* | *ARHGDIB* | 3779,4 | 2,5 | 0,028061375 |
| *ENSG00000198569* | *SLC34A3* | 48,5 | 2,5 | 0,02880328 |
| *ENSG00000116299* | *KIAA1324* | 66,7 | 2,2 | 0,029025336 |
| *ENSG00000205221* | *VIT* | 132,7 | 4,0 | 0,029184801 |
| *ENSG00000013588* | *GPRC5A* | 8289,0 | 2,7 | 0,029247581 |
| *ENSG00000180448* | *ARHGAP45* | 779,8 | 1,6 | 0,029247581 |
| *ENSG00000226690* | | 2,1 | 3,9 | 0,029349841 |
| *ENSG00000158104* | *HPD* | 16,4 | 2,3 | 0,02937684 |
| *ENSG00000019186* | *CYP24A1* | 612,6 | 3,8 | 0,029976436 |
| *ENSG00000111859* | *NEDD9* | 1880,7 | 2,3 | 0,029980956 |
| *ENSG00000165752* | *STK32C* | 383,6 | 1,2 | 0,029989739 |
| *ENSG00000074370* | *ATP2A3* | 140,7 | 3,0 | 0,030141923 |
| *ENSG00000130675* | *MNX1* | 22,7 | 3,6 | 0,030162258 |
| *ENSG00000058866* | *DGKG* | 119,7 | 3,3 | 0,030642628 |
| *ENSG00000215018* | *COL28A1* | 48,5 | 3,6 | 0,030702287 |
| *ENSG00000105929* | *ATP6V0A4* | 16,7 | 2,7 | 0,030747882 |
| *ENSG00000124772* | *CPNE5* | 15,8 | 3,1 | 0,031075523 |
| *ENSG00000180660* | *MAB21L1* | 15,4 | 5,1 | 0,031131113 |
| *ENSG00000162676* | *GFI1* | 22,6 | 2,4 | 0,031426421 |
| *ENSG00000110446* | *SLC15A3* | 844,5 | 2,8 | 0,031455558 |
| *ENSG00000183833* | *MAATS1* | 15,8 | 3,8 | 0,032005099 |
| *ENSG00000196935* | *SRGAP1* | 1429,0 | 1,1 | 0,032175005 |
| *ENSG00000174514* | *MFSD4A* | 101,4 | 2,6 | 0,032214426 |
| *ENSG00000187210* | *GCNT1* | 614,2 | 1,9 | 0,032219488 |

| | | | | |
|---|---|---|---|---|
| *ENSG00000153064* | *BANK1* | 57,3 | 2,4 | 0,032707799 |
| *ENSG00000146938* | *NLGN4X* | 75,4 | 4,3 | 0,032844012 |
| *ENSG00000170962* | *PDGFD* | 253,0 | 4,0 | 0,032874637 |
| *ENSG00000134470* | *IL15RA* | 518,1 | 1,8 | 0,032988605 |
| *ENSG00000044524* | *EPHA3* | 50,1 | 3,7 | 0,033132992 |
| *ENSG00000183873* | *SCN5A* | 25,3 | 2,9 | 0,033153026 |
| *ENSG00000116039* | *ATP6V1B1* | 25,9 | 2,7 | 0,033169749 |
| *ENSG00000182578* | *CSF1R* | 29,8 | 2,7 | 0,033274006 |
| *ENSG00000104783* | *KCNN4* | 1048,4 | 2,5 | 0,033292188 |
| *ENSG00000144230* | *GPR17* | 7,1 | 3,9 | 0,033802021 |
| *ENSG00000182103* | *FAM181B* | 342,6 | 4,2 | 0,033813187 |
| *ENSG00000099953* | *MMP11* | 73,1 | 2,0 | 0,034029042 |
| *ENSG00000198125* | *MB* | 8,4 | 4,8 | 0,034167547 |
| *ENSG00000121716* | *PILRB* | 388,4 | 2,0 | 0,034203538 |
| *ENSG00000179776* | *CDH5* | 23,2 | 5,4 | 0,03451553 |
| *ENSG00000164674* | *SYTL3* | 399,1 | 2,2 | 0,034878145 |
| *ENSG00000156206* | *CFAP161* | 6,4 | 4,7 | 0,035810112 |
| *ENSG00000146757* | *ZNF92* | 317,8 | 1,3 | 0,035810112 |
| *ENSG00000115919* | *KYNU* | 2124,3 | 3,0 | 0,035829762 |
| *ENSG00000147573* | *TRIM55* | 30,1 | 3,9 | 0,036048308 |
| *ENSG00000184845* | *DRD1* | 9,5 | 6,1 | 0,03613826 |
| *ENSG00000213397* | *HAUS7* | 27,7 | 1,4 | 0,036547613 |
| *ENSG00000197977* | *ELOVL2* | 149,6 | 4,1 | 0,036625673 |
| *ENSG00000168778* | *TCTN2* | 628,7 | 1,2 | 0,036691433 |
| *ENSG00000168702* | *LRP1B* | 79,6 | 3,6 | 0,036943016 |
| *ENSG00000158470* | *B4GALT5* | 4442,6 | 1,7 | 0,036968856 |
| *ENSG00000240065* | *PSMB9* | 1374,8 | 2,2 | 0,036985543 |
| *ENSG00000110492* | *MDK* | 3510,7 | 2,3 | 0,037197044 |
| *ENSG00000019582* | *CD74* | 2079,6 | 3,7 | 0,037457049 |
| *ENSG00000164849* | *GPR146* | 57,2 | 2,1 | 0,037953463 |
| *ENSG00000206531* | *CD200R1L* | 8,2 | 5,9 | 0,037985481 |
| *ENSG00000076351* | *SLC46A1* | 1236,5 | 1,8 | 0,038160828 |
| *ENSG00000172508* | *CARNS1* | 24,2 | 1,4 | 0,038187292 |
| *ENSG00000121005* | *CRISPLD1* | 137,9 | 4,3 | 0,038188577 |
| *ENSG00000178965* | *ERICH3* | 5,5 | 4,5 | 0,038209528 |
| *ENSG00000172322* | *CLEC12A* | 15,7 | 6,0 | 0,038220635 |
| *ENSG00000127399* | *LRRC61* | 1036,9 | 1,8 | 0,038372673 |
| *ENSG00000131238* | *PPT1* | 5047,0 | 1,3 | 0,038603703 |
| *ENSG00000162415* | *ZSWIM5* | 33,9 | 2,6 | 0,038654124 |
| *ENSG00000115738* | *ID2* | 704,5 | 2,5 | 0,038690757 |
| *ENSG00000168843* | *FSTL5* | 5,2 | 4,4 | 0,03870815 |
| *ENSG00000148120* | *C9orf3* | 3494,4 | 1,6 | 0,039737626 |
| *ENSG00000204296* | *C6orf10* | 8,2 | 5,2 | 0,03983201 |
| *ENSG00000188404* | *SELL* | 11,2 | 3,3 | 0,040156349 |
| *ENSG00000012171* | *SEMA3B* | 1337,5 | 3,1 | 0,040449719 |

| | | | | |
|---|---|---|---|---|
| *ENSG00000149089* | *APIP* | 954,9 | 1,6 | 0,040472039 |
| *ENSG00000111700* | *SLCO1B3* | 8,7 | 6,0 | 0,040472039 |
| *ENSG00000157064* | *NMNAT2* | 365,7 | 1,6 | 0,040601987 |
| *ENSG00000148200* | *NR6A1* | 391,7 | 1,2 | 0,04069547 |
| *ENSG00000197457* | *STMN3* | 478,0 | 1,9 | 0,040866112 |
| *ENSG00000145945* | *FAM50B* | 427,9 | 1,2 | 0,04114348 |
| *ENSG00000160345* | *C9orf116* | 115,1 | 1,5 | 0,04114348 |
| *ENSG00000198829* | *SUCNR1* | 6,5 | 3,0 | 0,041190265 |
| *ENSG00000205220* | *PSMB10* | 364,8 | 1,7 | 0,041221349 |
| *ENSG00000119888* | *EPCAM* | 1996,3 | 2,3 | 0,041221349 |
| *ENSG00000177519* | *RPRM* | 2,6 | 3,4 | 0,041628765 |
| *ENSG00000139055* | *ERP27* | 154,5 | 4,2 | 0,042141111 |
| *ENSG00000198246* | *SLC29A3* | 631,8 | 3,2 | 0,042323343 |
| *ENSG00000182957* | *SPATA13* | 255,0 | 2,0 | 0,042821827 |
| *ENSG00000175920* | *DOK7* | 190,1 | 2,8 | 0,043108098 |
| *ENSG00000162687* | *KCNT2* | 132,9 | 4,9 | 0,043179377 |
| *ENSG00000138347* | *MYPN* | 99,1 | 3,0 | 0,043533225 |
| *ENSG00000111110* | *PPM1H* | 110,3 | 2,8 | 0,044210903 |
| *ENSG00000136167* | *LCP1* | 1538,2 | 4,1 | 0,044296742 |
| *ENSG00000162391* | *FAM151A* | 186,5 | 1,8 | 0,044296742 |
| *ENSG00000206140* | *TMEM191C* | 18,0 | 1,8 | 0,044354143 |
| *ENSG00000166450* | *PRTG* | 129,8 | 4,5 | 0,044366056 |
| *ENSG00000172461* | *FUT9* | 30,2 | 4,9 | 0,044548287 |
| *ENSG00000163171* | *CDC42EP3* | 4880,7 | 2,2 | 0,044708858 |
| *ENSG00000162390* | *ACOT11* | 541,3 | 1,9 | 0,046182226 |
| *ENSG00000136514* | *RTP4* | 414,0 | 2,7 | 0,046284288 |
| *ENSG00000153208* | *MERTK* | 30,2 | 2,1 | 0,046485764 |
| *ENSG00000250305* | *TRMT9B* | 88,5 | 3,4 | 0,046951421 |
| *ENSG00000060140* | *STYK1* | 194,8 | 1,4 | 0,04725477 |
| *ENSG00000164161* | *HHIP* | 70,9 | 3,7 | 0,047521805 |
| *ENSG00000168993* | *CPLX1* | 34,5 | 2,6 | 0,047653064 |
| *ENSG00000164530* | *PI16* | 5,9 | 4,0 | 0,047717215 |
| *ENSG00000131019* | *ULBP3* | 122,6 | 1,0 | 0,047731978 |
| *ENSG00000185920* | *PTCH1* | 913,2 | 1,6 | 0,04773314 |
| *ENSG00000162946* | *DISC1* | 84,6 | 2,2 | 0,047766939 |
| *ENSG00000214078* | *CPNE1* | 4806,8 | 1,1 | 0,047898686 |
| *ENSG00000105643* | *ARRDC2* | 518,9 | 1,1 | 0,048018792 |
| *ENSG00000163131* | *CTSS* | 752,0 | 2,9 | 0,048030845 |
| *ENSG00000101280* | *ANGPT4* | 19,4 | 2,0 | 0,048030845 |
| *ENSG00000105967* | *TFEC* | 19,2 | 3,9 | 0,048412155 |
| *ENSG00000118004* | *COLEC11* | 14,2 | 4,4 | 0,048594321 |
| *ENSG00000105613* | *MAST1* | 64,9 | 2,3 | 0,048676078 |
| *ENSG00000196776* | *CD47* | 4913,2 | 1,1 | 0,048730581 |
| *ENSG00000162591* | *MEGF6* | 1574,3 | 2,8 | 0,048814386 |
| *ENSG00000138411* | *HECW2* | 239,3 | 3,0 | 0,048984364 |

| | | | | |
|---|---|---|---|---|
| *ENSG00000226479* | *TMEM185B* | 1135,8 | 1,1 | 0,04917418 |
| *ENSG00000149021* | *SCGB1A1* | 10,1 | 4,7 | 0,049218739 |
| *ENSG00000168811* | *IL12A* | 36,8 | 2,6 | 0,049365563 |
| *ENSG00000278828* | *HIST1H3H* | 149,0 | 2,2 | 0,049454437 |
| *ENSG00000186470* | *BTN3A2* | 1217,7 | 1,5 | 0,049517784 |
| *ENSG00000095637* | *SORBS1* | 125,6 | 1,5 | 0,049517784 |
| *ENSG00000065609* | *SNAP91* | 13,5 | 4,9 | 0,049995155 |
| *ENSG00000244300* | *GATA2-AS1* | 357,1 | 8,5 | 2,42271E-10 |
| *ENSG00000232934* | | 86,4 | 6,0 | 1,09373E-05 |
| *ENSG00000244675* | | 178,3 | 6,4 | 1,29013E-05 |
| *ENSG00000223829* | | 56,8 | 7,9 | 1,54737E-05 |
| *ENSG00000229967* | *MAGEA4-AS1* | 139,6 | 10,0 | 3,30675E-05 |
| *ENSG00000280734* | *LINC01232* | 304,3 | 3,3 | 3,34984E-05 |
| *ENSG00000250920* | | 171,9 | 9,6 | 5,33374E-05 |
| *ENSG00000233251* | | 263,3 | 7,0 | 6,36382E-05 |
| *ENSG00000231346* | *LINC01160* | 167,6 | 5,6 | 6,37996E-05 |
| *ENSG00000260604* | | 158,7 | 5,5 | 8,87514E-05 |
| *ENSG00000229214* | *LINC00242* | 72,4 | 3,4 | 0,000206183 |
| *ENSG00000231690* | *LINC00574* | 30,4 | 4,2 | 0,000233362 |
| *ENSG00000230061* | *TRPM2-AS* | 97,4 | 4,8 | 0,000307063 |
| *ENSG00000268531* | | 42,0 | 8,3 | 0,00033776 |
| *ENSG00000224516* | | 344,9 | 8,8 | 0,00034106 |
| *ENSG00000257636* | *G2E3-AS1* | 26,6 | 7,6 | 0,000353979 |
| *ENSG00000275437* | | 60,5 | 1,4 | 0,000370791 |
| *ENSG00000255571* | *MIR9-3HG* | 87,9 | 6,6 | 0,000621335 |
| *ENSG00000250682* | *LINC00491* | 58,6 | 6,7 | 0,000746548 |
| *ENSG00000251169* | *LINC01843* | 20,1 | 6,4 | 0,001029448 |
| *ENSG00000213057* | *C1orf220* | 41,6 | 2,0 | 0,001268659 |
| *ENSG00000228412* | | 35,8 | 5,1 | 0,001376677 |
| *ENSG00000222033* | *LINC01124* | 9,2 | 5,4 | 0,001559979 |
| *ENSG00000266283* | | 58,4 | 3,2 | 0,001712198 |
| *ENSG00000250564* | | 35,5 | 7,3 | 0,001824812 |
| *ENSG00000272463* | | 23,2 | 3,8 | 0,001969142 |
| *ENSG00000271646* | | 84,6 | 1,8 | 0,002013424 |
| *ENSG00000233515* | *LINC01518* | 74,7 | 8,4 | 0,002094643 |
| *ENSG00000243766* | *HOTTIP* | 7,6 | 5,8 | 0,002364294 |
| *ENSG00000206344* | *HCG27* | 52,1 | 1,9 | 0,002441575 |
| *ENSG00000279873* | *LINC01126* | 25,9 | 2,0 | 0,002480544 |
| *ENSG00000258661* | | 30,6 | 5,3 | 0,002592234 |
| *ENSG00000238160* | | 15,5 | 5,7 | 0,003433127 |
| *ENSG00000249001* | | 21,4 | 7,3 | 0,00360993 |
| *ENSG00000257702* | *LBX2-AS1* | 172,4 | 1,7 | 0,003734684 |
| *ENSG00000225361* | *PPP1R26-AS1* | 170,6 | 2,1 | 0,003765042 |
| *ENSG00000250546* | | 25,2 | 7,5 | 0,003965154 |
| *ENSG00000255446* | | 9,6 | 5,4 | 0,004120155 |

| | | | | |
|---|---|---|---|---|
| *ENSG00000231187* | | 57,4 | 2,8 | 0,004245671 |
| *ENSG00000283982* | | 12,9 | 4,6 | 0,004418177 |
| *ENSG00000261654* | | 27,3 | 3,0 | 0,004557692 |
| *ENSG00000273486* | | 50,8 | 1,4 | 0,004755762 |
| *ENSG00000234695* | | 7,9 | 5,1 | 0,004812881 |
| *ENSG00000259672* | | 89,5 | 8,6 | 0,005378909 |
| *ENSG00000225489* | | 29,6 | 2,6 | 0,005860221 |
| *ENSG00000250320* | | 66,8 | 4,4 | 0,005891465 |
| *ENSG00000261379* | | 38,0 | 4,0 | 0,005891465 |
| *ENSG00000232555* | | 16,8 | 3,9 | 0,005946517 |
| *ENSG00000237978* | *KCNMB2-AS1* | 48,5 | 5,0 | 0,006018271 |
| *ENSG00000259953* | | 48,2 | 2,1 | 0,006043695 |
| *ENSG00000268364* | *SMC5-AS1* | 70,4 | 1,8 | 0,006091582 |
| *ENSG00000272622* | | 39,6 | 3,8 | 0,006210431 |
| *ENSG00000272405* | | 76,5 | 3,1 | 0,006251769 |
| *ENSG00000124915* | | 18,9 | 4,8 | 0,006354459 |
| *ENSG00000246526* | *LINC002481* | 11,5 | 5,0 | 0,006361627 |
| *ENSG00000182648* | *LINC01006* | 201,7 | 2,2 | 0,006522036 |
| *ENSG00000223764* | *LINC02593* | 47,1 | 4,3 | 0,00661947 |
| *ENSG00000261211* | | 56,7 | 2,5 | 0,006724738 |
| *ENSG00000270607* | | 29,9 | 5,1 | 0,007036814 |
| *ENSG00000236449* | | 46,7 | 4,6 | 0,007272109 |
| *ENSG00000253931* | | 10,5 | 6,3 | 0,007296581 |
| *ENSG00000228434* | | 39,1 | 1,6 | 0,007396021 |
| *ENSG00000266010* | *GATA6-AS1* | 9,4 | 3,2 | 0,007593681 |
| *ENSG00000228801* | | 147,0 | 1,3 | 0,007722197 |
| *ENSG00000203650* | *LINC01285* | 26,9 | 4,9 | 0,00820742 |
| *ENSG00000260633* | | 15,8 | 2,6 | 0,008333086 |
| *ENSG00000224764* | | 19,5 | 2,7 | 0,008669777 |
| *ENSG00000271314* | | 10,1 | 5,5 | 0,008947344 |
| *ENSG00000260976* | *LINC01633* | 7,3 | 5,7 | 0,00954983 |
| *ENSG00000268996* | *MAN1B1-DT* | 99,0 | 2,0 | 0,010047915 |
| *ENSG00000276707* | | 48,5 | 6,1 | 0,010327664 |
| *ENSG00000248890* | *HHIP-AS1* | 62,8 | 4,4 | 0,010443389 |
| *ENSG00000197536* | *C5orf56* | 652,6 | 2,2 | 0,010742245 |
| *ENSG00000196366* | *C9orf163* | 93,0 | 1,2 | 0,010811453 |
| *ENSG00000227076* | | 25,2 | 4,8 | 0,011141855 |
| *ENSG00000237491* | | 86,4 | 1,5 | 0,011326571 |
| *ENSG00000242474* | | 69,5 | 1,7 | 0,012134402 |
| *ENSG00000229152* | *ANKRD10-IT1* | 209,3 | 1,5 | 0,012138719 |
| *ENSG00000263235* | | 39,5 | 1,2 | 0,012283661 |
| *ENSG00000251577* | | 29,9 | 7,1 | 0,012542655 |
| *ENSG00000268812* | | 161,5 | 3,2 | 0,012597354 |
| *ENSG00000179743* | | 240,9 | 1,4 | 0,012723096 |
| *ENSG00000272694* | | 130,3 | 3,3 | 0,013054112 |

| | | | | |
|---|---|---|---|---|
| *ENSG00000259409* | *BMF-AS1* | 42,3 | 2,7 | 0,013485635 |
| *ENSG00000235277* | | 11,3 | 5,0 | 0,013485635 |
| *ENSG00000229953* | | 30,2 | 3,5 | 0,013671407 |
| *ENSG00000242611* | | 10,8 | 5,5 | 0,01383483 |
| *ENSG00000251350* | *LINC02475* | 26,2 | 7,6 | 0,013918649 |
| *ENSG00000228680* | | 5,6 | 4,6 | 0,013920676 |
| *ENSG00000225411* | | 8,5 | 2,7 | 0,014102704 |
| *ENSG00000264569* | | 3,6 | 4,0 | 0,014144061 |
| *ENSG00000260495* | | 14,0 | 2,1 | 0,014400472 |
| *ENSG00000255045* | | 33,7 | 3,3 | 0,014450177 |
| *ENSG00000259826* | | 26,4 | 1,6 | 0,0145852 |
| *ENSG00000266495* | | 49,8 | 1,8 | 0,014783249 |
| *ENSG00000250220* | | 13,2 | 1,9 | 0,015451967 |
| *ENSG00000254287* | | 18,6 | 6,3 | 0,015877082 |
| *ENSG00000224905* | | 20,7 | 2,6 | 0,015945589 |
| *ENSG00000249867* | | 12,7 | 5,2 | 0,016103362 |
| *ENSG00000264464* | | 16,4 | 6,9 | 0,016194043 |
| *ENSG00000247402* | | 8,8 | 5,2 | 0,016226471 |
| *ENSG00000213468* | *FIRRE* | 26,9 | 5,6 | 0,016506642 |
| *ENSG00000267327* | | 19,1 | 6,0 | 0,016598168 |
| *ENSG00000276075* | | 11,0 | 2,4 | 0,016604674 |
| *ENSG00000250328* | | 27,6 | 5,0 | 0,016812058 |
| *ENSG00000254338* | *MAFA-AS1* | 12,5 | 5,4 | 0,017008587 |
| *ENSG00000188242* | | 949,0 | 1,3 | 0,017440649 |
| *ENSG00000273301* | | 3,2 | 4,5 | 0,017775366 |
| *ENSG00000253552* | *HOXA-AS2* | 511,8 | 2,1 | 0,018201928 |
| *ENSG00000257906* | *LINC02156* | 3,0 | 4,4 | 0,018447949 |
| *ENSG00000268894* | *PLCE1-AS1* | 9,7 | 4,8 | 0,018447949 |
| *ENSG00000258654* | | 18,8 | 7,1 | 0,019119191 |
| *ENSG00000267284* | | 18,8 | 6,0 | 0,019187488 |
| *ENSG00000233101* | *HOXB-AS3* | 207,9 | 2,7 | 0,019748046 |
| *ENSG00000244040* | *IL12A-AS1* | 14,0 | 3,3 | 0,020128883 |
| *ENSG00000250697* | | 60,7 | 2,6 | 0,02029801 |
| *ENSG00000258701* | *LINC00638* | 77,3 | 1,6 | 0,020345445 |
| *ENSG00000239332* | *LINC01119* | 30,9 | 2,5 | 0,020403494 |
| *ENSG00000249082* | *C5orf66-AS1* | 3,7 | 4,0 | 0,020612838 |
| *ENSG00000271930* | | 11,9 | 3,0 | 0,020938483 |
| *ENSG00000259460* | | 40,0 | 1,5 | 0,021261932 |
| *ENSG00000177337* | *DLGAP1-AS1* | 217,9 | 1,6 | 0,021890083 |
| *ENSG00000284428* | | 63,9 | 1,5 | 0,021979424 |
| *ENSG00000272764* | | 19,7 | 1,8 | 0,022046482 |
| *ENSG00000237074* | | 4,9 | 5,2 | 0,022093684 |
| *ENSG00000272620* | *AFAP1-AS1* | 921,1 | 2,7 | 0,022317586 |
| *ENSG00000260742* | | 12,7 | 1,7 | 0,022436591 |
| *ENSG00000260704* | *LINC00543* | 5,5 | 2,4 | 0,022553716 |

| | | | | |
|---|---|---|---|---|
| *ENSG00000251365* | | 4,7 | 4,3 | 0,022822854 |
| *ENSG00000224848* | | 14,8 | 4,5 | 0,023103455 |
| *ENSG00000254363* | | 30,7 | 2,9 | 0,023248889 |
| *ENSG00000243902* | | 16,1 | 3,2 | 0,023714745 |
| *ENSG00000239523* | *MYLK-AS1* | 261,3 | 1,8 | 0,023738538 |
| *ENSG00000249916* | | 11,8 | 5,6 | 0,023905976 |
| *ENSG00000259368* | | 58,8 | 2,4 | 0,023944162 |
| *ENSG00000261798* | | 3,9 | 4,8 | 0,023992763 |
| *ENSG00000273473* | | 5,8 | 3,8 | 0,024778371 |
| *ENSG00000272372* | | 4,0 | 4,8 | 0,024919584 |
| *ENSG00000236242* | *MYO16-AS1* | 28,7 | 2,3 | 0,024952074 |
| *ENSG00000259884* | | 36,6 | 2,1 | 0,025023505 |
| *ENSG00000270823* | | 422,9 | 2,8 | 0,025422713 |
| *ENSG00000271966* | | 17,8 | 1,8 | 0,02546948 |
| *ENSG00000230149* | | 201,0 | 1,2 | 0,025511886 |
| *ENSG00000254812* | | 14,5 | 1,9 | 0,025736178 |
| *ENSG00000259180* | | 5,2 | 5,2 | 0,026081279 |
| *ENSG00000225087* | | 18,8 | 5,1 | 0,026081279 |
| *ENSG00000248079* | *DPH6-DT* | 37,4 | 2,2 | 0,026193491 |
| *ENSG00000277619* | | 8,0 | 2,3 | 0,02630431 |
| *ENSG00000259523* | | 21,0 | 1,4 | 0,026414041 |
| *ENSG00000253210* | | 37,0 | 1,6 | 0,026414041 |
| *ENSG00000258168* | | 45,5 | 2,6 | 0,027397395 |
| *ENSG00000230333* | | 4,1 | 4,1 | 0,027542795 |
| *ENSG00000272769* | | 4,0 | 4,1 | 0,028265735 |
| *ENSG00000276603* | | 66,5 | 1,9 | 0,029976436 |
| *ENSG00000257607* | | 35,8 | 1,8 | 0,030162258 |
| *ENSG00000278071* | | 18,8 | 4,7 | 0,03022934 |
| *ENSG00000247934* | | 177,0 | 1,0 | 0,030494454 |
| *ENSG00000257515* | | 3,8 | 4,1 | 0,031171525 |
| *ENSG00000259341* | | 39,5 | 3,5 | 0,031452357 |
| *ENSG00000247317* | *LY6E-DT* | 112,8 | 2,4 | 0,031462979 |
| *ENSG00000255629* | | 101,5 | 1,1 | 0,031910745 |
| *ENSG00000237862* | | 7,2 | 5,7 | 0,032707799 |
| *ENSG00000272068* | | 8,7 | 3,6 | 0,033055762 |
| *ENSG00000146521* | *LINC01558* | 4,2 | 4,9 | 0,033069753 |
| *ENSG00000250635* | *CXXC5-AS1* | 9,2 | 3,3 | 0,033105844 |
| *ENSG00000255498* | | 44,5 | 1,8 | 0,033226954 |
| *ENSG00000234076* | *TPRG1-AS1* | 15,0 | 2,7 | 0,033292188 |
| *ENSG00000236268* | *LINC01361* | 7,7 | 5,0 | 0,033451197 |
| *ENSG00000205334* | *LINC01460* | 15,7 | 3,4 | 0,034276683 |
| *ENSG00000273100* | | 10,2 | 3,5 | 0,035125572 |
| *ENSG00000253806* | | 176,4 | 1,5 | 0,035271779 |
| *ENSG00000250421* | | 13,7 | 4,6 | 0,03635629 |
| *ENSG00000236423* | *LINC01134* | 38,5 | 1,6 | 0,036383792 |

| | | | | |
|---|---|---|---|---|
| *ENSG00000231764* | *DLX6-AS1* | 67,6 | 2,5 | 0,036761608 |
| *ENSG00000235741* | | 15,6 | 4,0 | 0,036761608 |
| *ENSG00000275322* | | 23,0 | 2,9 | 0,036985543 |
| *ENSG00000258534* | | 26,0 | 1,7 | 0,037228358 |
| *ENSG00000231560* | *CLEC12A-AS1* | 6,1 | 5,5 | 0,037752218 |
| *ENSG00000268686* | | 18,1 | 3,9 | 0,037964732 |
| *ENSG00000271776* | | 5,2 | 4,4 | 0,03839163 |
| *ENSG00000281756* | *C2-AS1* | 4,2 | 3,5 | 0,038552204 |
| *ENSG00000272767* | *JMJD1C-AS1* | 76,3 | 1,5 | 0,038726511 |
| *ENSG00000233117* | *LINC00702* | 70,8 | 2,9 | 0,039658335 |
| *ENSG00000258789* | | 17,9 | 1,4 | 0,03967463 |
| *ENSG00000227479* | | 15,6 | 3,3 | 0,040449719 |
| *ENSG00000264456* | | 61,4 | 1,1 | 0,040742326 |
| *ENSG00000241202* | *ZIC4-AS1* | 8,6 | 5,2 | 0,041026579 |
| *ENSG00000273329* | | 130,6 | 1,3 | 0,041543877 |
| *ENSG00000274922* | | 26,8 | 2,0 | 0,04176694 |
| *ENSG00000225032* | | 240,1 | 2,1 | 0,041951427 |
| *ENSG00000258754* | *LINC01579* | 34,6 | 5,2 | 0,042918871 |
| *ENSG00000204706* | *MAMDC2-AS1* | 60,4 | 1,9 | 0,043209904 |
| *ENSG00000283554* | *LINC02341* | 21,0 | 3,1 | 0,043709054 |
| *ENSG00000259877* | | 103,4 | 1,4 | 0,043709054 |
| *ENSG00000230648* | | 5,5 | 2,9 | 0,044210903 |
| *ENSG00000272328* | | 3,8 | 4,8 | 0,044954274 |
| *ENSG00000272180* | | 23,6 | 4,4 | 0,045746335 |
| *ENSG00000275450* | | 3,4 | 3,8 | 0,046063651 |
| *ENSG00000271122* | | 615,1 | 1,3 | 0,046114492 |
| *ENSG00000226659* | | 20,8 | 2,4 | 0,046119589 |
| *ENSG00000278383* | | 18,6 | 1,6 | 0,047619638 |
| *ENSG00000250994* | | 6,4 | 4,9 | 0,047653251 |
| *ENSG00000254389* | *RHPN1-AS1* | 25,6 | 2,1 | 0,04847901 |
| *ENSG00000206532* | | 20,1 | 4,3 | 0,04847901 |
| *ENSG00000260302* | | 10,3 | 4,2 | 0,049059113 |
| *ENSG00000228919* | | 95,3 | 1,6 | 0,049503279 |
| *ENSG00000227803* | | 28,7 | 2,0 | 0,049621281 |
| *ENSG00000253395* | | 13,8 | 2,1 | 0,049731469 |

**Supplement table S6.** Cluster 1 genes annotated with potential upstream regulator transcription factors (TFs).

| **HGNC.symbol** | **TFs** |
|---|---|
| *C16orf54* | E2F1, FLI1, RUNX1, MYC, SPI1, KLF4, SIN3B, ERG, GFI1B, TRIM28, SOX9, TFCP2L1, MECOM, SIN3A, LMO2, NACC1, NR0B1, CHD2, CTCF, EP300, ETS1, GATA1, HCFC1, MAZ, MXI1, MYOD1, MYOG, NELFE, POLR2A, TAL1, TBP, TCF12, TCF3, ZMIZ1 |
| *MMP13* | ETV4, FOS, JUN, NFKB1, SMAD3, TP53, STAT3, SMAD4, SOX2, SALL4, SOX17, SMAD2, POU3F2, BACH1, CEBPB, E2F4, EZH2, FOSL2, H2AFZ, JUND, MAFF, MAFK, POLR2A, RBBP5, SAP30, SETDB1, TRIM28 |

| | |
|---|---|
| *OLR1* | AR, SMAD4, TP63, NANOG, TCF3, SOX9, SRY, SMAD2, SMAD3, MECOM, STAT4, GATA4, RELA, BACH1, CEBPB, BRCA1, CBX2, CHD2, CTCF, E2F4, EP300, ESR1, EZH2, FOS, FOSL1, FOXA1, GATA1, GATA3, GTF2F1, GTF3C2, H2AFZ, IRF3, JUND, MAFK, MAX, MYB, MYC, NR2F2, POLR2A, PRDM1, RAD21, RCOR1, SAP30, SMC3, SP1, STAT3, TBP, TCF7L2, TFAP2A, TFAP2C, USF1, USF2, ZKSCAN1 |
| *MAGEA4* | ETS1, SMAD4, GATA2, REST, PPARD, BACH1, CTCF, CUX1, EGR1, EZH2, HDAC1, HDAC2, RAD21, SMC3, SPI1 |
| *TRPM2* | E2F1, E2F2, MYC, SPI1, SOX2, EGR1, TP53, HNF4A, SETDB1, SOX9, SOX17, NR3C1, SCLY, TCF7, JARID2, SUZ12, PAX6, BHLHE40, CBX2, CEBPB, CHD1, CHD2, CTCF, CUX1, EBF1, ELF1, EP300, EZH2, FLI1, FOXM1, GATA1, H2AFZ, HDAC1, HNF4G, JUND, KDM5A, KDM5B, MAX, MAZ, MTA3, MXI1, NFIC, PAX5, PHF8, PML, POLR2A, RBBP5, RCOR1, RELA, REST, RUNX3, SAP30, SIN3A, SMC3, SP1, STAT3, TAL1, TBP, TCF12, TCF3, YY1, ZNF143, ZNF263 |
| *CFI* | MYC, PPARG, HNF4A, SIN3B, FOXP1, PPARD, TRIM28, SOX9, SRY, TEAD4, SIN3A, ARID3A, BHLHE40, CEBPB, CHD7, CTCF, CUX1, E2F4, EP300, ETS1, EZH2, FOS, FOXA1, GATA1, H2AFZ, HDAC1, HDAC6, HNF4G, JUND, KDM5A, MAFF, MAFK, MAZ, MXI1, MYBL2, NFIC, POLR2A, RAD21, RBBP5, SP1, STAT3, USF2, YY1, ZEB1, ZNF384 |
| *WDR72* | ZNF217, AR, STAT3, MYC, KLF4, SOX2, TP53, REST, SOX9, SMARCA4, TEAD4, RUNX2, BMI1, ESRRB, CHD1, CHD2, CTCF, E2F6, EP300, EZH2, H2AFZ, HDAC2, KDM4A, MAFF, MAFK, MAX, MAZ, MXI1, NRF1, RBBP5, SAP30, SIN3A, TAF1, ZNF143 |
| *ACSL5* | E2F1, RUNX1, MYC, SPI1, SOX2, PPARG, FOXA2, HNF4A, GATA1, PPARD, ERG, ESR1, SALL4, ZNF281, TEAD4, KLF1, NR3C1, CUX1, CCND1, MYBL2, MECOM, CTNNB1, TBX3, PBX1, IRF1, IRF8, CLOCK, ARID3A, BHLHE40, BRCA1, CEBPB, CEBPD, CHD1, CHD2, CTCF, E2F4, EBF1, ELF1, ELK1, EP300, ETS1, EZH2, FOS, FOXA1, GTF2F1, H2AFZ, HCFC1, HDAC2, HNF4G, IRF3, IRF4, JUND, KAT2A, KDM5B, MAX, MAZ, MBD4, MTA3, MXI1, NELFE, NFIC, NFYA, NFYB, NR2F2, PML, POLR2A, PRDM1, RAD21, RBBP5, RCOR1, RELA, REST, RFX5, RUNX3, RXRA, SIN3A, SMARCC1, SMC3, SP1, SRF, STAT1, TAF1, TBL1XR1, TBP, TCF7L2, TFAP2A, TFAP2C, USF2, YY1, ZEB1, ZKSCAN1, ZMIZ1, ZNF143, ZNF384 |
| *ATOH8* | AR, STAT3, CREM, RUNX1, MYC, POU5F1, SOX2, TCF3, MITF, PPARG, HNF4A, PPARD, TRIM28, SOX9, OLIG2, SMARCA4, WT1, DMRT1, NR3C1, RUNX2, GATA4, TCF7, BMI1, EED, PHC1, EZH2, RNF2, JARID2, MTF2, SUZ12, CTNNB1, SREBF2, NR1I2, NFIB, FOXO1, BACH1, BHLHE40, BRCA1, CHD1, CHD2, CREB1, CTBP2, CTCF, E2F4, E2F6, EBF1, ELF1, EP300, ETS1, FOXA1, FOXA2, FOXP2, GABPA, H2AFZ, HCFC1, HDAC1, HDAC2, HDAC6, HMGN3, JUND, KDM4A, MAFK, MAX, MAZ, MXI1, MYOG, NELFE, PHF8, POLR2A, RAD21, RBBP5, RCOR1, REST, SIN3A, SMC3, SP4, TAF1, TBP, TCF12, TCF7L2, UBTF, USF1, USF2, ZBTB7A, ZC3H11A, ZMIZ1, ZNF143, ZNF263 |
| *GBP4* | STAT3, SMAD4, FLI1, SPI1, NANOG, GATA1, REST, SETDB1, STAT4, CEBPB, IRF8, ATF1, ATF2, BATF, BCL11A, BCLAF1, BHLHE40, CHD1, CTCF, EBF1, EP300, ETS1, EZH2, FOXM1, FOXP2, GATA2, GATA3, H2AFZ, HCFC1, HDAC1, HDAC2, IKZF1, IRF4, JUND, KDM5A, KDM5B, MAFK, MAZ, MEF2A, MTA3, MXI1, MYC, NELFE, NFIC, NR2F2, PAX5, PBX3, PML, POLR2A, POU2F2, RAD21, RBBP5, RCOR1, RELA, RUNX3, SIN3A, SP1, STAT1, STAT5A, TAF1, TAL1, TBL1XR1, TBP, TCF12, TEAD4, WRNIP1, ZC3H11A, ZMIZ1, ZNF143, ZNF384 |
| *NUP210* | SP1, WT1, PAX3, AR, SMAD4, FLI1, RUNX1, MYC, SPI1, TCF3, EGR1, MITF, TP53, HNF4A, GATA1, GATA2, ZFX, RCOR3, SIN3B, REST, SETDB1, TFAP2C, MYCN, TRIM28, EP300, ESR1, TFCP2L1, RUNX2, MECOM, STAT4, FOXP2, PBX1, MYB, RCOR1, TFEB, PDX1, ELF1, NR1H3, STAT6, HIF1A, ARID3A, ATF2, ATF3, BACH1, BCL11A, BCL3, BCLAF1, BHLHE40, BRCA1, CBX3, CCNT2, CEBPB, CEBPD, CHD1, CHD2, CHD4, CHD7, CREB1, CTCF, CTCFL, E2F1, E2F4, E2F6, EBF1, ELK1, ESRRA, ETS1, EZH2, FOXM1, GABPA, GTF2B, GTF2F1, H2AFZ, HCFC1, HDAC1, HDAC2, HDAC6, HMGN3, HNF4G, IKZF1, IRF1, IRF4, JUN, JUND, KAT2A, KAT2B, KDM1A, KDM4A, KDM5A, KDM5B, MAFF, MAFK, MAX, MAZ, MBD4, MTA3, MXI1, MYBL2, MYOD1, MYOG, NANOG, NELFE, NFATC1, NFE2, NFIC, NFYA, NFYB, NR2F2, NR3C1, NRF1, PAX5, PBX3, PHF8, PML, POLR2A, POU2F2, RAD21, RBBP5, RELA, RFX5, RNF2, RUNX3, SAP30, SIN3A, SIRT6, SMARCB1, SMC3, STAT1, STAT5A, SUZ12, TAF1, TAL1, TBL1XR1, TBP, TCF12, TCF7L2, TEAD4, UBTF, USF1, USF2, WHSC1, WRNIP1, YY1, ZBTB7A, ZC3H11A, ZEB1, ZKSCAN1, ZMIZ1, ZNF143, ZNF263, ZNF384 |
| *SPOCK3* | STAT3, SMAD4, MYC, NANOG, FOXA2, HNF4A, REST, EOMES, TFAP2C, PPARD, SOX9, OLIG2, SMAD3, EED, EZH2, RNF2, JARID2, MTF2, SUZ12, JUN, POU3F2, RCOR1, GATA3, BACH1, BATF, BCL11A, BHLHE40, CHD1, CHD2, CHD7, CTBP2, CTCF, E2F4, EP300, H2AFZ, HDAC2, KDM4A, KDM5A, MAX, MAZ, MXI1, PAX5, PHF8, POLR2A, RAD21, RBBP5, RUNX3, SAP30, SIN3A, SMC3, TAF1, TBL1XR1, TBP, TCF12, TEAD4, WRNIP1, ZNF143 |
| *ALPP* | AR, RARA, RARB, RARG, TP53, SMAD3, RUNX2, BHLHE40, CHD2, CTCF, EGR1, EP300, ESR1, EZH2, FOS, HDAC6, NR2F2, NR3C1, POLR2A, REST, RFX5, RXRA |
| *EXOC3L4* | AR, TP63, RUNX1, EGR1, HNF4A, GATA1, SETDB1, PPARD, SOX9, SRY, YAP1, MTF2, SUZ12, ARID3A, BHLHE40, BRCA1, CBX3, CCNT2, CEBPB, CEBPD, CHD1, CHD2, CREB1, CTCF, CUX1, E2F6, EBF1, ELF1, EP300, FOS, FOSL2, FOXA1, FOXA2, GATA2, GTF2B, GTF2F1, H2AFZ, HCFC1, HDAC1, HDAC2, HMGN3, HNF4G, IRF1, IRF4, JUND, KDM5A, KDM5B, MAFK, MAX, MAZ, MBD4, MEF2A, MTA3, MXI1, MYBL2, MYC, MYOG, NCOR1, NFIC, NR2F2, PAX5, PHF8, POLR2A, RAD21, RCOR1, REST, RFX5, RUNX3, RXRA, SAP30, SIN3A, SMC3, SP1, SPI1, STAT3, STAT5A, TAF1, TAL1, TBL1XR1, TBP, TCF12, TCF7L2, TEAD4, TFAP2A, TFAP2C, UBTF, USF1, YY1, ZBTB7A, ZEB1, ZKSCAN1, ZMIZ1, ZNF143, ZNF384 |
| *AMTN* | AR, TP63, NANOG, TP53, TFAP2C, CEBPB, E2F4, EP300, EZH2, FOS, FOSL2, H2AFZ, JUN, JUND, POLR2A, RCOR1, STAT3, TAF1, USF2 |
| *RHOBTB3* | ZNF217, TCF4, SMAD4, CREM, FLI1, RUNX1, SPI1, NANOG, TCF3, MITF, TP53, FOXA2, HNF4A, GATA1, GATA2, ZFX, TFAP2C, SALL4, WT1, ZNF281, KLF1, TFCP2L1, SMAD2, SMAD3, RUNX2, CUX1, HOXB4, MTF2, SUZ12, ESRRB, PBX1, POU3F2, RCOR1, CEBPB, PRDM5, PADI4, STAT1, ESR2, RARG, ARID3A, ATF1, BACH1, BCLAF1, BHLHE40, BRCA1, CBX3, CCNT2, CEBPD, CHD1, CHD2, CHD7, CREB1, CTBP2, CTCF, CTCFL, E2F4, E2F6, EBF1, EGR1, ELF1, ELK1, EP300, ESR1, ETS1, EZH2, FOS, FOXA1, FOXM1, FOXP2, GABPA, GATA3, GTF2B, GTF2F1, H2AFZ, HCFC1, HDAC1, HDAC2, HDAC6, |

| | |
|---|---|
| | HMGN3, IRF1, JUN, JUND, KDM4A, KDM5B, MAFF, MAFK, MAX, MAZ, MEF2A, MTA3, MXI1, MYBL2, MYC, MYOG, NCOR1, NFIC, NR2F2, NRF1, PAX5, PBX3, PHF8, PML, POLR2A, POU2F2, RAD21, RBBP5, REST, RFX5, RUNX3, RXRA, SAP30, SIN3A, SMARCB1, SMARCC1, SMC3, SP1, STAT3, STAT5A, TAF1, TAF7, TAL1, TBL1XR1, TBP, TCF12, TCF7L2, TEAD4, THAP1, TRIM28, UBTF, USF1, USF2, WRNIP1, YY1, ZBTB33, ZBTB7A, ZC3H11A, ZEB1, ZKSCAN1, ZMIZ1, ZNF143, ZNF263, ZNF274, ZNF384 |
| *KRTAP4-1* | EZH2, H2AFZ, MYC, MYOD1, MYOG, POLR2A, TCF3, USF1 |
| *NBEA* | AHR, ARNT, ZNF217, CEBPD, AR, STAT3, TCF4, SMAD4, TP63, CREM, RUNX1, MYC, SOX2, MITF, TP53, FOXA2, HNF4A, EOMES, EP300, ESR1, SALL4, DMRT1, RUNX2, GATA4, JARID2, MTF2, SUZ12, POU3F2, ESR2, BACH1, BRCA1, CEBPB, CHD1, CHD2, CHD7, CREB1, CTCF, E2F4, E2F6, EBF1, EGR1, EZH2, FOXA1, FOXP2, GTF2F1, H2AFZ, HCFC1, HDAC1, HDAC2, HDAC6, HMGN3, JUND, KDM4A, KDM5A, KDM5B, MAFK, MAX, MAZ, MXI1, MYOD1, MYOG, NANOG, NR2F2, NR3C1, NRF1, PAX5, PHF8, POLR2A, RAD21, RBBP5, RCOR1, REST, RFX5, RUNX3, SAP30, SIN3A, SMC3, SP1, STAT1, TAF1, TAF7, TBP, TCF12, TCF3, TCF7L2, TEAD4, TRIM28, USF1, USF2, WRNIP1, YY1, ZEB1, ZKSCAN1, ZNF143, ZNF263 |
| *PCDHGB1* | TP63, MYC, SPI1, POU5F1, SOX2, TP53, KDM5B, TET1, TRIM28, ATF3, SMAD2, SMAD3, MYBL2, ZFP42, CTCF, SUZ12, RCOR1, ETS2, HOXD13, ARID3A, BHLHE40, CEBPB, E2F6, ELF1, EZH2, FOXA1, GABPA, GATA2, GATA3, GTF2F1, H2AFZ, HDAC2, HDAC6, JUND, KDM4A, KDM5A, MAFF, MAFK, MAX, MAZ, MXI1, PHF8, POLR2A, RAD21, RBBP5, SETDB1, SIN3A, SMC3, SP1, SRF, TAF1, TAL1, TBP, TCF12, UBTF, ZNF143 |
| *SYT15* | TP63, FLI1, MYC, EGR1, FOXA2, E2F4, RCOR3, SETDB1, MYCN, TBX5, SUZ12, ELF5, ATF2, BACH1, BCL3, BRCA1, CEBPB, CHD1, CHD2, CREB1, CREBBP, CTBP2, CTCF, E2F6, ELK1, EP300, EZH2, FOSL2, GABPA, GATA2, GATA3, GTF2F1, H2AFZ, HDAC2, HDAC6, HMGN3, IRF1, JUN, JUND, KDM4A, KDM5B, MAX, MAZ, MXI1, NANOG, NRF1, PHF8, POLR2A, RAD21, RCOR1, RFX5, SAP30, SIN3A, SIRT6, SP4, TBP, TCF12, TCF7L2, TEAD4, UBTF, USF2, YY1, ZBTB7A, ZKSCAN1, ZNF143, ZNF274 |
| *INHBB* | AHR, ARNT, NFE2L2, AR, E2F1, FLI1, KLF4, POU5F1, SOX2, TCF3, EGR1, PPARG, KDM5B, ZFX, RCOR3, SIN3B, REST, ASH2L, TFAP2C, SOX9, ESR1, WT1, ZNF281, DMRT1, PRDM14, RUNX2, MEF2A, FOXP2, EZH2, RNF2, JARID2, MTF2, SUZ12, BACH1, CBX2, CEBPB, CHD1, CHD7, CREB1, CTBP2, CTCF, E2F6, EP300, FOXA1, FOXA2, GABPA, GATA3, H2AFZ, HCFC1, HDAC2, HMGN3, JUND, KDM4A, MAX, MAZ, MXI1, MYC, MYOG, NR3C1, PHF8, POLR2A, RAD21, RCOR1, SIN3A, SIRT6, TAF1, TCF12, TCF7L2, USF1, YY1, ZNF143, ZNF263 |
| *KCNH4* | AR, BRCA1, CEBPA, EGR1, MYB, MYC, NFKB1, SP1, TP53, GATA2, TRIM28, SUZ12, MEIS1, BCL3, RBPJ, ATF2, ATF3, BACH1, BCLAF1, BHLHE40, CBX3, CCNT2, CEBPB, CHD1, CHD2, CHD7, CREB1, CTCF, CTCFL, CUX1, E2F4, E2F6, EBF1, ELF1, EP300, ESRRA, ETS1, EZH2, FOSL2, FOXA1, FOXA2, FOXM1, GABPA, GTF2F1, H2AFZ, HCFC1, HDAC1, HDAC2, HDAC6, HMGN3, IRF1, JUN, JUND, KAT2A, KAT2B, KDM1A, KDM4A, KDM5A, KDM5B, MAFF, MAFK, MAX, MAZ, MTA3, MXI1, NELFE, NR2F2, PAX5, PBX3, PHF8, PML, POLR2A, POU2F2, RAD21, RBBP5, RCOR1, RELA, REST, RFX5, RUNX3, SAP30, SETDB1, SIN3A, SMC3, SP4, STAT1, STAT3, STAT5A, TBL1XR1, TBP, TCF12, TEAD4, UBTF, USF1, USF2, WHSC1, WRNIP1, YY1, ZBTB7A, ZC3H11A, ZMIZ1, ZNF143, ZNF263, ZNF384 |
| *MUC4* | AR, CREB1, ETV4, JUN, RARA, SP1, STAT3, FLI1, RUNX1, NANOG, POU5F1, EGR1, TET1, SMARCA4, RUNX2, SCLY, CTNNB1, DROSHA, IKZF1, BHLHE40, CHD1, CTCF, EP300, EZH2, H2AFZ, HDAC1, HDAC2, HDAC6, HMGN3, JUND, KDM5B, MAX, MAZ, PHF8, POLR2A, RBBP5, RCOR1, SAP30, UBTF, WHSC1 |
| *SECTM1* | TTF2, CREB1, FLI1, MYC, SOX2, EGR1, HNF4A, GATA2, SRY, SALL4, SOX17, CTNNB1, STAT5A, SOX11, IRF8, ATF1, ATF3, BACH1, BCL3, BHLHE40, BRCA1, CBX3, CCNT2, CEBPB, CHD1, CHD2, CHD7, CTCF, E2F4, E2F6, EBF1, ELF1, ELK1, EP300, ETS1, EZH2, FOS, FOSL2, FOXA1, GABPA, GATA3, GTF2F1, H2AFZ, HCFC1, HDAC1, HDAC2, HDAC6, HMGN3, HNF4G, IRF1, JUND, KDM1A, KDM4A, KDM5A, MAFF, MAFK, MAX, MAZ, MXI1, NFIC, NR2F2, NRF1, PAX5, PHF8, PML, POLR2A, PRDM1, RAD21, RBBP5, RCOR1, REST, RFX5, SAP30, SETDB1, SIN3A, SMARCB1, SMC3, SPI1, STAT1, STAT3, SUZ12, TAF1, TAL1, TBL1XR1, TBP, TCF12, TCF7L2, TEAD4, THAP1, TRIM28, UBTF, YY1, ZBTB7A, ZC3H11A, ZKSCAN1, ZMIZ1, ZNF143, ZNF263 |
| *DISP2* | TP63, KLF4, EGR1, HNF4A, GATA1, RCOR3, SIN3B, REST, SETDB1, FOXP1, TET1, OLIG2, NR3C1, TBX5, EZH2, SUZ12, SIN3A, ATF3, BHLHE40, CCNT2, CEBPB, CHD1, CHD2, CREB1, CTCF, E2F4, E2F6, ELF1, EP300, ETS1, FOS, GABPA, H2AFZ, HCFC1, HDAC1, HDAC2, HMGN3, JUND, KDM4A, KDM5A, KDM5B, MAX, MAZ, MXI1, MYC, MYOD1, MYOG, NELFE, NFYA, NFYB, NRF1, PAX5, PHF8, POLR2A, RAD21, RBBP5, RCOR1, RFX5, RUNX3, SAP30, SMARCB1, SMC3, SP1, SP4, STAT1, STAT3, STAT5A, TAF1, TAL1, TBP, TCF12, TCF3, TEAD4, THAP1, UBTF, USF2, YY1, ZBTB7A, ZC3H11A, ZEB1, ZKSCAN1, ZNF143, ZNF263 |
| *BIRC3* | NFKB1, RELA, TP53, FOXP3, TP63, E2F1, RUNX1, MYC, SPI1, GATA1, ZFX, E2F4, RCOR3, SETDB1, PPARD, TAL1, SOX9, SALL4, ZNF281, SMAD2, SMAD3, CUX1, MECOM, STAT4, CTCF, ESRRB, POU3F2, RCOR1, STAT5A, DNAJC2, NOTCH1, ARID3A, ATF1, ATF2, ATF3, BACH1, BATF, BCL11A, BCL3, BCLAF1, BHLHE40, BRCA1, CBX3, CCNT2, CEBPB, CHD1, CHD2, CHD7, CREB1, CTCFL, E2F6, EBF1, ELF1, ELK1, ELK4, EP300, ETS1, EZH2, FOS, FOSL2, FOXM1, FOXP2, GATA2, GATA3, GTF2F1, H2AFZ, HCFC1, HDAC1, HDAC2, HDAC6, HMGN3, IKZF1, IRF1, IRF4, JUN, JUND, KAT2A, KDM1A, KDM4A, KDM5A, KDM5B, MAFF, MAFK, MAX, MAZ, MEF2A, MEF2C, MTA3, MXI1, MYBL2, NELFE, NFATC1, NFIC, NR3C1, NRF1, PAX5, PBX3, PHF8, PML, POLR2A, POU2F2, RAD21, RBBP5, REST, RFX5, RNF2, RUNX3, RXRA, SAP30, SIN3A, SMC3, SP1, SRF, STAT1, STAT3, TAF1, TBL1XR1, TBP, TCF12, TCF3, TCF7L2, TEAD4, TFAP2A, TFAP2C, TRIM28, UBTF, USF1, USF2, WRNIP1, YY1, ZC3H11A, ZEB1, ZKSCAN1, ZMIZ1, ZNF143, ZNF263, ZNF384 |

| | |
|---|---|
| *IQCE* | HOXC9, FOXP3, AR, CREM, MYC, SPI1, KLF4, SOX2, EGR1, MITF, TP53, GATA2, ZFX, SETDB1, TRIM28, SOX9, SCLY, MYBL2, ZFP42, GATA4, TCF7, SUZ12, JUN, HSF1, TBP, ARID3A, BCLAF1, BHLHE40, BRCA1, CBX3, CCNT2, CEBPB, CHD1, CHD2, CREB1, CTCF, E2F4, E2F6, EBF1, ELF1, ELK1, EP300, ESR1, ESRRA, ETS1, EZH2, FOSL2, FOXA1, FOXA2, FOXM1, FOXP2, GABPA, GATA1, GATA3, GTF2F1, GTF3C2, H2AFZ, HCFC1, HDAC1, HDAC2, HMGN3, IRF1, JUND, KDM1A, KDM4A, KDM5A, KDM5B, MAFK, MAX, MAZ, MTA3, MXI1, MYOG, NANOG, NELFE, NFATC1, NFIC, NR2F2, NR3C1, NRF1, PAX5, PHF8, POLR2A, RAD21, RBBP5, RCOR1, REST, RFX5, RUNX3, RXRA, SAP30, SIN3A, SIX5, SMARCB1, SMARCC1, SMC3, SP1, SP4, SREBF1, SRF, STAT1, STAT3, TAF1, TAL1, TBL1XR1, TCF12, TCF3, TCF7L2, TEAD4, TFAP2A, TFAP2C, UBTF, USF2, YY1, ZBTB7A, ZC3H11A, ZKSCAN1, ZMIZ1, ZNF143, ZNF263, ZNF384 |
| *ABCD1* | TP63, SPI1, KLF4, EGR1, MITF, PPARG, ASH2L, SOX9, CTCF, JUN, HSF1, PDX1, IRF8, NR1H3, ARID3A, BACH1, BHLHE40, BRCA1, CBX3, CCNT2, CEBPB, CEBPD, CHD1, CHD2, CREB1, CTCFL, E2F4, E2F6, EBF1, ELF1, ELK1, ELK4, EP300, ESR1, ETS1, EZH2, FLI1, FOS, FOSL2, FOXA1, FOXA2, FOXP2, GABPA, GTF2B, GTF2F1, H2AFZ, HCFC1, HDAC1, HDAC2, HMGN3, HNF4A, IRF1, JUND, KAT2A, KAT2B, KDM4A, KDM5A, MAFF, MAFK, MAX, MAZ, MXI1, MYB, MYBL2, MYC, NELFE, NR2F2, NR3C1, NRF1, PAX5, PBX3, PHF8, PML, POLR2A, RAD21, RBBP5, RCOR1, REST, RFX5, RNF2, RUNX3, RXRA, SAP30, SIN3A, SMARCB1, SMC3, SP1, STAT1, STAT5A, TAF1, TAL1, TBL1XR1, TBP, TCF12, TCF3, TCF7L2, TEAD4, TFAP2A, TFAP2C, TRIM28, UBTF, USF1, USF2, WRNIP1, YY1, ZBTB33, ZBTB7A, ZC3H11A, ZEB1, ZKSCAN1, ZMIZ1, ZNF143, ZNF263, ZNF384 |
| *HOXB7* | SP1, SP3, USF1, KLF4, NANOG, POU5F1, TCF3, MITF, KDM5B, SETDB1, TET1, BMI1, EED, PHC1, EZH2, RNF2, JARID2, MTF2, SUZ12, BACH1, HSF1, SOX11, CDX2, TFEB, ATF3, BCL3, BHLHE40, CCNT2, CHD1, CHD2, CHD7, CREB1, CTBP2, CTCF, CUX1, E2F1, E2F6, EBF1, EGR1, ELF1, ELK1, EP300, ESR1, ETS1, FOS, FOSL2, FOXA1, FOXA2, FOXM1, GABPA, GATA3, GTF2F1, H2AFZ, HCFC1, HDAC6, HMGN3, JUN, JUND, KDM4A, KDM5A, MAFF, MAFK, MAX, MAZ, MTA3, MXI1, MYC, NFE2, NFIC, NFYA, NFYB, NRF1, PAX5, PHF8, PML, POLR2A, RAD21, RBBP5, RCOR1, REST, RFX5, RUNX3, SAP30, SIN3A, SIRT6, SIX5, SMARCB1, SMARCC2, STAT1, STAT5A, TAF1, TBL1XR1, TBP, TCF12, TCF7L2, TEAD4, THAP1, TRIM28, UBTF, USF2, YY1, ZBTB33, ZBTB7A, ZEB1, ZKSCAN1, ZNF143, ZNF263 |
| *GATA6* | AR, STAT3, SMAD4, TP63, MYC, KLF4, NANOG, POU5F1, SOX2, TCF3, EGR1, MITF, TP53, PPARG, FOXA2, HNF4A, GATA1, GATA2, KDM5B, SIN3B, SETDB1, TFAP2C, TET1, TRIM28, SOX9, FOXO3, ESR1, OLIG2, SALL4, SOX17, DMRT1, SMAD3, RUNX2, CUX1, MYBL2, CTCF, BMI1, EED, PHC1, EZH2, RNF2, JARID2, MTF2, SUZ12, TFAP2A, JUN, NR0B1, CDX2, SREBF2, GATA3, FOXO1, HTT, ARID3A, BACH1, BHLHE40, BRCA1, CBX3, CBX8, CCNT2, CEBPB, CEBPD, CHD1, CHD2, CHD7, CREB1, CTBP2, E2F1, E2F4, E2F6, ELF1, ELK1, ELK4, EP300, FOS, FOXA1, FOXP2, GTF2F1, H2AFZ, HCFC1, HDAC1, HDAC2, HDAC6, HMGN3, JUND, KDM1A, KDM4A, KDM5A, MAFK, MAX, MAZ, MXI1, NFYB, NR2F2, PHF8, POLR2A, PRDM1, RAD21, RBBP5, RCOR1, RFX5, SAP30, SIN3A, SIRT6, SMARCB1, SMC3, SP1, TAF1, TAL1, TBL1XR1, TBP, TCF12, TCF7L2, TEAD4, UBTF, WHSC1, YY1, ZBTB7A, ZKSCAN1, ZMIZ1, ZNF143, ZNF263, ZNF384 |
| *NR5A2* | ZNF217, PAX3, FOXP3, AR, STAT3, TP63, MYC, NANOG, POU5F1, SOX2, TCF3, EGR1, TP53, HNF4A, KDM5B, RCOR3, REST, SETDB1, ASH2L, PPARD, TET1, SMARCA4, YAP1, SALL4, SOX17, DMRT1, TEAD4, PRDM14, NR3C1, TBX3, SMAD1, GBX2, HOXD13, ARID3A, ATF3, BCL3, BHLHE40, CEBPB, CHD1, CHD2, CHD7, CREB1, CTBP2, CTCF, E2F4, ELF1, ELK1, EP300, EZH2, FOS, FOSL2, FOXA1, FOXA2, GABPA, GATA1, GATA3, GTF2F1, H2AFZ, HDAC2, KDM4A, KDM5A, MAX, MAZ, MXI1, NRF1, PHF8, POLR2A, RBBP5, RCOR1, RFX5, SAP30, SIN3A, SMC3, SP1, SUZ12, TAF1, TBP, TCF12, TRIM28, USF1, USF2, YY1, ZKSCAN1, ZNF143, ZNF263, ZNF384 |
| *VTCN1* | ZNF217, PAX3, AR, TCF4, SMAD4, CREB1, NANOG, EGR1, MITF, RCOR3, SETDB1, SOX9, SRY, ESR1, RUNX2, RCOR1, ELF5, NFIB, ESR2, CEBPB, CREBBP, CTCF, E2F4, ELF1, EP300, EZH2, FOS, FOXA1, GATA2, GATA3, H2AFZ, HDAC2, HDAC6, KDM5A, KDM5B, MAX, MYC, POLR2A, RBBP5, REST, SAP30, SIN3A, SMC3, STAT3, TBP |
| *ADAM28* | AR, PPARG, PAX3, TCF4, SMAD4, SPI1, NANOG, SOX2, EGR1, TET1, SMAD3, FOXP2, BACH1, CEBPA, CEBPB, ARID3A, ATF2, BATF, BCL11A, BHLHE40, CEBPD, CHD1, CHD2, CHD7, CTCF, E2F4, EBF1, ELF1, ELK1, EP300, FOS, FOXA1, FOXA2, GABPA, GTF2F1, H2AFZ, HDAC2, IRF4, MAX, MAZ, MBD4, MEF2A, MTA3, MXI1, MYC, NRF1, PAX5, POLR2A, RBBP5, RCOR1, REST, RUNX3, SIN3A, SMC3, STAT3, TAF1, TBL1XR1, TCF12, WRNIP1, YY1, ZNF143 |
| *THSD7B* | ZNF217, STAT3, TCF4, SMAD4, TP63, CREB1, TP53, HNF4A, GATA1, PPARD, SOX9, OLIG2, SMARCA4, YAP1, SMAD3, TCF7, EED, JARID2, MTF2, SUZ12, TBX3, CEBPB, CTCF, EP300, ETS1, EZH2, POLR2A, SAP30, SMC3 |
| *TMEM71* | VDR, FOXP3, AR, STAT3, FLI1, RUNX1, MITF, PPARD, SALL4, TEAD4, ATF3, MECOM, STAT4, EWSR1, STAT6, BHLHE40, CBX8, CEBPB, CTCF, E2F4, EP300, ETS1, EZH2, FOS, FOXM1, FOXP2, GATA1, H2AFZ, HDAC1, HDAC6, JUN, JUND, MAX, MXI1, MYC, POLR2A, RBBP5, SETDB1, SIN3A, SPI1, TBP, TRIM28, ZNF143 |
| *TBC1D10C* | FLI1, SPI1, KLF4, EGR1, CHD1, CEBPB, SREBF2, RBPJ, SREBF1, ATF2, BACH1, BCL11A, BCLAF1, BHLHE40, CHD2, CHD4, CTCF, EBF1, ELF1, ELK1, EP300, ETS1, EZH2, FOXM1, FOXP2, GATA1, GTF2B, GTF2F1, H2AFZ, HCFC1, HDAC1, HDAC2, HDAC6, HMGN3, IKZF1, IRF1, IRF4, JUN, JUND, KAT2A, KDM5B, MAFF, MAX, MAZ, MTA3, MXI1, MYBL2, MYC, MYOG, NELFE, NFIC, NR2C2, NR2F2, NRF1, PAX5, PHF8, PML, POLR2A, POU2F2, RAD21, RBBP5, RCOR1, REST, RFX5, RUNX3, RXRA, SAP30, SIN3A, SMC3, SP1, SP4, STAT1, STAT3, STAT5A, TAF1, TBL1XR1, TBP, TCF12, TCF3, TCF7L2, TEAD4, UBTF, USF1, USF2, WRNIP1, YY1, ZBTB7A, ZC3H11A, ZEB1, ZMIZ1, ZNF143, ZNF384 |

| | |
|---|---|
| *IGF2BP1* | TTF2, STAT3, SMAD4, E2F1, MYC, SPI1, KLF4, NANOG, POU5F1, SOX2, TCF3, EGR1, FOXA2, HNF4A, KDM5B, ZFX, RCOR3, REST, ASH2L, TFAP2C, PPARD, ERG, SRY, EP300, WT1, ZNF281, DMRT1, TEAD4, PRDM14, RUNX2, CCND1, MYBL2, BMI1, EZH2, CTNNB1, RELA, SOX11, ZIC3, ARID3A, ATF2, ATF3, BACH1, BCL3, BCLAF1, BHLHE40, BRCA1, CBX3, CCNT2, CEBPB, CEBPD, CHD1, CHD2, CHD4, CHD7, CREB1, CTBP2, CTCF, CUX1, E2F4, E2F6, ELF1, ELK1, ELK4, ETS1, FOSL2, FOXA1, FOXP2, GABPA, GATA3, GTF2B, GTF2F1, H2AFZ, HCFC1, HDAC1, HDAC2, HDAC6, HMGN3, HNF4G, IRF1, JUN, JUND, KAT2A, KAT2B, KDM1A, KDM4A, MAFK, MAX, MAZ, MBD4, MXI1, NCOR1, NFIC, NR2F2, NR3C1, PBX3, PHF8, PML, POLR2A, RAD21, RBBP5, RCOR1, RFX5, RNF2, RUNX3, SAP30, SETDB1, SIN3A, SIRT6, SIX5, SMARCB1, SMC3, SP1, SP4, STAT5A, SUZ12, TAF1, TAF7, TAL1, TBL1XR1, TBP, TCF12, TCF7L2, THAP1, TRIM28, UBTF, USF1, USF2, YY1, ZBTB33, ZBTB7A, ZC3H11A, ZKSCAN1, ZMIZ1, ZNF143, ZNF263, ZNF384 |
| *PLA2G16* | TCF4, FLI1, MYC, NANOG, SOX2, PPARG, HNF4A, GATA1, KLF1, ATF3, TBX5, SUZ12, BACH1, TAF7L, TFEB, ESR2, ZNF263, ATF1, BHLHE40, BRCA1, CCNT2, CEBPB, CHD1, CHD2, CHD7, CREB1, CTBP2, CTCF, E2F1, E2F6, EBF1, EGR1, ELF1, ELK1, EP300, ETS1, EZH2, FOS, FOSL2, FOXA1, GABPA, GATA2, GATA3, GTF2F1, H2AFZ, HCFC1, HDAC1, HDAC2, HDAC6, HMGN3, JUN, JUND, KDM1A, KDM4A, KDM5A, KDM5B, MAFK, MAX, MAZ, MXI1, MYOD1, MYOG, NELFE, NFYA, NFYB, NR2F2, PHF8, PML, POLR2A, RAD21, RBBP5, RCOR1, REST, RFX5, RNF2, RUNX3, RXRA, SAP30, SIN3A, SMARCB1, SMARCC1, SMC3, SP1, STAT1, STAT3, STAT5A, TAF1, TAL1, TBL1XR1, TBP, TCF12, TCF3, TCF7L2, TEAD4, TRIM28, UBTF, USF1, USF2, WRNIP1, YY1, ZBTB7A, ZKSCAN1, ZMIZ1, ZNF143 |
| *CATSPER1* | NFE2L2, TP63, FLI1, RUNX1, MYC, EGR1, ERG, FOXP2, RBPJ, BHLHE40, CTCF, EZH2, GATA2, H2AFZ, KDM5A, KDM5B, POLR2A, RBBP5, RCOR1, REST, SAP30, STAT3 |
| *HIST1H2BH* | VDR, TP63, MYC, NANOG, POU5F1, SOX2, EGR1, PPARG, GATA2, ZFX, RCOR3, SIN3B, TET1, ERG, CNOT3, MYBL2, ZFP42, TCF7, RNF2, EWSR1, TBP, CDX2, CEBPA, ARID3A, ATF1, ATF2, BACH1, BATF, BCL3, BCLAF1, BHLHE40, BRCA1, CBX3, CCNT2, CEBPB, CEBPD, CHD1, CHD2, CHD7, CREB1, CTCF, CUX1, E2F4, E2F6, EBF1, ELF1, ELK1, ELK4, EP300, ETS1, EZH2, FLI1, FOS, FOXA1, FOXA2, FOXM1, GABPA, GATA1, GATA3, GTF2B, GTF2F1, H2AFZ, HCFC1, HDAC1, HDAC2, HDAC6, HMGN3, IKZF1, IRF1, IRF3, IRF4, JUN, JUND, KAT2A, KDM1A, KDM4A, KDM5A, KDM5B, MAFF, MAFK, MAX, MAZ, MBD4, MTA3, MXI1, NELFE, NFATC1, NFE2, NFIC, NFYA, NFYB, NR2F2, NRF1, PAX5, PBX3, PHF8, PML, POLR2A, POU2F2, RAD21, RBBP5, RCOR1, RELA, REST, RFX5, RUNX3, RXRA, SAP30, SIN3A, SIRT6, SMC3, SP1, SP4, STAT1, STAT3, STAT5A, SUZ12, TAF1, TAF7, TAL1, TBL1XR1, TCF12, TCF3, TCF7L2, TEAD4, TRIM28, UBTF, USF2, WRNIP1, YY1, ZBTB33, ZEB1, ZKSCAN1, ZMIZ1, ZNF143, ZNF263, ZNF384 |
| *SLC5A12* | AR, SMAD4, EOMES, FOXP2, SUZ12, STAT5A, RBPJ, IRF8, CTCF, EP300, EZH2, FOS, H2AFZ, MAFF, MAFK, MYC, POLR2A, STAT3 |
| *CCDC85A* | STAT3, E2F1, NANOG, SOX2, TCF3, TP53, SIN3B, ASH2L, EOMES, PPARD, OLIG2, SMARCA4, NR3C1, CTCF, MTF2, SUZ12, ESRRB, TBX3, NACC1, NR0B1, GBX2, ATF3, BACH1, BCL11A, BHLHE40, BRCA1, CBX3, CCNT2, CHD1, CHD2, CHD7, CREB1, CTBP2, CUX1, E2F4, E2F6, EBF1, EGR1, ELF1, ELK1, EP300, EZH2, FOXA1, FOXM1, GABPA, GATA3, GTF2F1, H2AFZ, HCFC1, HDAC2, HDAC6, HMGN3, JUND, KDM4A, KDM5B, MAFF, MAFK, MAX, MAZ, MXI1, MYC, NFIC, PBX3, PHF8, POLR2A, POU2F2, POU5F1, RAD21, RBBP5, RCOR1, REST, RFX5, RUNX3, SAP30, SIN3A, SIRT6, SMC3, SP1, STAT1, TAF1, TBL1XR1, TBP, TCF12, TEAD4, UBTF, WRNIP1, YY1, ZBTB7A, ZC3H11A, ZKSCAN1, ZMIZ1, ZNF143, ZNF263, ZNF384 |
| *B3GNT7* | STAT3, TCF4, E2F1, FLI1, RUNX1, MYC, SPI1, KLF4, NANOG, POU5F1, SOX2, TCF3, MITF, TP53, GATA2, SIN3B, SETDB1, EOMES, TFAP2C, PPARD, TET1, MYCN, TAL1, SRY, EP300, SMARCA4, YAP1, SALL4, PRDM14, KLF1, RUNX2, MECOM, TCF7, TBX3, SIN3A, MYB, LMO2, HSF1, NACC1, NR0B1, SMAD1, RBPJ, ATF2, BACH1, BATF, BCL11A, BCL3, BCLAF1, BHLHE40, CEBPB, CHD1, CHD2, CREB1, CTCF, CUX1, EBF1, EGR1, ELF1, ELK1, EZH2, FOXM1, GATA1, GTF2F1, H2AFZ, HCFC1, IKZF1, IRF3, IRF4, JUND, KDM4A, KDM5A, MAX, MAZ, MTA3, MXI1, MYOD1, MYOG, NFATC1, NFE2, NFIC, NFYB, NRF1, PAX5, PHF8, PML, POLR2A, POU2F2, RAD21, RCOR1, RELA, RFX5, RUNX3, SAP30, SMC3, SP1, SRF, STAT1, STAT5A, TAF1, TBL1XR1, TBP, TCF12, USF2, WRNIP1, YY1, ZBTB7A, ZC3H11A, ZEB1, ZNF143, ZNF384 |
| *ANK1* | AR, STAT3, TP63, E2F1, RUNX1, MYC, SPI1, POU5F1, EGR1, TP53, GATA1, GATA2, KDM5B, RCOR3, SIN3B, REST, SETDB1, TFAP2C, SOX9, SRY, EP300, YAP1, SALL4, KLF1, SCLY, JARID2, MTF2, SUZ12, TBX3, BACH1, SOX11, CDX2, CEBPB, HOXD13, ATF2, BRCA1, CBX3, CCNT2, CHD1, CHD2, CHD7, CTCF, E2F6, ELF1, ETS1, EZH2, FOXP2, GABPA, GTF2F1, H2AFZ, HCFC1, HDAC2, HMGN3, JUND, KDM4A, KDM5A, MAFK, MAX, MAZ, MXI1, NANOG, NR2F2, PAX5, PHF8, POLR2A, RAD21, RBBP5, RCOR1, RXRA, SAP30, SIN3A, SIRT6, SP4, TBL1XR1, TBP, TCF12, TCF3, TEAD4, TRIM28, UBTF, USF1, USF2, YY1, ZBTB7A, ZNF143, ZNF263, ZNF384 |
| *TAS2R38* | STAT3, MYC, FOXP1, PPARD, SALL4, SOX17, PBX1, SOX11, IRF8, HTT, BHLHE40, CHD1, CHD2, CHD7, CTCF, CUX1, EP300, ETS1, EZH2, GTF2F1, H2AFZ, HDAC1, HDAC2, KAT2B, MAX, MAZ, MXI1, PHF8, POLR2A, RAD21, RBBP5, RCOR1, SAP30, SIN3A, SMC3, SUZ12, USF2, ZNF143 |
| *CFHR3* | FOXP1, POU3F2, BATF, CEBPB, CTCF, EZH2, POLR2A, STAT3 |
| *SVEP1* | AR, TCF4, TP63, NANOG, TP53, GATA1, GATA2, PPARD, SOX9, SRY, DMRT1, PRDM14, RUNX2, GATA4, EED, PHC1, EZH2, RNF2, JARID2, MTF2, SUZ12, LMO2, SOX11, BACH1, CEBPB, CHD1, CHD7, CTBP2, CTCF, E2F4, E2F6, EP300, FOS, FOXP2, GABPA, GATA3, H2AFZ, HDAC1, HDAC2, HDAC6, JUND, KDM4A, KDM5B, MAX, MXI1, MYC, NFIC, PBX3, PHF8, POLR2A, RAD21, RBBP5, REST, RFX5, SAP30, SIN3A, STAT3, TBP, TCF12, TEAD4, TRIM28, ZNF143, ZNF263 |
| *TMPRSS3* | ZNF217, STAT3, E2F1, SOX2, EGR1, FOXA2, HNF4A, REST, CUX1, TFAP2A, EWSR1, BACH1, PADI4, BHLHE40, BRCA1, CEBPB, CHD2, CTCF, E2F4, E2F6, EBF1, ELK1, EP300, EZH2, FOS, GTF2F1, GTF3C2, H2AFZ, HDAC2, HDAC6, JUND, KDM1A, KDM5A, KDM5B, MAFF, MAFK, MAX, MAZ, MXI1, MYC, NFYB, PHF8, POLR2A, RAD21, RBBP5, RCOR1, RFX5, SAP30, SMC3, STAT1, TAF1, TAL1, TBL1XR1, TBP, TCF7L2, TFAP2C, USF2, ZMIZ1 |

| | |
|---|---|
| *SUSD2* | STAT3, E2F1, KLF4, NANOG, POU5F1, SOX2, MITF, SETDB1, ASH2L, FOXP1, PPARD, TAL1, SOX9, SRY, ZFP42, FOXP2, RAD21, ESRRB, NR0B1, CEBPD, CHD1, CHD2, CHD7, CTCF, EP300, ESR1, EZH2, FOSL2, GABPA, GATA1, H2AFZ, HDAC2, HNF4A, HNF4G, KDM5A, MAFF, MAFK, MAX, MAZ, MXI1, MYBL2, MYC, NFIC, NR3C1, PHF8, POLR2A, RBBP5, RCOR1, RFX5, SIRT6, SMC3, TAF1, TBP, TCF12, TEAD4, ZBTB7A, ZNF143, ZNF263 |
| *SNCAIP* | CREM, SPI1, SOX2, REST, SRY, EP300, ESR1, SALL4, WT1, TEAD4, GATA4, MEF2A, MTF2, SUZ12, JUN, RELA, BACH1, CBX2, CBX8, CEBPB, CHD1, CHD2, CREB1, CTBP2, CTCF, E2F4, EGR1, EZH2, FOS, GABPA, GATA1, GATA2, H2AFZ, HDAC1, HDAC2, HDAC6, HMGN3, JUND, KDM4A, KDM5A, MAX, MAZ, MXI1, MYC, PAX5, PHF8, POLR2A, RAD21, RBBP5, RNF2, SAP30, SIN3A, STAT3, TAF1, TBP, TRIM28, UBTF, USF1, USF2, ZBTB7A, ZNF143, ZNF263 |
| *ALDH3B1* | KDM5A, STAT3, RUNX1, MYC, SPI1, SOX2, TP53, PPARG, HNF4A, FOXP1, SOX9, SRY, OLIG2, ATF3, MECOM, RAD21, TFAP2A, LMO2, HSF1, STAT5A, CEBPB, TFEB, IRF8, ARID3A, ATF1, BHLHE40, BRCA1, CBX3, CCNT2, CEBPD, CHD1, CHD2, CREB1, CTCF, CTCFL, CUX1, E2F4, E2F6, ELF1, ELK1, ELK4, EP300, ESR1, ETS1, EZH2, FOS, FOSL2, FOXA1, FOXA2, FOXP2, GABPA, GATA1, GATA2, GTF2F1, H2AFZ, HDAC2, HDAC6, HMGN3, HNF4G, IRF1, JUND, KDM1A, KDM5B, MAFF, MAFK, MAX, MAZ, MBD4, MXI1, MYBL2, MYOD1, MYOG, NELFE, NFIC, NR2F2, NR3C1, PBX3, PHF8, POLR2A, PRDM1, RBBP5, RCOR1, REST, RFX5, SAP30, SIN3A, SMARCB1, SMARCC2, SMC3, SP1, TAF1, TAL1, TBL1XR1, TBP, TCF3, TEAD4, TFAP2C, TRIM28, UBTF, USF1, USF2, YY1, ZBTB33, ZBTB7A, ZEB1, ZKSCAN1, ZMIZ1, ZNF143, ZNF263, ZNF384 |
| *GBP7* | SPI1, E2F4, SOX9, STAT4, POU3F2, STAT5A, CEBPA, IRF8, BHLHE40, CHD2, CTCF, CUX1, EP300, ETS1, EZH2, H2AFZ, JUN, JUND, KDM5A, MAFK, MAX, MAZ, MXI1, POLR2A, PRDM1, RCOR1, SIN3A, STAT3, SUPT20H, USF2, ZNF384 |
| *FAM178B* | AHR, ARNT, ZNF217, TCF4, SMAD4, TP63, CREM, FLI1, MYC, MITF, GATA2, RCOR3, SETDB1, TET1, EP300, SMARCA4, TEAD4, SRF, RBPJ, BHLHE40, CHD1, CTCF, E2F6, EZH2, GABPA, GATA1, H2AFZ, HDAC2, MAX, MYOG, NELFE, POLR2A, RAD21, RBBP5, REST, RNF2, SIRT6, SMC3, ZC3H11A, ZNF143 |
| *TNFAIP2* | AR, RARA, NFE2L2, PHF8, TCF4, TP63, CREM, SPI1, SOX2, EGR1, MITF, GATA1, EOMES, PPARD, TAL1, CNOT3, SOX9, ESR1, PRDM14, MECOM, EZH2, RNF2, JARID2, MTF2, SUZ12, RELA, TAF7L, BACH1, BHLHE40, CCNT2, CEBPB, CHD1, CTCF, E2F4, E2F6, EBF1, EP300, ETS1, GABPA, GATA2, H2AFZ, HDAC2, KDM4A, KDM5A, KDM5B, MAX, MAZ, MXI1, MYB, MYC, MYOG, NELFE, PBX3, POLR2A, RAD21, RCOR1, REST, SIN3A, SMC3, STAT5A, TAF1, TBP, TCF12, USF2, YY1, ZMIZ1, ZNF143 |
| *SORCS2* | VDR, AR, STAT3, TCF4, SMAD4, TP63, CREB1, FLI1, RUNX1, MYC, KLF4, SOX2, EGR1, MITF, TP53, PPARG, GATA2, REST, TET1, TRIM28, EP300, ESR1, TEAD4, RUNX2, SCLY, MEF2A, CTCF, MTF2, SUZ12, JUN, TCF7L2, HOXD13, BACH1, CBX3, CCNT2, CHD1, CHD2, CHD7, CTBP2, CTCFL, E2F6, EZH2, FOXP2, GABPA, GTF2F1, H2AFZ, HCFC1, HDAC1, HDAC2, HDAC6, HMGN3, JUND, KDM4A, MAFK, MAX, MAZ, MXI1, MYOD1, MYOG, NELFE, NFIC, POLR2A, RAD21, RBBP5, RCOR1, RFX5, SAP30, SIN3A, SMC3, SRF, TBP, TCF12, UBTF, USF2, YY1, ZBTB7A, ZC3H11A, ZKSCAN1, ZMIZ1, ZNF143, ZNF263 |
| *SUSD3* | TP63, RUNX1, SPI1, EGR1, KDM5B, FOXP1, TAL1, SRY, EP300, SALL4, ZNF281, MTF2, SUZ12, NR1H3, BACH1, BCL3, BHLHE40, CBX3, CCNT2, CEBPD, CHD1, CHD2, CHD4, CHD7, CTCF, CUX1, E2F6, EBF1, ELF1, ELK1, ETS1, EZH2, GABPA, GATA1, H2AFZ, HCFC1, HDAC1, HDAC2, HDAC6, HMGN3, HNF4A, IKZF1, IRF1, JUN, JUND, KDM1A, KDM4A, KDM5A, MAX, MAZ, MTA3, MXI1, MYC, MYOD1, MYOG, NCOR1, NELFE, NFATC1, NFIC, NR2F2, NRF1, PAX5, PHF8, PML, POLR2A, RAD21, RBBP5, RCOR1, RFX5, RUNX3, RXRA, SAP30, SETDB1, SIN3A, SIRT6, SMC3, SP1, SP4, STAT1, STAT5A, TAF1, TBL1XR1, TBP, TCF12, TCF3, TEAD4, UBTF, USF1, USF2, WRNIP1, YY1, ZBTB7A, ZC3H11A, ZEB1, ZMIZ1, ZNF143, ZNF384 |
| *BOLL* | CREM, MYC, POU5F1, SOX2, ZFX, SETDB1, TET1, MYCN, ERG, MTF2, SUZ12, RCOR1, DNAJC2, ATF2, BACH1, BCL3, CEBPB, CHD1, CHD2, CREB1, CTBP2, CTCF, E2F6, ELF1, EP300, EZH2, GABPA, GTF2F1, H2AFZ, HCFC1, HDAC2, JUND, KDM4A, MAX, MXI1, PBX3, PHF8, POLR2A, RAD21, RBBP5, SIN3A, SMC3, TBP, TCF12, TCF3, TEAD4, USF2, YY1, ZEB1, ZNF143 |
| *LHX1* | NFE2L2, STAT3, TCF4, SMAD4, CREB1, E2F1, RUNX1, MYC, KLF4, NANOG, POU5F1, SOX2, TP53, KDM5B, REST, SETDB1, EOMES, TET1, ERG, WT1, ZNF281, SMAD3, RUNX2, MYBL2, TBX5, BMI1, EZH2, RNF2, JARID2, MTF2, SUZ12, ZIC3, BACH1, BCL3, CBX3, CBX8, CEBPB, CHD1, CHD2, CTBP2, CTCF, E2F4, E2F6, EGR1, ELF1, EP300, ETS1, FOXA2, GABPA, GTF2F1, H2AFZ, HDAC1, HDAC2, HDAC6, HMGN3, HNF4A, JUN, JUND, KDM4A, MAX, MAZ, MXI1, NFYB, PHF8, PML, POLR2A, RAD21, RBBP5, RCOR1, SAP30, SIN3A, SMARCC1, SMC3, SP1, TAF1, TBP, TCF12, TCF7L2, TRIM28, UBTF, USF2, YY1, ZBTB7A, ZKSCAN1, ZNF143, ZNF263, ZNF384 |
| *TSPAN2* | ZNF217, NFE2L2, VDR, TCF4, SMAD4, TP63, RUNX1, SOX2, TCF3, GATA2, SETDB1, ERG, TRIM28, SOX9, SMARCA4, RUNX2, TCF7, SUZ12, MEIS1, EWSR1, ATF2, BACH1, BHLHE40, CCNT2, CHD1, CHD2, CREB1, CTCF, EGR1, EP300, ETS1, EZH2, GATA3, GTF2F1, H2AFZ, HDAC1, HDAC2, HDAC6, JUN, JUND, KDM4A, KDM5A, MAFK, MAX, MAZ, MXI1, MYC, MYOG, PHF8, POLR2A, RAD21, RBBP5, RCOR1, REST, SAP30, SIN3A, SMC3, SP1, SRF, TAF1, TAF7, TBP, TCF12, UBTF, USF2, ZBTB7A, ZNF143, ZNF263 |
| *CSAG3* | CTNNB1, CTCF |
| *NEURL3* | STAT3, FLI1, RUNX1, GATA2, SIN3B, PPARD, GFI1B, TAL1, FOXO3, HOXB4, BMI1, SUZ12, CTNNB1, LMO2, LYL1, MEIS1, HSF1, ASXL1, ARID3A, ATF3, BACH1, BATF, BCL3, BCLAF1, BHLHE40, BRCA1, CBX3, CCNT2, CEBPB, CHD1, CHD2, CHD7, CREB1, CTCF, CTCFL, CUX1, E2F4, E2F6, EBF1, ELF1, ELK1, EP300, ESR1, EZH2, FOS, FOSL2, FOXM1, FOXP2, GATA1, GTF2F1, H2AFZ, HCFC1, HDAC1, HDAC2, HDAC6, HMGN3, HNF4A, HNF4G, IRF1, IRF3, IRF4, JUN, JUND, KDM4A, KDM5B, MAFK, MAX, MAZ, MBD4, MTA3, MXI1, MYBL2, MYC, NANOG, NFIC, NFYB, NRF1, PAX5, PML, POLR2A, POU2F2, RAD21, RBBP5, RCOR1, RELA, RFX5, RNF2, RUNX3, SAP30, SETDB1, SIN3A, SIRT6, SMC3, SP1, SPI1, STAT1, STAT5A, TBL1XR1, TBP, TCF12, TCF3, TEAD4, TRIM28, UBTF, USF2, WHSC1, WRNIP1, YY1, ZBTB7A, ZC3H11A, ZEB1, ZKSCAN1, ZNF143, ZNF263, ZNF384 |

| | |
|---|---|
| *RGL3* | SP1, TFAP2A, TP53, SETDB1, TFAP2C, TET1, BMI1, EED, PHC1, EZH2, RNF2, JARID2, MTF2, SUZ12, RBPJ, BACH1, BHLHE40, BRCA1, CEBPB, CEBPD, CHD1, CHD2, CTCF, EBF1, EGR1, ELF1, EP300, ESR1, GATA2, GATA3, H2AFZ, HDAC1, HDAC2, HMGN3, JUN, JUND, KDM4A, KDM5A, KDM5B, MAFK, MAX, MAZ, MXI1, MYC, NANOG, PHF8, POLR2A, RAD21, RBBP5, RCOR1, RFX5, SAP30, SIN3A, SMC3, TAF1, TBP, TCF12, TEAD4, TRIM28, USF1, USF2, WRNIP1, YY1, ZBTB7A, ZEB1, ZKSCAN1, ZNF143 |
| *SPP1* | ETS1, ETS2, ETV4, JUN, LEF1, POU2F1, POU2F2, SMAD1, SP1, CEBPD, E2F1, RUNX1, SPI1, KLF4, NANOG, POU5F1, SOX2, TCF3, MITF, PPARG, GATA2, ASH2L, FOXP1, YAP1, SALL4, PRDM14, ATF3, MYBL2, CHD1, POU3F2, RELA, NACC1, NR0B1, RBPJ, NR1H3, BACH1, CBX2, CBX8, CEBPB, CHD7, CTCF, CUX1, E2F4, EBF1, EP300, EZH2, FOS, FOXP2, GTF2F1, H2AFZ, HDAC1, HDAC2, MAFF, MAFK, MAX, MAZ, MXI1, MYC, MYOD1, NFIC, POLR2A, RAD21, RBBP5, RCOR1, RFX5, RNF2, RUNX3, SAP30, SIN3A, SMC3, STAT3, TAF1, TBP, TCF12, TEAD4, USF1, YY1, ZNF143, ZNF384 |
| *CDH18* | ZNF217, CEBPD, ELK1, AR, STAT3, TCF4, TP63, SOX2, TP53, PPARG, FOXA2, HNF4A, PPARD, MYCN, YAP1, RUNX2, SUZ12, JUN, BACH1, CEBPB, CHD1, CTCF, EP300, ESR1, EZH2, FOXP2, H2AFZ, HDAC1, HDAC2, HDAC6, JUND, KDM4A, KDM5B, MAX, NFYB, PBX3, POLR2A, RAD21, RBBP5, REST, SAP30, SIN3A, SPI1, TAF1, TBP, TCF12, ZNF143 |
| *CLDN3* | NFE2L2, AR, RUNX1, KLF4, NANOG, POU5F1, SOX2, TCF3, EGR1, FOXA2, HNF4A, GATA2, KDM5B, ZFX, RCOR3, SIN3B, ASH2L, EOMES, TFAP2C, TET1, TAL1, CNOT3, SRY, SMARCA4, YAP1, ZNF281, TFCP2L1, SMAD2, SMAD3, SCLY, BMI1, EED, PHC1, EZH2, RNF2, JARID2, MTF2, SUZ12, BACH1, SMAD1, GATA3, STAT1, SREBF1, ATF2, BHLHE40, BRCA1, CCNT2, CHD1, CHD2, CHD7, CREB1, CTCF, E2F6, ELF1, EP300, FOXA1, GABPA, GTF2F1, H2AFZ, HCFC1, HDAC2, HDAC6, HMGN3, JUND, KDM4A, MAX, MAZ, MXI1, MYC, MYOG, NFIC, NRF1, PHF8, PML, POLR2A, RAD21, RBBP5, RFX5, SAP30, SIN3A, SIRT6, SP1, SP4, TAF1, TAF7, TBP, TCF12, TCF7L2, TEAD4, USF1, USF2, YY1, ZBTB7A, ZNF143, ZNF263 |
| *SUSD1* | AHR, ARNT, NFE2L2, STAT3, TCF4, SMAD4, TP63, SPI1, NANOG, EGR1, MITF, TP53, PPARG, FOXA2, HNF4A, GATA2, PPARD, FOXO3, SMAD2, SMAD3, JARID2, SUZ12, NFIB, FOXO1, ZNF274, ARID3A, ATF2, BACH1, BHLHE40, BRCA1, CBX3, CCNT2, CEBPB, CEBPD, CHD1, CHD2, CHD4, CHD7, CREB1, CREBBP, CTBP2, CTCF, CUX1, E2F4, E2F6, EBF1, ELF1, ELK1, ELK4, EP300, ETS1, EZH2, FOS, FOXA1, FOXP2, GABPA, GATA1, GATA3, GTF2B, GTF2F1, H2AFZ, HCFC1, HDAC1, HDAC2, HDAC6, HMGN3, IRF1, JUN, JUND, KDM1A, KDM4A, KDM5B, MAFK, MAX, MAZ, MEF2A, MTA3, MXI1, MYC, NCOR1, NELFE, NFIC, NFYB, NR2F2, NR3C1, NRF1, PAX5, PHF8, PML, POLR2A, RAD21, RBBP5, RCOR1, RELA, REST, RFX5, RUNX3, SAP30, SETDB1, SIN3A, SIRT6, SMC3, SRF, TAF1, TAL1, TBL1XR1, TBP, TCF12, TCF3, TCF7L2, TEAD4, UBTF, USF2, YY1, ZBTB7A, ZC3H11A, ZEB1, ZMIZ1, ZNF143, ZNF384 |
| *ADD2* | AHR, ARNT, AR, MYC, POU5F1, EGR1, GATA1, GATA2, FOXP1, TAL1, YAP1, KLF1, RUNX2, SRF, SUZ12, LMO2, CRX, IRF8, BACH1, BHLHE40, CBX3, CCNT2, CEBPB, CHD1, CHD2, CHD7, CREB1, CTCF, CTCFL, E2F4, E2F6, ELF1, ELK1, EP300, EZH2, FOSL2, FOXM1, GABPA, GATA3, GTF2B, GTF2F1, H2AFZ, HCFC1, HDAC1, HDAC2, HDAC6, HMGN3, JUN, JUND, KDM1A, KDM4A, KDM5B, MAFF, MAFK, MAX, MAZ, MXI1, NFYB, PAX5, PHF8, POLR2A, RAD21, RBBP5, RCOR1, REST, RFX5, RNF2, RXRA, SAP30, SIN3A, SIRT6, SMC3, SP1, SP4, SPI1, STAT3, TAF1, TAF7, TBL1XR1, TBP, TCF12, TEAD4, TRIM28, UBTF, USF1, USF2, YY1, ZBTB7A, ZC3H11A, ZMIZ1, ZNF143, ZNF263 |
| *SPTA1* | AR, RUNX1, SPI1, POU5F1, GATA1, GATA2, REST, ERG, GFI1B, TAL1, CUX1, YY1, GATA4, RAD21, LMO2, LYL1, POU3F2, RCOR1, ELF1, ARID3A, ATF1, ATF3, BACH1, CCNT2, CEBPB, CEBPD, CHD1, CTCF, EP300, ETS1, EZH2, FOS, GABPA, GTF2B, H2AFZ, HDAC1, HMGN3, IRF1, JUN, JUND, KDM1A, KDM5B, MAFF, MAFK, MYC, NFE2, PHF8, POLR2A, RBBP5, RFX5, SAP30, STAT5A, TBL1XR1, TBP, TEAD4, TRIM28, UBTF, USF2, ZMIZ1, ZNF384 |
| *IRF1* | NFKB1, STAT1, STAT3, STAT4, NFE2L2, PHF8, AR, TCF4, TP63, CREB1, E2F1, FLI1, RUNX1, MYC, SPI1, KLF4, NANOG, POU5F1, SOX2, KDM5B, EOMES, TFAP2C, FOXP1, TET1, ERG, GFI1B, SOX9, SRY, FOXO3, ZNF281, TFCP2L1, MYB, MEIS1, NUCKS1, RELA, ELF5, STAT5A, ZIC3, RBPJ, CRX, CHD7, ARID3A, ATF1, ATF2, ATF3, BACH1, BCL3, BCLAF1, BHLHE40, BRCA1, CBX3, CCNT2, CEBPB, CEBPD, CHD1, CHD2, CHD4, CTBP2, CTCF, CTCFL, CUX1, E2F4, E2F6, EBF1, EGR1, ELF1, ELK1, EP300, ETS1, EZH2, FOS, FOSL2, FOXA1, FOXA2, FOXM1, FOXP2, GABPA, GATA1, GTF2B, GTF2F1, H2AFZ, HCFC1, HDAC1, HDAC2, HDAC6, HMGN3, IRF1, IRF3, JUN, JUND, KAT2A, KDM4A, KDM5A, MAFK, MAX, MAZ, MTA3, MXI1, MYBL2, MYOD1, MYOG, NELFE, NFATC1, NFIC, NFYA, NFYB, NR2F2, NRF1, PAX5, PBX3, PML, POLR2A, POU2F2, RAD21, RBBP5, RCOR1, REST, RFX5, RNF2, RUNX3, SAP30, SIN3A, SMARCB1, SMC3, SP1, SP4, SRF, STAT2, SUZ12, TAF1, TAF7, TAL1, TBL1XR1, TBP, TCF12, TCF3, TCF7L2, TEAD4, TFAP2A, UBTF, USF1, USF2, WRNIP1, YY1, ZBTB7A, ZC3H11A, ZEB1, ZKSCAN1, ZMIZ1, ZNF143, ZNF263, ZNF384 |
| *IL12RB1* | E2F1, AR, STAT3, FLI1, RUNX1, POU5F1, TP53, KDM5B, REST, SRY, STAT4, MEIS1, RBPJ, IRF1, IRF8, RCOR2, FOXO1, DROSHA, ATF2, BATF, BCL11A, BCLAF1, BHLHE40, CEBPB, CHD1, CHD2, CTCF, EBF1, ELF1, ELK1, EP300, ETS1, EZH2, FOXM1, H2AFZ, HDAC2, IKZF1, IRF4, JUND, MAX, MAZ, MTA3, MXI1, MYC, NELFE, NFATC1, NFIC, PAX5, PML, POLR2A, POU2F2, RBBP5, RCOR1, RELA, RUNX3, SAP30, SIN3A, SMC3, SP1, SPI1, STAT1, STAT5A, TAF1, TBL1XR1, TBP, USF2, YY1, ZC3H11A, ZEB1, ZNF143, ZNF384 |
| *GRIN2D* | ESR1, ESR2, ETS1, KDM5A, CREM, E2F1, FLI1, RUNX1, MYC, SPI1, TCF3, EGR1, HNF4A, ZFX, SIN3B, REST, SETDB1, SRY, SALL4, YY1, JARID2, SUZ12, ESRRB, RCOR1, RBPJ, NR4A2, BCL3, BHLHE40, CBX3, CCNT2, CEBPB, CHD1, CHD2, CHD7, CREB1, CTBP2, CTCF, E2F6, ELF1, EP300, EZH2, FOSL2, GABPA, GATA1, H2AFZ, HCFC1, HDAC1, HDAC2, HDAC6, HMGN3, JUND, KDM4A, KDM5B, MAX, MAZ, MXI1, MYOD1, MYOG, NELFE, NR3C1, PBX3, PHF8, POLR2A, RAD21, RBBP5, RNF2, SAP30, SIN3A, STAT5A, TAF1, TBP, TCF12, UBTF, ZBTB7A, ZNF263 |
| *TTC6* | E2F1, FOXA2, HNF4A, POU3F2, ELF5 |
| *FRMD3* | AR, STAT3, TCF4, SMAD4, FLI1, RUNX1, EGR1, MITF, TP53, FOXA2, HNF4A, GATA1, KDM5B, REST, TFAP2C, OLIG2, SALL4, TEAD4, MTF2, SUZ12, SOX11, DNAJC2, ARID3A, BACH1, BRCA1, CEBPB, CEBPD, CHD1, CHD2, CHD7, CREB1, CTBP2, CTCF, E2F4, E2F6, ELK1, EP300, EZH2, FOXP2, GABPA, GATA3, GTF2F1, H2AFZ, HCFC1, HDAC2, HNF4G, JUND, |

| | |
|---|---|
| | KDM4A, KDM5A, MAFK, MAX, MAZ, MXI1, MYBL2, MYC, NFIC, NFYA, NR3C1, PHF8, POLR2A, RAD21, RBBP5, RCOR1, RFX5, SAP30, SETDB1, SIN3A, SIRT6, SMC3, TAF1, TBP, TCF12, TCF7L2, TFAP2A, TRIM28, USF1, YY1, ZBTB7A, ZKSCAN1, ZNF143, ZNF263 |
| *UNC13D* | DACH1, TCF4, FLI1, KLF4, TCF3, EGR1, MITF, TP53, FOXA2, HNF4A, GATA1, GATA2, SETDB1, FOXP1, TET1, TAL1, ESR1, MYBL2, YY1, MECOM, TCF7, EZH2, TFAP2A, RELA, SOX11, SMAD1, SREBF2, TFEB, STAT1, ARID3A, ATF1, ATF2, BCL3, BCLAF1, BHLHE40, BRCA1, CBX3, CCNT2, CEBPB, CEBPD, CHD1, CHD2, CHD4, CHD7, CREB1, CTCF, CUX1, E2F4, E2F6, EBF1, ELF1, ELK1, ELK4, EP300, ETS1, FOS, FOSL2, FOXA1, FOXM1, GABPA, GATA3, GTF2B, GTF2F1, H2AFZ, HCFC1, HDAC1, HDAC2, HDAC6, HMGN3, IRF1, IRF4, JUN, JUND, KAT2B, KDM1A, KDM5A, KDM5B, MAFK, MAX, MAZ, MBD4, MTA3, MXI1, MYC, NCOR1, NELFE, NFIC, NFYA, NR2F2, NR3C1, NRF1, PAX5, PBX3, PHF8, PML, POLR2A, POU2F2, PRDM1, RAD21, RBBP5, RCOR1, REST, RFX5, RUNX3, SAP30, SIN3A, SIRT6, SIX5, SMARCB1, SMARCC1, SMC3, SP1, SPI1, SRF, STAT3, STAT5A, TAF1, TBL1XR1, TBP, TCF12, TCF7L2, TEAD4, TFAP2C, TRIM28, UBTF, USF1, USF2, WRNIP1, ZBTB33, ZBTB7A, ZC3H11A, ZEB1, ZKSCAN1, ZMIZ1, ZNF143, ZNF217, ZNF263, ZNF384 |
| *PIGR* | AR, NFKB1, RELA, RELB, USF1, USF2, TCF4, TP63, SPI1, GATA1, SALL4, SOX17, STAT5A, CDX2, PRDM16, ATF3, BHLHE40, CHD1, CTCF, EZH2, H2AFZ, HDAC2, HDAC6, KDM5B, MAX, MAZ, POLR2A, RCOR1, TAL1, TEAD4, YY1, ZKSCAN1 |
| *EFHD1* | ZNF217, TCF4, TP63, RUNX1, POU5F1, EGR1, MITF, TP53, TFAP2C, PPARD, TRIM28, ESR1, SCLY, BMI1, EED, PHC1, EZH2, RNF2, JARID2, MTF2, SUZ12, SMAD1, RBPJ, CLOCK, ESR2, ARID3A, ATF2, BACH1, BCL3, BCLAF1, BHLHE40, BRCA1, CCNT2, CEBPB, CHD1, CHD2, CHD7, CREB1, CTCF, CTCFL, CUX1, E2F4, EBF1, ELF1, ELK1, EP300, FOS, FOSL2, FOXA2, FOXM1, FOXP2, GABPA, GATA3, GTF2F1, H2AFZ, HCFC1, HDAC1, HDAC2, HMGN3, JUN, JUND, KDM4A, KDM5A, KDM5B, MAFF, MAFK, MAX, MAZ, MBD4, MTA3, MXI1, MYBL2, MYC, NFIC, NFYB, NR2F2, NRF1, PAX5, PHF8, PML, POLR2A, POU2F2, RAD21, RBBP5, RCOR1, REST, RFX5, RUNX3, SETDB1, SIN3A, SIX5, SMC3, SP1, SP4, SRF, STAT1, STAT3, STAT5A, TAF1, TBL1XR1, TBP, TCF12, TCF7L2, TEAD4, UBTF, USF2, WRNIP1, YY1, ZC3H11A, ZEB1, ZKSCAN1, ZMIZ1, ZNF143, ZNF263, ZNF384 |
| *CCDC87* | DACH1, FLI1, MYC, PPARG, HNF4A, E2F4, SIN3B, TET1, ERG, TFCP2L1, SRF, ESRRB, NR0B1, SMAD1, ARID3A, ATF2, ATF3, BACH1, BCL3, BCLAF1, BHLHE40, BRCA1, CBX3, CCNT2, CEBPB, CEBPD, CHD1, CHD2, CHD4, CHD7, CREB1, CTCF, CTCFL, CUX1, E2F6, EBF1, ELF1, ELK1, EP300, ETS1, EZH2, FOS, FOSL2, FOXA1, FOXA2, FOXM1, FOXP2, GATA1, GTF2B, GTF2F1, H2AFZ, HCFC1, HDAC1, HDAC2, HDAC6, HMGN3, HNF4G, JUN, JUND, KAT2A, KAT2B, KDM4A, KDM5B, MAFF, MAFK, MAX, MAZ, MBD4, MTA3, MXI1, MYB, MYBL2, MYOG, NELFE, NFE2, NFIC, NFYA, NFYB, NR2F2, NRF1, PAX5, PHF8, PML, POLR2A, POU2F2, RAD21, RBBP5, RCOR1, RELA, REST, RFX5, RUNX3, SAP30, SETDB1, SIN3A, SMARCB1, SMARCC1, SMC3, SP1, SP4, SPI1, STAT1, STAT3, STAT5A, TAF1, TAF7, TAL1, TBL1XR1, TBP, TCF12, TCF3, TCF7L2, TEAD4, THAP1, TRIM28, UBTF, USF1, USF2, WRNIP1, YY1, ZBTB7A, ZC3H11A, ZEB1, ZKSCAN1, ZMIZ1, ZNF143, ZNF263, ZNF384 |
| *OR2B6* | FLI1, MYC, POU3F2, RCOR1, CEBPB, HTT, EZH2, H2AFZ, MAFK, POLR2A, RBBP5 |
| *DAGLA* | ZNF217, AR, STAT3, E2F1, RUNX1, MYC, SPI1, NANOG, EGR1, TP53, FOXA2, HNF4A, REST, TFAP2C, SOX9, DMRT1, RAD21, MTF2, SUZ12, TFAP2A, TBX3, TAF7L, DNAJC2, FOXO1, ARID3A, ATF3, BACH1, BHLHE40, BRCA1, CBX3, CCNT2, CEBPB, CHD2, CREB1, CTCF, CTCFL, E2F4, E2F6, EBF1, ELF1, ELK1, EP300, ETS1, EZH2, FOS, FOSL1, FOSL2, FOXA1, FOXM1, FOXP2, GABPA, GATA2, GATA3, GTF2B, GTF2F1, H2AFZ, HCFC1, HDAC1, HDAC2, HDAC6, HMGN3, HNF4G, IRF1, JUN, JUND, KDM4A, KDM5A, MAFF, MAFK, MAX, MAZ, MBD4, MTA3, MXI1, MYBL2, MYOD1, MYOG, NFIC, NR2F2, NRF1, PAX5, PHF8, POLR2A, POU2F2, RBBP5, RCOR1, RFX5, RUNX3, RXRA, SETDB1, SIN3A, SIX5, SMC3, SP1, SP4, STAT1, TAF1, TBL1XR1, TBP, TCF12, TEAD4, TRIM28, UBTF, USF1, USF2, WRNIP1, YY1, ZBTB7A, ZC3H11A, ZEB1, ZKSCAN1, ZMIZ1, ZNF143, ZNF263 |
| *HIST1H3G* | AR, TP63, FLI1, MYC, SPI1, KLF4, POU5F1, SOX2, EGR1, ZFX, E2F4, EOMES, ERG, TAL1, CNOT3, SRY, CUX1, CCND1, HOXB4, MYBL2, CHD1, ZFP42, EED, RNF2, HSF1, TBP, SMAD1, CEBPB, IRF8, ARID3A, ATF2, BACH1, BATF, BCL11A, BCL3, BCLAF1, BHLHE40, BRCA1, CBX3, CHD2, CHD7, CREB1, CTCF, EBF1, ELF1, ELK1, ELK4, EP300, ETS1, EZH2, FOS, FOXA2, FOXM1, GATA1, GATA2, GTF2F1, H2AFZ, HCFC1, HDAC2, HDAC6, HMGN3, HNF4G, IRF3, IRF4, JUND, KAT2A, KDM4A, KDM5A, KDM5B, MAFF, MAFK, MAX, MAZ, MBD4, MEF2A, MTA3, MXI1, NELFE, NFATC1, NFIC, NFYA, NFYB, NR2F2, NRF1, PAX5, PBX3, PHF8, PML, POLR2A, POU2F2, RAD21, RBBP5, RCOR1, RELA, REST, RFX5, RUNX3, RXRA, SAP30, SETDB1, SIN3A, SIRT6, SIX5, SMC3, SP1, SP4, SRF, STAT1, STAT3, STAT5A, SUPT20H, TAF1, TAF7, TBL1XR1, TCF12, TCF3, TCF7L2, UBTF, USF1, USF2, WRNIP1, YY1, ZBTB33, ZEB1, ZKSCAN1, ZMIZ1, ZNF143, ZNF384 |
| *APOL3* | AR, E2F1, SOX2, PPARD, NR3C1, RUNX2, CUX1, TFAP2A, STAT1, IRF8, CBX2, CBX8, CEBPB, EP300, EZH2, FOXA1, FOXA2, H2AFZ, HDAC6, JUN, JUND, KDM5A, MAFF, MAFK, POLR2A, RNF2, SMC3 |
| *LGALS9* | CREB1, MYC, SPI1, NANOG, POU5F1, SOX2, EGR1, REST, ASH2L, GFI1B, ESR1, SMARCA4, SMAD2, SMAD3, HOXB4, YY1, MECOM, SRF, RAD21, JUN, PBX1, BATF, BCL11A, BCL3, BCLAF1, BHLHE40, CEBPB, CHD1, CTCF, E2F6, EBF1, ELF1, EP300, ETS1, EZH2, FOS, GATA1, H2AFZ, HDAC1, HDAC2, IRF1, IRF4, MAFK, MAX, MAZ, MTA3, MXI1, NELFE, PAX5, PBX3, PML, POLR2A, POU2F2, RBBP5, RCOR1, RUNX3, SIN3A, SMC3, SP1, STAT2, TAF1, TBP, TCF12, TCF3, USF1, USF2, WHSC1, WRNIP1, ZMIZ1, ZNF143 |
| *PHF2* | TCF4, FLI1, RUNX1, EGR1, TP53, GATA1, GATA2, RCOR3, REST, ASH2L, TFAP2C, PPARD, TAL1, EP300, YAP1, RUNX2, MTF2, CTNNB1, JUN, PRDM5, DROSHA, BACH1, BHLHE40, BRCA1, CCNT2, CEBPB, CHD1, CHD2, CHD7, CREB1, CTBP2, CTCF, E2F4, E2F6, EBF1, ELF1, ELK1, ETS1, EZH2, FOXA1, GATA3, GTF2B, GTF2F1, H2AFZ, HCFC1, HDAC1, HDAC2, HDAC6, HMGN3, HNF4A, IRF1, JUND, KAT2A, KDM4A, KDM5B, MAX, MAZ, MTA3, MXI1, MYC, MYOD1, NELFE, NFATC1, NFIC, NR2F2, PAX5, PHF8, PML, POLR2A, RAD21, RBBP5, RCOR1, RELA, RFX5, RNF2, RUNX3, SAP30, SIN3A, SMARCB1, SMC3, SPI1, STAT3, STAT5A, SUZ12, TAF1, TBL1XR1, TBP, TCF12, TEAD4, UBTF, WRNIP1, YY1, ZBTB7A, ZC3H11A, ZMIZ1, ZNF143, ZNF217, ZNF263, ZNF384 |
| *WISP3* | AR, STAT3, TP63, TP53, MYCN, SMARCA4, PRDM14, LMO2, HIF1A, CBX2, CBX8, CEBPB, CHD1, CTCF, EZH2, H2AFZ, IRF1, MAFK, POLR2A, RBBP5, RNF2, SAP30 |

| | |
|---|---|
| *IGFBP3* | E2F2, FLI1, SMAD1, SP1, SP3, TFAP2A, TP53, ZNF217, NFE2L2, AR, STAT3, SMAD4, TP63, E2F1, NANOG, POU5F1, SOX2, TCF3, GATA2, SETDB1, EOMES, PPARD, TRIM28, EP300, ESR1, SMARCA4, YAP1, SALL4, PRDM14, ATF3, SMAD2, SMAD3, NR3C1, CTCF, RAD21, MTF2, SUZ12, RCOR1, BACH1, HIF1A, ARID3A, BCL3, BRCA1, CEBPB, CHD1, CHD2, CHD7, CREB1, CTBP2, E2F4, E2F6, EBF1, EGR1, ELF1, ELK1, EZH2, FOS, FOSL2, FOXA1, FOXA2, FOXM1, FOXP2, GATA3, GTF2F1, H2AFZ, HCFC1, HDAC2, HDAC6, JUND, KDM4A, KDM5B, MAFK, MAX, MAZ, MXI1, MYC, MYOG, NFIC, NRF1, PAX5, PHF8, POLR2A, RBBP5, REST, RFX5, SAP30, SIN3A, SIRT6, SIX5, SMARCB1, SMC3, STAT1, TAF1, TBP, TCF12, TCF7L2, TEAD4, USF1, USF2, WRNIP1, YY1, ZBTB33, ZBTB7A, ZC3H11A, ZEB1, ZKSCAN1, ZNF143, ZNF263 |
| *UNC93B1* | AR, TP63, RUNX1, MYC, SPI1, EGR1, MITF, HNF4A, E2F4, FOXP1, TET1, SOX9, SRY, ZNF281, RUNX2, CUX1, TCF7, MTF2, SUZ12, HSF1, CEBPB, NR1H3, ARID3A, ATF2, ATF3, BACH1, BCL3, BCLAF1, BHLHE40, BRCA1, CBX3, CCNT2, CHD1, CHD2, CREB1, CTCF, E2F6, EBF1, ELF1, ELK1, EP300, ETS1, EZH2, FOS, FOSL2, FOXA1, FOXA2, FOXP2, GABPA, GATA1, GATA2, GTF2F1, GTF3C2, H2AFZ, HCFC1, HDAC1, HDAC2, HDAC6, HMGN3, IRF1, JUND, KAT2A, KDM1A, KDM4A, KDM5A, KDM5B, MAFK, MAX, MAZ, MBD4, MTA3, MXI1, MYBL2, MYOG, NELFE, NFIC, NFYB, NRF1, PAX5, PHF8, PML, POLR2A, POU2F2, RAD21, RBBP5, RCOR1, RELA, REST, RFX5, RUNX3, SAP30, SIN3A, SMARCB1, SMARCC1, SMC3, SP1, STAT1, STAT2, STAT5A, TAF1, TAL1, TBL1XR1, TBP, TCF12, TCF3, TEAD4, TRIM28, UBTF, USF1, USF2, WRNIP1, YY1, ZBTB33, ZBTB7A, ZC3H11A, ZEB1, ZKSCAN1, ZMIZ1, ZNF143, ZNF384 |
| *PLCB1* | ZNF217, TCF4, SMAD4, E2F1, MYC, SOX2, MITF, TP53, PPARG, FOXA2, HNF4A, GATA1, TRIM28, SOX9, SRY, FOXO3, OLIG2, SMARCA4, TEAD4, TFCP2L1, SMAD3, RUNX2, CUX1, YY1, MTF2, SUZ12, TBX3, TAF7L, NR1I2, TCF7L2, HTT, ZNF274, ARID3A, BATF, BHLHE40, BRCA1, CEBPB, CEBPD, CHD1, CHD2, CHD7, CREB1, CTBP2, CTCF, E2F6, EBF1, ELF1, EP300, ESR1, EZH2, FOXA1, FOXM1, FOXP2, GATA2, GATA3, GTF2F1, H2AFZ, HCFC1, HDAC2, JUND, KDM4A, KDM5A, MAFF, MAFK, MAX, MAZ, MXI1, MYBL2, MYOG, NELFE, NFIC, NRF1, PAX5, PBX3, PHF8, POLR2A, POU2F2, RAD21, RBBP5, RCOR1, REST, RFX5, SAP30, SIN3A, SMC3, SP1, STAT1, STAT3, TAF1, TBP, TCF12, TCF3, USF2, WRNIP1, ZBTB7A, ZC3H11A, ZNF143 |
| *ZIC2* | AR, SMAD4, MYC, KLF4, NANOG, POU5F1, SOX2, TCF3, EGR1, MITF, TP53, KDM5B, SIN3B, SETDB1, EOMES, TFAP2C, FOXP1, TET1, TRIM28, EP300, ZNF281, TEAD4, CCND1, MYBL2, GATA4, BMI1, PHC1, EZH2, RNF2, JARID2, MTF2, SUZ12, EWSR1, SOX11, NACC1, NR0B1, ATF2, BACH1, BHLHE40, CBX3, CCNT2, CEBPB, CHD1, CHD2, CHD7, CREB1, CTBP2, CTCF, E2F1, E2F4, E2F6, ELF1, ELK1, ESR1, FOXA1, FOXA2, FOXP2, GABPA, GTF2B, GTF2F1, H2AFZ, HCFC1, HDAC1, HDAC2, HDAC6, HMGN3, IRF1, JUND, KDM1A, KDM4A, MAFK, MAX, MAZ, MXI1, MYOG, NFIC, NR2F2, NRF1, PHF8, PML, POLR2A, RAD21, RBBP5, RCOR1, REST, RFX5, SAP30, SIN3A, SMARCB1, SMARCC1, SMC3, SPI1, TAF1, TAF7, TAL1, TBL1XR1, TBP, TCF12, TCF7L2, TFAP2A, UBTF, USF1, USF2, WHSC1, YY1, ZBTB7A, ZKSCAN1, ZMIZ1, ZNF143, ZNF263, ZNF384 |
| *ERBB4* | ZNF217, PAX3, AR, STAT3, TCF4, TP63, MYC, POU5F1, SOX2, TCF3, EGR1, TP53, FOXA2, HNF4A, PPARD, MYCN, ERG, TRIM28, EP300, OLIG2, SMARCA4, TEAD4, GATA4, MEF2A, JARID2, MTF2, SUZ12, CTNNB1, TBX3, BACH1, BRCA1, CEBPB, CHD1, CHD2, CHD7, CTBP2, CTCF, E2F6, EZH2, FOXA1, H2AFZ, HDAC2, JUND, KDM4A, MAFF, MAFK, MAX, MAZ, MXI1, NFYA, PHF8, POLR2A, RAD21, RBBP5, RCOR1, RFX5, SAP30, SIN3A, SMARCB1, SMARCC1, SMC3, SP1, SPI1, TAF1, TBP, TCF12, TCF7L2, YY1, ZBTB7A, ZKSCAN1, ZNF143 |
| *MAFA* | DACH1, AR, TP63, CREM, FLI1, NANOG, POU5F1, SOX2, TCF3, EGR1, TP53, KDM5B, TET1, ERG, SALL4, ZNF281, CTCF, BMI1, EED, PHC1, EZH2, RNF2, JARID2, MTF2, SUZ12, RCOR1, EWSR1, SREBF2, DNAJC2, ARID3A, ATF3, BACH1, BCL3, BCLAF1, BHLHE40, BRCA1, CBX3, CCNT2, CEBPB, CEBPD, CHD2, CHD4, CHD7, CREB1, CTBP2, CTCFL, CUX1, E2F4, E2F6, EBF1, ELF1, ELK1, EP300, ESR1, ESRRA, ETS1, FOS, FOSL1, FOSL2, FOXA1, FOXA2, FOXM1, FOXP2, GABPA, GATA3, GTF2B, GTF2F1, H2AFZ, HCFC1, HDAC2, HDAC6, HMGN3, HNF4G, IRF1, JUN, JUND, KDM4A, MAFK, MAX, MAZ, MEF2A, MXI1, MYC, MYOD1, MYOG, NELFE, NFIC, NFYA, NFYB, NR2F2, NR3C1, NRF1, PAX5, PBX3, PHF8, POLR2A, POU2F2, RAD21, RBBP5, RELA, REST, RFX5, RUNX3, SAP30, SIN3A, SMARCB1, SMC3, SP1, SP2, SP4, SREBF1, SRF, STAT1, STAT3, TAF1, TAL1, TBL1XR1, TBP, TCF12, TCF7L2, TEAD4, TFAP2A, TFAP2C, THAP1, TRIM28, UBTF, USF1, USF2, WHSC1, WRNIP1, YY1, ZBTB7A, ZC3H11A, ZKSCAN1, ZMIZ1, ZNF143, ZNF263, ZNF384 |
| *PLCXD3* | HOXC9, AR, STAT3, TCF4, SMAD4, MITF, REST, SETDB1, TFAP2C, MTF2, SUZ12, HTT, CHD1, CHD2, CHD7, CTCF, E2F6, EP300, EZH2, FOXA1, FOXA2, FOXP2, GTF2F1, H2AFZ, HDAC2, KDM4A, KDM5B, MAX, MXI1, MYC, MYOG, PHF8, POLR2A, RAD21, RBBP5, SAP30, SIN3A, STAT1, TBP, TCF12, TEAD4, ZKSCAN1, ZNF143, ZNF263 |
| *FGF13* | AR, EGR1, HOXB7, MYB, STAT3, SMAD4, RUNX1, SOX2, TCF3, MITF, SOX9, SRY, SALL4, SMAD3, BMI1, MTF2, SUZ12, TFAP2A, JUN, BACH1, ARID3A, BHLHE40, CBX3, CCNT2, CHD1, CHD4, CREB1, CTBP2, CTCF, E2F4, E2F6, EP300, EZH2, GTF2B, GTF2F1, H2AFZ, HCFC1, HDAC1, HDAC2, HDAC6, HMGN3, IRF1, JUND, KDM4A, KDM5B, MAX, MAZ, MYC, NR2F2, PAX5, PHF8, PML, POLR2A, RAD21, RBBP5, RCOR1, REST, SAP30, SIN3A, SMC3, SPI1, TAF1, TBL1XR1, TBP, UBTF, WHSC1, YY1, ZBTB7A, ZC3H11A, ZMIZ1, ZNF143, ZNF263, ZNF384 |
| *EDIL3* | E2F1, AHR, ARNT, CEBPD, STAT3, SMAD4, TP63, MYC, POU5F1, SOX2, EGR1, FOXA2, HNF4A, REST, PPARD, ERG, SRY, EP300, SMAD2, SMAD3, FOXP2, PHC1, EZH2, RNF2, MTF2, SUZ12, ESRRB, TBX3, DNAJC2, HOXD13, ATF2, BACH1, BRCA1, CEBPB, CHD1, CHD2, CHD7, CREB1, CTBP2, CTCF, E2F6, FOXA1, GTF2F1, H2AFZ, HDAC2, JUND, KDM4A, KDM5B, MAX, MXI1, MYOG, PHF8, POLR2A, RAD21, RBBP5, SAP30, SIN3A, SP1, TAF1, TAF7, TBP, TEAD4, USF2, YY1, ZNF143, ZNF263 |

| | |
|---|---|
| *ZNF782* | ETS1, TCF4, HNF4A, FOXP1, TFEB, ATF1, ATF2, ATF3, BACH1, BCL3, BCLAF1, BHLHE40, BRCA1, CBX3, CCNT2, CEBPB, CEBPD, CHD1, CHD2, CHD7, CREB1, CTCF, CUX1, E2F4, E2F6, EBF1, EGR1, ELF1, ELK1, ELK4, EP300, EZH2, FOS, FOXM1, GABPA, GTF2F1, H2AFZ, HCFC1, HDAC1, HDAC2, HDAC6, HMGN3, IRF1, IRF3, JUN, JUND, KDM1A, KDM4A, KDM5A, KDM5B, MAFF, MAFK, MAX, MAZ, MTA3, MXI1, MYBL2, MYC, NFE2, NRF1, PAX5, PHF8, PML, POLR2A, POU2F2, RAD21, RBBP5, RCOR1, RELA, REST, RFX5, RUNX3, SAP30, SETDB1, SIN3A, SIRT6, SIX5, SMC3, SP1, SP4, SRF, STAT1, STAT3, TAF1, TAL1, TBL1XR1, TBP, TCF12, TCF7L2, UBTF, USF1, USF2, WHSC1, WRNIP1, YY1, ZBTB7A, ZC3H11A, ZMIZ1, ZNF143, ZNF384 |
| *TBX1* | AR, TP63, FLI1, SPI1, KLF4, POU5F1, TCF3, EGR1, TP53, GATA1, KDM5B, SIN3B, SETDB1, ERG, SOX9, MYBL2, BMI1, EZH2, RNF2, JARID2, MTF2, SUZ12, JUN, TBP, IRF1, IRF8, ARID3A, ATF3, BACH1, BHLHE40, CBX3, CCNT2, CEBPB, CHD2, CHD4, CHD7, CTBP2, CTCF, CTCFL, E2F4, E2F6, EBF1, ELF1, ELK1, EP300, ETS1, GABPA, GATA2, GATA3, GTF2F1, H2AFZ, HCFC1, HDAC2, HMGN3, JUND, KDM5A, MAFK, MAX, MAZ, MXI1, MYC, MYOG, NCOR1, NR2F2, PAX5, POLR2A, PRDM1, RAD21, RCOR1, REST, SAP30, SIN3A, SMC3, TBL1XR1, TEAD4, THAP1, UBTF, YY1, ZBTB7A, ZC3H11A, ZMIZ1, ZNF143, ZNF263, ZNF384 |
| *TSPAN12* | TCF4, TP63, MYC, POU5F1, SOX2, EGR1, TP53, PPARG, TRIM28, EP300, OLIG2, SMARCA4, SRF, MTF2, SUZ12, ESRRB, SREBF2, ARID3A, ATF2, BACH1, BATF, BCL11A, BCL3, BCLAF1, BHLHE40, BRCA1, CBX3, CCNT2, CEBPB, CHD1, CHD2, CHD7, CREB1, CTBP2, CTCF, CTCFL, CUX1, E2F4, E2F6, EBF1, ELF1, ELK1, ETS1, EZH2, FOSL1, FOXA1, FOXM1, GABPA, GATA3, GTF2F1, H2AFZ, HCFC1, HDAC1, HDAC2, HDAC6, HMGN3, IKZF1, IRF4, JUN, JUND, KAT2B, KDM1A, KDM4A, KDM5A, KDM5B, MAFK, MAX, MAZ, MTA3, MXI1, MYOD1, MYOG, NANOG, NFIC, NRF1, PAX5, PBX3, PHF8, POLR2A, POU2F2, RAD21, RBBP5, RCOR1, RELA, REST, RFX5, RNF2, RUNX3, SAP30, SIN3A, SIRT6, SMARCB1, SMC3, SP1, SP4, SPI1, STAT1, STAT3, SUPT20H, TAF1, TAF7, TAL1, TBL1XR1, TBP, TCF12, TCF3, TCF7L2, TEAD4, UBTF, USF2, WHSC1, WRNIP1, YY1, ZBTB7A, ZC3H11A, ZKSCAN1, ZMIZ1, ZNF143, ZNF263, ZNF384 |
| *SIX2* | PHF8, STAT3, TCF4, TP63, MYC, KLF4, POU5F1, SOX2, EGR1, TP53, KDM5B, SIN3B, SETDB1, FOXP1, YAP1, SALL4, WT1, ZNF281, PRDM14, NR3C1, SCLY, MYBL2, BMI1, EED, PHC1, EZH2, RNF2, JARID2, MTF2, SUZ12, NR0B1, ATF2, BACH1, BCL3, BHLHE40, BRCA1, CEBPB, CHD1, CHD2, CREB1, CTBP2, CTCF, E2F4, E2F6, ELK1, EP300, FOS, FOXA1, GABPA, GATA1, GATA3, GTF2F1, H2AFZ, HCFC1, HDAC2, HDAC6, IRF3, JUND, KDM4A, MAFK, MAX, MAZ, MXI1, MYOG, NANOG, NELFE, NFIC, NFYA, NFYB, NR2F2, PBX3, POLR2A, RAD21, RBBP5, RCOR1, REST, RFX5, SAP30, SIN3A, SIX5, SMARCB1, SMARCC1, SMC3, SP1, SP2, SP4, SRF, TAF1, TBP, TCF12, TCF7L2, TEAD4, TFAP2A, TFAP2C, TRIM28, UBTF, USF1, USF2, YY1, ZBTB7A, ZKSCAN1, ZNF143, ZNF217, ZNF263 |
| *NOS1AP* | ZNF217, NFE2L2, AR, STAT3, SMAD4, RUNX1, POU5F1, SOX2, EGR1, MITF, TP53, FOXA2, HNF4A, GATA1, SETDB1, TFAP2C, TAL1, TRIM28, EP300, YAP1, TEAD4, PRDM14, ATF3, MECOM, TCF7, JARID2, MTF2, SUZ12, TBX3, STAT5A, NR1I2, PADI4, BACH1, BRCA1, CBX2, CBX8, CCNT2, CEBPB, CEBPD, CHD1, CHD2, CHD7, CREB1, CTBP2, CTCF, CUX1, E2F4, E2F6, EBF1, ELF1, ELK1, EZH2, FOS, FOXA1, FOXP2, GATA2, GATA3, GTF2F1, H2AFZ, HCFC1, HDAC1, HDAC2, HDAC6, HMGN3, HNF4G, IRF1, JUND, KDM4A, KDM5A, KDM5B, MAFF, MAFK, MAX, MAZ, MXI1, MYC, MYOG, NELFE, PAX5, PHF8, POLR2A, PRDM1, RAD21, RBBP5, RCOR1, RFX5, RNF2, SAP30, SIN3A, SMARCB1, SMARCC1, SMC3, SP1, STAT1, SUPT20H, TAF1, TBL1XR1, TBP, TCF12, TCF7L2, UBTF, YY1, ZBTB7A, ZKSCAN1, ZMIZ1, ZNF143, ZNF263 |
| *FOXA1* | AR, ZNF217, TP63, E2F1, MYC, KLF4, POU5F1, SOX2, TCF3, EGR1, TP53, FOXA2, HNF4A, GATA2, KDM5B, SETDB1, EOMES, TFAP2C, PPARD, TRIM28, SOX9, YAP1, SMAD2, SMAD3, RAD21, BMI1, PHC1, EZH2, RNF2, JARID2, MTF2, SUZ12, PBX1, LMO2, POU3F2, ELF5, BACH1, HSF1, NR0B1, PADI4, ZNF263, ARID3A, ATF1, BCL3, BHLHE40, BRCA1, CBX3, CCNT2, CEBPB, CEBPD, CHD1, CHD2, CHD4, CHD7, CREB1, CTBP2, CTCF, CUX1, E2F4, E2F6, ELF1, ELK1, EP300, FOS, FOSL2, FOXA1, FOXM1, GABPA, GATA1, GATA3, GTF2F1, H2AFZ, HCFC1, HDAC1, HDAC2, HMGN3, HNF4G, IRF1, JUN, JUND, KDM1A, KDM4A, MAFF, MAFK, MAX, MAZ, MBD4, MXI1, MYBL2, NANOG, NCOR1, NFIC, NFYB, NR2F2, PHF8, PML, POLR2A, RBBP5, RCOR1, REST, RFX5, RXRA, SAP30, SIN3A, SIRT6, SMARCB1, SMC3, SP1, SREBF1, SRF, STAT1, STAT3, TAF1, TAL1, TBL1XR1, TBP, TCF12, TCF7L2, TEAD4, UBTF, USF2, WHSC1, YY1, ZBTB7A, ZC3H11A, ZKSCAN1, ZMIZ1, ZNF143, ZNF384 |
| *ZHX2* | ZNF217, PAX3, VDR, STAT3, SMAD4, TP63, CREB1, RUNX1, MYC, SPI1, NANOG, POU5F1, SOX2, EGR1, TP53, HNF4A, GATA2, KDM5B, TFAP2C, PPARD, TET1, GFI1B, SOX9, SRY, YAP1, WT1, DMRT1, SMAD3, NR3C1, RUNX2, CUX1, YY1, TCF7, EZH2, RNF2, JARID2, MTF2, SUZ12, CTNNB1, TBX3, PBX1, RELA, ELF5, EWSR1, SOX11, CEBPB, SREBF2, NR1I2, STAT6, ARID3A, ATF2, BACH1, BATF, BCL11A, BCLAF1, BHLHE40, BRCA1, CEBPD, CHD1, CHD2, CHD7, CTBP2, CTCF, E2F4, E2F6, EBF1, ELF1, ELK1, EP300, ETS1, FOS, FOSL2, FOXA1, FOXA2, FOXM1, FOXP2, GABPA, GATA3, GTF2F1, H2AFZ, HCFC1, HDAC2, HDAC6, HMGN3, HNF4G, IRF3, IRF4, JUN, JUND, KDM4A, KDM5A, MAFF, MAFK, MAX, MAZ, MBD4, MTA3, MXI1, MYBL2, MYOD1, MYOG, NELFE, NFATC1, NFIC, NFYB, NR2F2, NRF1, PAX5, PBX3, PHF8, PML, POLR2A, POU2F2, RAD21, RBBP5, RCOR1, REST, RFX5, RUNX3, RXRA, SAP30, SIN3A, SMARCB1, SMARCC1, SMC3, STAT1, STAT5A, TAF1, TAF7, TBL1XR1, TBP, TCF12, TCF3, TCF7L2, TEAD4, TRIM28, UBTF, USF1, USF2, WRNIP1, ZBTB7A, ZEB1, ZKSCAN1, ZNF143, ZNF263, ZNF384 |
| *RAB19* | FOXP3, STAT3, TCF4, E2F1, RUNX1, SPI1, NANOG, POU5F1, EGR1, PPARG, SETDB1, ASH2L, TFAP2C, FOXP1, PPARD, GFI1B, TAL1, TEAD4, SCLY, HOXB4, STAT4, LMO2, ARID3A, BHLHE40, CEBPB, CHD1, CHD2, CHD7, CTCF, E2F4, E2F6, ELF1, ELK1, EP300, EZH2, FOS, GATA1, GATA3, GTF2F1, H2AFZ, HCFC1, HDAC1, HDAC2, IRF1, JUND, KDM1A, KDM4A, KDM5A, KDM5B, MAFK, MAX, MAZ, MXI1, MYC, NCOR1, NR2F2, PHF8, POLR2A, RAD21, RBBP5, RCOR1, RFX5, SAP30, SIN3A, SIX5, SMC3, SUZ12, TAF1, TBL1XR1, TBP, TCF12, THAP1, USF1, USF2, YY1, ZBTB7A, ZEB1, ZKSCAN1, ZMIZ1, ZNF143, ZNF384 |

| | |
|---|---|
| *TMEM255A* | AR, TP63, TP53, HNF4A, GATA1, GATA2, REST, MYCN, TEAD4, SUZ12, TFAP2A, ARID3A, ATF2, ATF3, BACH1, BATF, BCL11A, BCL3, BCLAF1, BHLHE40, BRCA1, CBX3, CEBPB, CHD1, CHD2, CREB1, CTBP2, CTCF, E2F4, E2F6, EBF1, ELK1, EP300, EZH2, FOS, FOSL2, FOXA2, FOXM1, FOXP2, GABPA, GATA3, GTF2F1, H2AFZ, HDAC2, IRF4, JUND, KDM4A, KDM5A, KDM5B, MAX, MAZ, MTA3, MXI1, MYBL2, MYC, NFATC1, NFIC, NRF1, PAX5, PBX3, PHF8, PML, POLR2A, POU2F2, RAD21, RBBP5, RCOR1, RELA, RFX5, RUNX3, SAP30, SIN3A, SMC3, SP1, SPI1, STAT1, STAT3, STAT5A, TAF1, TBL1XR1, TBP, TCF12, TCF3, TCF7L2, TRIM28, USF1, USF2, WRNIP1, YY1, ZBTB33, ZEB1, ZNF143, ZNF263, ZNF384, ZZZ3 |
| *HPDL* | TCF4, SMAD4, RUNX1, MYC, SPI1, EGR1, MITF, TP53, E2F4, RCOR3, EOMES, TET1, ERG, CNOT3, TRIM28, TFCP2L1, ESRRB, NR0B1, PDX1, ESR2, RARG, ATF2, BACH1, BATF, BCL3, BCLAF1, BHLHE40, BRCA1, CCNT2, CEBPB, CHD1, CHD2, CHD7, CREB1, CTCF, E2F6, EBF1, ELF1, ELK1, EP300, EZH2, FOXA2, FOXM1, GABPA, GATA1, GTF2F1, H2AFZ, HDAC2, HDAC6, HMGN3, JUN, JUND, KDM4A, KDM5A, KDM5B, MAFK, MAX, MAZ, MTA3, MXI1, MYOG, NELFE, NFATC1, NFIC, NFYB, NRF1, PAX5, PHF8, PML, POLR2A, POU2F2, RAD21, RBBP5, RCOR1, REST, RFX5, RUNX3, SAP30, SIN3A, SMARCB1, SMC3, SP1, SP4, STAT1, STAT5A, TAF1, TAF7, TBL1XR1, TBP, TCF12, TCF3, TCF7L2, TEAD4, USF1, USF2, WRNIP1, YY1, ZBTB33, ZKSCAN1, ZMIZ1, ZNF143, ZNF384 |
| *RHPN2* | ZNF217, AR, SMAD4, CREB1, E2F1, SPI1, NANOG, POU5F1, SOX2, TCF3, TP53, HNF4A, RCOR3, SIN3B, REST, TFAP2C, FOXP1, MYCN, GFI1B, FOXO3, SOX17, WT1, ZNF281, PRDM14, KLF1, TFCP2L1, MEF2A, TBX5, FOXP2, BMI1, TBX3, SIN3A, ZIC3, CDX2, IRF8, ESR2, BACH1, BCL3, BHLHE40, BRCA1, CCNT2, CEBPB, CEBPD, CHD1, CHD2, CTCF, CUX1, E2F4, E2F6, EGR1, ELF1, ELK1, EP300, ETS1, EZH2, FOSL2, FOXA2, GABPA, GATA1, GATA2, GATA3, GTF2B, GTF2F1, H2AFZ, HCFC1, HDAC1, HDAC2, HDAC6, HMGN3, IRF1, IRF4, JUND, KDM4A, KDM5A, KDM5B, MAFK, MAX, MAZ, MEF2C, MTA3, MXI1, MYC, NFIC, NRF1, PBX3, PHF8, POLR2A, PRDM1, RAD21, RBBP5, RCOR1, RFX5, SAP30, SMARCB1, SMC3, SP1, STAT3, SUZ12, TAF1, TBL1XR1, TBP, TCF12, TCF7L2, TFAP2A, UBTF, USF1, YY1, ZBTB33, ZBTB7A, ZEB1, ZKSCAN1, ZMIZ1, ZNF143, ZNF263 |
| *KIAA0319* | NANOG, POU5F1, SOX2, TCF3, SIN3B, TBX5, BACH1, SREBF2, DROSHA, BHLHE40, BRCA1, CBX3, CCNT2, CEBPB, CHD1, CHD2, CHD7, CREB1, CTBP2, CTCF, E2F1, E2F4, E2F6, EBF1, EGR1, ELF1, ELK1, ELK4, EP300, EZH2, FOS, FOSL2, FOXA2, GATA3, GTF2F1, H2AFZ, HCFC1, HDAC1, HDAC2, HDAC6, HMGN3, JUND, KDM1A, KDM4A, KDM5A, KDM5B, MAFK, MAX, MAZ, MXI1, MYC, MYOG, NR2F2, NRF1, PAX5, PHF8, POLR2A, RAD21, RBBP5, RCOR1, RELA, RFX5, RNF2, RUNX3, SAP30, SETDB1, SIN3A, SMARCB1, SMARCC1, SMC3, SP4, SPI1, STAT1, STAT3, SUZ12, TAL1, TBL1XR1, TBP, TCF12, TCF7L2, TFAP2A, TFAP2C, TRIM28, UBTF, USF2, YY1, ZBTB7A, ZEB1, ZKSCAN1, ZNF143, ZNF217, ZNF384 |
| *NXNL2* | KLF2, KLF5, ELK1, STAT3, TP63, KLF4, TCF3, EGR1, REST, PPARD, SOX9, TCF7, SUZ12, ESRRB, TBX3, PADI4, BACH1, BCL3, BRCA1, CEBPB, CHD1, CHD2, CREB1, CTBP2, CTCF, E2F4, E2F6, ELF1, EP300, ESR1, ETS1, EZH2, FOSL2, FOXA1, FOXP2, GATA3, GTF2F1, H2AFZ, HDAC1, HDAC2, HDAC6, HMGN3, IRF1, JUND, KDM4A, KDM5B, MAFK, MAX, MAZ, MXI1, MYC, NANOG, NFIC, NR2F2, NR3C1, PBX3, PHF8, PML, POLR2A, PRDM1, RAD21, RBBP5, RCOR1, RFX5, SAP30, SIN3A, SIRT6, SIX5, SMARCB1, SMC3, SP1, SP4, SPI1, TAF1, TBL1XR1, TBP, TCF12, TEAD4, TFAP2C, TRIM28, UBTF, USF1, YY1, ZBTB7A, ZC3H11A, ZEB1, ZKSCAN1, ZNF143, ZNF217, ZNF263 |
| *APOLD1* | FOXP3, PHF8, SMAD4, FLI1, RUNX1, SPI1, MITF, TP53, FOXA2, GATA1, E2F4, SETDB1, PRDM14, KLF1, ATF3, SMAD3, MTF2, SUZ12, JUN, FOXM1, ARID3A, ATF1, ATF2, BACH1, BATF, BCL3, BCLAF1, BHLHE40, BRCA1, CBX3, CCNT2, CEBPB, CEBPD, CHD1, CHD2, CREB1, CTBP2, CTCF, CUX1, E2F1, E2F6, EBF1, EGR1, ELF1, ELK1, EP300, ETS1, EZH2, FOS, FOSL1, FOSL2, FOXA1, FOXP2, GABPA, GATA2, GATA3, GTF2B, GTF2F1, H2AFZ, HCFC1, HDAC1, HDAC2, HDAC6, HMGN3, HNF4A, HNF4G, IRF1, IRF3, JUND, KAT2A, KDM1A, KDM4A, KDM5B, MAFF, MAFK, MAX, MAZ, MBD4, MEF2A, MEF2C, MTA3, MXI1, MYBL2, MYC, MYOG, NANOG, NELFE, NFATC1, NFIC, NFYA, NFYB, NR2F2, NR3C1, NRF1, PAX5, PBX3, PML, POLR2A, POU2F2, RAD21, RBBP5, RCOR1, RELA, REST, RFX5, RNF2, RUNX3, RXRA, SAP30, SIN3A, SIX5, SMARCB1, SMC3, SP1, SREBF2, SRF, STAT1, STAT3, STAT5A, SUPT20H, TAF1, TAF7, TAL1, TBL1XR1, TBP, TCF12, TCF3, TCF7L2, TEAD4, TRIM28, UBTF, USF1, USF2, WRNIP1, YY1, ZBTB33, ZBTB7A, ZEB1, ZKSCAN1, ZMIZ1, ZNF143, ZNF263, ZNF384 |
| *MAPK8IP1* | DACH1, TP63, CREB1, CREM, RUNX1, MYC, SOX2, PPARG, RCOR3, SIN3B, REST, FOXP1, PPARD, ERG, SOX9, SRY, EP300, SALL4, MTF2, SIN3A, POU3F2, RCOR1, BACH1, RBPJ, ARID3A, BHLHE40, CBX3, CCNT2, CHD1, CHD2, CHD7, CTCF, CUX1, E2F4, EBF1, EGR1, ELF1, ELK1, ETS1, EZH2, GTF2F1, H2AFZ, HCFC1, HDAC1, HDAC2, HMGN3, IRF1, JUN, JUND, KDM4A, KDM5A, KDM5B, MAX, MAZ, MXI1, NELFE, PAX5, PHF8, POLR2A, RAD21, RBBP5, RUNX3, SAP30, SMARCB1, SMC3, SP4, STAT1, SUZ12, TAF1, TAF7, TAL1, TBP, TCF12, TEAD4, THAP1, USF1, USF2, YY1, ZBTB7A, ZC3H11A, ZKSCAN1, ZMIZ1, ZNF143, ZNF384 |
| *MARC2* | TTF2, ELK1, PHF8, TP63, E2F1, KLF4, POU5F1, PPARG, HNF4A, ZFX, EOMES, CNOT3, TRIM28, SOX9, MECOM, SOX11, NACC1, PDX1, CLOCK, ARID3A, ATF2, ATF3, BCL3, BHLHE40, BRCA1, CEBPB, CEBPD, CHD1, CHD2, CHD7, CREB1, CTBP2, CTCF, E2F4, E2F6, ELF1, EP300, ETS1, EZH2, FOSL2, FOXA1, FOXA2, GABPA, GATA1, GATA3, GTF2F1, H2AFZ, HCFC1, HDAC2, HDAC6, HNF4G, JUND, KAT2A, KDM4A, KDM5A, KDM5B, MAFF, MAFK, MAX, MAZ, MXI1, MYBL2, MYC, MYOD1, MYOG, NELFE, NFIC, NFYA, NRF1, PAX5, PML, POLR2A, RAD21, RBBP5, RCOR1, REST, RFX5, SAP30, SIN3A, SIRT6, SMARCB1, SMARCC1, SMC3, SP1, STAT1, SUZ12, TAF1, TAF7, TBP, TCF12, TCF7L2, TEAD4, UBTF, USF1, USF2, WRNIP1, YY1, ZBTB7A, ZKSCAN1, ZMIZ1, ZNF143, ZNF263 |

| | |
|---|---|
| *HOXB13* | KLF2, KLF5, STAT3, SMAD4, CREB1, KLF4, NANOG, POU5F1, SOX2, TCF3, TP53, HNF4A, KDM5B, SIN3B, SETDB1, ASH2L, TET1, SRY, SMARCA4, SALL4, ZNF281, DMRT1, PRDM14, YY1, RAD21, BMI1, EED, PHC1, EZH2, RNF2, MTF2, SUZ12, TBX3, SOX11, NR0B1, ZIC3, SMAD1, ELF1, CBX3, CEBPB, CHD1, CHD7, CTCF, EBF1, EGR1, EP300, FOSL2, FOXP2, GATA1, GTF2F1, H2AFZ, HCFC1, HDAC1, HDAC2, HDAC6, JUND, KDM4A, KDM5A, MAFK, MAX, MAZ, MXI1, MYC, NELFE, NFYA, NFYB, POLR2A, RBBP5, REST, RFX5, SIN3A, SP1, TBP, TCF12, TRIM28, UBTF, ZBTB7A, ZMIZ1, ZNF143, ZNF263, ZZZ3 |
| *DUSP9* | AR, E2F1, RUNX1, MYC, SOX2, EGR1, MITF, HNF4A, SRY, SALL4, MTF2, SUZ12, TFAP2A, PRDM5, ARID3A, BACH1, BHLHE40, CBX2, CBX3, CCNT2, CEBPD, CHD1, CHD2, CREB1, CTCF, CTCFL, E2F4, E2F6, EBF1, ELF1, EP300, EZH2, FOXP2, H2AFZ, HCFC1, HDAC1, HDAC2, HDAC6, HMGN3, JUND, KDM4A, KDM5A, KDM5B, MAFF, MAFK, MAX, MAZ, MXI1, MYOG, NELFE, PHF8, POLR2A, RAD21, RBBP5, RCOR1, RFX5, RXRA, SAP30, SMARCB1, SMC3, SP1, STAT3, TAF1, TBP, UBTF, USF1, YY1, ZBTB7A, ZNF143, ZNF263 |
| *ACOT4* | E2F1, POU5F1, EGR1, GATA2, E2F4, SETDB1, GFI1B, TAL1, CNOT3, TRIM28, FOXO3, ESR1, HOXB4, LMO2, MEIS1, ELF5, CEBPA, CEBPB, ESR2, BACH1, BHLHE40, BRCA1, CHD1, CHD2, CTCF, CUX1, E2F6, EBF1, ELF1, ELK1, EP300, EZH2, GABPA, GATA1, H2AFZ, HDAC2, JUND, KDM4A, KDM5B, MAX, MAZ, MTA3, MXI1, MYC, NANOG, NFIC, NFYB, NR2F2, NRF1, PAX5, PHF8, PML, POLR2A, RAD21, RBBP5, RCOR1, RUNX3, SIN3A, SMC3, SP1, SPI1, STAT1, STAT3, SUZ12, TAF1, TBP, TCF12, TCF3, TEAD4, WRNIP1, YY1, ZBTB7A, ZEB1, ZNF143, ZNF384 |
| *CAGE1* | PHF8, TP63, FLI1, RUNX1, SPI1, PPARG, GATA2, PPARD, TET1, TAL1, SOX9, CUX1, MEF2A, RAD21, LMO2, MEIS1, TBP, DCP1A, ELF1, ARID3A, ATF2, ATF3, BACH1, BCL3, BCLAF1, BHLHE40, BRCA1, CBX3, CCNT2, CEBPB, CEBPD, CHD1, CHD2, CHD7, CREB1, CREBBP, CTCF, E2F4, E2F6, EBF1, EGR1, ELK1, ELK4, EP300, ETS1, EZH2, FOS, FOSL2, FOXA1, FOXM1, FOXP2, GABPA, GATA3, GTF2B, GTF2F1, H2AFZ, HCFC1, HDAC1, HDAC2, HDAC6, HMGN3, IRF1, IRF3, IRF4, JUN, JUND, KAT2A, KAT2B, KDM1A, KDM4A, KDM5A, KDM5B, MAFF, MAFK, MAX, MAZ, MBD4, MTA3, MXI1, MYB, MYBL2, MYC, MYOG, NELFE, NFATC1, NFIC, NFYB, NR2F2, NRF1, PAX5, PBX3, PML, POLR2A, POU2F2, PRDM1, RBBP5, RCOR1, RELA, REST, RFX5, RUNX3, SAP30, SETDB1, SIN3A, SIRT6, SMARCB1, SMC3, SP1, SP2, SP4, SRF, STAT1, STAT3, STAT5A, SUZ12, TAF1, TAF7, TBL1XR1, TCF12, TCF3, TCF7L2, TEAD4, TRIM28, UBTF, USF1, USF2, WHSC1, WRNIP1, YY1, ZBTB7A, ZC3H11A, ZEB1, ZKSCAN1, ZMIZ1, ZNF143, ZNF263, ZNF384 |
| *ABHD11* | XRN2, AR, CREM, E2F1, MYC, SPI1, KLF4, NANOG, POU5F1, SOX2, TCF3, PPARG, SIN3B, TFAP2C, FOXP1, TET1, ERG, TAL1, ESR1, DMRT1, PRDM14, KLF1, SMAD2, SMAD3, NR3C1, SCLY, CCND1, HOXB4, STAT4, SRF, TBX5, ESRRB, MYB, STAT5A, NR0B1, CDX2, CEBPA, PDX1, RBPJ, SREBF1, ARID3A, ATF1, ATF2, ATF3, BCL3, BHLHE40, BRCA1, CBX3, CCNT2, CEBPB, CEBPD, CHD1, CHD2, CHD7, CREB1, CTCF, CUX1, E2F4, E2F6, EBF1, EGR1, ELF1, ELK1, ELK4, EP300, ETS1, EZH2, FOS, FOXA1, FOXM1, FOXP2, GABPA, GATA3, GTF2B, GTF2F1, H2AFZ, HCFC1, HDAC1, HDAC2, HDAC6, HMGN3, IRF1, JUN, JUND, KDM4A, KDM5B, MAFK, MAX, MAZ, MTA3, MXI1, MYBL2, NELFE, NFIC, NFYA, NFYB, NR2F2, NRF1, PAX5, PHF8, PML, POLR2A, POU2F2, RAD21, RBBP5, RCOR1, REST, RFX5, RUNX3, SAP30, SETDB1, SIN3A, SIRT6, SMARCB1, SMC3, SP1, SP4, STAT1, STAT3, SUZ12, TAF1, TAF7, TBL1XR1, TBP, TCF12, TCF7L2, TEAD4, TFAP2A, THAP1, TRIM28, UBTF, USF1, USF2, WHSC1, YY1, ZBTB33, ZBTB7A, ZC3H11A, ZEB1, ZKSCAN1, ZMIZ1, ZNF143, ZNF263, ZNF384 |
| *NKX2-5* | SMAD1, SMAD4, STAT3, MYC, KLF4, POU5F1, EGR1, TP53, KDM5B, REST, GATA4, MEF2A, TBX5, BMI1, EED, PHC1, EZH2, RNF2, JARID2, MTF2, SUZ12, LMO2, NOTCH1, RCOR2, BACH1, CHD1, CREB1, CTBP2, CTCF, E2F4, E2F6, EBF1, ELK4, EP300, FOS, GABPA, H2AFZ, HDAC1, HDAC2, HDAC6, JUND, KDM4A, KDM5A, MAX, NFIC, POLR2A, RAD21, RBBP5, SIN3A, SIRT6, SP1, SRF, TAF1, TBP, TCF12, TRIM28, YY1, ZBTB7A, ZNF143, ZNF263, ZNF384 |
| *NPTXR* | CREM, SPI1, KLF4, TCF3, MITF, HNF4A, RCOR3, SIN3B, REST, ASH2L, CNOT3, TRIM28, SALL4, NR3C1, TCF7, JARID2, MTF2, SUZ12, BACH1, TCF7L2, STAT1, NFIB, BCLAF1, BHLHE40, CBX3, CCNT2, CEBPB, CEBPD, CHD1, CHD2, CHD4, CREB1, CTCF, CTCFL, E2F1, E2F4, E2F6, EBF1, ELF1, EP300, ETS1, EZH2, FOS, FOXA1, FOXA2, FOXP2, GABPA, H2AFZ, HCFC1, HDAC1, HDAC2, HDAC6, HMGN3, HNF4G, JUND, KAT2A, KDM1A, KDM4A, KDM5B, MAX, MAZ, MBD4, MXI1, MYBL2, MYC, MYOD1, MYOG, NELFE, NFIC, NR2F2, NRF1, PHF8, POLR2A, RAD21, RBBP5, RCOR1, RFX5, SAP30, SIN3A, SMC3, SP1, TAF1, TAL1, TBL1XR1, TBP, TCF12, UBTF, YY1, ZBTB7A, ZKSCAN1, ZMIZ1, ZNF143, ZNF263, ZNF384 |
| *HHIPL2* | AR, TCF4, SOX2, HNF4A, PPARD, CTNNB1, ATF3, BCL3, CEBPB, CHD1, CHD2, CREB1, CTCF, CUX1, ELK1, EP300, EZH2, FOS, FOSL2, FOXA1, FOXA2, FOXP2, GATA3, H2AFZ, HDAC1, HDAC2, JUND, KDM5A, KDM5B, MAFK, MAX, MAZ, MXI1, MYC, NR3C1, PBX3, POLR2A, RAD21, RBBP5, RCOR1, REST, RFX5, SETDB1, SMC3, SP1, STAT3, TBL1XR1, TBP, TCF12, TCF7L2, TFAP2A, TFAP2C, TRIM28, UBTF, USF2, YY1, ZKSCAN1, ZNF143, ZNF263 |
| *FOXB1* | ARNT, AR, STAT3, KLF4, NANOG, POU5F1, SOX2, TCF3, TP53, KDM5B, SIN3B, REST, FOXP1, TRIM28, EP300, ESR1, OLIG2, SMARCA4, ZNF281, MYBL2, BMI1, EZH2, RNF2, JARID2, MTF2, SUZ12, BACH1, BRCA1, CBX2, CBX8, CEBPB, CHD1, CHD2, CHD7, CREB1, CTBP2, CTCF, E2F6, EBF1, EGR1, FOXA2, GTF2F1, H2AFZ, HDAC2, HMGN3, JUND, KDM4A, KDM5A, MAX, MAZ, MXI1, MYC, PHF8, POLR2A, RAD21, RFX5, SAP30, SIN3A, TAF1, TBP, TCF12, TCF7L2, TEAD4, USF2, YY1, ZNF143, ZNF263 |
| *GABRA3* | AR, SMAD4, E2F1, ZNF281, SCLY, CBX3, CHD1, CHD2, CTCF, EZH2, GATA3, GTF2F1, H2AFZ, HDAC1, HDAC2, HDAC6, JUND, KDM4A, KDM5A, KDM5B, MAFK, MAX, MAZ, MYC, NFYB, POLR2A, RBBP5, RCOR1, REST, RFX5, SAP30, SIN3A, TAF1, TBL1XR1, TBP, TCF7L2, WHSC1, ZNF143 |
| *CD38* | AR, RARA, VDR, STAT3, TP63, E2F1, RUNX1, KLF4, NANOG, POU5F1, SOX2, TCF3, TP53, GATA1, ASH2L, TAL1, TRIM28, SALL4, ZNF281, RUNX2, ESRRB, PBX1, RELA, NACC1, NR0B1, ZIC3, SMAD1, ARID3A, ATF2, BATF, BCL11A, BCLAF1, BHLHE40, CBX2, CCNT2, CEBPB, CHD1, CHD2, CHD7, CTBP2, CTCF, CUX1, E2F4, EBF1, EGR1, ELF1, ELK1, EP300, EZH2, FOS, FOXA1, FOXA2, FOXM1, FOXP2, H2AFZ, HDAC2, IRF4, KDM4A, MAFK, MAX, MAZ, MEF2A, MTA3, MXI1, MYC, NELFE, NFATC1, NFE2, NFIC, NFYB, NR3C1, PAX5, PHF8, PML, POLR2A, POU2F2, RAD21, RBBP5, RCOR1, REST, RFX5, |

| | |
|---|---|
| | RNF2, RUNX3, SAP30, SIN3A, SP1, SPI1, STAT1, STAT5A, SUZ12, TAF1, TBL1XR1, TBP, TCF12, USF2, WRNIP1, YY1, ZBTB7A, ZEB1, ZNF143, ZNF384 |
| *CCRL2* | MYC, FOXP3, TCF4, SMAD4, TP63, SPI1, SOX2, GATA1, GATA2, TAL1, SOX9, KLF1, CUX1, MECOM, GATA4, SRF, TBX5, FOXP2, TCF7, SUZ12, LMO2, IRF8, ARID3A, ATF1, ATF3, BACH1, BATF, BHLHE40, CBX3, CCNT2, CEBPB, CEBPD, CHD1, CHD2, CHD7, CREB1, CTCF, E2F6, EGR1, ELF1, ELK1, ELK4, EP300, ETS1, EZH2, FOS, FOSL1, FOXA1, GABPA, GTF2B, GTF2F1, H2AFZ, HCFC1, HDAC1, HDAC2, HDAC6, HMGN3, IRF1, JUN, JUND, KDM1A, KDM5A, KDM5B, MAFF, MAFK, MAX, MAZ, MTA3, MXI1, NCOR1, NELFE, NR2F2, PHF8, PML, POLR2A, RAD21, RBBP5, RCOR1, REST, RFX5, RNF2, RUNX3, SAP30, SETDB1, SIN3A, SIRT6, SMC3, STAT5A, TAF1, TBL1XR1, TBP, TCF7L2, TEAD4, TRIM28, UBTF, USF1, USF2, WHSC1, YY1, ZBTB7A, ZC3H11A, ZMIZ1, ZNF143, ZNF384 |
| *PLCL1* | PAX3, STAT3, TCF4, FLI1, SPI1, SOX2, PPARD, TET1, GFI1B, SOX9, ESR1, OLIG2, SMARCA4, DMRT1, SCLY, TBX5, TCF7, RAD21, JARID2, MTF2, SUZ12, POU3F2, BACH1, STAT5A, ARID3A, BCLAF1, BHLHE40, BRCA1, CCNT2, CEBPB, CHD1, CHD2, CHD7, CREB1, CTBP2, CTCF, CTCFL, CUX1, E2F4, E2F6, EBF1, ELF1, ELK1, EP300, ETS1, EZH2, FOS, GABPA, GATA2, GTF2F1, H2AFZ, HCFC1, HDAC1, HDAC2, HDAC6, HMGN3, IRF1, IRF3, JUND, KAT2A, KDM1A, KDM4A, KDM5A, KDM5B, MAX, MAZ, MXI1, MYC, MYOD1, MYOG, NANOG, NELFE, NFIC, NFYA, NFYB, NR2F2, NRF1, PAX5, PBX3, PHF8, POLR2A, RBBP5, RCOR1, REST, RFX5, SAP30, SIN3A, SMC3, SP1, SP2, STAT1, TAF1, TBL1XR1, TBP, TCF7L2, TEAD4, UBTF, USF2, WRNIP1, YY1, ZBTB7A, ZC3H11A, ZMIZ1, ZNF143 |
| *CYTH4* | VDR, STAT3, TP63, FLI1, RUNX1, SPI1, SOX2, MITF, PPARG, RCOR3, PPARD, ERG, TAL1, TFAP2A, RELA, RBPJ, ATF1, BHLHE40, BRCA1, CBX3, CHD1, CHD2, CHD7, CTCF, CUX1, EBF1, ELF1, ELK1, EP300, ETS1, EZH2, FOS, FOXM1, GATA1, H2AFZ, HDAC1, HDAC2, HMGN3, JUND, KAT2A, KDM5A, KDM5B, MAFF, MAFK, MAX, MAZ, MTA3, MXI1, NELFE, NRF1, PAX5, PHF8, PML, POLR2A, RAD21, RBBP5, RCOR1, RFX5, RUNX3, SAP30, SIN3A, SMC3, STAT1, STAT5A, TAF1, TBL1XR1, TBP, TCF12, TCF3, USF2, WRNIP1, YY1, ZBTB7A, ZEB1, ZMIZ1, ZNF143, ZNF384 |
| *PTPRH* | STAT3, FLI1, RUNX1, SPI1, SOX2, HNF4A, GATA2, SIN3B, FOXP1, MEIS1, BACH1, ARID3A, ATF3, BCL3, BHLHE40, BRCA1, CEBPB, CEBPD, CHD1, CHD2, CTCF, CUX1, E2F6, ELF1, ELK1, EP300, ETS1, EZH2, FOS, FOSL1, FOSL2, FOXA1, FOXA2, GABPA, GATA1, GATA3, GTF3C2, H2AFZ, HDAC1, HDAC2, JUN, JUND, KDM1A, KDM5B, MAFF, MAFK, MAX, MAZ, MBD4, MEF2A, MXI1, MYBL2, MYC, NFIC, NR2F2, PBX3, POLR2A, POU2F2, RBBP5, RCOR1, REST, RFX5, RXRA, SAP30, SIN3A, SIX5, SMARCC1, SMC3, SP1, TBP, TCF12, TCF7L2, TEAD4, USF1, USF2, WHSC1, YY1, ZBTB33, ZBTB7A, ZEB1, ZKSCAN1, ZNF217, ZNF263, ZNF384 |
| *CASP2* | KDM5A, AR, TCF4, TP63, CREB1, CREM, E2F1, FLI1, RUNX1, MYC, SPI1, POU5F1, EGR1, PPARG, GATA1, KDM5B, ZFX, TFAP2C, PPARD, WT1, CUX1, MECOM, CTCF, RAD21, PRDM5, ARID3A, ATF3, BACH1, BCLAF1, BHLHE40, BRCA1, CBX3, CCNT2, CEBPB, CEBPD, CHD1, CHD2, CHD4, CHD7, CTCFL, E2F4, E2F6, EBF1, ELF1, ELK1, ELK4, EP300, ETS1, EZH2, FOS, FOXA1, FOXM1, FOXP2, GABPA, GATA3, GTF2B, GTF2F1, H2AFZ, HCFC1, HDAC1, HDAC2, HDAC6, HMGN3, IRF1, IRF3, JUN, JUND, KAT2A, KAT2B, KDM1A, KDM4A, MAFF, MAFK, MAX, MAZ, MTA3, MXI1, MYBL2, MYOD1, MYOG, NANOG, NELFE, NFIC, NFYA, NFYB, NR2F2, NRF1, PAX5, PBX3, PHF8, PML, POLR2A, POU2F2, RBBP5, RCOR1, RELA, REST, RFX5, RNF2, RUNX3, SAP30, SETDB1, SIN3A, SMARCB1, SMC3, SP1, SP2, SP4, STAT1, STAT3, STAT5A, TAF1, TAF7, TAL1, TBL1XR1, TBP, TCF12, TCF3, TCF7L2, TEAD4, THAP1, TRIM28, UBTF, USF2, WRNIP1, YY1, ZBTB7A, ZC3H11A, ZKSCAN1, ZMIZ1, ZNF143, ZNF263, ZNF384 |
| *RAB39A* | CREM, E2F1, FLI1, SPI1, NANOG, EGR1, SIN3B, REST, ASH2L, FOXP1, TET1, TEAD4, TFCP2L1, SIN3A, ARID3A, ATF2, ATF3, BACH1, BHLHE40, BRCA1, CBX3, CCNT2, CEBPB, CHD1, CHD2, CHD7, CREB1, CTBP2, CTCF, CUX1, E2F4, E2F6, EBF1, ELF1, EP300, ETS1, EZH2, FOS, FOXA1, FOXP2, GABPA, GATA2, GATA3, GTF2F1, H2AFZ, HCFC1, HDAC1, HDAC2, HMGN3, IRF1, IRF3, JUN, JUND, KAT2A, KDM1A, KDM4A, KDM5A, KDM5B, MAFF, MAFK, MAX, MAZ, MTA3, MXI1, MYC, MYOG, NELFE, NFYA, NFYB, NR2F2, NRF1, PAX5, PBX3, PHF8, PML, POLR2A, RAD21, RBBP5, RCOR1, RELA, RFX5, RUNX3, SAP30, SMC3, SP1, SP2, SP4, STAT1, STAT3, STAT5A, SUZ12, TAF1, TAF7, TBL1XR1, TBP, TCF12, TCF3, TCF7L2, TRIM28, UBTF, USF1, USF2, WRNIP1, YY1, ZBTB7A, ZC3H11A, ZMIZ1, ZNF143, ZNF263, ZNF384 |
| *ALG1L* | ATF3, ARID3A, BATF, BHLHE40, CEBPB, CEBPD, CHD2, CTCF, EP300, EZH2, FOS, FOSL2, FOXA1, FOXA2, GTF2F1, H2AFZ, IRF1, JUND, MAX, MAZ, MXI1, MYBL2, MYC, NFIC, NFYB, POLR2A, RAD21, RCOR1, RFX5, RXRA, SETDB1, SMC3, SP1, STAT3, TBP, TCF7L2, TRIM28, USF2, YY1, ZNF143 |
| *HOXA10* | ESR1, AR, STAT3, KLF4, NANOG, POU5F1, SOX2, TCF3, TP53, KDM5B, EOMES, TET1, TRIM28, SRY, EP300, WT1, SMAD2, SMAD3, BMI1, EED, PHC1, EZH2, RNF2, JARID2, MTF2, SUZ12, BACH1, CDX2, SREBF2, GATA3, RARG, CHD7, ARID3A, BHLHE40, BRCA1, CEBPB, CHD1, CHD2, CREB1, CTBP2, CTCF, CTCFL, CUX1, E2F4, ELK1, ETS1, GABPA, GTF2F1, H2AFZ, HCFC1, HDAC1, HDAC2, HMGN3, JUN, JUND, KAT2A, MAFK, MAX, MAZ, MXI1, MYC, NELFE, PBX3, POLR2A, POU2F2, RAD21, RBBP5, RCOR1, REST, RFX5, SAP30, SIN3A, SMARCB1, SMC3, SP1, TBL1XR1, TBP, TCF12, TCF7L2, TEAD4, UBTF, WRNIP1, YY1, ZC3H11A, ZNF143, ZNF263, ZNF384 |
| *MPPED1* | AR, SMAD4, TP63, CREB1, MYC, NANOG, SOX2, EGR1, MITF, TP53, HNF4A, GATA1, RCOR3, REST, SOX9, SRY, ESR1, CUX1, BMI1, JARID2, MTF2, SUZ12, CHD1, CTCF, EP300, EZH2, GATA2, H2AFZ, HDAC2, HDAC6, KDM4A, PHF8, POLR2A, RAD21, RBBP5, SAP30, SIN3A, SMC3 |
| *HSD17B3* | AR, STAT3, TP63, SPI1, NANOG, POU5F1, SOX2, TP53, OLIG2, SMARCA4, DMRT1, PRDM14, FOXP2, TBX3, BACH1, EP300, EZH2, FOS, GATA3, H2AFZ, MAX, MXI1, PAX5, PBX3, POLR2A, RBBP5, RUNX3, TCF12, TCF7L2, YY1 |

| | |
|---|---|
| *TMPRSS2* | AR, PPARG, HNF4A, TET1, ERG, ESR1, YAP1, SOX17, DMRT1, RUNX2, EED, MTF2, SUZ12, CTNNB1, CDX2, RBPJ, PAX6, BACH1, BRCA1, CHD1, CHD2, CHD7, CTBP2, CTCF, E2F6, EGR1, EP300, EZH2, GABPA, GATA3, H2AFZ, HDAC2, HDAC6, JUND, KDM4A, KDM5A, KDM5B, MAFK, MAX, MYC, PHF8, POLR2A, RBBP5, SAP30, SIN3A, TAF1, TBP, TCF12, ZBTB7A, ZNF143, ZNF263 |
| *RGS6* | ARNT, ZNF217, AR, STAT3, TCF4, SMAD4, FLI1, RUNX1, NANOG, SOX2, TP53, GATA1, E2F4, RCOR3, TFAP2C, FOXP1, TAL1, SRY, EP300, ESR1, YAP1, DMRT1, TEAD4, RUNX2, SCLY, GATA4, SRF, EZH2, RNF2, JARID2, MTF2, SUZ12, POU3F2, NR1I2, BATF, BCL3, BCLAF1, BHLHE40, BRCA1, CBX3, CEBPB, CHD1, CHD2, CHD7, CTBP2, CTCF, E2F6, EBF1, ELF1, FOSL1, FOXA1, FOXM1, GABPA, GATA3, GTF2F1, H2AFZ, HDAC2, HMGN3, IKZF1, JUN, JUND, KDM4A, MAFK, MAX, MAZ, MTA3, MXI1, MYC, NFATC1, NFIC, NRF1, PAX5, PHF8, PML, POLR2A, RAD21, RBBP5, RCOR1, RUNX3, SAP30, SETDB1, SIN3A, SMC3, STAT5A, TAF1, TAF7, TBP, TCF12, TCF3, USF1, USF2, WRNIP1, YY1, ZEB1, ZNF143, ZNF384 |
| *RARRES3* | AR, RARA, RARB, RARG, SOX2, TP53, ERG, SCLY, IRF1, ARID3A, ATF2, BATF, BCLAF1, BHLHE40, CEBPB, CHD1, CHD2, CHD7, CTCF, CUX1, EBF1, ELK1, EP300, EZH2, FOS, FOXM1, GATA2, H2AFZ, HDAC1, HDAC2, HDAC6, IRF3, IRF4, JUND, KAT2A, KDM1A, KDM5B, MAFF, MAFK, MAX, MAZ, MXI1, MYC, NFIC, PAX5, PHF8, POLR2A, PRDM1, RAD21, RBBP5, RCOR1, RELA, REST, RFX5, RUNX3, SAP30, SMC3, SP1, SRF, STAT1, STAT3, STAT5A, TAL1, TBL1XR1, TBP, TCF12, TEAD4, UBTF, USF1, WHSC1, WRNIP1, YY1, ZMIZ1, ZNF143 |
| *OXTR* | E2F1, JUN, SP1, STAT3, TFAP2A, POU5F1, SOX2, GATA1, GATA2, SETDB1, FOXP1, PPARD, SOX9, ESR1, YAP1, SCLY, EZH2, RNF2, JARID2, MTF2, SUZ12, LMO2, RELA, EWSR1, PAX6, ATF2, ATF3, BACH1, BATF, BCL11A, BCL3, BCLAF1, BHLHE40, BRCA1, CBX2, CBX3, CCNT2, CEBPB, CHD1, CHD2, CHD7, CREB1, CTCF, CTCFL, CUX1, E2F4, E2F6, EBF1, ELF1, ELK1, EP300, FOSL2, FOXA1, FOXA2, FOXM1, FOXP2, GABPA, GATA3, GTF2F1, H2AFZ, HCFC1, HDAC1, HDAC2, HDAC6, HMGN3, IKZF1, JUND, KDM1A, KDM4A, KDM5A, KDM5B, MAFK, MAX, MAZ, MEF2A, MEF2C, MTA3, MXI1, MYBL2, MYC, MYOG, NANOG, NFIC, NR3C1, NRF1, PAX5, POLR2A, POU2F2, RAD21, RBBP5, RCOR1, REST, RFX5, RUNX3, RXRA, SAP30, SIN3A, SMC3, SP4, SRF, STAT1, STAT5A, TBL1XR1, TBP, TCF12, TCF3, TEAD4, THAP1, USF2, WHSC1, WRNIP1, YY1, ZBTB7A, ZC3H11A, ZEB1, ZMIZ1, ZNF143, ZNF384 |
| *DMBX1* | STAT3, TCF4, SMAD4, TP63, FLI1, SPI1, KLF4, SOX2, EGR1, TP53, GATA2, SETDB1, TET1, SRY, YAP1, SMAD3, CUX1, BMI1, EED, PHC1, EZH2, RNF2, JARID2, MTF2, SUZ12, ARID3A, ATF1, ATF3, BACH1, BHLHE40, CBX3, CCNT2, CEBPB, CEBPD, CHD1, CHD4, CHD7, CTCF, E2F4, ELF1, EP300, FOS, FOSL1, FOSL2, FOXA1, GATA1, HDAC2, HMGN3, IRF1, JUN, JUND, KDM1A, MAFF, MAFK, MAX, MAZ, MYC, NR2F2, POLR2A, RAD21, RBBP5, RCOR1, SAP30, SRF, STAT5A, TAL1, TBL1XR1, TEAD4, UBTF, ZBTB7A, ZMIZ1, ZNF263, ZNF384 |
| *PAX5* | POU2F1, STAT5A, STAT5B, AHR, ARNT, VDR, AR, STAT3, TCF4, TP63, CREM, MYC, SOX2, EGR1, TP53, FOXA2, HNF4A, GATA1, KDM5B, E2F4, RCOR3, SIN3B, REST, FOXP1, PPARD, SRY, ZNF281, PRDM14, MYBL2, FOXP2, CTCF, BMI1, PHC1, EZH2, RNF2, JARID2, MTF2, SUZ12, BACH1, SOX11, DNAJC2, ATF2, BCL3, BCLAF1, BHLHE40, CEBPB, CHD1, CHD2, CHD7, CREB1, CTBP2, CTCFL, E2F6, EBF1, ELF1, ELK1, EP300, ETS1, FOXM1, GTF2F1, H2AFZ, HDAC1, HDAC2, JUND, KDM4A, MAX, MAZ, MTA3, MXI1, NELFE, NFATC1, NFIC, NRF1, PAX5, PML, POLR2A, POU2F2, RAD21, RBBP5, RCOR1, RELA, RUNX3, SIN3A, SMC3, SP1, SPI1, STAT1, TAF1, TBL1XR1, TBP, TCF12, TCF3, TCF7L2, UBTF, USF1, USF2, WRNIP1, YY1, ZBTB7A, ZEB1, ZMIZ1, ZNF143, ZNF263, ZNF384 |
| *PDZK1* | ZNF217, STAT3, SMAD4, MYC, NANOG, POU5F1, SOX2, HNF4A, GATA1, EOMES, TAL1, ESR1, ATF3, SMAD3, CDX2, ESR2, ARID3A, ATF1, BRCA1, CBX2, CBX8, CEBPB, CHD2, CREB1, CTCF, E2F4, EGR1, ELF1, EP300, EZH2, FOS, FOSL2, FOXA1, FOXA2, FOXM1, GABPA, GATA2, GATA3, GTF2F1, H2AFZ, HDAC2, HNF4G, JUN, JUND, KDM1A, KDM4A, MAFK, MAX, MAZ, MXI1, MYBL2, NFIC, NR2F2, NR3C1, POLR2A, POU2F2, RAD21, RBBP5, RCOR1, REST, RFX5, RUNX3, RXRA, SIN3A, SMC3, SP1, TBP, TCF12, TCF7L2, TEAD4, TFAP2A, TFAP2C, USF1, USF2, YY1, ZEB1, ZNF143 |
| *ESAM* | FLI1, RUNX1, SPI1, NANOG, SOX2, SETDB1, FOXP1, TAL1, EP300, ESR1, OLIG2, SALL4, SOX17, MTF2, SUZ12, ARID3A, BACH1, BCL3, BCLAF1, BHLHE40, BRCA1, CBX3, CCNT2, CHD1, CHD2, CHD7, CREB1, CTBP2, CTCF, CTCFL, E2F1, E2F4, E2F6, EBF1, EGR1, ELF1, ETS1, EZH2, FOS, FOSL2, FOXA1, FOXA2, FOXM1, GATA2, GATA3, GTF2F1, H2AFZ, HCFC1, HDAC1, HDAC2, HDAC6, HMGN3, IKZF1, JUND, KDM1A, KDM4A, KDM5A, KDM5B, MAFK, MAX, MAZ, MEF2A, MTA3, MXI1, MYC, NFATC1, NFIC, NR2F2, NRF1, PAX5, PHF8, PML, POLR2A, POU2F2, RAD21, RBBP5, RCOR1, RELA, REST, RFX5, RUNX3, SAP30, SIN3A, SIRT6, SMC3, SP1, STAT1, STAT3, STAT5A, TAF1, TBL1XR1, TBP, TCF12, TCF3, TCF7L2, TEAD4, TRIM28, UBTF, USF2, WHSC1, WRNIP1, YY1, ZBTB7A, ZC3H11A, ZEB1, ZKSCAN1, ZMIZ1, ZNF143, ZNF217, ZNF263, ZNF384 |
| *PDX1* | USF1, USF2, MYC, NANOG, POU5F1, SOX2, TP53, KDM5B, SETDB1, SOX9, SRY, SALL4, BMI1, EED, PHC1, RNF2, MTF2, SUZ12, EWSR1, SOX11, PDX1, ARID3A, ATF3, CBX2, CBX8, CEBPB, CHD1, CHD2, CHD7, CREB1, CTBP2, CTCF, E2F4, E2F6, EBF1, EP300, EZH2, FOS, FOXA1, FOXA2, H2AFZ, HDAC2, HDAC6, HNF4A, JUN, KDM4A, MAX, MAZ, NFYB, POLR2A, RAD21, RBBP5, RXRA, SIN3A, SIRT6, SP1, TCF12, ZBTB7A, ZNF143, ZNF263, ZNF384 |
| *P2RX5* | TP53, MYC, SPI1, SOX2, PPARG, TBX5, MTF2, SUZ12, MEIS1, RCOR1, ZIC3, RBPJ, PAX6, NOTCH1, THRA, BACH1, BCLAF1, BHLHE40, BRCA1, CEBPB, CHD1, CHD2, CREB1, CTBP2, CTCF, E2F1, E2F4, E2F6, EBF1, EGR1, ELF1, ELK1, EP300, EZH2, GTF2F1, H2AFZ, HCFC1, HDAC1, HDAC2, HDAC6, HMGN3, JUND, KDM4A, KDM5A, MAFK, MAX, MAZ, MTA3, MXI1, MYOD1, MYOG, NFIC, NRF1, PAX5, PHF8, POLR2A, POU2F2, RAD21, RBBP5, REST, RFX5, RUNX3, SAP30, SETDB1, SIN3A, SMARCB1, SMARCC1, SMC3, STAT3, STAT5A, TAF1, TBL1XR1, TBP, TCF12, TCF3, TCF7L2, TEAD4, UBTF, USF1, USF2, WRNIP1, YY1, ZBTB7A, ZKSCAN1, ZNF143, ZNF263 |

| | |
|---|---|
| *SOAT2* | TCF4, E2F1, FLI1, SPI1, HNF4A, GATA1, GATA2, RCOR3, SIN3B, ERG, SRY, ZNF281, CUX1, MYBL2, MECOM, STAT4, EZH2, MTF2, SIN3A, LYL1, CEBPB, GATA3, ARID3A, BHLHE40, CEBPD, CHD1, CHD2, CTCF, EBF1, ELF1, EP300, FOSL2, FOXA1, FOXA2, H2AFZ, HCFC1, HDAC1, HDAC2, HDAC6, HNF4G, IKZF1, JUND, KDM1A, KDM5B, MAFK, MAX, MAZ, MBD4, MTA3, MXI1, MYC, NELFE, NFIC, NR2C2, NR2F2, PBX3, POLR2A, POU2F2, RBBP5, RCOR1, REST, RFX5, RUNX3, RXRA, SP1, SRF, SUZ12, TAF1, TAL1, TBP, TCF12, TCF3, TCF7L2, TEAD4, USF1, WRNIP1, YY1, ZBTB7A, ZMIZ1, ZNF384 |
| *ALOX5* | EGR1, MYB, SP1, AR, STAT3, TP63, FLI1, RUNX1, MYC, SPI1, SOX2, TP53, GATA1, E2F4, EOMES, OLIG2, SMARCA4, TCF7, SUZ12, RELA, GATA3, DROSHA, BACH1, BCLAF1, BHLHE40, BRCA1, CCNT2, CEBPB, CHD1, CHD2, CHD4, CTCF, CUX1, E2F6, EBF1, ELF1, ELK1, EP300, ETS1, EZH2, FOSL2, GTF2F1, H2AFZ, HCFC1, HDAC1, HDAC2, HMGN3, IRF1, IRF3, JUN, JUND, KAT2B, KDM1A, KDM4A, KDM5A, KDM5B, MAFK, MAX, MAZ, MTA3, MXI1, MYOG, NFIC, PAX5, PHF8, PML, POLR2A, RAD21, RBBP5, RCOR1, REST, RFX5, RNF2, RUNX3, SAP30, SIN3A, SMARCB1, SMARCC1, SMC3, STAT1, STAT5A, SUPT20H, TAL1, TBL1XR1, TBP, TCF3, TEAD4, TRIM28, UBTF, USF2, WRNIP1, YY1, ZBTB7A, ZC3H11A, ZEB1, ZKSCAN1, ZMIZ1, ZNF143, ZNF263, ZNF384 |
| *NPBWR1* | SMAD4, EGR1, GATA2, ERG, SMAD3, RUNX2, RAD21, JARID2, SUZ12, BACH1, BHLHE40, BRCA1, CBX3, CEBPB, CHD1, CHD2, CHD7, CREB1, CTBP2, CTCF, CTCFL, CUX1, E2F4, E2F6, ELK1, EP300, EZH2, FOS, FOXA1, FOXP2, GABPA, GATA3, GTF2F1, H2AFZ, HDAC2, JUND, KDM4A, KDM5A, KDM5B, MAFK, MAX, MAZ, MXI1, MYC, MYOG, PAX5, PHF8, POLR2A, RBBP5, RCOR1, REST, RFX5, RUNX3, SAP30, SIN3A, SMC3, SP4, STAT1, TAF7, TBP, TCF12, TCF7L2, TEAD4, TRIM28, USF2, YY1, ZC3H11A, ZNF143, ZNF263 |
| *PLAT* | AR, ATF2, CREB1, ELK1, JUN, NFIC, RARA, RARB, TFAP2A, PAX3, VDR, STAT3, TP63, FLI1, RUNX1, SPI1, NANOG, MITF, FOXA2, GATA2, KDM5B, REST, TFAP2C, ERG, TRIM28, YAP1, SALL4, SOX17, WT1, ATF3, JARID2, PBX1, EWSR1, BACH1, CLOCK, ATF1, BHLHE40, BRCA1, CEBPB, CHD1, CHD2, CHD7, CTBP2, CTCF, E2F4, E2F6, EP300, ESR1, EZH2, FOS, FOSL2, GATA1, GTF2F1, H2AFZ, HDAC2, JUND, KDM1A, KDM4A, KDM5A, MAX, MAZ, MXI1, MYC, NR3C1, PBX3, POLR2A, RAD21, RBBP5, RCOR1, RFX5, RXRA, SETDB1, SIN3A, SMC3, SUZ12, TAF1, TCF12, TCF7L2, TEAD4, UBTF, USF1, USF2, YY1, ZKSCAN1, ZNF263 |
| *FAM20C* | ZNF217, DACH1, AR, FLI1, RUNX1, NANOG, SOX2, EGR1, MITF, TP53, PPARG, HNF4A, TET1, SRY, RUNX2, SUZ12, MYB, TCF7L2, NR1H3, ARID3A, BACH1, BHLHE40, BRCA1, CEBPB, CEBPD, CHD1, CHD2, CREB1, CTCF, E2F1, E2F4, E2F6, ELF1, EP300, ESR1, EZH2, FOS, FOXA1, FOXA2, FOXP2, GABPA, GATA2, GATA3, H2AFZ, HDAC1, HDAC2, HDAC6, HNF4G, JUND, KDM4A, KDM5A, KDM5B, MAX, MAZ, MXI1, MYBL2, MYC, MYOD1, MYOG, NFIC, NFYA, NFYB, PHF8, PML, POLR2A, RAD21, RCOR1, REST, RFX5, SAP30, SIN3A, SMC3, SP1, TAF1, TBP, TCF12, TCF3, TEAD4, TRIM28, UBTF, USF1, USF2, YY1, ZBTB7A, ZNF263 |
| *TBX4* | TTF2, TCF4, TP63, SPI1, KLF4, NANOG, POU5F1, SOX2, TCF3, EGR1, MITF, TP53, FOXA2, HNF4A, GATA1, TET1, ERG, TRIM28, SRY, EP300, SALL4, BMI1, EED, PHC1, EZH2, RNF2, JARID2, MTF2, SUZ12, POU3F2, RCOR1, CRX, ARID3A, BHLHE40, BRCA1, CBX3, CEBPB, CEBPD, CHD1, CHD2, CTBP2, CTCF, E2F6, ELF1, FOXA1, GATA3, H2AFZ, HDAC2, HNF4G, JUND, KDM4A, KDM5A, MAX, MAZ, MYBL2, MYC, NFIC, POLR2A, RAD21, RBBP5, RFX5, RXRA, SAP30, SP1, TBP, TCF12, TEAD4, YY1, ZBTB7A |
| *TAPBPL* | TCF4, E2F1, FLI1, SPI1, ASH2L, CNOT3, SRY, CUX1, IRF1, IRF8, ATF2, BACH1, BATF, BCL11A, BCL3, BCLAF1, BHLHE40, BRCA1, CEBPB, CEBPD, CHD1, CHD2, CHD7, CTBP2, CTCF, E2F4, E2F6, EBF1, EGR1, ELF1, ELK1, EP300, ESRRA, ETS1, EZH2, FOXA1, FOXA2, FOXM1, GABPA, GATA1, GTF2F1, H2AFZ, HCFC1, HDAC1, HDAC2, HDAC6, IKZF1, JUN, JUND, KDM4A, KDM5A, KDM5B, MAFK, MAX, MAZ, MBD4, MTA3, MXI1, MYBL2, MYC, NELFE, NFATC1, NFIC, NFYB, NRF1, PAX5, PBX3, PHF8, PML, POLR2A, POU2F2, PRDM1, RAD21, RBBP5, RCOR1, RELA, REST, RFX5, RUNX3, SAP30, SIN3A, SMC3, SP1, SRF, STAT1, STAT3, STAT5A, TAF1, TAL1, TBL1XR1, TBP, TCF12, TCF3, TEAD4, THAP1, TRIM28, UBTF, USF1, USF2, WRNIP1, YY1, ZMIZ1, ZNF143, ZNF384 |
| *ELMO1* | PAX3, FOXP3, AR, STAT3, TCF4, SMAD4, TP63, CREM, FLI1, RUNX1, MYC, SPI1, KLF4, NANOG, POU5F1, SOX2, TCF3, EGR1, TP53, FOXA2, HNF4A, GATA1, GATA2, KDM5B, RCOR3, REST, ASH2L, GFI1B, TAL1, SOX9, ESR1, OLIG2, SMARCA4, SALL4, TEAD4, KLF1, SMAD3, CUX1, SCLY, MECOM, STAT4, CTNNB1, PBX1, LMO2, LYL1, MEIS1, POU3F2, EWSR1, BACH1, NR0B1, NR1I2, TCF7L2, IRF8, GBX2, FOXM1, ATF1, ATF2, BCL11A, BCLAF1, BHLHE40, CEBPB, CHD1, CHD2, CHD7, CREB1, CTBP2, CTCF, E2F4, E2F6, EBF1, ELF1, ELK1, EP300, ETS1, EZH2, GABPA, GTF2F1, H2AFZ, HCFC1, HDAC1, HDAC2, HDAC6, HMGN3, IRF4, JUN, JUND, KDM4A, KDM5A, MAX, MAZ, MEF2A, MTA3, MXI1, NELFE, NFIC, NRF1, PAX5, PHF8, PML, POLR2A, POU2F2, RAD21, RBBP5, RCOR1, RELA, RFX5, RUNX3, SAP30, SIN3A, SMC3, STAT1, STAT5A, SUZ12, TAF1, TAF7, TBL1XR1, TBP, TCF12, TRIM28, UBTF, USF2, YY1, ZC3H11A, ZNF143, ZNF263, ZNF384 |
| *NRGN* | ETV4, SP1, TFAP2A, TP63, E2F1, FLI1, RUNX1, MYC, SPI1, POU5F1, SOX2, EGR1, GATA2, E2F4, FOXP1, TET1, KLF1, SCLY, HOXB4, MECOM, EZH2, RNF2, MTF2, SUZ12, CTNNB1, LMO2, LYL1, BACH1, BHLHE40, BRCA1, CBX3, CCNT2, CEBPB, CHD1, CHD2, CREBBP, CTBP2, CTCF, E2F6, EBF1, ELF1, EP300, ETS1, FOS, FOXA1, FOXA2, FOXP2, GATA1, GATA3, H2AFZ, HDAC1, HDAC2, HDAC6, HMGN3, IRF1, JUND, KDM4A, KDM5B, MAX, MAZ, MXI1, NFYA, NFYB, NR2F2, NRF1, PHF8, POLR2A, RAD21, RBBP5, RCOR1, RFX5, RUNX3, SAP30, SIN3A, SIRT6, SIX5, SMC3, STAT1, STAT3, TAF1, TAL1, TBL1XR1, TBP, UBTF, USF1, USF2, WHSC1, YY1, ZBTB7A, ZNF143, ZNF263, ZNF384 |

| | |
|---|---|
| *HIST1H2AG* | ELK1, CREB1, RUNX1, MYC, NANOG, POU5F1, SOX2, TCF3, EGR1, TP53, E2F4, TAL1, TFCP2L1, ATF3, CUX1, HOXB4, MYBL2, STAT4, RNF2, TBX3, HSF1, TBP, NACC1, NR0B1, SMAD1, KDM6A, PADI4, IRF8, THAP11, FOXM1, ARID3A, ATF1, ATF2, BACH1, BATF, BCL11A, BCL3, BCLAF1, BHLHE40, BRCA1, CBX3, CCNT2, CEBPB, CEBPD, CHD1, CHD2, CHD7, CREBBP, CTCF, E2F1, E2F6, EBF1, ELF1, ELK4, EP300, ETS1, EZH2, FLI1, FOS, FOSL1, FOSL2, FOXA1, FOXA2, FOXP2, GABPA, GATA1, GATA2, GATA3, GTF2B, GTF2F1, GTF3C2, H2AFZ, HCFC1, HDAC2, HDAC6, HMGN3, IKZF1, IRF1, IRF3, IRF4, JUN, JUND, KAT2A, KAT2B, KDM4A, KDM5A, MAFF, MAFK, MAX, MAZ, MEF2A, MEF2C, MTA3, MXI1, NELFE, NFATC1, NFE2, NFIC, NFYA, NFYB, NR2C2, NR2F2, NR3C1, NRF1, PAX5, PBX3, PHF8, PML, POLR2A, POU2F2, RAD21, RBBP5, RCOR1, RELA, REST, RFX5, RUNX3, RXRA, SAP30, SETDB1, SIN3A, SIRT6, SIX5, SMARCB1, SMC3, SP1, SP2, SP4, SPI1, SRF, STAT1, STAT3, STAT5A, TAF1, TAF7, TBL1XR1, TCF12, TCF7L2, TEAD4, TRIM28, UBTF, USF1, USF2, WRNIP1, YY1, ZBTB33, ZBTB7A, ZC3H11A, ZEB1, ZKSCAN1, ZMIZ1, ZNF143, ZNF217, ZNF263, ZNF274, ZNF384 |
| *TPRN* | KDM5A, XRN2, DACH1, PHF8, CREB1, E2F1, MYC, SPI1, NANOG, SOX2, EGR1, MITF, SIN3B, REST, TFAP2C, TET1, ERG, TRIM28, ESR1, ZNF281, MYBL2, GATA4, RCOR1, NR0B1, CEBPB, PRDM5, ARID3A, ATF2, ATF3, BACH1, BCL3, BHLHE40, BRCA1, CBX3, CCNT2, CEBPD, CEBPZ, CHD1, CHD2, CHD4, CTCF, E2F4, E2F6, EBF1, ELF1, ELK1, ELK4, EP300, ESRRA, ETS1, EZH2, FOS, FOSL2, FOXA1, FOXA2, FOXP2, GABPA, GATA1, GTF2B, GTF2F1, H2AFZ, HCFC1, HDAC1, HDAC2, HDAC6, HMGN3, HNF4A, HNF4G, IRF1, IRF3, JUND, KAT2A, KDM1A, KDM4A, KDM5B, MAFF, MAFK, MAX, MAZ, MBD4, MTA3, MXI1, MYOG, NELFE, NFATC1, NFIC, NFYA, NFYB, NR2F2, NRF1, PAX5, PBX3, PML, POLR2A, POU2F2, RAD21, RBBP5, RELA, RFX5, RUNX3, SAP30, SIN3A, SIRT6, SIX5, SMARCB1, SMC3, SP1, SP2, SP4, SRF, STAT1, STAT3, STAT5A, SUZ12, TAF1, TAF7, TBL1XR1, TBP, TCF12, TCF3, TCF7L2, THAP1, UBTF, USF1, USF2, WRNIP1, YY1, ZBTB33, ZBTB7A, ZC3H11A, ZKSCAN1, ZMIZ1, ZNF143, ZNF263, ZNF384 |
| *PFKFB3* | HIF1A, NFIC, PGR, TFAP2A, ZNF217, AR, TP63, E2F1, FLI1, RUNX1, MYC, SOX2, EGR1, MITF, PPARG, E2F4, FOXP1, PPARD, TET1, CNOT3, TRIM28, SOX9, ESR1, SMARCA4, SALL4, DMRT1, ATF3, CCND1, CTCF, MTF2, SUZ12, NUCKS1, ELF5, TBP, GATA3, CLOCK, NR1H3, ESR2, ARID3A, BACH1, BHLHE40, BRCA1, CBX3, CCNT2, CEBPB, CHD1, CHD2, CHD4, CREB1, CTBP2, E2F6, EBF1, ELF1, ELK1, ELK4, EP300, ETS1, EZH2, FOXA1, FOXP2, GABPA, H2AFZ, HCFC1, HDAC1, HDAC2, HDAC6, HMGN3, IKZF1, IRF1, JUN, JUND, KAT2A, KDM4A, KDM5B, MAFK, MAX, MAZ, MTA3, MXI1, NELFE, PAX5, PHF8, PML, POLR2A, RAD21, RBBP5, RCOR1, REST, RFX5, RUNX3, SAP30, SIN3A, SMARCB1, SMC3, SP1, SRF, STAT1, STAT5A, TAF1, TAF7, TBL1XR1, TCF12, TCF3, TCF7L2, TEAD4, TFAP2C, UBTF, USF1, USF2, WRNIP1, YY1, ZBTB33, ZBTB7A, ZC3H11A, ZEB1, ZKSCAN1, ZMIZ1, ZNF143, ZNF263, ZNF384 |
| *NR4A1* | HIF1A, RELA, VDR, ELK1, PHF8, AR, CREB1, CREM, RUNX1, MYC, KLF4, TCF3, EGR1, MITF, TP53, RCOR3, SIN3B, REST, EOMES, TFAP2C, FOXP1, PPARD, MYCN, SRY, EP300, WT1, ZNF281, DMRT1, PRDM14, EZH2, TFAP2A, CTNNB1, SIN3A, NACC1, GATA3, CLOCK, THRA, ATF2, BACH1, BCL3, BHLHE40, BRCA1, CBX2, CBX3, CCNT2, CEBPB, CHD1, CHD2, CHD4, CHD7, CTCF, CTCFL, CUX1, E2F4, E2F6, EBF1, ELF1, ETS1, FOS, FOSL2, FOXP2, GABPA, GTF2F1, H2AFZ, HCFC1, HDAC1, HDAC2, HDAC6, HMGN3, IRF1, JUN, JUND, KAT2A, KDM1A, KDM4A, KDM5B, MAFF, MAFK, MAX, MAZ, MXI1, MYOG, NFIC, NFYA, NRF1, PAX5, POLR2A, RAD21, RBBP5, RCOR1, RFX5, RUNX3, RXRA, SAP30, SMARCB1, SMC3, SP1, SP4, SPI1, SRF, STAT1, TAF1, TBL1XR1, TBP, TCF12, TRIM28, UBTF, USF2, WRNIP1, YY1, ZBTB7A, ZC3H11A, ZEB1, ZKSCAN1, ZMIZ1, ZNF143, ZNF263, ZNF384 |
| *BCL2L14* | NFE2L2, ELK1, STAT3, TP63, SPI1, NANOG, POU5F1, SOX2, TCF3, EGR1, HNF4A, PPARD, ERG, RUNX2, BACH1, NR1I2, CHD1, CHD2, CTCF, EBF1, ELF1, EP300, EZH2, FOXA1, FOXA2, H2AFZ, HDAC2, IRF1, KDM5A, KDM5B, MAX, MAZ, MTA3, MXI1, MYC, NRF1, POLR2A, RAD21, RBBP5, SMC3, SRF, TAF1, TBP, UBTF, WRNIP1, ZEB1, ZNF143 |
| *MYO5C* | ZNF217, ELK1, STAT3, TCF4, TP63, FLI1, MYC, KLF4, SOX2, PPARG, FOXA2, HNF4A, TFAP2C, TET1, TAL1, TRIM28, EP300, ESR1, BMI1, ESRRB, HSF1, BACH1, BHLHE40, CEBPD, CHD1, CHD2, CHD7, CREB1, CTBP2, CTCF, E2F4, E2F6, ELF1, EZH2, GABPA, GATA1, GATA3, GTF2F1, H2AFZ, HCFC1, HDAC2, HDAC6, JUND, KDM4A, KDM5A, KDM5B, MAFK, MAX, MAZ, MXI1, MYBL2, MYOG, NFIC, NFYA, NFYB, NR2F2, NRF1, PAX5, PHF8, PML, POLR2A, RAD21, RBBP5, RCOR1, RFX5, RNF2, SAP30, SIN3A, SP1, SUZ12, TAF1, TAF7, TBP, TCF12, TCF3, TCF7L2, USF1, USF2, WRNIP1, YY1, ZBTB7A, ZEB1, ZNF143 |
| *SCHIP1* | ZNF217, AR, TCF4, SMAD4, RUNX1, MYC, NANOG, POU5F1, SOX2, TCF3, EGR1, TP53, FOXA2, GATA1, ZFX, RCOR3, REST, SETDB1, ASH2L, TAL1, EP300, OLIG2, SMARCA4, YAP1, SALL4, TFCP2L1, MYB, POU3F2, SMAD1, PDX1, ATF1, CHD7, CREB1, CTCF, E2F4, EZH2, FOXA1, GATA2, GATA3, HDAC2, HDAC6, JUN, JUND, MAX, MXI1, PHF8, POLR2A, RAD21, RCOR1, RFX5, SAP30, SIN3A, TEAD4, USF1, ZNF143 |
| *GPRC5B* | STAT3, TCF4, CREM, NANOG, POU5F1, SOX2, TCF3, EGR1, MITF, PPARG, RCOR3, REST, SETDB1, EOMES, TFAP2C, SOX9, PRDM14, RUNX2, EZH2, RNF2, MTF2, SUZ12, PBX1, BACH1, ARID3A, BHLHE40, BRCA1, CBX3, CCNT2, CEBPB, CHD1, CHD2, CHD4, CREB1, CTCF, CTCFL, E2F4, E2F6, ELF1, EP300, FOS, FOXA1, FOXA2, FOXP2, GABPA, GTF2F1, H2AFZ, HCFC1, HDAC1, HDAC2, HMGN3, IRF4, JUND, KDM4A, KDM5A, KDM5B, MAFF, MAFK, MAX, MAZ, MXI1, MYBL2, MYC, MYOD1, MYOG, NFE2, NFIC, PAX5, PHF8, POLR2A, POU2F2, RAD21, RBBP5, RCOR1, RFX5, RUNX3, SAP30, SIN3A, SMC3, SP1, SP4, STAT1, TAF1, TAL1, TBL1XR1, TBP, TCF12, THAP1, UBTF, USF1, USF2, WHSC1, WRNIP1, YY1, ZBTB7A, ZC3H11A, ZEB1, ZNF143, ZNF263, ZNF384 |

| | |
|---|---|
| *LANCL2* | PHF8, AR, TP63, CREB1, CREM, E2F1, RUNX1, MYC, SPI1, POU5F1, SOX2, MITF, ZFX, SIN3B, PPARD, DMRT1, KLF1, SCLY, MECOM, FOXP2, RCOR1, PRDM5, TCF7L2, PAX6, NFIB, ARID3A, ATF1, ATF2, ATF3, BACH1, BCL3, BCLAF1, BHLHE40, BRCA1, CBX3, CCNT2, CEBPB, CEBPD, CHD1, CHD2, CHD7, CTBP2, CTCF, CTCFL, CUX1, E2F4, E2F6, EBF1, EGR1, ELF1, ELK1, ELK4, EP300, ETS1, EZH2, FOS, FOSL2, FOXA1, FOXA2, FOXM1, GABPA, GATA1, GATA2, GATA3, GTF2B, GTF2F1, H2AFZ, HCFC1, HDAC1, HDAC2, HDAC6, HMGN3, IRF1, IRF3, IRF4, JUN, JUND, KAT2A, KAT2B, KDM1A, KDM4A, KDM5A, KDM5B, MAFF, MAFK, MAX, MAZ, MTA3, MXI1, MYBL2, MYOG, NELFE, NFATC1, NFIC, NFYA, NR2F2, NR3C1, NRF1, PAX5, PBX3, PML, POLR2A, POU2F2, RAD21, RBBP5, RELA, REST, RFX5, RNF2, RUNX3, SAP30, SETDB1, SIN3A, SIRT6, SIX5, SMARCB1, SMC3, SP1, SP4, SREBF1, SRF, STAT1, STAT3, STAT5A, SUZ12, TAF1, TAF7, TAL1, TBL1XR1, TBP, TCF12, TCF3, TEAD4, THAP1, TRIM28, UBTF, USF1, USF2, WHSC1, WRNIP1, YY1, ZBTB33, ZBTB7A, ZC3H11A, ZEB1, ZKSCAN1, ZMIZ1, ZNF143, ZNF263, ZNF384 |
| *RNF224* | RUNX1, TCF3, TP53, GATA2, FOXP1, TAL1, SOX9, SRY, TFCP2L1, LMO2, ARID3A, BACH1, BHLHE40, BRCA1, CBX3, CCNT2, CEBPB, CEBPD, CHD1, CHD2, CTCF, CUX1, EBF1, EGR1, ELF1, ELK1, ELK4, EP300, ESR1, EZH2, FOS, FOXA1, FOXM1, GABPA, GATA1, GATA3, GTF2F1, H2AFZ, HCFC1, HDAC2, HDAC6, HMGN3, IRF4, JUN, JUND, KDM1A, KDM5A, MAFK, MAX, MAZ, MTA3, MXI1, MYBL2, MYC, NFIC, NFYA, NR2F2, NR3C1, NRF1, PAX5, PHF8, POLR2A, POU2F2, RAD21, RBBP5, RCOR1, REST, RFX5, RUNX3, SAP30, SIN3A, SMARCB1, SMARCC1, SMC3, SP1, SP4, STAT1, STAT3, STAT5A, SUZ12, TAF1, TBL1XR1, TBP, TCF12, TCF7L2, TEAD4, TFAP2A, TFAP2C, UBTF, USF2, WHSC1, WRNIP1, YY1, ZBTB7A, ZC3H11A, ZEB1, ZKSCAN1, ZMIZ1, ZNF143, ZNF217, ZNF263, ZNF384 |
| *ZBP1* | AR, CREM, RUNX1, NANOG, POU5F1, MITF, ASH2L, SRY, FOXO3, ESR1, SMAD2, SMAD3, STAT4, CTCF, PBX1, BACH1, STAT5A, GATA3, IRF1, IRF8, STAT6, RARG, THRA, ATF2, BATF, BCL11A, BCL3, BCLAF1, BHLHE40, BRCA1, CBX2, CBX8, CEBPB, CHD1, CHD2, CREB1, CUX1, E2F4, EBF1, EGR1, ELF1, ELK1, EP300, ETS1, EZH2, FOS, FOXM1, H2AFZ, IKZF1, IRF4, JUN, JUND, KDM5B, MAFK, MAX, MAZ, MEF2A, MEF2C, MTA3, MXI1, MYC, MYOD1, MYOG, NFATC1, NFIC, NFYA, NFYB, NRF1, PAX5, PBX3, PML, POLR2A, POU2F2, RAD21, RCOR1, RELA, RUNX3, SAP30, SETDB1, SIN3A, SMC3, SP1, SPI1, SRF, STAT1, STAT3, SUZ12, TBL1XR1, TBP, TCF12, TCF3, USF2, WRNIP1, YY1, ZEB1, ZNF143, ZNF384 |
| *JRK* | E2F1, ETS1, KDM5A, VDR, STAT3, TCF4, CREM, MYC, POU5F1, EGR1, HNF4A, SIN3B, REST, TET1, MYCN, ERG, SRY, FOXO3, YY1, CTCF, RAD21, ELF5, BACH1, HSF1, BCL3, PRDM5, DCP1A, ELF1, THAP11, RARG, ARID3A, ATF2, ATF3, BATF, BCLAF1, BHLHE40, BRCA1, CBX3, CCNT2, CEBPB, CEBPD, CHD1, CHD2, CHD7, CREB1, CTBP2, CTCFL, CUX1, E2F4, E2F6, EBF1, ELK1, ELK4, EP300, ESRRA, EZH2, FOS, FOSL2, FOXA1, FOXA2, FOXM1, FOXP2, GABPA, GATA1, GATA3, GTF2B, GTF2F1, H2AFZ, HCFC1, HDAC1, HDAC2, HDAC6, HMGN3, IRF1, JUN, JUND, KAT2A, KAT2B, KDM4A, MAFF, MAFK, MAX, MAZ, MBD4, MEF2C, MTA3, MXI1, MYBL2, MYOG, NANOG, NELFE, NFIC, NFYA, NR2F2, NR3C1, NRF1, PAX5, PBX3, PHF8, PML, POLR2A, POU2F2, RBBP5, RCOR1, RELA, RFX5, RUNX3, SAP30, SETDB1, SIN3A, SIX5, SMARCB1, SMC3, SP1, SP2, SP4, SREBF1, SREBF2, SRF, STAT1, STAT5A, SUPT20H, TAF1, TAF7, TAL1, TBL1XR1, TBP, TCF12, TCF3, TCF7L2, TEAD4, THAP1, TRIM28, UBTF, USF1, USF2, WRNIP1, ZBTB33, ZBTB7A, ZC3H11A, ZEB1, ZKSCAN1, ZMIZ1, ZNF143, ZNF384 |
| *CELSR3* | HOXC9, ELK1, STAT3, CREB1, CREM, E2F1, FLI1, RUNX1, MYC, NANOG, POU5F1, SOX2, MITF, RCOR3, SIN3B, REST, ASH2L, FOXP1, ERG, GFI1B, TRIM28, SRY, SOX17, WT1, TFCP2L1, YY1, FOXP2, BMI1, EZH2, SUZ12, SIN3A, TFEB, DNAJC2, CRX, RCOR2, ARID3A, ATF1, ATF2, ATF3, BACH1, BATF, BCL3, BCLAF1, BHLHE40, BRCA1, CBX3, CCNT2, CEBPB, CEBPD, CHD1, CHD2, CHD7, CTBP2, CTCF, CTCFL, E2F4, E2F6, EBF1, EGR1, ELF1, ELK4, EP300, ESRRA, ETS1, FOS, FOSL1, FOSL2, FOXA1, FOXA2, FOXM1, GABPA, GATA2, GATA3, GTF2B, GTF2F1, H2AFZ, HCFC1, HDAC1, HDAC2, HDAC6, HMGN3, HNF4A, HNF4G, IRF1, IRF3, IRF4, JUN, JUND, KDM1A, KDM4A, KDM5A, KDM5B, MAFK, MAX, MAZ, MBD4, MTA3, MXI1, MYBL2, NFATC1, NFIC, NFYA, NFYB, NR2F2, NR3C1, NRF1, PAX5, PBX3, PHF8, PML, POLR2A, POU2F2, RAD21, RBBP5, RCOR1, RELA, RFX5, RUNX3, SAP30, SIRT6, SIX5, SMARCB1, SMC3, SP1, SP2, SP4, SPI1, SREBF1, SRF, STAT1, STAT5A, TAF1, TAF7, TBL1XR1, TBP, TCF12, TCF3, TCF7L2, TEAD4, THAP1, UBTF, USF1, USF2, WRNIP1, ZBTB33, ZBTB7A, ZC3H11A, ZEB1, ZKSCAN1, ZMIZ1, ZNF143, ZNF263, ZNF384 |
| *GUSB* | TFAP2A, ETS1, AR, STAT3, E2F1, MYC, SPI1, HNF4A, SETDB1, PPARD, TET1, ZNF281, KLF1, CUX1, YY1, SRF, TBX5, SUZ12, PRDM5, NR1H3, ARID3A, ATF1, ATF2, ATF3, BCL11A, BCL3, BCLAF1, BHLHE40, BRCA1, CBX3, CCNT2, CEBPB, CEBPD, CHD1, CHD2, CHD7, CREB1, CTCF, CTCFL, E2F4, E2F6, EBF1, EGR1, ELF1, ELK1, EP300, ESRRA, EZH2, FOS, FOSL1, FOSL2, FOXA1, FOXA2, FOXM1, FOXP2, GABPA, GATA3, GTF2B, GTF2F1, GTF3C2, H2AFZ, HCFC1, HDAC1, HDAC2, HMGN3, IKZF1, IRF1, IRF4, JUN, JUND, KAT2A, KDM4A, KDM5B, MAFF, MAFK, MAX, MAZ, MBD4, MTA3, MXI1, MYBL2, MYOD1, MYOG, NANOG, NELFE, NFATC1, NFE2, NFIC, NR2C2, NR2F2, NR3C1, NRF1, PAX5, PBX3, PHF8, PML, POLR2A, POU2F2, RAD21, RBBP5, RCOR1, RELA, REST, RFX5, RUNX3, RXRA, SAP30, SIN3A, SIX5, SMARCB1, SMC3, SP1, SP4, STAT1, STAT5A, TAF1, TAF7, TAL1, TBL1XR1, TBP, TCF12, TCF3, TCF7L2, TEAD4, THAP1, TRIM28, UBTF, USF1, USF2, WHSC1, WRNIP1, ZBTB33, ZBTB7A, ZC3H11A, ZEB1, ZKSCAN1, ZMIZ1, ZNF143, ZNF263, ZNF384 |
| *CD69* | EGR1, EGR3, FOS, JUN, NFKB1, VDR, FOXP3, ELK1, AR, STAT3, TCF4, RUNX1, SPI1, PPARG, FOXO3, HOXB4, MECOM, STAT4, PBX1, MYB, RELA, BACH1, IRF8, STAT6, ARID3A, ATF1, ATF2, BCL11A, BCL3, BCLAF1, BHLHE40, CCNT2, CEBPB, CEBPD, CHD1, CHD2, CREB1, CTCF, CUX1, EBF1, EP300, ETS1, EZH2, FOXM1, GATA3, GTF2B, GTF2F1, H2AFZ, HCFC1, HDAC1, HDAC2, HMGN3, IKZF1, IRF1, IRF3, JUND, KAT2A, KDM5A, KDM5B, MAFF, MAFK, MAX, MAZ, MEF2A, MTA3, MXI1, MYC, NELFE, NFATC1, NFE2, NFIC, PAX5, PHF8, PML, POLR2A, POU2F2, RBBP5, RCOR1, REST, RFX5, RUNX3, SAP30, SIN3A, SIRT6, SMC3, SP1, STAT1, STAT5A, TAF1, TAF7, TBL1XR1, TBP, TCF3, TEAD4, UBTF, USF2, WHSC1, WRNIP1, YY1, ZMIZ1, ZNF143, ZNF384 |

| | |
|---|---|
| *IQSEC3* | VDR, TTF2, CREB1, KLF4, POU5F1, EGR1, GATA1, RCOR3, SIN3B, REST, FOXP1, TET1, CNOT3, TRIM28, ESR1, OLIG2, DMRT1, SCLY, YY1, JARID2, SUZ12, SIN3A, BATF, BCL11A, BHLHE40, CBX3, CEBPB, CEBPD, CHD1, CTCF, E2F4, EBF1, ETS1, EZH2, FOS, FOSL2, FOXA1, FOXA2, FOXP2, GATA2, H2AFZ, HCFC1, HDAC1, HDAC2, JUND, KDM4A, MAX, MAZ, MXI1, MYBL2, MYC, NR2F2, NR3C1, PAX5, PBX3, PHF8, POLR2A, RBBP5, RCOR1, RXRA, SAP30, SP1, SPI1, STAT3, STAT5A, TAF1, TBP, TCF12, TCF3, TEAD4, UBTF, USF1, ZBTB33, ZNF143, ZNF263 |
| *SHH* | GLI3, AHR, ARNT, ZNF217, STAT3, SMAD4, TP63, RUNX1, MYC, KLF4, POU5F1, SOX2, TCF3, EGR1, MITF, TP53, PPARG, HNF4A, KDM5B, REST, FOXP1, SRY, YAP1, SOX17, RUNX2, BMI1, EED, PHC1, EZH2, RNF2, JARID2, MTF2, SUZ12, JUN, CDX2, ARID3A, BACH1, CEBPB, CHD1, CHD2, CTBP2, CTCF, FOS, FOSL2, FOXA1, H2AFZ, HDAC2, HDAC6, HNF4G, JUND, MAX, MAZ, MXI1, MYBL2, NRF1, POLR2A, RAD21, RCOR1, RXRA, SIN3A, SP1, TCF7L2, YY1, ZBTB7A, ZNF143 |
| *ZNF556* | MYCN, SCLY, BACH1, ZNF274, ARID3A, ATF1, ATF3, BHLHE40, BRCA1, CBX3, CCNT2, CEBPB, CEBPD, CEBPZ, CHD1, CHD2, CTCF, CUX1, ELF1, EP300, EZH2, FOS, GABPA, GATA1, GATA2, GTF2F1, H2AFZ, HCFC1, HDAC2, HNF4A, HNF4G, IRF1, IRF3, JUN, JUND, KDM1A, MAFF, MAFK, MAX, MAZ, MBD4, MXI1, MYBL2, MYC, NFYA, NFYB, NR2F2, PBX3, PML, POLR2A, RBBP5, RCOR1, REST, RFX5, SIN3A, SMC3, SP1, SP2, SREBF1, SRF, STAT5A, TAF1, TAL1, TBL1XR1, TBP, TEAD4, UBTF, USF1, USF2, YY1, ZBTB7A, ZMIZ1, ZNF143 |
| *APOL1* | SOX2, HNF4A, FOXP1, PPARD, CUX1, TFAP2A, DNAJC2, IRF1, IRF8, ARID3A, BATF, BHLHE40, CEBPB, CEBPD, CHD1, CHD2, CTCF, E2F6, EBF1, EGR1, ELF1, EP300, EZH2, FOS, FOSL2, FOXA1, FOXA2, H2AFZ, HCFC1, HDAC1, HDAC2, HDAC6, HNF4G, IRF4, JUN, JUND, KDM5B, MAFF, MAFK, MAX, MAZ, MBD4, MTA3, MXI1, MYBL2, MYC, NFIC, NR3C1, PAX5, PHF8, POLR2A, PRDM1, RBBP5, RCOR1, REST, RUNX3, RXRA, SAP30, SIN3A, SP1, STAT1, STAT2, STAT3, STAT5A, TAF1, TAL1, TBL1XR1, TBP, TCF12, TEAD4, UBTF, USF1, WRNIP1, YY1, ZBTB7A, ZNF143 |
| *SOWAHA* | RUNX1, TCF3, MITF, TP53, HNF4A, RCOR3, SIN3B, TRIM28, YAP1, KLF1, EZH2, RNF2, JARID2, MTF2, SUZ12, ESRRB, RCOR1, CDX2, NOTCH1, BACH1, BHLHE40, CHD1, CHD2, CHD7, CREB1, CTBP2, CTCF, E2F4, E2F6, EBF1, EGR1, ELF1, EP300, FOXA2, FOXP2, GABPA, GATA1, GTF2F1, H2AFZ, HDAC2, HMGN3, HNF4G, JUND, KDM4A, MAX, MAZ, MYC, MYOG, NFYA, PAX5, PHF8, POLR2A, RAD21, RBBP5, RFX5, SAP30, SIN3A, SPI1, TAL1, TBP, TCF12, YY1, ZBTB7A, ZNF143, ZNF263, ZNF384 |
| *PLCE1* | FOXP3, AR, STAT3, TP63, RUNX1, NANOG, POU5F1, SOX2, MITF, PPARG, FOXA2, HNF4A, GATA1, KDM5B, RCOR3, SIN3B, REST, EOMES, ESR1, YAP1, NR3C1, RNF2, RCOR1, RELA, ATF3, BACH1, BCL3, CCNT2, CEBPB, CHD1, CHD7, CREB1, CTBP2, CTCF, E2F1, E2F6, EGR1, EP300, EZH2, FOS, FOSL1, FOSL2, FOXA1, GATA3, H2AFZ, HDAC2, HMGN3, JUN, JUND, KDM4A, MAX, MAZ, MXI1, MYC, NFIC, NRF1, PHF8, POLR2A, RAD21, RBBP5, SIN3A, SP1, TAF1, TBP, TCF12, TCF7L2, TEAD4, TRIM28, UBTF, USF1, YY1, ZBTB7A, ZNF143, ZNF263 |
| *TCTE3* | TTF2, DACH1, CREB1, FLI1, RUNX1, SPI1, NANOG, POU5F1, EGR1, PPARG, REST, FOXP1, TET1, EP300, SMARCA4, CTNNB1, LMO2, MEIS1, MYBL1, ARID3A, ATF1, ATF2, ATF3, BACH1, BCLAF1, BHLHE40, BRCA1, CBX3, CCNT2, CEBPB, CEBPD, CHD1, CHD2, CHD7, CTCF, CTCFL, CUX1, E2F1, E2F4, E2F6, EBF1, ELF1, ELK1, ELK4, ETS1, EZH2, FOS, FOSL1, FOSL2, FOXA1, FOXA2, FOXM1, FOXP2, GABPA, GATA1, GATA2, GATA3, GTF2B, GTF2F1, H2AFZ, HCFC1, HDAC2, HDAC6, HMGN3, HNF4A, IRF1, IRF3, IRF4, JUN, JUND, KAT2A, KDM4A, KDM5A, KDM5B, MAFK, MAX, MAZ, MBD4, MEF2A, MTA3, MXI1, MYBL2, MYC, NELFE, NFATC1, NFIC, NFYA, NFYB, NR2F2, NRF1, PAX5, PBX3, PHF8, PML, POLR2A, POU2F2, RAD21, RBBP5, RCOR1, RELA, RFX5, RNF2, RUNX3, SAP30, SETDB1, SIN3A, SIRT6, SIX5, SMARCB1, SMC3, SP1, SP4, SRF, STAT1, STAT3, STAT5A, SUZ12, TAF1, TAF7, TAL1, TBL1XR1, TBP, TCF12, TCF3, TCF7L2, TEAD4, THAP1, TRIM28, UBTF, USF1, USF2, WHSC1, WRNIP1, YY1, ZBTB33, ZBTB7A, ZC3H11A, ZEB1, ZKSCAN1, ZMIZ1, ZNF143, ZNF263, ZNF384 |
| *LEKR1* | AR, STAT3, SMAD4, TP63, CREB1, CREM, FLI1, RUNX1, SPI1, NANOG, POU5F1, SOX2, TCF3, MITF, PPARG, HNF4A, ZFX, SIN3B, ASH2L, PPARD, EP300, ESR1, PRDM14, RUNX2, ESRRB, SMAD1, PADI4, ARID3A, ATF1, ATF2, ATF3, BACH1, BCL3, BCLAF1, BHLHE40, BRCA1, CBX3, CCNT2, CEBPB, CEBPD, CHD1, CHD2, CHD7, CTBP2, CTCF, CUX1, E2F4, E2F6, EBF1, EGR1, ELF1, ELK1, ELK4, ETS1, EZH2, FOS, FOSL1, FOSL2, FOXA1, FOXM1, GABPA, GATA1, GATA2, GATA3, GTF2B, GTF2F1, H2AFZ, HCFC1, HDAC1, HDAC2, HDAC6, HMGN3, IKZF1, IRF1, IRF4, JUN, JUND, KDM1A, KDM4A, KDM5A, KDM5B, MAFK, MAX, MAZ, MTA3, MXI1, MYBL2, MYC, NCOR1, NELFE, NFATC1, NFIC, NFYA, NFYB, NR2F2, NR3C1, NRF1, PAX5, PHF8, PML, POLR2A, POU2F2, RAD21, RBBP5, RCOR1, RELA, REST, RFX5, RUNX3, SAP30, SETDB1, SIN3A, SIRT6, SMARCB1, SMARCC1, SMC3, SP1, STAT1, STAT5A, SUZ12, TAF1, TAL1, TBL1XR1, TBP, TCF12, TCF7L2, TEAD4, THAP1, TRIM28, UBTF, USF1, USF2, WHSC1, WRNIP1, YY1, ZBTB7A, ZC3H11A, ZEB1, ZKSCAN1, ZMIZ1, ZNF143, ZNF263, ZNF384 |
| *ATF7IP2* | AR, TP63, RUNX1, MYC, NANOG, SOX2, FOXA2, HNF4A, GATA2, RCOR3, ASH2L, PPARD, TET1, PRDM14, RUNX2, JUN, PBX1, BACH1, RCOR2, ARID3A, ATF1, BCL3, BCLAF1, BHLHE40, BRCA1, CBX3, CCNT2, CEBPB, CEBPD, CHD1, CHD2, CHD7, CREB1, CTBP2, CTCF, CUX1, E2F4, E2F6, EBF1, EGR1, ELF1, ELK1, EP300, ETS1, EZH2, FOS, FOXA1, FOXM1, FOXP2, GABPA, GTF2B, GTF2F1, H2AFZ, HCFC1, HDAC1, HDAC2, HDAC6, HMGN3, IRF1, IRF3, IRF4, JUND, KDM1A, KDM4A, KDM5B, MAFF, MAFK, MAX, MAZ, MBD4, MTA3, MXI1, MYBL2, NFE2, NFIC, NFYA, NFYB, NR2F2, NRF1, PAX5, PHF8, PML, POLR2A, POU2F2, RAD21, RBBP5, RCOR1, RELA, REST, RFX5, RUNX3, SAP30, SETDB1, SIN3A, SIRT6, SMC3, SP1, SP2, SP4, SPI1, STAT1, STAT3, STAT5A, TAF1, TAF7, TAL1, TBL1XR1, TBP, TCF12, TCF3, TCF7L2, TEAD4, TRIM28, UBTF, USF1, USF2, WRNIP1, YY1, ZBTB7A, ZC3H11A, ZMIZ1, ZNF143, ZNF274, ZNF384 |

| | |
|---|---|
| *CYP26A1* | AR, RARA, RARG, SP1, SP3, TP63, CREB1, KLF4, POU5F1, TP53, REST, EOMES, SRY, ESR1, TFCP2L1, BMI1, EED, PHC1, EZH2, RNF2, JARID2, MTF2, SUZ12, ARID3A, BACH1, BHLHE40, BRCA1, CBX3, CCNT2, CEBPD, CHD1, CHD2, CHD4, CHD7, CTBP2, CTCF, CTCFL, CUX1, E2F6, EGR1, ELF1, ELK1, EP300, FOXA1, FOXA2, FOXP2, GABPA, GATA2, GATA3, GTF2B, GTF2F1, H2AFZ, HCFC1, HDAC1, HDAC2, HDAC6, HMGN3, HNF4A, IRF1, JUND, KAT2B, KDM1A, KDM4A, KDM5B, MAFK, MAX, MAZ, MBD4, MXI1, MYBL2, MYC, MYOG, NCOR1, NFIC, NFYB, NR2F2, PHF8, PML, POLR2A, RAD21, RBBP5, RCOR1, RFX5, RXRA, SAP30, SETDB1, SIN3A, SIRT6, SMC3, SPI1, SRF, STAT5A, TAF1, TAL1, TBL1XR1, TBP, TCF12, TCF7L2, TEAD4, UBTF, USF1, USF2, WHSC1, YY1, ZBTB7A, ZC3H11A, ZEB1, ZMIZ1, ZNF143, ZNF263, ZNF384 |
| *PRAME* | STAT3, SMAD4, SPI1, EGR1, MITF, TP53, WT1, CUX1, JUN, ARID3A, ATF1, ATF3, BACH1, BHLHE40, BRCA1, CBX2, CBX3, CCNT2, CEBPB, CEBPD, CHD2, CHD4, CREB1, CTBP2, CTCF, CTCFL, E2F4, E2F6, EBF1, ELF1, ELK1, ELK4, EP300, ETS1, EZH2, FOSL2, FOXA1, FOXA2, FOXP2, GABPA, GTF2B, GTF2F1, GTF3C2, H2AFZ, HCFC1, HDAC2, HMGN3, IRF1, JUND, MAFF, MAFK, MAX, MAZ, MXI1, MYC, NCOR1, NFYA, NFYB, NR2F2, NR3C1, NRF1, PAX5, PHF8, PML, POLR2A, POU2F2, RAD21, RBBP5, RCOR1, REST, RFX5, RNF2, RUNX3, SETDB1, SIN3A, SIX5, SMARCA4, SMARCB1, SMC3, SP1, SP2, SRF, STAT1, STAT2, STAT5A, SUZ12, TAF1, TAF7, TAL1, TBL1XR1, TBP, TCF12, TCF7L2, TEAD4, THAP1, TRIM28, UBTF, USF1, USF2, YY1, ZBTB33, ZBTB7A, ZC3H11A, ZKSCAN1, ZMIZ1, ZNF143, ZNF263, ZNF384 |
| *SYT12* | AHR, ARNT, AR, STAT3, TCF4, RUNX1, SPI1, EGR1, TP53, PPARG, HNF4A, TFAP2C, ERG, TRIM28, ESR1, WT1, TEAD4, ATF3, RUNX2, EZH2, RNF2, JARID2, MTF2, SUZ12, PBX1, ELF5, EWSR1, BACH1, NR1I2, ELF1, RBPJ, ATF2, BATF, BCL11A, BCL3, BHLHE40, BRCA1, CBX3, CEBPB, CHD1, CHD2, CHD7, CTCF, CTCFL, E2F1, E2F4, E2F6, EBF1, EP300, ETS1, FOS, FOSL2, FOXM1, GABPA, GATA1, GATA3, GTF2F1, H2AFZ, HCFC1, HDAC1, HDAC2, IRF4, JUN, JUND, KDM4A, KDM5A, MAFK, MAX, MAZ, MEF2C, MTA3, MXI1, MYC, MYOD1, MYOG, NFIC, NR2F2, PAX5, PHF8, POLR2A, POU2F2, PRDM1, RAD21, RBBP5, RCOR1, RFX5, RUNX3, SAP30, SIN3A, SMARCB1, SMARCC1, SMC3, SP1, STAT1, STAT5A, TBL1XR1, TBP, TCF12, TCF3, TCF7L2, TFAP2A, USF2, YY1, ZBTB7A, ZC3H11A, ZKSCAN1, ZNF143, ZNF217, ZNF384 |
| *ANKRD10* | AR, SMAD4, TP63, CREM, RUNX1, MYC, KLF4, NANOG, POU5F1, SOX2, TCF3, MITF, TP53, KDM5B, E2F4, SETDB1, TFAP2C, FOXP1, PPARD, CNOT3, TRIM28, SOX9, FOXO3, EP300, YAP1, SALL4, DMRT1, TEAD4, ATF3, SMAD2, SMAD3, RUNX2, CUX1, MEF2A, RAD21, MTF2, TFAP2A, JUN, STAT5A, NACC1, NR0B1, ZIC3, SREBF2, ARID3A, BACH1, BCL3, BCLAF1, BHLHE40, BRCA1, CBX3, CCNT2, CEBPB, CEBPD, CHD1, CHD2, CHD7, CREB1, CTCF, CTCFL, E2F1, E2F6, EBF1, EGR1, ELF1, ELK1, ELK4, ETS1, EZH2, FOS, FOXA1, FOXA2, FOXM1, FOXP2, GABPA, GATA1, GATA2, GATA3, GTF2F1, H2AFZ, HCFC1, HDAC1, HDAC2, HDAC6, HMGN3, IRF1, IRF3, JUND, KAT2A, KAT2B, KDM4A, KDM5A, MAFK, MAX, MAZ, MTA3, MXI1, MYBL2, MYOD1, MYOG, NELFE, NFATC1, NFE2, NFIC, NR2F2, NR3C1, NRF1, PAX5, PBX3, PHF8, PML, POLR2A, POU2F2, RBBP5, RCOR1, RELA, REST, RFX5, RNF2, RUNX3, SAP30, SIN3A, SIX5, SMARCB1, SMC3, SP1, SP4, SPI1, SRF, STAT1, STAT3, TAF1, TAF7, TAL1, TBL1XR1, TBP, TCF12, TCF7L2, UBTF, USF1, USF2, WRNIP1, YY1, ZBTB7A, ZC3H11A, ZEB1, ZKSCAN1, ZMIZ1, ZNF143, ZNF263, ZNF384 |
| *FAM69B* | TP63, KLF4, EGR1, MITF, TP53, PPARG, ZFX, ASH2L, WT1, TFCP2L1, ESRRB, NR0B1, ARID3A, ATF3, BACH1, BHLHE40, CBX3, CCNT2, CEBPB, CHD1, CHD4, CHD7, CTCF, CUX1, E2F6, EP300, EZH2, FOS, FOSL1, FOSL2, FOXA1, FOXA2, GABPA, GATA2, GATA3, GTF3C2, H2AFZ, HDAC1, HDAC2, HDAC6, HMGN3, IRF1, JUN, JUND, KDM1A, KDM4A, MAX, MAZ, MXI1, MYC, MYOG, NFIC, NR2F2, NR3C1, PHF8, POLR2A, POLR3A, RAD21, RBBP5, RCOR1, REST, SAP30, SIN3A, SIRT6, SP1, SP4, SUZ12, TAF1, TBP, TCF12, TCF7L2, UBTF, USF2, YY1, ZBTB7A, ZNF143, ZNF263, ZNF384 |
| *WNK2* | ZNF217, AR, STAT3, TP63, FLI1, MYC, EGR1, MITF, HNF4A, GATA2, TFAP2C, ERG, YAP1, NR3C1, SCLY, TCF7, EZH2, RNF2, JARID2, MTF2, SUZ12, PRDM5, GATA3, STAT1, BACH1, CCNT2, CHD1, CHD2, CREB1, CTBP2, CTCF, E2F6, ELF1, EP300, FOXP2, GABPA, GTF2F1, H2AFZ, HCFC1, HDAC2, HDAC6, HMGN3, IRF1, JUND, KDM4A, KDM5A, KDM5B, MAX, MAZ, MXI1, NFIC, NRF1, PHF8, POLR2A, RAD21, RBBP5, RCOR1, REST, RFX5, SAP30, SIN3A, SMC3, TAF1, TBP, TCF12, TCF7L2, UBTF, YY1, ZBTB7A, ZNF143, ZNF263 |
| *RASSF4* | PAX3, RUNX1, NANOG, SOX2, FOXA2, HNF4A, E2F4, TFAP2C, FOXP1, TAL1, EP300, STAT4, STAT5A, CDX2, NR1I2, NFIB, ARID3A, BACH1, BCLAF1, BHLHE40, BRCA1, CBX2, CCNT2, CEBPB, CEBPD, CHD1, CHD2, CHD4, CREB1, CTCF, CUX1, E2F6, EBF1, EGR1, ELF1, ELK1, ETS1, EZH2, FOS, FOSL2, FOXA1, GABPA, GATA1, GATA2, GATA3, GTF2F1, H2AFZ, HCFC1, HDAC1, HDAC2, HMGN3, HNF4G, IRF1, JUN, JUND, KAT2A, KDM1A, KDM4A, KDM5A, KDM5B, MAFK, MAX, MAZ, MTA3, MXI1, MYC, MYOG, NELFE, NFYB, NR2F2, NRF1, PAX5, PBX3, PHF8, PML, POLR2A, PRDM1, RAD21, RBBP5, RCOR1, RELA, REST, RFX5, RUNX3, RXRA, SAP30, SIN3A, SMC3, SP1, SPI1, SRF, STAT1, STAT3, TAF1, TBL1XR1, TBP, TCF12, TCF3, TCF7L2, TEAD4, TRIM28, UBTF, USF2, WRNIP1, YY1, ZBTB7A, ZC3H11A, ZEB1, ZKSCAN1, ZMIZ1, ZNF143, ZNF384 |
| *IL1RAPL2* | AHR, ARNT, AR, STAT3, TCF4, E2F1, SOX2, TP53, FOXA2, SMARCA4, NR3C1, RUNX2, MTF2, SUZ12, TFAP2A, PBX1, NR1I2, PRDM16, CHD1, CHD2, CHD7, CTBP2, CTCF, EP300, EZH2, GTF2F1, H2AFZ, HDAC1, HDAC2, HDAC6, IRF1, JUND, KDM4A, MAZ, MYC, PHF8, POLR2A, RAD21, RBBP5, SAP30, SIN3A, SMC3, TBP, ZNF143, ZNF263, ZNF384 |
| *BMP3* | ERG, OLIG2, NR3C1, RUNX2, EED, EZH2, RNF2, JARID2, MTF2, SUZ12, JUN, PBX1, BACH1, CHD1, CHD7, CTBP2, CTCF, E2F6, EP300, FOXP2, GABPA, GTF2F1, H2AFZ, HDAC2, HDAC6, KDM4A, MAX, MXI1, MYOG, NRF1, PHF8, POLR2A, RBBP5, RFX5, SAP30, SETDB1, SIN3A, SIRT6, TBP, TRIM28, ZNF143, ZNF263 |
| *HOXB8* | E2F1, STAT3, MYC, SPI1, POU5F1, TCF3, KDM5B, SIN3B, REST, SETDB1, RAD21, BMI1, EED, PHC1, EZH2, RNF2, JARID2, MTF2, SUZ12, EWSR1, CDX2, ELF1, RBPJ, BACH1, BCL3, CCNT2, CEBPB, CHD1, CHD2, CREB1, CTBP2, CTCF, E2F6, EP300, ETS1, FOSL2, GABPA, GTF2F1, H2AFZ, HDAC2, HDAC6, JUN, JUND, MAX, MAZ, NFYA, PAX5, POLR2A, RBBP5, RCOR1, RFX5, SAP30, SIN3A, SMC3, TAF1, TBP, TCF12, TCF7L2, WHSC1, YY1, ZNF143, ZNF263, ZNF384 |

| | |
|---|---|
| *ZIC5* | ELK1, PHF8, AR, STAT3, TCF4, CREM, MYC, KLF4, NANOG, POU5F1, SOX2, TCF3, EGR1, MITF, ZFX, SETDB1, EOMES, TFAP2C, FOXP1, TRIM28, SOX9, SMARCA4, ZNF281, CCND1, MYBL2, BMI1, EZH2, RNF2, JARID2, MTF2, SUZ12, TBX3, RCOR1, NACC1, NR0B1, RCOR2, ATF2, BACH1, BHLHE40, BRCA1, CBX3, CCNT2, CEBPB, CHD1, CHD2, CREB1, CTBP2, CTCF, E2F1, E2F4, E2F6, ELF1, EP300, ETS1, FOXP2, GABPA, GATA2, GTF2F1, H2AFZ, HCFC1, HDAC1, HDAC2, HDAC6, HMGN3, IRF1, JUND, KAT2A, KDM4A, KDM5B, MAFK, MAX, MAZ, MXI1, NELFE, NFIC, NFYA, NFYB, NRF1, PAX5, PBX3, POLR2A, RAD21, RBBP5, RFX5, SAP30, SIN3A, SMARCB1, SMC3, SP1, TAF1, TAF7, TAL1, TBL1XR1, TBP, TCF12, TCF7L2, TEAD4, UBTF, USF1, USF2, YY1, ZBTB33, ZBTB7A, ZKSCAN1, ZMIZ1, ZNF143, ZNF384 |
| *PTGIS* | ZNF217, PAX3, AR, STAT3, TCF4, TP63, NANOG, POU5F1, SOX2, PPARG, RCOR3, SIN3B, REST, SETDB1, ERG, SOX9, SRY, ZNF281, RUNX2, EED, SUZ12, SIN3A, RELA, CEBPB, SREBF2, NFIB, DROSHA, BACH1, CBX2, CBX8, CHD1, CHD2, CHD7, CTBP2, CTCF, E2F6, EGR1, EP300, EZH2, FOXA1, FOXA2, GTF2F1, H2AFZ, HDAC2, HDAC6, JUN, JUND, KDM4A, KDM5A, KDM5B, MAX, MXI1, MYC, MYOG, PHF8, POLR2A, RAD21, RBBP5, RNF2, SAP30, SIRT6, SP1, TAF1, TBP, TCF12, TCF7L2, TEAD4, YY1, ZBTB7A, ZNF143, ZNF263 |
| *MPPED2* | E2F1, E2F2, ZNF217, PAX3, AR, STAT3, CREB1, MYC, NANOG, POU5F1, MITF, TP53, FOXA2, HNF4A, REST, ASH2L, TET1, ERG, EP300, SALL4, BMI1, SUZ12, TFAP2A, POU3F2, ELF5, SOX11, BACH1, CHD1, CHD2, CHD7, CTBP2, CTCF, EGR1, ELF1, EZH2, GTF2F1, H2AFZ, HCFC1, HDAC2, HDAC6, JUND, KDM4A, KDM5A, KDM5B, MAX, MAZ, MXI1, MYOG, NFIC, NR2F2, PHF8, POLR2A, RAD21, RBBP5, SAP30, SIN3A, TAF1, TAL1, TBP, TCF12, TCF7L2, TEAD4, TRIM28, YY1, ZBTB7A, ZNF143, ZNF263, ZNF384 |
| *ACOT1* | E2F1, MYC, NANOG, POU5F1, SOX2, TCF3, TP53, PPARG, REST, SETDB1, EOMES, TFAP2C, CNOT3, TRIM28, SMARCA4, YAP1, HOXB4, MTF2, SUZ12, STAT5A, SOX11, NACC1, CEBPA, CLOCK, ARID3A, BACH1, BHLHE40, CEBPB, CEBPD, CHD1, CHD2, CHD7, CTCF, E2F4, E2F6, EBF1, ELF1, EP300, EZH2, FOS, FOSL2, FOXA1, FOXA2, FOXM1, GABPA, GATA1, GATA2, GATA3, GTF2F1, H2AFZ, HDAC2, HNF4A, HNF4G, IRF4, JUN, JUND, KDM4A, MAFF, MAFK, MAX, MAZ, MXI1, MYBL2, NFIC, NFYA, NR3C1, PAX5, PBX3, PHF8, POLR2A, POU2F2, RAD21, RBBP5, RCOR1, RFX5, RXRA, SIN3A, SMARCB1, SMC3, SP1, SPI1, SREBF1, SRF, STAT3, TAF1, TAF7, TAL1, TBP, TCF12, TEAD4, USF1, USF2, YY1, ZBTB7A, ZEB1, ZKSCAN1, ZNF143, ZNF384 |
| *FAM133A* | SMAD4, RUNX2, BRCA1, CHD1, CHD2, CHD7, CTBP2, CTCF, EBF1, EP300, EZH2, GTF2F1, H2AFZ, HDAC2, JUND, KDM4A, MYC, NFYB, PBX3, PHF8, POLR2A, RAD21, RBBP5, REST, SAP30, SIN3A, SIRT6, SP1, SP2, STAT3, TAF1, TBP, TCF12, TEAD4, YY1, ZNF143 |
| *CABLES1* | TP53, TP73, AHR, ARNT, PAX3, TTF2, XRN2, AR, STAT3, TCF4, CREM, E2F1, FLI1, MYC, SPI1, NANOG, POU5F1, SOX2, EGR1, MITF, FOXA2, HNF4A, EOMES, TFAP2C, ERG, CNOT3, TRIM28, OLIG2, SMARCA4, PHC1, EZH2, RNF2, JARID2, MTF2, SUZ12, TFAP2A, RCOR1, NUCKS1, ARID3A, BACH1, BCLAF1, BHLHE40, BRCA1, CBX2, CBX8, CEBPB, CEBPD, CHD1, CHD2, CREB1, CTCF, CUX1, E2F4, E2F6, EBF1, ELF1, ELK1, EP300, ETS1, FOS, FOXA1, FOXP2, GABPA, GATA2, GATA3, GTF2F1, H2AFZ, HDAC1, HDAC2, HMGN3, IKZF1, JUND, KAT2A, KDM4A, MAFK, MAX, MAZ, MTA3, MXI1, MYBL2, MYOD1, MYOG, NFIC, NR2F2, NRF1, PAX5, PHF8, PML, POLR2A, POU2F2, RAD21, RBBP5, RELA, REST, RFX5, RUNX3, RXRA, SAP30, SETDB1, SIN3A, SIX5, SMARCB1, SMC3, SP1, STAT1, STAT5A, TAF1, TAF7, TBL1XR1, TBP, TCF12, TCF3, TCF7L2, TEAD4, UBTF, USF1, USF2, WRNIP1, YY1, ZBTB7A, ZEB1, ZKSCAN1, ZNF143, ZNF263 |
| *KLRC2* | REST, ATF3, SMAD2, SMAD3, HOXB4, STAT4, RCOR1, RBPJ, PAX6, HTT, CEBPB, CHD2, ELK1, EP300, ESR1, FOS, FOSL2, FOXA1, FOXA2, GABPA, GATA2, GTF2F1, H2AFZ, IRF3, JUND, MAFK, MAX, MYC, NR2F2, NR3C1, PBX3, POLR2A, PRDM1, RAD21, RFX5, RXRA, SMC3, SP1, STAT1, STAT3, SUPT20H, TAF1, TAL1, TBP, TCF12, TCF7L2, TEAD4, TRIM28, USF2, YY1, ZKSCAN1, ZNF143 |
| *SAMD15* | HOXC9, FLI1, E2F4, SALL4, PBX1, ARID3A, BACH1, BCLAF1, BHLHE40, BRCA1, CBX3, CCNT2, CEBPB, CHD1, CHD2, CHD7, CTCF, CTCFL, CUX1, E2F6, EBF1, ELF1, ELK1, ELK4, EP300, ETS1, EZH2, FOXA1, FOXA2, FOXM1, FOXP2, GABPA, GATA1, GATA3, GTF2F1, H2AFZ, HCFC1, HDAC1, HDAC2, HDAC6, HMGN3, HNF4G, JUND, KDM4A, KDM5A, KDM5B, MAX, MAZ, MTA3, MXI1, MYC, NR2F2, NRF1, PAX5, PHF8, PML, POLR2A, RAD21, RBBP5, RCOR1, REST, RFX5, RUNX3, SAP30, SETDB1, SIN3A, SIRT6, SMARCB1, SMC3, STAT1, STAT3, STAT5A, SUZ12, TAF1, TBL1XR1, TBP, TCF12, TCF3, TCF7L2, TRIM28, UBTF, USF2, WRNIP1, YY1, ZBTB7A, ZC3H11A, ZEB1, ZKSCAN1, ZMIZ1, ZNF143, ZNF384 |
| *MUC16* | SMAD4, E2F1, TFAP2A, DROSHA, GBX2, CEBPB, EZH2, MAFK, POLR2A |
| *FOXK1* | ZNF217, NFE2L2, KDM5A, VDR, DACH1, AR, STAT3, TP63, CREM, E2F1, FLI1, RUNX1, SPI1, KLF4, EGR1, MITF, TP53, FOXA2, RCOR3, SIN3B, REST, TFAP2C, PPARD, ESR1, DMRT1, PRDM14, NR3C1, SCLY, TBX5, EZH2, SUZ12, CTNNB1, RELA, RCOR2, ETS2, ARID3A, ATF2, ATF3, BCL3, BCLAF1, BHLHE40, CBX3, CCNT2, CEBPB, CEBPD, CHD1, CHD2, CREB1, CTCF, CTCFL, CUX1, E2F4, E2F6, EBF1, ELF1, ELK1, EP300, ETS1, FOS, FOXA1, FOXM1, FOXP2, GABPA, GATA2, GATA3, GTF2F1, H2AFZ, HCFC1, HDAC1, HDAC2, HDAC6, HMGN3, IRF1, JUN, JUND, KAT2A, KDM1A, KDM4A, KDM5B, MAFK, MAX, MAZ, MTA3, MXI1, MYC, MYOG, NCOR1, NELFE, NFIC, NFYA, NR2F2, NRF1, PAX5, PHF8, PML, POLR2A, POU2F2, RAD21, RBBP5, RCOR1, RFX5, RUNX3, RXRA, SAP30, SETDB1, SIN3A, SMARCB1, SMARCC1, SMC3, SP1, SRF, STAT1, STAT5A, TAF1, TAL1, TBL1XR1, TBP, TCF12, TCF3, TCF7L2, TEAD4, UBTF, USF1, USF2, WHSC1, WRNIP1, YY1, ZBTB7A, ZC3H11A, ZKSCAN1, ZMIZ1, ZNF143, ZNF263, ZNF384 |

| | |
|---|---|
| *CEP70* | NFE2L2, XRN2, CREM, RUNX1, MYC, SPI1, NANOG, GATA1, GATA2, RCOR3, SIN3B, REST, SETDB1, TAL1, SRY, FOXO3, DMRT1, CUX1, SCLY, RAD21, JUN, SIN3A, LMO2, MEIS1, NACC1, NR0B1, ZIC3, CDX2, PAX6, RCOR2, ARID3A, ATF1, ATF2, BACH1, BCL3, BCLAF1, BHLHE40, BRCA1, CBX3, CCNT2, CEBPB, CEBPD, CHD1, CHD2, CHD7, CREB1, CTCF, E2F4, E2F6, EBF1, EGR1, ELF1, ELK1, EP300, ETS1, EZH2, FOS, FOXA1, FOXA2, FOXM1, FOXP2, GABPA, GATA3, GTF2B, GTF2F1, H2AFZ, HCFC1, HDAC1, HDAC2, HDAC6, HMGN3, IRF1, JUND, KAT2B, KDM1A, KDM4A, KDM5A, KDM5B, MAFF, MAFK, MAX, MAZ, MBD4, MTA3, MXI1, MYB, MYBL2, MYOD1, MYOG, NELFE, NFE2, NFIC, NFYA, NFYB, NR2F2, NRF1, PAX5, PBX3, PHF8, PML, POLR2A, POU2F2, PRDM1, RBBP5, RCOR1, RELA, RFX5, RUNX3, RXRA, SAP30, SMARCB1, SMC3, SP1, SP4, SRF, STAT1, STAT3, SUZ12, TAF1, TAF7, TBL1XR1, TBP, TCF12, TCF3, TCF7L2, TEAD4, TRIM28, UBTF, USF2, WRNIP1, YY1, ZBTB33, ZBTB7A, ZC3H11A, ZEB1, ZKSCAN1, ZMIZ1, ZNF143, ZNF263, ZNF384 |
| *EPS8* | PHF8, AR, STAT3, CREM, E2F1, MYC, NANOG, TP53, PPARG, HNF4A, GATA2, E2F4, REST, SETDB1, TFAP2C, FOXP1, TRIM28, DMRT1, PRDM14, SMAD2, SMAD3, NR3C1, CTCF, TBX3, RELA, SMAD1, CDX2, ARID3A, ATF3, BACH1, BCL3, BCLAF1, BHLHE40, BRCA1, CEBPB, CEBPD, CHD1, CHD2, CHD7, CREB1, CTBP2, E2F6, EBF1, EGR1, ELF1, ELK1, EP300, ETS1, EZH2, FOS, FOSL2, FOXA1, FOXA2, FOXM1, FOXP2, GABPA, GATA1, GATA3, GTF2F1, H2AFZ, HCFC1, HDAC1, HDAC2, HNF4G, JUND, KDM4A, KDM5A, KDM5B, MAFK, MAX, MAZ, MBD4, MXI1, MYBL2, NFIC, NFYB, NR2F2, NRF1, PAX5, PBX3, PML, POLR2A, POU2F2, RAD21, RBBP5, RCOR1, RFX5, RUNX3, SAP30, SIN3A, SMARCB1, SMARCC1, SMC3, SP1, SP4, SRF, STAT1, STAT5A, SUZ12, TAF1, TAF7, TBL1XR1, TBP, TCF12, TCF3, TCF7L2, TEAD4, USF1, USF2, WRNIP1, YY1, ZBTB33, ZBTB7A, ZEB1, ZKSCAN1, ZNF143, ZNF217, ZNF263, ZNF384 |
| *DGKB* | TP53, PAX3, AR, STAT3, TCF4, SMAD4, MYC, NANOG, POU5F1, SOX2, TCF3, MITF, PPARD, SOX9, EP300, SALL4, BACH1, GBX2, CEBPB, CTCF, EZH2, FOS, FOSL2, GATA3, H2AFZ, JUND, KDM5A, MAFK, NFIC, PBX3, POLR2A, TCF12 |
| *SLCO6A1* | AR, STAT3, TP63, CREM, REST, PRDM14, RUNX2, POU3F2, RCOR1, CDX2, CEBPB, CTCF, CTCFL, E2F6, EGR1, EZH2, FOS, H2AFZ, HDAC2, KDM5A, MAFF, MAFK, PHF8, POLR2A, RAD21, RBBP5 |
| *RARB* | AR, CREB1, MYC, PPARG, RARA, RARB, RARG, ELK1, STAT3, TCF4, TP63, NANOG, POU5F1, MITF, REST, TFAP2C, PPARD, CNOT3, EP300, ESR1, SALL4, SOX17, TEAD4, ATF3, SMAD2, SMAD3, RUNX2, CUX1, SCLY, GATA4, SRF, TBX5, RAD21, MTF2, SUZ12, POU3F2, BACH1, NR0B1, NR1I2, DNAJC2, IRF1, NFIB, DROSHA, ARID3A, ATF1, ATF2, BCL3, BHLHE40, CBX2, CBX8, CEBPB, CHD1, CHD2, CHD7, CTBP2, CTCF, ELF1, ETS1, EZH2, FOS, FOSL2, FOXA1, FOXA2, GATA2, GATA3, H2AFZ, HCFC1, HDAC1, HDAC2, HDAC6, HMGN3, JUND, KDM1A, KDM4A, KDM5A, KDM5B, MAFF, MAFK, MAX, MAZ, MXI1, NFIC, NR3C1, PBX3, PHF8, PML, POLR2A, RBBP5, RCOR1, RNF2, RXRA, SAP30, SIN3A, SIX5, SP1, TAF1, TBL1XR1, TBP, TCF12, TCF7L2, TRIM28, USF1, YY1, ZBTB33, ZBTB7A, ZMIZ1, ZNF143, ZNF263 |
| *CCL5* | ATF1, CEBPA, CEBPB, CEBPD, CREB1, FOS, FOSB, JUN, NFKB1, REL, RELA, SPI1, STAT3, TCF4, FLI1, GATA2, SCLY, MYB, LMO2, BACH1, NACC1, DNAJC2, IRF8, NR1H3, ARID3A, ATF2, BATF, BCL3, BCLAF1, BHLHE40, CHD1, CHD2, CTCF, CUX1, E2F6, EBF1, EGR1, EP300, EZH2, FOXM1, GABPA, H2AFZ, HDAC2, JUND, MAFF, MAFK, MAX, MAZ, MEF2A, MEF2C, MTA3, MXI1, MYC, NFATC1, NFIC, NFYB, PAX5, PHF8, PML, POLR2A, POU2F2, RCOR1, RUNX3, RXRA, SP1, SRF, STAT1, STAT5A, TAF1, TAL1, TBP, TCF12, TCF3, USF2, WHSC1, WRNIP1, YY1, ZBTB33, ZEB1, ZNF143, ZNF384 |
| *PAX9* | AR, STAT3, SMAD4, E2F1, KLF4, NANOG, POU5F1, SOX2, TCF3, TP53, KDM5B, SIN3B, SETDB1, TRIM28, TFCP2L1, SMAD3, BMI1, EED, PHC1, EZH2, RNF2, JARID2, MTF2, SUZ12, PBX1, ZIC3, HTT, ZNF274, ATF2, BACH1, BHLHE40, CHD1, CHD2, CHD7, CREB1, CTBP2, CTCF, CTCFL, E2F6, EGR1, ELF1, EP300, FOXA1, FOXA2, FOXP2, GABPA, GATA1, GATA3, H2AFZ, HCFC1, HDAC1, HDAC2, HDAC6, JUN, JUND, KDM4A, KDM5A, MAFK, MAX, MAZ, MXI1, MYC, POLR2A, RAD21, RBBP5, RCOR1, SAP30, SIN3A, SMC3, SP1, SPI1, TBP, TCF12, UBTF, USF2, YY1, ZNF143 |
| *LIF* | ETS1, ETS2, ZNF217, STAT3, TP63, RUNX1, NANOG, POU5F1, EGR1, TP53, HNF4A, GATA1, KDM5B, RCOR3, REST, SETDB1, ASH2L, TFAP2C, TET1, MYCN, TAL1, SOX9, SRY, EP300, YAP1, ZNF281, PRDM14, TFCP2L1, SMAD2, SMAD3, SCLY, STAT4, TBX5, FOXP2, TCF7, SUZ12, ESRRB, RCOR1, RBPJ, BACH1, BHLHE40, CEBPB, CHD1, CHD2, CTBP2, CTCF, E2F4, E2F6, EBF1, ELF1, EZH2, FOS, GABPA, GATA2, GTF2F1, H2AFZ, HDAC1, HDAC2, HMGN3, JUND, KDM4A, KDM5A, MAX, MAZ, MXI1, MYC, NELFE, NR2F2, NRF1, PBX3, PHF8, POLR2A, PRDM1, RAD21, RBBP5, RNF2, RXRA, SAP30, SIN3A, SMC3, TBP, TCF12, TCF7L2, TRIM28, UBTF, USF2, YY1, ZBTB7A, ZC3H11A, ZEB1, ZNF143, ZNF263 |
| *ID3* | E2F1, E2F4, MYC, SMAD1, TTF2, XRN2, SMAD4, TP63, CREB1, FLI1, RUNX1, SPI1, KLF4, NANOG, POU5F1, SOX2, TCF3, MITF, TP53, PPARG, FOXA2, HNF4A, GATA1, GATA2, RCOR3, REST, SETDB1, TFAP2C, FOXP1, PPARD, GFI1B, EP300, ESR1, SALL4, ZNF281, PRDM14, TFCP2L1, ATF3, SMAD2, SMAD3, NR3C1, CUX1, CCND1, GATA4, EED, PHC1, EZH2, RNF2, MTF2, SUZ12, ESRRB, TBX3, NACC1, NR0B1, BCL3, ELF1, FOXO1, BCL11B, ARID3A, ATF2, BATF, BCL11A, BCLAF1, BHLHE40, BRCA1, CBX3, CCNT2, CEBPB, CEBPD, CHD1, CHD2, CTCF, E2F6, EBF1, EGR1, ELK1, ELK4, ETS1, FOS, FOSL2, FOXA1, FOXM1, FOXP2, GABPA, GATA3, GTF2B, GTF2F1, H2AFZ, HCFC1, HDAC2, HMGN3, HNF4G, IRF1, JUN, JUND, KAT2A, KDM1A, KDM4A, KDM5A, KDM5B, MAFF, MAFK, MAX, MAZ, MBD4, MEF2A, MTA3, MXI1, MYBL2, MYOG, NELFE, NFATC1, NFIC, NR2F2, NRF1, PAX5, PBX3, PHF8, PML, POLR2A, POU2F2, PRDM1, RAD21, RBBP5, RCOR1, RELA, RFX5, RUNX3, RXRA, SIN3A, SIRT6, SIX5, SMARCB1, SMARCC1, SMC3, SP1, SP4, SRF, STAT1, STAT3, STAT5A, TAF1, TAF7, TAL1, TBL1XR1, TBP, TCF12, TCF7L2, TEAD4, TRIM28, UBTF, USF1, USF2, WHSC1, WRNIP1, YY1, ZBTB33, ZBTB7A, ZC3H11A, ZEB1, ZKSCAN1, ZMIZ1, ZNF143, ZNF263, ZNF384 |

| | |
|---|---|
| *MEST* | E2F1, MYC, KLF4, EGR1, KDM5B, SETDB1, FOXP1, TET1, TRIM28, EP300, ZNF281, CUX1, CCND1, MYBL2, FOXP2, CTCF, RAD21, BMI1, SUZ12, NACC1, PRDM5, ARID3A, ATF3, BACH1, BATF, BCL3, BHLHE40, BRCA1, CBX3, CEBPB, CEBPD, CHD1, CHD2, CHD7, CREB1, E2F4, E2F6, ELF1, ELK1, ETS1, EZH2, FOS, FOXA2, FOXM1, GATA3, GTF2F1, H2AFZ, HCFC1, HDAC1, HDAC2, HMGN3, JUN, JUND, KDM4A, KDM5A, MAFK, MAX, MAZ, MBD4, MTA3, MXI1, MYOG, NCOR1, NFIC, NR2F2, NR3C1, NRF1, PHF8, PML, POLR2A, RBBP5, RCOR1, REST, RFX5, RUNX3, SAP30, SIN3A, SMARCB1, SMARCC1, SMC3, SP1, SP2, SP4, STAT1, STAT3, TAF1, TAF7, TBL1XR1, TBP, TCF12, TCF7L2, TEAD4, TFAP2A, TFAP2C, USF1, USF2, WRNIP1, YY1, ZBTB7A, ZEB1, ZKSCAN1, ZNF143, ZNF217, ZNF384 |
| *ELFN2* | TCF4, CREB1, CREM, MITF, TP53, REST, SETDB1, ERG, TFCP2L1, EZH2, RNF2, JARID2, MTF2, SUZ12, RBPJ, BACH1, BRCA1, CEBPB, CHD1, CHD2, CHD7, CTBP2, CTCF, CTCFL, ELF1, EP300, GABPA, GTF2F1, H2AFZ, HDAC1, HDAC2, HDAC6, JUN, JUND, KDM4A, KDM5A, KDM5B, MAFK, MAX, MAZ, MXI1, MYC, MYOG, NFIC, NR3C1, PAX5, POLR2A, RAD21, RCOR1, RFX5, SAP30, SIN3A, SMC3, STAT1, STAT3, TBP, TCF12, TEAD4, YY1, ZBTB7A, ZKSCAN1, ZNF143, ZNF263 |
| *NALCN* | NFE2L2, STAT3, TCF4, SMAD4, TP63, MYC, GATA1, ZFX, REST, ERG, SMARCA4, NR3C1, RUNX2, FOXP2, SUZ12, TFAP2A, POU3F2, RCOR1, PRDM16, ZNF322, BACH1, CEBPB, CHD1, CHD2, CHD7, CREB1, CTBP2, CTCF, E2F4, E2F6, EP300, EZH2, FOS, FOXA2, GATA3, H2AFZ, HCFC1, HDAC2, JUN, JUND, KDM4A, MAFK, MAX, MAZ, PHF8, POLR2A, RAD21, RBBP5, SIN3A, SMARCB1, TCF7L2, TRIM28, YY1, ZKSCAN1 |
| *ANKRD1* | EGR1, SMAD1, SMAD3, AR, TCF4, TP63, NANOG, POU5F1, SOX2, TCF3, FOXA2, HNF4A, GATA1, GATA2, REST, TRIM28, FOXO3, EP300, SMARCA4, SALL4, ATF3, SMAD2, NR3C1, MEF2A, RAD21, CTNNB1, MEIS1, EWSR1, ESR2, ARID3A, BACH1, BCL3, BHLHE40, BRCA1, CBX3, CEBPB, CEBPD, CHD1, CHD2, CHD7, CREB1, CTCF, E2F4, ELF1, ESR1, ETS1, EZH2, FOS, FOSL1, FOSL2, FOXA1, FOXM1, GABPA, GATA3, GTF2F1, H2AFZ, HDAC2, HNF4G, JUN, JUND, MAFF, MAFK, MAX, MAZ, MBD4, MXI1, MYBL2, MYC, MYOD1, MYOG, NFIC, NR2F2, PBX3, POLR2A, RBBP5, RCOR1, RFX5, RXRA, SAP30, SIN3A, SIX5, SMC3, SP1, SRF, STAT3, SUPT20H, TAF1, TBP, TCF12, TCF7L2, TEAD4, TFAP2A, TFAP2C, USF1, USF2, YY1, ZBTB33, ZNF217, ZNF384 |
| *EPHA6* | ZNF217, STAT3, TCF4, SMAD4, TP63, MITF, PPARG, SIN3B, TFAP2C, PPARD, SOX9, YAP1, ZNF281, RUNX2, FOXP2, EZH2, RNF2, JARID2, MTF2, SUZ12, GBX2, CDKN2AIP, BACH1, BCLAF1, BHLHE40, BRCA1, CEBPB, CHD1, CHD2, CHD7, CREB1, CTBP2, CTCF, E2F4, E2F6, ELF1, ELK1, EP300, FOS, FOXM1, GATA3, GTF2F1, H2AFZ, HDAC2, IRF4, JUND, KDM4A, KDM5A, MAFK, MAX, MAZ, MXI1, MYC, NRF1, PAX5, PHF8, POLR2A, RAD21, RBBP5, RCOR1, RELA, RFX5, RUNX3, SAP30, SIN3A, SIRT6, SMC3, SPI1, STAT1, TAF1, TBL1XR1, TBP, TCF12, TCF7L2, TEAD4, TRIM28, USF2, WRNIP1, YY1, ZBTB7A, ZEB1, ZKSCAN1, ZNF143, ZNF263, ZNF384 |
| *KCNQ1* | AR, TCF4, TP63, FLI1, RUNX1, SPI1, KLF4, SOX2, EGR1, MITF, TP53, FOXA2, HNF4A, GATA1, GATA2, KDM5B, CNOT3, TRIM28, SOX9, SRY, EP300, YAP1, SCLY, TCF7, EZH2, RNF2, SUZ12, TFAP2A, DROSHA, CHD1, CTCF, E2F6, EBF1, ETS1, GABPA, H2AFZ, HDAC2, HDAC6, IRF1, KDM4A, MAX, MAZ, MYB, NFIC, PHF8, POLR2A, SAP30, SETDB1, SIN3A, STAT3, TAF1, TAL1, TCF12, WHSC1, YY1, ZBTB7A, ZNF143 |
| *SLC16A14* | E2F1, AHR, ARNT, HOXC9, AR, RUNX1, HNF4A, RCOR3, REST, SETDB1, FOXP1, PPARD, TAL1, CNOT3, TRIM28, SOX9, YAP1, SALL4, TEAD4, RUNX2, YY1, TCF7, CTCF, BMI1, MTF2, SUZ12, ESRRB, LMO2, LYL1, ELF5, EWSR1, STAT5A, ELF1, STAT1, ARID3A, ATF2, BACH1, BATF, BCL3, BCLAF1, BHLHE40, CEBPB, CHD1, CHD2, CHD7, CREB1, CTBP2, E2F4, E2F6, EBF1, EP300, EZH2, FOXA1, FOXA2, FOXM1, GABPA, GATA3, GTF2F1, H2AFZ, HCFC1, HDAC1, HDAC2, IRF4, JUND, KDM4A, KDM5B, MAFK, MAX, MAZ, MTA3, MXI1, MYC, NELFE, NFATC1, NFIC, PAX5, PHF8, PML, POLR2A, POU2F2, RAD21, RBBP5, RCOR1, RFX5, RUNX3, SIN3A, SMC3, SP1, SPI1, SRF, TAF1, TBL1XR1, TBP, TCF12, TCF3, TCF7L2, UBTF, WRNIP1, ZBTB7A, ZEB1, ZNF143, ZNF263, ZNF384 |
| *AOC1* | REST |
| *TMEM130* | TP63, RUNX1, NANOG, EGR1, TP53, PPARG, GATA1, REST, SETDB1, RAD21, SUZ12, TBX3, JUN, SREBF2, ATF3, BACH1, BATF, BCL11A, BHLHE40, CHD1, CHD2, CHD7, CTCF, E2F6, EBF1, EP300, EZH2, FOS, FOSL2, H2AFZ, HDAC1, HDAC2, HMGN3, JUND, KDM4A, KDM5B, MAFK, MAX, MAZ, MXI1, MYC, NRF1, PAX5, PHF8, POLR2A, RBBP5, RCOR1, SAP30, SIN3A, SPI1, STAT3, TAF1, TBL1XR1, TCF12, UBTF, USF2, YY1, ZBTB7A, ZNF143, ZNF263, ZNF384 |
| *RP1L1* | AR, TP63, HNF4A, PPARD, SOX9, SMAD2, SMAD3, SCLY, ESRRB, ARID3A, ATF3, BACH1, BCL3, BHLHE40, BRCA1, CCNT2, CEBPB, CHD2, CTCF, E2F4, ELK1, EP300, EZH2, FOS, FOSL1, FOSL2, FOXA1, FOXA2, GABPA, GATA2, GATA3, GTF2F1, H2AFZ, HCFC1, HDAC1, HDAC2, HDAC6, JUND, KDM5B, MAFF, MAFK, MAX, MAZ, MXI1, MYC, PBX3, PHF8, POLR2A, RAD21, RBBP5, RCOR1, REST, RFX5, RNF2, SAP30, SMARCC1, SMC3, SP1, STAT3, SUZ12, TAF1, TAL1, TBL1XR1, TBP, TCF12, TCF7L2, TEAD4, TFAP2A, TFAP2C, TRIM28, UBTF, USF2, WHSC1, ZKSCAN1, ZMIZ1, ZNF143, ZNF263 |
| *SAA2* | CEBPB, E2F1, NFKB1, RELA, MYC, POU5F1, SOX2, EGR1, ASH2L, ESR1, RCOR1, CEBPA, ATF2, BATF, BCL11A, BCL3, BCLAF1, BHLHE40, CBX2, CHD1, CTCF, EBF1, EP300, ETS1, EZH2, FOS, FOXM1, H2AFZ, IKZF1, IRF4, JUND, MAFK, MAX, MTA3, MYOG, NFIC, PAX5, PBX3, POLR2A, POU2F2, RUNX3, SP1, SPI1, STAT3, TCF12, TCF3, ZEB1, ZNF143, ZNF384 |
| *PIWIL2* | TCF4, SMAD4, HNF4A, ZFX, SOX9, SMARCA4, ATF3, PBX1, PRDM5, STAT1, ARID3A, CEBPB, CHD2, CHD7, CTCF, E2F6, EP300, EZH2, FOS, FOSL2, FOXA1, FOXA2, GABPA, GATA2, GATA3, H2AFZ, HDAC2, JUN, JUND, KDM4A, MAFF, MAFK, MAX, MBD4, MYBL2, MYC, NFIC, NFYA, NFYB, PBX3, PHF8, POLR2A, RAD21, RBBP5, RCOR1, REST, RFX5, RXRA, SIN3A, SMC3, SP1, SP2, SPI1, STAT3, TCF12, TEAD4, USF2, ZBTB33 |

| | |
|---|---|
| *TMEM169* | HOXC9, VDR, STAT3, TCF4, MYC, SPI1, HNF4A, MYCN, RUNX2, TBX5, SIN3A, ARID3A, ATF3, BHLHE40, BRCA1, CCNT2, CEBPD, CHD1, CHD2, CHD7, CREB1, CTCF, E2F4, E2F6, ELF1, EP300, EZH2, FOXA1, FOXA2, GATA1, GATA3, GTF2B, GTF2F1, H2AFZ, HCFC1, HDAC1, HDAC2, HDAC6, HMGN3, JUN, JUND, KAT2A, KDM4A, KDM5A, KDM5B, MAFK, MAX, MAZ, MBD4, MTA3, MXI1, MYBL2, NELFE, NFIC, PAX5, PHF8, PML, POLR2A, POU2F2, RAD21, RBBP5, RCOR1, REST, RFX5, RNF2, RUNX3, SAP30, STAT1, TAF1, TAF7, TBL1XR1, TBP, TCF12, TCF3, TCF7L2, TEAD4, UBTF, USF1, USF2, WRNIP1, YY1, ZEB1, ZMIZ1, ZNF143, ZNF384 |
| *HTRA3* | TCF4, TP63, RUNX1, SPI1, TCF3, EGR1, MITF, PPARG, GATA1, DMRT1, TEAD4, RUNX2, MTF2, SUZ12, JUN, MEIS1, HSF1, NFIB, DROSHA, BHLHE40, CHD1, CREB1, CTBP2, CTCF, E2F6, EP300, EZH2, GABPA, H2AFZ, HDAC2, JUND, KDM4A, KDM5A, KDM5B, MAX, MAZ, MYC, MYOG, PHF8, POLR2A, RAD21, RFX5, SIN3A, SMC3, TAF1, TBP, USF1, YY1, ZNF143, ZNF263 |
| *UNC5C* | CEBPD, HOXC9, TTF2, STAT3, TCF4, SMAD4, SPI1, NANOG, POU5F1, SOX2, TCF3, HNF4A, REST, TRIM28, EP300, SMARCA4, YAP1, WT1, NR3C1, EED, PHC1, EZH2, RNF2, JARID2, MTF2, SUZ12, GATA3, ATF2, BATF, BCL11A, BCLAF1, BHLHE40, CEBPB, CHD1, CHD2, CHD7, CREB1, CTBP2, CTCF, CUX1, E2F4, EBF1, FOXA2, FOXP2, GATA2, H2AFZ, HDAC1, HDAC2, HDAC6, IKZF1, IRF3, IRF4, KDM4A, MAX, MAZ, MXI1, MYC, MYOG, NRF1, PAX5, PBX3, PHF8, POLR2A, POU2F2, RAD21, RBBP5, RCOR1, RFX5, RUNX3, SAP30, SIN3A, STAT1, TAF1, TBL1XR1, TBP, TCF12, TCF7L2, TEAD4, WRNIP1, YY1, ZNF143, ZNF263, ZNF384 |
| *THEM6* | VDR, DACH1, PHF8, CREB1, E2F1, MYC, SPI1, KLF4, MITF, GATA1, ZFX, SIN3B, FOXP1, ERG, CNOT3, TRIM28, SRY, TEAD4, CTCF, SUZ12, EWSR1, HSF1, DCP1A, ARID3A, ATF2, ATF3, BACH1, BCL3, BCLAF1, BHLHE40, BRCA1, CBX3, CCNT2, CEBPB, CEBPD, CHD1, CHD2, CHD7, CTCFL, CUX1, E2F4, E2F6, EBF1, EGR1, ELF1, ELK1, ELK4, EP300, ETS1, EZH2, FOS, FOSL1, FOSL2, FOXA1, FOXA2, FOXP2, GABPA, GATA3, GTF2F1, H2AFZ, HCFC1, HDAC1, HDAC2, HDAC6, HMGN3, HNF4A, IRF1, JUN, JUND, KAT2A, KDM1A, KDM4A, KDM5B, MAFF, MAFK, MAX, MAZ, MTA3, MXI1, MYBL2, MYOG, NELFE, NFIC, NFYA, NFYB, NR2F2, NRF1, PAX5, PML, POLR2A, POU2F2, RAD21, RBBP5, RCOR1, RELA, REST, RFX5, RNF2, RUNX3, SAP30, SETDB1, SIN3A, SIX5, SMARCB1, SMC3, SP1, STAT1, STAT3, STAT5A, SUPT20H, TAF1, TAL1, TBL1XR1, TBP, TCF12, TCF3, TCF7L2, TFAP2A, TFAP2C, THAP1, UBTF, USF1, USF2, WRNIP1, YY1, ZBTB33, ZBTB7A, ZC3H11A, ZEB1, ZKSCAN1, ZMIZ1, ZNF143, ZNF263, ZNF384 |
| *WNT7B* | NFE2L2, AR, STAT3, TP63, RUNX1, MYC, KLF4, NANOG, SOX2, EGR1, MITF, TP53, RCOR3, SIN3B, REST, SETDB1, FOXP1, PPARD, TET1, SOX9, SRY, EP300, OLIG2, YAP1, DMRT1, KLF1, NR3C1, SCLY, CCND1, FOXP2, EZH2, RNF2, JARID2, MTF2, SUZ12, BACH1, HSF1, GATA3, ESR2, DROSHA, ZNF322, BCL3, BHLHE40, BRCA1, CBX3, CCNT2, CEBPB, CHD2, CREB1, CTBP2, CTCF, E2F4, E2F6, ELF1, ELK1, FOXA2, GABPA, GTF2F1, H2AFZ, HCFC1, HDAC1, HDAC2, HDAC6, HMGN3, JUND, KDM4A, KDM5B, MAFK, MAX, MAZ, MXI1, MYOD1, MYOG, NELFE, NFIC, NRF1, PAX5, PHF8, POLR2A, RAD21, RBBP5, RCOR1, RFX5, RUNX3, SAP30, SIN3A, SMC3, SP1, STAT1, TAF1, TBL1XR1, TBP, TCF12, TCF3, TCF7L2, TEAD4, UBTF, USF1, YY1, ZBTB7A, ZC3H11A, ZEB1, ZKSCAN1, ZNF143 |
| *HOXB6* | STAT3, E2F1, KLF4, NANOG, POU5F1, TCF3, EGR1, TP53, KDM5B, SETDB1, TFAP2C, TET1, WT1, ZNF281, ZFP42, BMI1, EED, PHC1, EZH2, RNF2, JARID2, MTF2, SUZ12, TFAP2A, BACH1, HSF1, CDX2, ARID3A, ATF3, BCL3, BHLHE40, BRCA1, CBX3, CEBPB, CHD1, CHD2, CHD4, CHD7, CREB1, CTBP2, CTCF, E2F6, EBF1, ELF1, ELK1, ELK4, EP300, ETS1, FOSL2, FOXA1, FOXM1, GABPA, GATA2, GATA3, GTF2B, GTF2F1, H2AFZ, HCFC1, HDAC1, HDAC2, HMGN3, IRF1, JUND, KDM1A, KDM5A, MAFF, MAFK, MAX, MAZ, MXI1, MYC, NFIC, NFYA, NFYB, NR3C1, PHF8, POLR2A, PRDM1, RAD21, RBBP5, RCOR1, REST, RFX5, SIN3A, SIX5, SMC3, SP1, TAF1, TAL1, TBL1XR1, TBP, TCF12, TCF7L2, TEAD4, UBTF, USF1, USF2, WHSC1, YY1, ZBTB33, ZC3H11A, ZKSCAN1, ZMIZ1, ZNF143, ZNF263 |
| *UCP3* | AR, MYC, PPARA, PPARD, SPI1, TP53, PPARG, FOXA2, HNF4A, GATA1, GATA2, SOX9, EP300, SMARCA4, PRDM14, MYBL2, STAT5A, BCL3, ARID3A, BHLHE40, CBX3, CCNT2, CEBPB, CHD1, CHD2, CHD7, CTCF, CUX1, E2F6, ELF1, EZH2, FOXA1, H2AFZ, HDAC1, HDAC2, JUN, JUND, KDM1A, KDM5A, MAFF, MAFK, MAX, MAZ, MYOD1, MYOG, NR2F2, POLR2A, RAD21, RBBP5, RCOR1, REST, SAP30, SIRT6, SMC3, TAL1, TBL1XR1, TEAD4, UBTF, USF1, USF2, YY1, ZC3H11A, ZMIZ1, ZNF143, ZNF384 |
| *KRT81* | AHR, ARNT, ELK1, AR, SOX9, EWSR1, RBPJ, DNAJC2, CEBPB, CTCF, EBF1, EP300, EZH2, GATA1, H2AFZ, MAFK, POLR2A, RAD21, REST, TCF12, ZKSCAN1 |
| *KLHDC7A* | AR, SMAD4, E2F1, NANOG, POU5F1, SOX2, TCF3, TP53, PPARG, HNF4A, GATA2, RCOR3, SIN3B, REST, TET1, SRY, FOXO3, DMRT1, RUNX2, ESRRB, TBX3, EWSR1, STAT5A, NR0B1, ARID3A, ATF1, ATF2, CEBPB, CEBPD, CHD2, CTBP2, CTCF, E2F6, ELF1, EP300, ESR1, EZH2, FOS, FOSL2, FOXA1, FOXM1, GATA1, GATA3, GTF2F1, H2AFZ, HDAC1, HDAC2, HDAC6, HMGN3, HNF4G, IRF1, JUN, JUND, KDM5B, MAFK, MAX, MAZ, MXI1, MYC, NFIC, NR3C1, PAX5, PHF8, POLR2A, RAD21, RBBP5, RCOR1, RFX5, SAP30, SIN3A, SP1, SP4, SRF, STAT3, SUZ12, TAF1, TAF7, TAL1, TBL1XR1, TBP, TCF12, TEAD4, UBTF, USF1, YY1, ZBTB7A, ZMIZ1, ZNF143 |
| *NTRK2* | ZNF217, AR, STAT3, SMAD4, TP63, RUNX1, MYC, NANOG, POU5F1, SOX2, MITF, TP53, RCOR3, REST, ASH2L, TFAP2C, ERG, TRIM28, SRY, SMARCA4, YAP1, DMRT1, KLF1, NR3C1, RUNX2, MYBL2, FOXP2, CTCF, EED, PHC1, EZH2, RNF2, JARID2, MTF2, SUZ12, TBX3, JUN, BACH1, SOX11, TCF7L2, DNAJC2, SREBF1, BHLHE40, CEBPB, CHD1, CHD2, CHD7, CTBP2, CTCFL, CUX1, E2F6, EBF1, ELK1, EP300, GABPA, GTF2F1, H2AFZ, HDAC2, HDAC6, JUND, KDM4A, KDM5A, MAX, MAZ, MXI1, MYOG, NRF1, PAX5, PHF8, POLR2A, RAD21, RBBP5, RUNX3, SAP30, SIN3A, SMC3, SPI1, SRF, STAT1, TAF1, TAF7, TBP, TCF12, TCF3, TEAD4, USF2, WRNIP1, YY1, ZEB1, ZNF143, ZNF263 |

| | |
|---|---|
| *RAB17* | AR, SMAD4, FLI1, MYC, SPI1, NANOG, SOX2, MITF, PPARG, HNF4A, GATA2, EOMES, TFAP2C, FOXP1, GFI1B, TRIM28, SALL4, SOX17, ZNF281, TFCP2L1, SMAD3, FOXP2, RAD21, BMI1, JUN, BACH1, CEBPB, GATA3, ARID3A, BHLHE40, BRCA1, CBX2, CEBPD, CHD1, CHD2, CTCF, E2F4, E2F6, EGR1, ELF1, EP300, ESRRA, EZH2, FOS, FOSL2, FOXA1, FOXA2, GABPA, GATA1, GTF2F1, H2AFZ, HDAC2, HNF4G, JUND, KDM4A, KDM5A, MAFF, MAFK, MAX, MAZ, MBD4, MXI1, MYBL2, MYOG, NFIC, NR2C2, NR2F2, NR3C1, PML, POLR2A, RCOR1, REST, RFX5, RXRA, SAP30, SIN3A, SMC3, SP1, STAT3, TAF1, TAF7, TBP, TCF12, TCF7L2, TEAD4, USF1, USF2, YY1, ZBTB7A, ZEB1, ZKSCAN1, ZNF143, ZNF217 |
| *CDH23* | ELK1, AR, SMAD4, CREB1, FLI1, RUNX1, MYC, SPI1, SOX2, EGR1, MITF, TP53, FOXA2, HNF4A, GATA1, GATA2, SETDB1, SOX9, SRY, ZNF281, DMRT1, TEAD4, SMAD3, SCLY, CTCF, RAD21, JARID2, LMO2, RELA, CEBPB, RBPJ, NR1H3, BCL11B, BACH1, BHLHE40, CCNT2, CHD1, CHD2, CHD7, CTBP2, E2F6, ELF1, EP300, ETS1, EZH2, GABPA, GTF2F1, H2AFZ, HDAC1, HDAC2, HDAC6, HMGN3, JUND, KAT2A, KAT2B, KDM4A, KDM5B, MAX, MAZ, MXI1, NANOG, PHF8, POLR2A, RBBP5, RCOR1, REST, SAP30, SIN3A, SIRT6, SMC3, SUZ12, TBP, TCF12, TCF7L2, UBTF, USF2, YY1, ZBTB7A, ZC3H11A, ZNF143, ZNF263 |
| *CGB5* | TP63, CUX1, CTCF, EBF1, EZH2, FOS, FOSL2, H2AFZ, JUND, MAX, NFIC, NR3C1, PBX3, POLR2A, POU2F2, REST, TCF12, TCF3, ZBTB33 |
| *SEMA3D* | ZNF217, PAX3, AR, STAT3, RUNX1, MYC, NANOG, POU5F1, SOX2, PPARG, GATA2, RCOR3, PPARD, SMARCA4, PRDM14, RUNX2, YY1, CTCF, BMI1, EED, JARID2, MTF2, SUZ12, POU3F2, CEBPB, E2F4, H2AFZ, MYOD1, MYOG, POLR2A, TCF12, TCF3 |
| *IGFBP6* | CREB1, E2F1, EGR1, HNF4A, GATA1, EOMES, TET1, TRIM28, SALL4, WT1, ATF3, MYBL2, STAT4, MTF2, SUZ12, NR1H3, ARID3A, ATF2, BACH1, BATF, BCL3, BHLHE40, BRCA1, CBX3, CCNT2, CEBPB, CHD1, CHD2, CHD7, CTBP2, CTCF, E2F4, E2F6, EBF1, ELF1, ELK1, ELK4, EP300, ETS1, EZH2, FOS, FOSL1, FOSL2, FOXA1, FOXM1, FOXP2, GABPA, GATA2, GATA3, GTF2F1, H2AFZ, HCFC1, HDAC1, HDAC2, HDAC6, HMGN3, JUN, JUND, KDM1A, KDM4A, KDM5B, MAFF, MAFK, MAX, MAZ, MXI1, MYC, MYOD1, MYOG, NFIC, NR2F2, NR3C1, PBX3, PHF8, POLR2A, RAD21, RBBP5, RCOR1, REST, RFX5, RUNX3, RXRA, SAP30, SIN3A, SIRT6, SIX5, SMARCB1, SMARCC1, SMARCC2, SMC3, SP1, SP4, SRF, STAT1, STAT3, STAT5A, TAF1, TAL1, TBL1XR1, TBP, TCF12, TCF7L2, TEAD4, UBTF, USF1, USF2, WRNIP1, YY1, ZBTB7A, ZC3H11A, ZKSCAN1, ZMIZ1, ZNF143, ZNF263, ZNF384 |
| *RGPD2* | BHLHE40, CEBPB, FOSL2, FOXA1, FOXA2, JUND, MYBL2, POLR2A, POU2F2, RXRA, SIN3A, SP1, TAF1, TCF12, TCF3, TEAD4, USF1, ZBTB33 |
| *HOXA4* | AR, RARA, TFAP2A, KLF4, POU5F1, TP53, PPARG, TET1, ERG, WT1, SMAD2, SMAD3, YY1, BMI1, EED, PHC1, RNF2, JARID2, MTF2, SUZ12, SOX11, CDX2, CHD7, ARID3A, ATF3, BACH1, BHLHE40, BRCA1, CEBPD, CHD1, CHD2, CREB1, CTBP2, CTCF, CUX1, E2F4, E2F6, ELF1, EP300, EZH2, FOS, FOSL2, GABPA, GTF2F1, H2AFZ, HDAC2, HNF4A, HNF4G, JUND, KDM4A, KDM5A, MAFK, MAX, MAZ, MXI1, MYBL2, MYC, NANOG, NFIC, NFYA, NFYB, NR3C1, PML, POLR2A, RAD21, RBBP5, RCOR1, REST, RFX5, SAP30, SIN3A, SMC3, SP1, STAT3, TAF1, TBP, TCF12, TCF7L2, TEAD4, USF1, USF2, ZBTB7A, ZKSCAN1, ZNF143, ZNF263 |
| *RCSD1* | PAX3, VDR, TCF4, SMAD4, TP63, RUNX1, SPI1, EGR1, PPARG, FOXA2, GATA1, GATA2, TFAP2C, TAL1, EP300, WT1, DMRT1, TEAD4, SMAD2, SMAD3, MECOM, TCF7, EZH2, RNF2, MTF2, SUZ12, ESRRB, SOX11, GATA3, ELF1, ARID3A, ATF2, BACH1, BATF, BCL3, BCLAF1, BHLHE40, BRCA1, CBX3, CCNT2, CEBPB, CEBPD, CHD1, CHD2, CHD4, CHD7, CTBP2, CTCF, CUX1, E2F6, EBF1, ELK1, ETS1, FOXM1, GTF2B, GTF2F1, H2AFZ, HCFC1, HDAC1, HDAC2, HDAC6, HMGN3, IKZF1, IRF1, JUN, JUND, KAT2A, KAT2B, KDM1A, KDM4A, KDM5A, KDM5B, MAFF, MAFK, MAX, MAZ, MEF2A, MTA3, MXI1, MYB, MYC, NELFE, NFATC1, NFE2, NFIC, NR2F2, NRF1, PAX5, PHF8, PML, POLR2A, POU2F2, RAD21, RBBP5, RCOR1, RELA, REST, RFX5, RUNX3, SAP30, SIN3A, SMC3, SP1, SRF, STAT1, STAT3, STAT5A, TAF1, TBL1XR1, TBP, TCF12, TCF3, TCF7L2, THAP1, TRIM28, UBTF, USF2, WRNIP1, YY1, ZBTB7A, ZC3H11A, ZEB1, ZMIZ1, ZNF143, ZNF263, ZNF384 |
| *ULBP2* | NFE2L2, HOXC9, AR, TP63, E2F1, FLI1, FOXA2, EWSR1, RBPJ, ATF1, BACH1, BHLHE40, CBX3, CCNT2, CHD1, CHD2, CHD4, CHD7, CREB1, CTBP2, CTCF, E2F4, E2F6, ELF1, ELK1, EP300, ETS1, EZH2, FOS, FOXP2, GABPA, GTF2F1, H2AFZ, HDAC1, HDAC2, HDAC6, HMGN3, IRF1, JUN, JUND, KDM4A, KDM5B, MAFK, MAX, MAZ, MXI1, MYC, NFYA, NFYB, NR2F2, PAX5, PBX3, PHF8, POLR2A, RAD21, RBBP5, RCOR1, REST, RFX5, RNF2, SAP30, SIN3A, SIRT6, SP1, SP2, SP4, STAT1, STAT3, SUZ12, TAF1, TBL1XR1, TBP, TCF12, UBTF, USF1, USF2, YY1, ZBTB7A, ZC3H11A, ZMIZ1, ZNF143, ZNF263 |
| *LIMS2* | SOX2, EGR1, PPARG, TET1, ESR1, SOX17, SUZ12, IRF1, THRA, BHLHE40, CEBPB, CTCF, EP300, EZH2, GATA2, H2AFZ, HDAC6, KDM5B, MXI1, MYC, MYOG, POLR2A, RAD21, RBBP5, SAP30 |
| *STAT5A* | ESR1, ESR2, STAT3, CREB1, FLI1, RUNX1, SOX2, EGR1, MITF, PPARG, HNF4A, GATA1, GATA2, RCOR3, SETDB1, TFAP2C, TET1, ERG, MECOM, STAT4, MTF2, SUZ12, SIN3A, NUCKS1, RELA, TBP, NR0B1, TAF7L, STAT6, FOXO1, BACH1, BCL3, BCLAF1, BHLHE40, BRCA1, CBX3, CCNT2, CEBPB, CHD1, CHD2, CHD7, CTCF, E2F4, E2F6, EBF1, ELF1, ELK1, EP300, ETS1, EZH2, FOXM1, FOXP2, GABPA, GTF2B, GTF2F1, H2AFZ, HCFC1, HDAC1, HDAC2, HDAC6, HMGN3, IRF1, JUND, KAT2A, KDM1A, KDM4A, KDM5B, MAFF, MAFK, MAX, MAZ, MTA3, MXI1, MYC, MYOD1, MYOG, NELFE, NFATC1, NFIC, NFYA, NFYB, NR2F2, NRF1, PAX5, PHF8, PML, POLR2A, POU2F2, RAD21, RBBP5, RCOR1, REST, RFX5, RUNX3, SAP30, SMARCB1, SMARCC1, SMC3, SP1, SPI1, STAT1, STAT5A, TAF1, TBL1XR1, TCF12, TCF3, TCF7L2, TEAD4, UBTF, USF1, USF2, WRNIP1, YY1, ZBTB7A, ZC3H11A, ZKSCAN1, ZMIZ1, ZNF143, ZNF384 |
| *PALM3* | RUNX1, POU5F1, SOX2, TP53, RCOR3, TFAP2C, TET1, TEAD4, TFCP2L1, MTF2, ESRRB, ARID3A, BACH1, BHLHE40, CBX3, CCNT2, CEBPB, CEBPD, CHD1, CHD2, CHD4, CHD7, CREB1, CREBBP, CTCF, CTCFL, CUX1, E2F6, EGR1, ELF1, EP300, EZH2, FOXA2, GABPA, GATA1, GATA2, GATA3, H2AFZ, HCFC1, HDAC1, HDAC2, HDAC6, HMGN3, HNF4A, HNF4G, JUND, KDM1A, KDM4A, KDM5A, KDM5B, MAFF, MAFK, MAX, MAZ, MXI1, MYBL2, MYC, MYOD1, MYOG, NFIC, NRF1, PHF8, POLR2A, RAD21, RBBP5, RCOR1, RFX5, RNF2, RUNX3, RXRA, SAP30, SETDB1, SIN3A, SIRT6, SMC3, SP1, SP4, SRF, STAT3, TAL1, TBL1XR1, TBP, TCF12, TCF7L2, UBTF, USF1, USF2, YY1, ZBTB7A, ZC3H11A, ZNF143, ZNF263, ZNF384 |

| | |
|---|---|
| *DLX4* | TTF2, TCF4, TP63, MYC, KLF4, NANOG, POU5F1, SOX2, EGR1, TP53, HNF4A, GATA2, REST, YAP1, ATF3, TBX5, BMI1, EZH2, RNF2, JARID2, MTF2, SUZ12, ESRRB, CTNNB1, RCOR1, GATA3, DNAJC2, ATF2, BACH1, BCL3, BHLHE40, BRCA1, CBX3, CCNT2, CEBPB, CEBPD, CHD1, CHD2, CHD7, CREB1, CTBP2, CTCF, E2F1, E2F4, E2F6, EBF1, ELF1, ELK1, ELK4, EP300, FOS, FOSL1, FOSL2, FOXA1, FOXA2, FOXP2, GABPA, GTF2F1, H2AFZ, HCFC1, HDAC1, HDAC2, HDAC6, HMGN3, IRF1, JUN, JUND, KDM4A, KDM5A, KDM5B, MAFF, MAFK, MAX, MAZ, MXI1, MYBL2, MYOG, NELFE, NFIC, NFYA, NFYB, NR2F2, NRF1, PAX5, PHF8, PML, POLR2A, RAD21, RBBP5, RFX5, RUNX3, SAP30, SIN3A, SIRT6, SMARCB1, SMARCC1, SMC3, SP1, SP4, TAF1, TBP, TCF12, TCF3, TCF7L2, TEAD4, TFAP2A, TFAP2C, THAP1, TRIM28, UBTF, USF1, USF2, WHSC1, YY1, ZBTB33, ZBTB7A, ZKSCAN1, ZNF143, ZNF263 |
| *DMRT2* | ZNF217, PAX3, STAT3, SMAD4, E2F1, MYC, KLF4, TP53, PPARD, SMARCA4, MYBL2, BMI1, EED, PHC1, EZH2, RNF2, JARID2, MTF2, SUZ12, BACH1, BHLHE40, CHD1, CHD2, CREB1, CTCF, E2F4, E2F6, EBF1, EGR1, ELF1, ESR1, GABPA, GATA3, GTF2F1, H2AFZ, HDAC2, KDM4A, KDM5A, MAX, MAZ, MXI1, MYOG, NFIC, NFYB, NR2F2, PHF8, POLR2A, RAD21, RBBP5, RCOR1, RFX5, SAP30, SETDB1, SIN3A, SP1, TAF1, TBP, TCF3, TCF7L2, YY1, ZNF143 |
| *ASB2* | AR, RARA, ARNT, NFE2L2, TP63, MYC, PPARG, RCOR3, TFAP2C, ATF3, SMAD2, SMAD3, STAT4, FOXP2, ATF2, BATF, BCL11A, BCL3, BCLAF1, BHLHE40, BRCA1, CEBPB, CHD1, CHD2, CREB1, CTCF, CUX1, E2F4, EBF1, EGR1, ELF1, ELK1, EP300, EZH2, FOXM1, H2AFZ, IKZF1, IRF3, IRF4, JUND, KDM4A, KDM5A, KDM5B, MAX, MAZ, MEF2A, MEF2C, MTA3, MXI1, MYOD1, MYOG, NFATC1, NFE2, NFIC, NRF1, PAX5, PML, POLR2A, POU2F2, RAD21, RBBP5, RCOR1, RELA, RFX5, RUNX3, RXRA, SIN3A, SMC3, SP1, SPI1, SRF, STAT1, STAT3, STAT5A, TAF1, TBL1XR1, TBP, TCF12, TCF3, USF1, USF2, WRNIP1, YY1, ZEB1, ZNF143, ZNF384 |
| *HIST1H2BO* | VDR, DACH1, PHF8, CREB1, E2F1, MYC, NANOG, POU5F1, SOX2, TCF3, EGR1, TP53, E2F4, PPARD, CNOT3, SALL4, DMRT1, ATF3, HOXB4, MYBL2, RNF2, RCOR1, ZIC3, SMAD1, PRDM5, ELF1, PAX6, IRF8, ARID3A, ATF1, ATF2, BACH1, BATF, BCL11A, BCL3, BCLAF1, BHLHE40, BRCA1, CBX3, CCNT2, CEBPB, CEBPD, CHD1, CHD2, CHD7, CREBBP, CTCF, CTCFL, CUX1, E2F6, EBF1, ELK1, ELK4, EP300, ETS1, EZH2, FLI1, FOS, FOSL1, FOSL2, FOXA1, FOXA2, FOXM1, GABPA, GATA1, GATA2, GATA3, GTF2B, GTF2F1, GTF3C2, H2AFZ, HCFC1, HDAC1, HDAC2, HDAC6, HMGN3, HNF4G, IRF1, IRF3, IRF4, JUN, JUND, KAT2A, KAT2B, KDM1A, KDM4A, KDM5A, KDM5B, MAFF, MAFK, MAX, MAZ, MBD4, MEF2A, MEF2C, MTA3, MXI1, NCOR1, NELFE, NFATC1, NFE2, NFIC, NFYA, NFYB, NR2F2, NRF1, PAX5, PBX3, PML, POLR2A, POU2F2, PRDM1, RAD21, RBBP5, RELA, REST, RFX5, RUNX3, RXRA, SAP30, SETDB1, SIN3A, SIRT6, SIX5, SMC3, SP1, SP2, SP4, SPI1, SRF, STAT1, STAT3, STAT5A, TAF1, TAF7, TBL1XR1, TBP, TCF12, TCF7L2, TEAD4, TRIM28, UBTF, USF1, USF2, WRNIP1, YY1, ZBTB33, ZBTB7A, ZC3H11A, ZEB1, ZKSCAN1, ZMIZ1, ZNF143, ZNF274, ZNF384 |
| *PACSIN1* | RUNX1, POU5F1, EGR1, MITF, TP53, RCOR3, SIN3B, REST, SETDB1, TFAP2C, FOXP1, TRIM28, SOX9, SRY, ZNF281, SMAD3, MEF2A, TBX5, MTF2, SUZ12, SIN3A, PAX6, NR1H3, ESR2, ATF3, BHLHE40, BRCA1, CBX3, CCNT2, CHD1, CHD2, CREB1, CTCF, E2F6, ELF1, EP300, ETS1, EZH2, FOXA1, GATA1, GATA3, H2AFZ, HCFC1, HDAC1, HDAC2, HMGN3, JUN, JUND, KAT2A, KDM1A, KDM4A, KDM5A, KDM5B, MAFK, MAX, MAZ, MXI1, MYC, MYOD1, MYOG, NELFE, NRF1, PHF8, POLR2A, RAD21, RBBP5, RCOR1, SAP30, SMC3, TAF1, TBL1XR1, TBP, UBTF, USF2, YY1, ZBTB7A, ZNF143, ZNF263, ZNF384 |
| *GLRA3* | AR, STAT3, TCF4, TP63, E2F1, FLI1, MYC, KDM5B, RCOR3, SIN3B, REST, SRY, SALL4, NR3C1, CUX1, PBX1, POU3F2, RCOR1, PAX6, BACH1, CBX3, CEBPB, CHD1, CHD2, CHD7, CTBP2, CTCF, CTCFL, E2F4, E2F6, EGR1, ELF1, EP300, EZH2, GABPA, GTF2F1, H2AFZ, HDAC1, HDAC2, HDAC6, HMGN3, JUN, JUND, KDM4A, KDM5A, MAFF, MAFK, MAX, MXI1, NFYA, NFYB, PBX3, PHF8, PML, POLR2A, RAD21, RBBP5, SAP30, SIN3A, SMC3, SP1, SP2, SP4, TAF1, TAF7, TBP, TCF12, TEAD4, USF1, YY1, ZBTB7A, ZNF143 |
| *NKX2-8* | GLI1, AR, TP63, MYC, KLF4, NANOG, POU5F1, TCF3, TP53, HNF4A, SETDB1, ASH2L, SRY, YAP1, SALL4, ZNF281, BMI1, EED, PHC1, EZH2, RNF2, JARID2, MTF2, SUZ12, ZNF274, BACH1, CHD1, CREB1, CTBP2, CTCF, E2F4, E2F6, EGR1, ELF1, EP300, GABPA, H2AFZ, HDAC2, HDAC6, JUN, JUND, KDM4A, KDM5B, MAX, MXI1, MYOG, NR2F2, PHF8, POLR2A, RBBP5, SAP30, SIN3A, TAF1, TBP, TCF12, YY1, ZBTB7A, ZNF143 |
| *LONRF3* | AR, SMAD4, NANOG, POU5F1, SOX2, EGR1, TP53, TRIM28, WT1, NR3C1, RUNX2, MYBL2, EZH2, RNF2, JARID2, MTF2, SUZ12, RCOR1, CEBPB, ARID3A, ATF3, BCL3, BHLHE40, BRCA1, CBX2, CBX3, CBX8, CCNT2, CEBPD, CHD1, CHD2, CHD7, CREB1, CTCF, E2F1, E2F4, E2F6, EBF1, ELF1, ELK1, ELK4, EP300, ETS1, FOS, FOSL1, FOSL2, FOXA1, FOXA2, FOXP2, GABPA, GTF2F1, H2AFZ, HCFC1, HDAC1, HDAC2, HMGN3, HNF4A, HNF4G, IRF1, JUN, JUND, KDM4A, KDM5A, KDM5B, MAFK, MAX, MAZ, MXI1, MYC, NFIC, NFYA, NFYB, NR2F2, PHF8, POLR2A, RAD21, RBBP5, REST, RFX5, RXRA, SAP30, SIN3A, SMARCB1, SMC3, SP1, STAT3, TAF1, TAL1, TBL1XR1, TBP, TCF12, TCF3, TCF7L2, TEAD4, UBTF, USF1, USF2, YY1, ZBTB7A, ZKSCAN1, ZMIZ1, ZNF143, ZNF263, ZNF384 |
| *MAMDC2* | ZNF217, NFE2L2, STAT3, TCF4, TP63, FLI1, NANOG, SOX2, TP53, FOXA2, HNF4A, GATA2, KDM5B, REST, ASH2L, SOX9, ESR1, YAP1, TEAD4, CTNNB1, PBX1, LMO2, ATF2, BACH1, CHD1, CHD2, CHD7, CTBP2, CTCF, E2F4, E2F6, EP300, EZH2, FOS, FOSL2, GTF2F1, H2AFZ, HDAC2, HDAC6, JUN, JUND, KDM4A, MAX, MXI1, MYC, MYOD1, MYOG, PHF8, POLR2A, RAD21, RBBP5, SAP30, SIN3A, SIRT6, SMC3, SP1, SUZ12, TAF1, TBP, TCF12, TCF7L2, YY1, ZNF143 |
| *SPDYE1* | TP63, MYC, FOXP1, SMAD2, SMAD3, CDX2, EP300, FOSL2, GABPA, GATA2, IRF4, JUND, PAX5, PBX3, POLR2A, POU2F2, RXRA, SIX5, SP1, SPI1, TAF1, ZBTB33 |
| *TSPAN15* | ZNF217, AR, STAT3, SMAD4, E2F1, SPI1, KLF4, NANOG, POU5F1, SOX2, TCF3, MITF, FOXA2, ZFX, E2F4, RCOR3, REST, TFAP2C, PPARD, TET1, TAL1, TRIM28, FOXO3, SALL4, TEAD4, SMAD2, SMAD3, TBX5, TCF7, RAD21, EZH2, RELA, NR0B1, ARID3A, ATF3, BACH1, BCL3, BHLHE40, BRCA1, CCNT2, CEBPB, CHD1, CHD2, CHD7, CREB1, CTBP2, CTCF, E2F6, EBF1, EGR1, ELF1, ELK1, EP300, FOS, FOSL1, FOSL2, FOXA1, GATA3, GTF2F1, H2AFZ, HDAC2, HDAC6, HMGN3, IRF1, IRF3, JUND, KAT2B, KDM1A, KDM4A, KDM5A, KDM5B, MAFK, MAX, MAZ, MXI1, MYC, MYOG, NFIC, NFYA, NFYB, NR2F2, |

| | |
|---|---|
| | PAX5, PBX3, PHF8, PML, POLR2A, RBBP5, RCOR1, RFX5, RXRA, SAP30, SIN3A, SIX5, SMARCB1, SMC3, SP1, SP2, SP4, STAT1, SUZ12, TAF1, TBP, TCF12, TCF7L2, UBTF, USF2, YY1, ZBTB33, ZBTB7A, ZEB1, ZKSCAN1, ZNF143 |
| *SARM1* | TP63, TCF3, EGR1, REST, TFAP2C, CNOT3, TRIM28, CCND1, JARID2, SUZ12, PBX1, BACH1, TFEB, ARID3A, ATF1, ATF2, ATF3, BCL3, BCLAF1, BHLHE40, BRCA1, CCNT2, CEBPB, CEBPD, CHD1, CHD2, CHD7, CREB1, CTBP2, CTCF, CUX1, E2F4, E2F6, EBF1, ELF1, ELK1, EP300, EZH2, FOSL2, FOXA1, FOXA2, FOXP2, GABPA, GATA2, GATA3, GTF2F1, H2AFZ, HCFC1, HDAC1, HDAC2, HDAC6, HMGN3, HNF4A, HNF4G, IRF1, JUN, JUND, KDM1A, KDM4A, KDM5A, KDM5B, MAFF, MAFK, MAX, MAZ, MBD4, MTA3, MXI1, MYBL2, MYC, MYOG, NELFE, NFATC1, NFE2, NFIC, NFYA, NFYB, NR2F2, NRF1, PAX5, PHF8, PML, POLR2A, RAD21, RBBP5, RCOR1, RFX5, RUNX3, RXRA, SAP30, SETDB1, SIN3A, SIRT6, SMARCC1, SMC3, SP1, SPI1, SRF, STAT1, STAT3, STAT5A, TAF1, TAL1, TBL1XR1, TBP, TCF12, TCF7L2, TEAD4, TFAP2A, UBTF, USF1, USF2, WHSC1, YY1, ZBTB33, ZBTB7A, ZC3H11A, ZKSCAN1, ZMIZ1, ZNF143, ZNF263, ZNF384 |
| *ZBED6CL* | TTF2, HNF4A, SMAD2, SMAD3, PBX1, ATF2, BACH1, CCNT2, CHD2, CHD7, CTBP2, CTCF, EZH2, GABPA, GTF2F1, H2AFZ, HDAC2, HMGN3, JUND, KDM5B, MAFK, MAX, MXI1, MYC, NANOG, PHF8, POLR2A, RAD21, RBBP5, SAP30, SIN3A, TAF1, TBP, TCF12, USF2, ZNF143 |
| *SLFN14* | FLI1, RUNX1, SPI1, GATA1, GATA2, TAL1, SCLY, ATF1, BHLHE40, CEBPB, CHD2, CUX1, EP300, ETS1, EZH2, H2AFZ, HDAC1, HDAC2, IRF1, JUND, KDM5B, MAFF, MAFK, MAX, MYC, RCOR1, SAP30, TBL1XR1, TEAD4, WHSC1 |
| *ABO* | TP63, RUNX1, GATA2, SETDB1, BACH1, BCL3, BHLHE40, CCNT2, CHD1, CHD2, CHD4, CHD7, CTCF, CTCFL, E2F6, EGR1, ELF1, EP300, ETS1, EZH2, GABPA, GTF2F1, H2AFZ, HCFC1, HDAC1, HDAC2, HDAC6, HMGN3, IRF1, JUND, KDM4A, KDM5A, KDM5B, MAFF, MAFK, MAX, MAZ, MYC, PHF8, POLR2A, RAD21, RBBP5, RCOR1, REST, RNF2, SAP30, SIRT6, SMC3, STAT3, SUZ12, TAF1, TAL1, TBL1XR1, TBP, UBTF, USF2, YY1, ZBTB7A, ZC3H11A, ZMIZ1, ZNF143, ZNF384 |
| *SEC14L4* | AR, E2F1, SOX2, HNF4A, ZFX, RCOR3, TET1, SRY, ESR1, ARID3A, BHLHE40, CBX3, CCNT2, CEBPB, CEBPD, CHD1, CHD2, CHD7, CREB1, CTCF, CUX1, E2F6, EGR1, EP300, EZH2, FOXA1, FOXA2, GABPA, GATA1, GATA2, GATA3, GTF2F1, H2AFZ, HCFC1, HDAC2, HMGN3, HNF4G, JUND, KDM4A, MAFF, MAFK, MAX, MAZ, MBD4, MXI1, MYBL2, MYC, NFIC, NFYB, NR2F2, PHF8, POLR2A, RAD21, RBBP5, RCOR1, REST, RFX5, RXRA, SMC3, SP1, SRF, TAF1, TAL1, TBL1XR1, TBP, TCF12, TEAD4, UBTF, USF1, YY1, ZBTB7A, ZC3H11A, ZMIZ1, ZNF143, ZNF384 |
| *LDB3* | AR, SPI1, MITF, TP53, PPARG, HNF4A, YY1, GATA4, SRF, DNAJC2, BHLHE40, BRCA1, CEBPB, CHD2, CTCF, EBF1, ELK1, EP300, ESR1, EZH2, FOS, FOSL1, FOSL2, GATA1, GTF2F1, H2AFZ, HDAC2, JUND, MAFK, MAX, MAZ, MXI1, MYC, MYOD1, MYOG, NFIC, NR2F2, PAX5, POLR2A, POU2F2, RAD21, RBBP5, RCOR1, REST, RFX5, SMARCB1, SMARCC1, SMC3, TBP, TCF12, TCF3, TCF7L2, TEAD4, TFAP2A, TFAP2C, TRIM28, ZBTB7A, ZKSCAN1, ZNF143, ZNF263 |
| *PKN1* | AR, FLI1, SPI1, POU5F1, TCF3, MITF, ZFX, TFAP2C, PPARD, ESR1, SMARCA4, SALL4, KLF1, MECOM, CTNNB1, SIN3A, MYB, RCOR1, TBP, STAT5A, SREBF2, TFEB, ELF1, RBPJ, RARG, ARID3A, ATF3, BACH1, BHLHE40, BRCA1, CBX3, CCNT2, CEBPB, CEBPD, CHD1, CHD2, CHD4, CHD7, CREB1, CTCF, CTCFL, CUX1, E2F4, E2F6, EBF1, EGR1, ELK4, EP300, ETS1, EZH2, FOS, FOSL1, FOSL2, FOXM1, FOXP2, GABPA, GATA1, GATA2, GTF2F1, H2AFZ, HCFC1, HDAC1, HDAC2, HDAC6, HMGN3, IRF1, JUN, JUND, KAT2A, KAT2B, KDM1A, KDM4A, KDM5B, MAFK, MAX, MAZ, MTA3, MXI1, MYBL2, MYC, MYOD1, MYOG, NELFE, NFIC, NR2F2, NR3C1, NRF1, PAX5, PHF8, POLR2A, RAD21, RBBP5, RELA, REST, RFX5, RNF2, RUNX3, RXRA, SAP30, SETDB1, SMARCB1, SMC3, STAT1, SUZ12, TAF1, TBL1XR1, TCF12, TCF7L2, TEAD4, UBTF, USF1, USF2, WHSC1, WRNIP1, YY1, ZBTB7A, ZC3H11A, ZEB1, ZKSCAN1, ZMIZ1, ZNF143, ZNF263, ZNF384 |
| *EMID1* | AR, STAT3, FLI1, RUNX1, EGR1, HNF4A, GATA1, GATA2, TET1, SOX9, SMARCA4, YAP1, TFCP2L1, GATA4, TCF7, EZH2, RNF2, MTF2, SUZ12, PBX1, BACH1, BHLHE40, CBX3, CCNT2, CHD1, CHD2, CTCF, CTCFL, E2F6, EBF1, ELF1, EP300, ETS1, GABPA, GTF2F1, H2AFZ, HCFC1, HDAC1, HDAC2, HMGN3, IKZF1, IRF1, JUND, KAT2A, KDM4A, KDM5B, MAFK, MAX, MAZ, MTA3, MXI1, MYC, MYOD1, MYOG, NFIC, NRF1, PAX5, PHF8, PML, POLR2A, RAD21, RBBP5, RCOR1, RFX5, RUNX3, SAP30, SIN3A, SMC3, SPI1, STAT1, STAT5A, TAF1, TBL1XR1, TBP, TCF12, TCF3, UBTF, WHSC1, WRNIP1, YY1, ZBTB7A, ZEB1, ZMIZ1, ZNF143, ZNF263 |
| *GDA* | AR, TCF4, SMAD4, TP63, KLF4, SOX2, MITF, PPARG, HNF4A, TAL1, ATF3, CTCF, MTF2, SUZ12, PBX1, IRF8, BHLHE40, CEBPB, CHD1, CHD2, E2F6, EP300, ESR1, GTF2B, H2AFZ, HCFC1, IRF1, KDM5B, MAFK, MAX, MAZ, MYC, MYOG, PHF8, PML, POLR2A, RAD21, RBBP5, RCOR1, REST, SAP30, SMC3, SPI1, STAT3, TAF1, TBP, UBTF, YY1, ZNF143, ZNF384 |
| *EPB41L1* | SMAD4, TP63, CREM, E2F1, MYC, SPI1, KLF4, NANOG, POU5F1, SOX2, TCF3, EGR1, MITF, TP53, FOXA2, HNF4A, GATA1, GATA2, RCOR3, SETDB1, TFAP2C, PPARD, TET1, SOX9, SRY, FOXO3, SMARCA4, SALL4, TEAD4, RUNX2, CUX1, SCLY, RNF2, MTF2, SUZ12, TFAP2A, PADI4, PAX6, DNAJC2, NFIB, BHLHE40, CEBPB, CHD1, CTCF, EBF1, ELF1, EZH2, FOXA1, GATA3, H2AFZ, HDAC2, HDAC6, KDM5A, MAX, MYOG, NR2F2, POLR2A, RBBP5, REST, SAP30, TCF12, ZBTB7A |
| *CSF1* | CEBPA, E2F4, MYB, NFKB1, SPI1, STAT1, TP53, TTF2, AR, STAT3, SMAD4, TP63, FLI1, RUNX1, TCF3, EGR1, GATA2, TRIM28, EP300, SCLY, EZH2, JARID2, MTF2, SUZ12, NUCKS1, RELA, ELF5, BACH1, NFIB, ATF2, BCL3, BCLAF1, BHLHE40, CBX3, CCNT2, CEBPB, CHD1, CHD2, CTBP2, CTCF, CUX1, E2F6, EBF1, ELF1, ETS1, FOS, FOSL2, FOXM1, FOXP2, GABPA, GATA3, GTF2F1, H2AFZ, HCFC1, HDAC2, HDAC6, HMGN3, IRF1, JUN, JUND, KDM1A, KDM4A, KDM5B, MAFF, MAFK, MAX, MAZ, MTA3, MXI1, MYC, MYOG, NELFE, NFIC, NR2F2, PHF8, PML, POLR2A, POU2F2, PRDM1, RAD21, RBBP5, RCOR1, REST, RNF2, RUNX3, SAP30, SIN3A, SIRT6, SMC3, SP1, STAT2, STAT5A, TAF1, TAL1, TBL1XR1, TBP, TCF12, TCF7L2, TEAD4, THAP1, UBTF, USF1, USF2, WRNIP1, YY1, ZBTB7A, ZC3H11A, ZEB1, ZMIZ1, ZNF143, ZNF263, ZNF384 |
| *GRIA2* | AHR, ARNT, NFE2L2, TTF2, STAT3, TCF4, SMAD4, RUNX1, SOX2, TCF3, REST, OLIG2, SMARCA4, NR3C1, ZFP42, TCF7, EED, MTF2, SUZ12, POU3F2, BACH1, CHD1, CHD2, CHD7, CREB1, CTBP2, CTCF, E2F6, EGR1, EP300, EZH2, FOSL1, FOXP2, GATA2, GATA3, GTF2F1, H2AFZ, HDAC2, JUND, KDM4A, KDM5A, MAX, MAZ, MXI1, MYC, NRF1, PHF8, POLR2A, |

| | |
|---|---|
| | RAD21, RBBP5, RCOR1, SAP30, SIN3A, TAF1, TAF7, TBP, TCF12, TCF7L2, TEAD4, TRIM28, USF2, WRNIP1, YY1, ZBTB7A, ZNF143, ZNF263 |
| *BMF* | AHR, ARNT, FOXP3, AR, STAT3, CREB1, CREM, E2F1, RUNX1, MYC, SPI1, KLF4, NANOG, POU5F1, SOX2, TCF3, EGR1, MITF, PPARG, FOXA2, HNF4A, GATA1, E2F4, SIN3B, ASH2L, FOXP1, PPARD, TET1, TRIM28, SOX9, EP300, ESR1, SALL4, ZNF281, SMAD2, SMAD3, CUX1, GATA4, TFAP2A, PBX1, STAT5A, SOX11, NACC1, NR0B1, CEBPA, CEBPB, NFIB, SREBF1, ESR2, ARID3A, ATF1, ATF2, BACH1, BATF, BCL3, BCLAF1, BHLHE40, BRCA1, CBX3, CCNT2, CHD1, CHD2, CHD4, CHD7, CTBP2, CTCF, CTCFL, E2F6, EBF1, ELF1, ELK1, ETS1, EZH2, FOS, FOSL2, FOXA1, FOXM1, FOXP2, GABPA, GATA3, GTF2F1, H2AFZ, HCFC1, HDAC2, HMGN3, HNF4G, IKZF1, IRF1, IRF3, IRF4, JUN, JUND, KDM1A, KDM4A, MAFF, MAFK, MAX, MAZ, MEF2A, MTA3, MXI1, MYBL2, MYOD1, MYOG, NFATC1, NFE2, NFIC, NFYA, NFYB, NR2F2, NR3C1, PAX5, PBX3, PHF8, PML, POLR2A, POU2F2, RAD21, RBBP5, RCOR1, RELA, REST, RFX5, RNF2, RUNX3, RXRA, SAP30, SETDB1, SIN3A, SMARCB1, SMC3, SP1, SP4, SRF, STAT1, SUPT20H, SUZ12, TAF1, TAL1, TBL1XR1, TBP, TCF12, TCF7L2, TEAD4, UBTF, USF1, USF2, WRNIP1, YY1, ZBTB33, ZBTB7A, ZC3H11A, ZKSCAN1, ZMIZ1, ZNF143, ZNF263, ZNF384 |
| *ACOT2* | HOXC9, SOX2, TCF3, PPARG, HNF4A, ZFX, REST, SETDB1, FOXP1, CNOT3, TRIM28, ESR1, ATF3, SMAD2, SMAD3, CTCF, ELF5, ELF1, ARID3A, ATF2, BCL3, BCLAF1, BHLHE40, BRCA1, CEBPB, CEBPD, CHD1, CHD2, EBF1, ELK1, EP300, ETS1, EZH2, FOS, FOSL2, FOXA1, FOXA2, FOXM1, GATA2, GATA3, GTF2F1, H2AFZ, HCFC1, HDAC2, HDAC6, HNF4G, JUN, JUND, KDM4A, MAFF, MAFK, MAX, MAZ, MBD4, MTA3, MXI1, MYBL2, MYC, MYOG, NELFE, NFATC1, NFIC, NFYA, NFYB, PAX5, PHF8, PML, POLR2A, POU2F2, RAD21, RBBP5, RCOR1, RFX5, RUNX3, RXRA, SIN3A, SMC3, SP1, SRF, STAT1, STAT5A, TAF1, TBP, TCF12, TCF7L2, TEAD4, UBTF, USF1, USF2, WRNIP1, YY1, ZKSCAN1, ZMIZ1, ZNF143 |
| *SPERT* | STAT3, SMAD4, TP63, NANOG, REST, OLIG2, SMARCA4, TBX3, PBX1, EWSR1, DNAJC2, ATF2, BATF, BCLAF1, CEBPB, CHD1, CREB1, CTCF, EBF1, ELK1, EP300, EZH2, FOXM1, H2AFZ, IKZF1, IRF4, KDM5A, MAFK, MAX, MAZ, MEF2A, MTA3, MXI1, MYC, NFATC1, NFE2, NFIC, NFYB, PAX5, PML, POLR2A, POU2F2, RAD21, RCOR1, RELA, RFX5, RUNX3, SIN3A, SP1, STAT1, STAT5A, TAF1, TBL1XR1, TBP, TCF12, TCF3, WRNIP1, YY1, ZEB1, ZNF143, ZNF384 |
| *ADAM22* | NFE2L2, AR, STAT3, TCF4, SMAD4, TP63, TCF3, EGR1, TP53, FOXA2, HNF4A, KDM5B, PPARD, ERG, TRIM28, RUNX2, CUX1, MTF2, POU3F2, PRDM5, THAP11, ARID3A, ATF1, ATF2, BACH1, BCLAF1, BHLHE40, BRCA1, CBX8, CCNT2, CEBPB, CEBPD, CHD1, CHD2, CHD7, CREB1, CTBP2, CTCF, E2F1, E2F4, E2F6, EBF1, ELF1, ELK1, EP300, ETS1, EZH2, FOXM1, FOXP2, GABPA, GATA1, GATA2, GTF2F1, H2AFZ, HCFC1, HDAC1, HDAC2, HDAC6, HMGN3, IRF1, JUND, KDM4A, KDM5A, MAFK, MAX, MAZ, MEF2A, MTA3, MXI1, MYBL2, MYC, NFATC1, NFIC, NFYB, NRF1, PAX5, PHF8, PML, POLR2A, POU2F2, RAD21, RBBP5, RCOR1, RELA, REST, RFX5, RUNX3, SAP30, SIN3A, SIRT6, SIX5, SMC3, SP1, SREBF1, SRF, STAT1, STAT5A, SUZ12, TAF1, TAF7, TBL1XR1, TBP, TCF12, TCF7L2, TEAD4, UBTF, USF1, USF2, WRNIP1, YY1, ZBTB7A, ZEB1, ZKSCAN1, ZNF143, ZNF263, ZNF384 |
| *SLC41A2* | NFE2L2, AR, STAT3, TCF4, CREB1, CREM, E2F1, MYC, SPI1, NANOG, POU5F1, SOX2, TCF3, MITF, TP53, PPARG, GATA2, EP300, OLIG2, SMARCA4, TEAD4, JARID2, MTF2, SUZ12, PBX1, RELA, EWSR1, STAT5A, NR1I2, ARID3A, ATF2, BRCA1, CBX2, CHD2, CUX1, EBF1, EZH2, FOXA1, FOXM1, GATA3, H2AFZ, HNF4A, JUND, KDM5A, MTA3, MXI1, MYOG, NFIC, PBX3, POLR2A, REST, SIN3A, STAT1, TAL1, TBL1XR1, TCF12, UBTF, WRNIP1, YY1, ZKSCAN1, ZNF384 |
| *TCIRG1* | JUN, MYC, DACH1, AR, CREB1, SPI1, NANOG, TP53, HNF4A, ZFP42, GATA3, RBPJ, RARG, ARID3A, BACH1, BCL11A, BCL3, BCLAF1, BHLHE40, BRCA1, CBX3, CCNT2, CEBPB, CEBPD, CHD1, CHD2, CHD7, CTCF, CUX1, E2F1, E2F4, E2F6, EBF1, EGR1, ELF1, ELK1, EP300, ESR1, ESRRA, ETS1, EZH2, FLI1, FOSL1, FOSL2, FOXA1, FOXA2, FOXM1, FOXP2, GABPA, GATA1, GTF2B, GTF2F1, H2AFZ, HCFC1, HDAC1, HDAC2, HDAC6, HMGN3, HNF4G, IRF1, IRF4, JUND, KAT2A, KDM1A, KDM4A, KDM5B, MAFF, MAFK, MAX, MAZ, MBD4, MTA3, MXI1, MYBL2, NELFE, NFATC1, NFIC, NR2F2, NRF1, PAX5, PBX3, PHF8, PML, POLR2A, POU2F2, RAD21, RBBP5, RCOR1, RELA, REST, RFX5, RUNX3, RXRA, SAP30, SETDB1, SIN3A, SIX5, SMARCB1, SMC3, SP1, SP4, SRF, STAT1, STAT3, STAT5A, TAF1, TAF7, TAL1, TBL1XR1, TBP, TCF12, TCF3, TCF7L2, TEAD4, TFAP2C, THAP1, UBTF, USF1, USF2, WRNIP1, YY1, ZBTB7A, ZC3H11A, ZEB1, ZKSCAN1, ZMIZ1, ZNF143, ZNF384 |
| *FAT4* | PAX3, NFE2L2, DACH1, PPARG, PPARD, SOX9, OLIG2, SMARCA4, YAP1, SOX17, TEAD4, SMAD2, SMAD3, RAD21, EZH2, RNF2, JARID2, MTF2, SUZ12, POU3F2, STAT5A, ZNF652, ARID3A, BACH1, BRCA1, CBX3, CEBPB, CEBPD, CHD1, CHD2, CHD7, CREB1, CTBP2, CTCF, CTCFL, E2F4, E2F6, ELK1, EP300, FOS, FOXA1, FOXA2, FOXP2, GABPA, GATA2, GATA3, GTF2F1, H2AFZ, HDAC2, HDAC6, HMGN3, HNF4A, HNF4G, JUN, JUND, KDM4A, KDM5A, KDM5B, MAFK, MAX, MAZ, MXI1, MYBL2, MYC, MYOG, NFIC, NFYA, NFYB, PBX3, PHF8, POLR2A, RBBP5, RCOR1, REST, RFX5, SAP30, SIN3A, SIRT6, SMC3, SP1, SP2, SP4, STAT3, TAF1, TBP, TCF12, TCF7L2, TRIM28, USF2, YY1, ZBTB7A, ZKSCAN1, ZNF143, ZNF263 |
| *SLC16A12* | CREM, SOX2, TCF3, FOXA2, EP300, YAP1, SALL4, ATF3, MEF2A, RAD21, EED, MTF2, SUZ12, JUN, NR1I2, HTT, ATF2, CEBPB, CHD1, CHD2, CHD7, CTCF, E2F6, EZH2, GABPA, GTF2F1, H2AFZ, HDAC2, JUND, KDM4A, KDM5B, MAX, MXI1, MYC, MYOD1, MYOG, NANOG, NRF1, PHF8, POLR2A, RBBP5, SAP30, SIN3A, TAF1, TAF7, TBP, TCF12, TCF7L2, TEAD4, USF1, USF2, YY1, ZNF263 |
| *TMEM145* | POU5F1, SOX2, TCF3, EGR1, GATA1, RCOR3, SIN3B, REST, EOMES, FOXP1, TRIM28, TEAD4, MEF2A, SIN3A, NR0B1, ZNF274, ATF2, BACH1, BCL3, BHLHE40, CCNT2, CEBPD, CHD1, CHD4, CREB1, CTBP2, CTCF, CTCFL, E2F4, E2F6, ELF1, ELK1, EP300, ETS1, EZH2, FOS, FOXP2, GABPA, GTF2F1, H2AFZ, HCFC1, HDAC1, HDAC2, HDAC6, HMGN3, JUND, KDM4A, KDM5B, MAX, MAZ, MXI1, MYC, NRF1, PAX5, PHF8, POLR2A, RAD21, RBBP5, RCOR1, RFX5, SAP30, SMC3, SP1, SUZ12, TAF1, TBL1XR1, TBP, TCF12, UBTF, YY1, ZBTB7A, ZC3H11A, ZMIZ1, ZNF143, ZNF263, ZNF384 |

| | |
|---|---|
| *SYT13* | ETS1, AR, STAT3, TCF4, E2F1, NANOG, POU5F1, SOX2, MITF, TP53, PPARG, REST, TET1, TAL1, SRY, OLIG2, PRDM14, NR3C1, RUNX2, SUZ12, ESRRB, ATF3, BHLHE40, CEBPB, CHD1, CHD2, CHD7, CREB1, CTCF, E2F6, EGR1, EP300, EZH2, GABPA, GTF2F1, H2AFZ, HDAC2, HMGN3, JUND, KDM4A, KDM5A, KDM5B, MAX, MXI1, MYC, MYOD1, MYOG, PHF8, POLR2A, RBBP5, SAP30, SETDB1, SIN3A, TAF1, TBP, USF1, USF2, ZBTB7A, ZNF143, ZNF263 |
| *KIF5C* | AHR, ARNT, ZNF217, SMAD4, CREB1, CREM, FLI1, KLF4, EGR1, TP53, PPARG, REST, SETDB1, ESR1, OLIG2, WT1, DMRT1, TEAD4, TFCP2L1, RUNX2, RNF2, SUZ12, ESRRB, BACH1, THAP11, ARID3A, ATF1, ATF2, ATF3, BRCA1, CEBPB, CHD1, CHD2, CHD7, CTBP2, CTCF, E2F4, E2F6, ELF1, EP300, ETS1, EZH2, FOSL2, FOXA1, FOXA2, GABPA, GTF2F1, H2AFZ, HCFC1, HDAC1, HDAC2, HMGN3, IRF1, JUN, JUND, KDM4A, KDM5A, KDM5B, MAFF, MAFK, MAX, MAZ, MXI1, MYC, MYOG, NRF1, PAX5, PBX3, PHF8, POLR2A, RAD21, RBBP5, RCOR1, RFX5, SAP30, SIN3A, SIX5, SMC3, SP1, SP4, TAF1, TBP, TCF12, TCF7L2, TRIM28, USF2, YY1, ZEB1, ZNF143, ZNF263 |
| *ARHGDIB* | NFE2L2, AR, TP63, FLI1, RUNX1, SPI1, SOX2, TP53, GATA1, GATA2, PPARD, ERG, SALL4, SMAD2, SMAD3, CUX1, MECOM, LMO2, LYL1, ARID3A, ATF1, ATF2, BATF, BCL3, BCLAF1, BHLHE40, BRCA1, CEBPB, CHD1, CHD2, CREB1, CTCF, E2F4, E2F6, EBF1, EGR1, ELF1, ELK1, EP300, ETS1, EZH2, FOS, FOSL2, FOXA1, FOXM1, FOXP2, GABPA, GATA3, H2AFZ, HCFC1, IRF4, JUN, JUND, KAT2A, MAFF, MAFK, MAX, MAZ, MTA3, MXI1, MYC, MYOD1, MYOG, NELFE, NFATC1, NFE2, NFIC, NR3C1, NRF1, PBX3, PHF8, PML, POLR2A, POU2F2, RAD21, RBBP5, RCOR1, RELA, REST, RFX5, RUNX3, SAP30, SIN3A, SMC3, STAT1, STAT3, STAT5A, TAF1, TAL1, TBL1XR1, TBP, TCF12, TEAD4, TFAP2C, UBTF, USF2, WRNIP1, YY1, ZNF143, ZNF384 |
| *SLC34A3* | STAT3, FLI1, RUNX1, MYC, SPI1, TCF3, EGR1, TP53, RCOR3, FOXP1, TET1, TAL1, SOX9, SRY, MYBL2, MECOM, TCF7, RAD21, MEIS1, BHLHE40, CCNT2, CEBPD, CHD1, CHD2, CTCF, E2F6, EP300, ETS1, EZH2, FOSL2, FOXA1, FOXA2, GABPA, GATA1, H2AFZ, HDAC2, HDAC6, HMGN3, HNF4A, HNF4G, JUND, KDM1A, KDM4A, MAFF, MAFK, MAX, MAZ, MBD4, MXI1, NFIC, NR2C2, NR2F2, PBX3, PHF8, POLR2A, RCOR1, REST, RFX5, RXRA, SP1, SUZ12, TAF1, TCF12, UBTF, USF1, YY1, ZBTB7A, ZKSCAN1 |
| *KIAA1324* | ZNF217, AR, SMAD4, CREM, SPI1, KLF4, NANOG, POU5F1, SOX2, EGR1, MITF, TP53, FOXA2, HNF4A, GATA2, E2F4, RCOR3, TFAP2C, TEAD4, PRDM14, KLF1, CUX1, CTCF, BMI1, SUZ12, CEBPB, ZNF263, ARID3A, ATF2, ATF3, BACH1, BCL11A, BHLHE40, BRCA1, CBX3, CCNT2, CHD1, CHD2, CHD7, CREB1, CTBP2, CTCFL, E2F6, EBF1, ELF1, ELK1, EP300, ETS1, EZH2, FOS, FOXA1, FOXM1, FOXP2, GABPA, GATA1, GATA3, GTF2F1, H2AFZ, HCFC1, HDAC1, HDAC2, HMGN3, JUN, JUND, KDM1A, KDM4A, KDM5A, KDM5B, MAFF, MAFK, MAX, MAZ, MTA3, MXI1, MYC, NFIC, NFYB, NR2F2, PAX5, PHF8, PML, POLR2A, RAD21, RBBP5, RCOR1, RELA, REST, RFX5, RNF2, RUNX3, SAP30, SIN3A, SIRT6, SMC3, SP1, SP4, STAT1, STAT3, STAT5A, TAF1, TAL1, TBL1XR1, TBP, TCF12, TRIM28, UBTF, USF1, USF2, WRNIP1, YY1, ZBTB7A, ZC3H11A, ZEB1, ZKSCAN1, ZMIZ1, ZNF143, ZNF384, ZZZ3 |
| *VIT* | ZNF217, AR, TP63, E2F1, NANOG, POU5F1, SOX2, HNF4A, GATA1, GATA2, RCOR3, PPARD, SOX9, EP300, SALL4, SOX17, LMO2, LYL1, CEBPB, CTCF, E2F4, ELK1, EZH2, FOS, FOSL2, FOXA1, FOXA2, GATA3, H2AFZ, KDM5A, KDM5B, MAFF, MAFK, MAX, MYC, NFIC, PBX3, POLR2A, RAD21, RBBP5, RCOR1, REST, RFX5, SMC3, STAT3, TCF12, TCF7L2 |
| *GPRC5A* | PAX3, PHF8, STAT3, SMAD4, SPI1, KLF4, NANOG, SOX2, TCF3, MITF, TP53, GATA2, RCOR3, ASH2L, TFAP2C, SALL4, WT1, PRDM14, TFCP2L1, ATF3, SMAD2, SMAD3, TFAP2A, PBX1, RELA, NR0B1, CDX2, NFIB, ESR2, RARG, BACH1, BCL3, BHLHE40, BRCA1, CBX2, CBX3, CBX8, CCNT2, CEBPB, CHD1, CHD2, CHD7, CREB1, CTBP2, CTCF, E2F1, E2F4, E2F6, EBF1, EGR1, ELF1, ELK1, ELK4, EP300, ETS1, EZH2, FOS, FOSL2, FOXA1, FOXA2, FOXM1, FOXP2, GABPA, GATA3, GTF2F1, H2AFZ, HCFC1, HDAC1, HDAC2, HDAC6, HMGN3, JUND, KAT2A, KDM1A, KDM4A, KDM5A, KDM5B, MAFK, MAX, MAZ, MXI1, MYC, NFIC, NFYA, NFYB, NR2F2, NR3C1, PBX3, PML, POLR2A, PRDM1, RAD21, RBBP5, RCOR1, REST, RFX5, RNF2, RUNX3, RXRA, SAP30, SIN3A, SIX5, SMC3, SP1, SRF, SUZ12, TAF1, TBL1XR1, TBP, TCF12, TCF7L2, TEAD4, TRIM28, UBTF, USF1, USF2, WHSC1, YY1, ZBTB33, ZBTB7A, ZKSCAN1, ZNF143, ZNF217, ZNF263, ZNF384 |
| *HPD* | NFE2L2, AR, RUNX1, NANOG, POU5F1, EGR1, MITF, FOXA2, HNF4A, GATA1, GATA2, REST, FOXP1, PPARD, TAL1, KLF1, ATF3, RAD21, BMI1, ELF1, ARID3A, ATF1, ATF2, BACH1, BATF, BCL11A, BCL3, BCLAF1, BHLHE40, BRCA1, CBX3, CCNT2, CEBPB, CEBPD, CHD1, CHD2, CHD4, CHD7, CREB1, CTBP2, CTCF, CTCFL, CUX1, E2F4, E2F6, EBF1, ELK1, ELK4, EP300, ETS1, EZH2, FOS, FOSL1, FOSL2, FOXA1, FOXM1, FOXP2, GABPA, GATA3, GTF2F1, GTF3C2, H2AFZ, HCFC1, HDAC1, HDAC2, HDAC6, HMGN3, HNF4G, IRF1, IRF4, JUN, JUND, KAT2B, KDM1A, KDM4A, KDM5A, KDM5B, MAFF, MAFK, MAX, MAZ, MBD4, MTA3, MXI1, MYBL2, MYC, NFATC1, NFE2, NFIC, NR2C2, NR2F2, NR3C1, NRF1, PAX5, PBX3, PHF8, PML, POLR2A, POU2F2, RBBP5, RCOR1, RELA, RFX5, RUNX3, RXRA, SAP30, SETDB1, SIN3A, SIRT6, SIX5, SMARCB1, SMARCC2, SMC3, SP1, SP4, SRF, STAT1, STAT3, STAT5A, SUZ12, TAF1, TAF7, TBL1XR1, TBP, TCF12, TCF3, TCF7L2, TEAD4, TRIM28, UBTF, USF1, USF2, WHSC1, WRNIP1, YY1, ZBTB33, ZBTB7A, ZC3H11A, ZEB1, ZKSCAN1, ZMIZ1, ZNF143, ZNF263, ZNF384 |
| *CYP24A1* | VDR, ELK1, AR, SMAD4, TP63, CREM, FLI1, RUNX1, NANOG, SOX2, TP53, SIN3B, REST, PPARD, ERG, FOXO3, SMARCA4, TFCP2L1, SMAD3, EED, PHC1, EZH2, RNF2, JARID2, MTF2, SUZ12, BACH1, FOXO1, ATF3, CEBPB, CEBPD, CHD1, CHD2, CHD7, CREB1, CTBP2, CTCF, E2F4, E2F6, ELF1, EP300, ETS1, FOS, FOSL2, FOXA2, FOXP2, GABPA, GATA2, GATA3, GTF2F1, H2AFZ, HDAC1, HDAC2, HDAC6, JUND, KDM4A, KDM5A, KDM5B, MAFK, MAX, MAZ, MEF2A, MXI1, MYC, MYOG, NFYB, NR3C1, NRF1, PBX3, PHF8, POLR2A, RBBP5, RCOR1, RFX5, SIN3A, SIX5, SMC3, SP1, STAT3, TAF1, TBP, TCF12, TCF7L2, TEAD4, USF1, USF2, YY1, ZBTB33, ZNF143, ZNF263 |

| | |
|---|---|
| *NEDD9* | AR, RARA, AHR, ARNT, ZNF217, VDR, FOXP3, TCF4, SMAD4, TP63, FLI1, RUNX1, NANOG, SOX2, TCF3, EGR1, MITF, TP53, PPARG, FOXA2, HNF4A, GATA1, SIN3B, REST, ASH2L, TFAP2C, FOXP1, PPARD, ERG, SOX9, SRY, ATF3, SMAD3, CUX1, HOXB4, YY1, MECOM, SUZ12, CTNNB1, MYB, CDX2, SREBF2, NR1I2, TCF7L2, PADI4, RBPJ, PAX6, CLOCK, ATF2, BATF, BCL3, BCLAF1, BHLHE40, BRCA1, CBX2, CBX8, CEBPB, CHD1, CHD2, CREB1, CTCF, EBF1, ELF1, ELK1, EP300, ESR1, ETS1, EZH2, FOXM1, FOXP2, GABPA, H2AFZ, HCFC1, HDAC2, IKZF1, IRF4, KAT2A, KDM5A, KDM5B, MAX, MAZ, MEF2A, MTA3, MXI1, MYC, NELFE, NFATC1, NFIC, NFYB, NR3C1, NRF1, PAX5, PBX3, PML, POLR2A, POU2F2, RAD21, RBBP5, RCOR1, RELA, RFX5, RUNX3, SIN3A, SMC3, SP1, SPI1, SRF, STAT1, STAT3, STAT5A, TAF1, TBL1XR1, TBP, TCF12, TEAD4, USF2, WRNIP1, ZEB1, ZNF143, ZNF384 |
| *STK32C* | DACH1, CREM, FLI1, NANOG, SOX2, EGR1, E2F4, RCOR3, REST, SOX9, TEAD4, PRDM14, NR3C1, SCLY, EZH2, RNF2, MTF2, SUZ12, BCL3, ELF1, RBPJ, NR1H3, BACH1, BHLHE40, BRCA1, CBX8, CCNT2, CEBPB, CEBPD, CHD1, CHD2, CREB1, CTBP2, CTCF, E2F1, E2F6, EBF1, ELK1, EP300, GABPA, GTF2F1, H2AFZ, HCFC1, HDAC1, HDAC2, HDAC6, HMGN3, IRF1, JUND, KDM4A, KDM5B, MAFK, MAX, MAZ, MXI1, MYC, MYOD1, MYOG, NELFE, NR2F2, NRF1, PAX5, PHF8, POLR2A, RAD21, RBBP5, RCOR1, RFX5, RUNX3, SAP30, SIN3A, SMARCB1, SMC3, SP1, SP4, STAT1, STAT3, TAF1, TAF7, TBL1XR1, TBP, TCF12, TCF7L2, TFAP2C, UBTF, USF1, USF2, YY1, ZBTB7A, ZKSCAN1, ZMIZ1, ZNF143, ZNF263 |
| *ATP2A3* | ETS1, TFAP2A, AR, STAT3, CREB1, FLI1, RUNX1, SPI1, KLF4, POU5F1, SOX2, EGR1, GATA1, GATA2, E2F4, SIN3B, TFAP2C, TET1, GFI1B, TAL1, TRIM28, SOX9, SRY, YAP1, SCLY, MECOM, CTCF, MTF2, SUZ12, LMO2, LYL1, MEIS1, ARID3A, ATF2, BCL3, BCLAF1, BHLHE40, BRCA1, CBX3, CCNT2, CEBPB, CHD1, CHD2, CHD7, CTBP2, CUX1, E2F6, EBF1, ELF1, ELK1, EP300, EZH2, FOXM1, GABPA, GATA3, GTF2F1, H2AFZ, HCFC1, HDAC1, HDAC2, HDAC6, HMGN3, IRF1, JUN, JUND, KAT2A, KDM1A, KDM4A, KDM5B, MAFK, MAX, MAZ, MTA3, MXI1, MYC, MYOD1, MYOG, NCOR1, NELFE, NFATC1, NR2F2, NRF1, PAX5, PHF8, PML, POLR2A, POU2F2, RAD21, RBBP5, RCOR1, RELA, REST, RFX5, RUNX3, SAP30, SETDB1, SIN3A, SMARCB1, SMC3, SP1, SRF, STAT1, STAT5A, TAF1, TBL1XR1, TBP, TCF12, TCF3, TEAD4, UBTF, USF2, WHSC1, WRNIP1, YY1, ZBTB7A, ZC3H11A, ZEB1, ZKSCAN1, ZMIZ1, ZNF143, ZNF384 |
| *MNX1* | MYC, TTF2, SMAD4, TP63, KLF4, NANOG, POU5F1, SOX2, TCF3, EGR1, MITF, TP53, FOXA2, KDM5B, SIN3B, SETDB1, SRY, ZNF281, RUNX2, YY1, BMI1, EED, PHC1, EZH2, RNF2, JARID2, MTF2, SUZ12, BACH1, CDX2, ELF1, ARID3A, ATF3, BCL3, BCLAF1, BHLHE40, BRCA1, CBX3, CCNT2, CEBPB, CEBPD, CHD1, CHD2, CHD7, CREB1, CTBP2, CTCF, CUX1, E2F1, E2F4, E2F6, EBF1, ELK1, EP300, ETS1, FOSL1, FOXA1, FOXP2, GABPA, GATA3, GTF2F1, H2AFZ, HCFC1, HDAC1, HDAC2, HMGN3, IRF1, IRF3, JUN, JUND, KDM1A, KDM4A, KDM5A, MAFK, MAX, MAZ, MBD4, MTA3, MXI1, MYBL2, NCOR1, NFIC, NFYA, NFYB, NR2F2, NRF1, PAX5, PBX3, PHF8, PML, POLR2A, RAD21, RBBP5, RCOR1, RELA, REST, RFX5, RUNX3, RXRA, SAP30, SIN3A, SIRT6, SMARCB1, SMARCC1, SMC3, SP1, SP2, STAT1, STAT5A, TAF1, TAL1, TBL1XR1, TBP, TCF12, TCF7L2, TEAD4, TRIM28, UBTF, USF1, USF2, WHSC1, WRNIP1, ZBTB7A, ZC3H11A, ZEB1, ZKSCAN1, ZMIZ1, ZNF143, ZNF263, ZNF384 |
| *DGKG* | TP53, AR, STAT3, TCF4, SMAD4, TP63, RUNX1, MYC, SPI1, POU5F1, MITF, GATA1, SIN3B, REST, YAP1, PRDM14, RUNX2, YY1, MECOM, STAT4, TCF7, EZH2, RNF2, MTF2, SUZ12, MYB, PAX6, FOXO1, BACH1, BHLHE40, CBX2, CBX8, CHD1, CHD2, CHD7, CTBP2, CTCF, EBF1, EGR1, ELF1, EP300, ETS1, FOXA1, FOXA2, FOXM1, GABPA, H2AFZ, HCFC1, HDAC1, HDAC2, HNF4A, HNF4G, JUN, JUND, KAT2A, KDM4A, KDM5B, MAFK, MAX, MAZ, MTA3, MXI1, NANOG, NELFE, NFIC, NRF1, PBX3, PHF8, PML, POLR2A, RAD21, RBBP5, RCOR1, RUNX3, SAP30, SIN3A, SMC3, SRF, TBP, UBTF, USF1, USF2, ZBTB7A, ZMIZ1, ZNF143, ZNF263, ZNF384 |
| *COL28A1* | PAX3, AR, TCF4, SMAD4, TP63, MYC, RUNX2, MEF2A, RBPJ, CTCF, EP300, EZH2, H2AFZ, KDM5A, NFIC, POLR2A |
| *ATP6V0A4* | AR, FLI1, RUNX1, MYC, NANOG, POU5F1, EGR1, MITF, TP53, PPARG, TRIM28, ESR1, SMARCA4, DMRT1, TEAD4, ESR2, ATF2, CBX2, CHD1, CHD2, CHD7, CTCF, ELF1, EP300, EZH2, FOXA2, GATA1, GTF2F1, H2AFZ, HDAC1, HDAC2, HDAC6, JUND, KAT2B, KDM4A, KDM5A, MAFF, MAFK, MAX, PHF8, POLR2A, RAD21, RBBP5, REST, SIN3A, SIRT6, SUZ12, TBP, TCF12, USF2, YY1, ZNF143, ZNF384 |
| *CPNE5* | AR, STAT3, TP63, MYC, NANOG, POU5F1, TCF3, EGR1, MITF, TP53, PPARG, FOXA2, HNF4A, GATA2, RCOR3, SETDB1, ASH2L, PPARD, ZNF281, DMRT1, PRDM14, NR3C1, RUNX2, SCLY, RNF2, MTF2, SUZ12, ESRRB, JUN, LMO2, LYL1, HSF1, ARID3A, BHLHE40, BRCA1, CBX3, CBX8, CCNT2, CEBPB, CHD1, CHD2, CREB1, CTBP2, CTCF, CTCFL, E2F4, E2F6, EBF1, ELF1, EP300, EZH2, FOXA1, GABPA, H2AFZ, HCFC1, HDAC2, HDAC6, HMGN3, JUND, KDM1A, KDM4A, KDM5B, MAFK, MAX, MAZ, MYOG, PAX5, PHF8, POLR2A, RAD21, RBBP5, RCOR1, RFX5, SAP30, SIN3A, TAL1, TBP, TEAD4, UBTF, USF2, YY1, ZBTB7A, ZMIZ1, ZNF143, ZNF263, ZNF384 |
| *MAB21L1* | PAX3, HOXC9, RUNX1, SPI1, NANOG, POU5F1, SOX2, TP53, PPARG, EP300, SMARCA4, YAP1, NR3C1, EED, PHC1, EZH2, RNF2, JARID2, MTF2, SUZ12, TFAP2A, PBX1, POU3F2, GATA3, DNAJC2, BACH1, CBX2, CBX8, CHD1, CHD2, CHD7, CTBP2, CTCF, EGR1, GATA2, GTF2F1, H2AFZ, HDAC2, JUND, KDM4A, KDM5A, KDM5B, MAFK, MAX, MXI1, MYC, NFIC, PBX3, POLR2A, RAD21, RBBP5, REST, SAP30, SIN3A, SP1, STAT3, TAF1, TBP, TCF12, TEAD4, TRIM28, USF1, WRNIP1, YY1, ZNF143, ZNF263 |
| *GFI1* | RUNX1, MYC, SOX2, EGR1, TP53, GATA2, E2F4, ERG, TRIM28, SRY, STAT4, BMI1, EZH2, RNF2, JARID2, MTF2, SUZ12, MYB, MEIS1, POU3F2, ARID3A, ATF2, BACH1, BCL11A, BCL3, BCLAF1, BHLHE40, BRCA1, CBX2, CBX8, CCNT2, CEBPB, CHD1, CHD2, CHD7, CREB1, CTCF, CTCFL, CUX1, E2F6, EBF1, ELF1, ELK1, EP300, ETS1, FOXM1, GABPA, GTF2F1, H2AFZ, HCFC1, HDAC1, HDAC2, HDAC6, HMGN3, IKZF1, JUN, JUND, KDM1A, KDM5B, MAFF, MAFK, MAX, MAZ, MEF2A, MEF2C, MTA3, MXI1, MYBL2, MYOD1, MYOG, NANOG, NELFE, NFATC1, NFIC, NFYB, NRF1, PAX5, PHF8, PML, POLR2A, POU2F2, RAD21, RBBP5, RCOR1, RELA, REST, RFX5, RUNX3, SAP30, SIN3A, SMC3, SP2, SPI1, STAT1, STAT5A, TAF1, TAL1, TBL1XR1, TBP, TCF12, TCF3, TCF7L2, UBTF, USF2, WRNIP1, YY1, ZBTB7A, ZEB1, ZMIZ1, ZNF143, ZNF263, ZNF384 |

| | |
|---|---|
| *SLC15A3* | FLI1, RUNX1, SPI1, KLF4, NANOG, POU5F1, SOX2, EGR1, PPARG, E2F4, TFAP2C, PPARD, GFI1B, SRY, MECOM, LMO2, MEIS1, ASXL1, IRF1, IRF8, ATF2, BATF, BCLAF1, BHLHE40, BRCA1, CBX2, CBX8, CEBPB, CHD1, CHD2, CHD7, CTBP2, CTCF, CUX1, E2F6, EBF1, ELF1, ELK1, EP300, ETS1, EZH2, FOXM1, GTF2F1, H2AFZ, HDAC1, HDAC2, IKZF1, IRF4, JUND, KDM4A, KDM5A, KDM5B, MAX, MAZ, MTA3, MXI1, MYC, NFATC1, NFE2, NFIC, NFYA, NFYB, NRF1, PAX5, PBX3, PHF8, PML, POLR2A, POU2F2, RBBP5, RCOR1, RELA, REST, RFX5, RNF2, RUNX3, SAP30, SIN3A, SMC3, SP1, STAT1, STAT5A, SUZ12, TAF1, TBL1XR1, TBP, TCF12, TEAD4, UBTF, USF1, USF2, WRNIP1, YY1, ZEB1, ZMIZ1, ZNF143 |
| *MAATS1* | AR, STAT3, CREM, MYC, SOX2, PPARG, HNF4A, PPARD, BACH1, BHLHE40, BRCA1, CEBPB, CHD1, CHD2, CHD7, CTCF, E2F4, E2F6, ELF1, EP300, EZH2, FOXA1, FOXA2, GTF2F1, H2AFZ, HDAC2, IRF1, JUND, KDM4A, KDM5A, MAX, MAZ, MTA3, MXI1, NFE2, NR2F2, PHF8, POLR2A, RAD21, RBBP5, RCOR1, RELA, REST, RFX5, RUNX3, SAP30, SIN3A, SP1, SP4, SPI1, TAF1, TAL1, TBP, TCF12, TEAD4, USF1, USF2, WRNIP1, YY1, ZBTB7A, ZNF143 |
| *SRGAP1* | AHR, ARNT, ZNF217, DACH1, STAT3, SMAD4, TP63, CREM, FLI1, RUNX1, KLF4, POU5F1, SOX2, MITF, TP53, FOXA2, HNF4A, E2F4, SETDB1, TFAP2C, PPARD, SOX9, ESR1, SMARCA4, NR3C1, RUNX2, YY1, CTCF, ESRRB, TFAP2A, RELA, SOX11, CRX, ARID3A, ATF3, BACH1, BATF, BCL3, BCLAF1, BHLHE40, BRCA1, CBX2, CBX3, CBX8, CCNT2, CEBPB, CHD1, CHD2, CHD7, CREB1, CTBP2, CTCFL, CUX1, E2F1, E2F6, EBF1, EGR1, ELF1, ELK1, EP300, ETS1, EZH2, FOS, FOSL2, FOXA1, FOXM1, FOXP2, GABPA, GATA2, GATA3, GTF2F1, H2AFZ, HCFC1, HDAC1, HDAC2, HDAC6, HMGN3, IRF3, JUN, JUND, KAT2B, KDM4A, KDM5A, KDM5B, MAFK, MAX, MAZ, MTA3, MXI1, MYBL2, MYC, NFATC1, NFIC, NFYA, NFYB, NR2F2, NRF1, PAX5, PBX3, PHF8, PML, POLR2A, POU2F2, RAD21, RBBP5, RCOR1, REST, RFX5, RNF2, RUNX3, SAP30, SIN3A, SMARCB1, SMARCC1, SMC3, SP1, SP2, SP4, SPI1, SRF, STAT1, STAT5A, SUZ12, TAF1, TAF7, TBL1XR1, TBP, TCF12, TCF3, TCF7L2, TEAD4, TRIM28, UBTF, USF1, USF2, WHSC1, WRNIP1, ZBTB33, ZBTB7A, ZC3H11A, ZKSCAN1, ZMIZ1, ZNF143, ZNF263, ZNF384 |
| *GCNT1* | ZNF217, CEBPD, NFE2L2, STAT3, SMAD4, CREB1, E2F1, RUNX1, MYC, SPI1, KLF4, SOX2, PPARG, HNF4A, GATA1, SETDB1, ASH2L, PPARD, TAL1, EP300, SMARCA4, SALL4, SOX17, TEAD4, KLF1, SMAD3, CTCF, TBX3, LMO2, RELA, CEBPB, ATF2, BATF, BCL11A, BCL3, BCLAF1, BHLHE40, CHD1, CHD2, CUX1, EBF1, ESR1, EZH2, FOS, FOSL2, FOXM1, GATA2, GATA3, H2AFZ, HDAC2, IKZF1, IRF4, JUN, JUND, KDM5A, MAFF, MAFK, MAX, MAZ, MEF2A, MTA3, MXI1, NFATC1, NFIC, PAX5, PML, POLR2A, POU2F2, RCOR1, RUNX3, SMC3, SP1, STAT5A, TBL1XR1, TBP, TRIM28, WRNIP1, ZNF384 |
| *BANK1* | ZNF217, CEBPD, STAT3, RUNX1, MYC, SOX2, EGR1, FOXA2, MYB, POU3F2, IRF8, ATF2, BATF, BCL11A, BCL3, BCLAF1, BHLHE40, CEBPB, CHD1, CHD2, CHD7, CTBP2, CTCF, CUX1, E2F6, EBF1, ELF1, ELK1, EP300, EZH2, FLI1, FOXM1, GTF2F1, H2AFZ, HCFC1, HDAC2, HMGN3, IRF4, JUND, KAT2A, KDM4A, KDM5A, MAX, MAZ, MEF2A, MTA3, MXI1, NANOG, NELFE, NFATC1, NFIC, NRF1, PAX5, PHF8, PML, POLR2A, POU2F2, RAD21, RBBP5, RCOR1, RELA, REST, RUNX3, SAP30, SIN3A, SMC3, SP1, SP4, SPI1, SRF, STAT1, STAT5A, TAF1, TBL1XR1, TBP, TCF12, TCF3, TCF7L2, TEAD4, UBTF, USF2, WRNIP1, YY1, ZBTB7A, ZEB1, ZNF143, ZNF384 |
| *NLGN4X* | PAX3, AR, TCF4, SMAD4, FLI1, MYC, SOX2, TFAP2A, BACH1, NOTCH1, CEBPB, CHD1, CHD2, CHD7, CTCF, E2F6, EP300, EZH2, GTF2F1, H2AFZ, HDAC2, KDM4A, MAFK, MAX, MXI1, NFYB, PHF8, POLR2A, RAD21, RBBP5, REST, SAP30, SIN3A, TAF1, TBP, TCF12, USF1, YY1, ZNF143 |
| *PDGFD* | TCF4, SMAD4, MYC, SOX2, EGR1, REST, FOXP1, POU3F2, STAT5A, STAT1, RARG, BACH1, BCL3, BRCA1, CEBPB, CHD1, CHD2, CHD7, CREB1, CTBP2, CTCF, E2F6, ELF1, EP300, EZH2, FOXA2, FOXP2, GABPA, GATA2, GATA3, GTF2F1, H2AFZ, HDAC2, HDAC6, HMGN3, JUND, KDM4A, KDM5A, MAFK, MAX, MAZ, MXI1, NANOG, NRF1, PHF8, POLR2A, RAD21, RBBP5, RFX5, SAP30, SIN3A, SIRT6, SMC3, SP1, STAT3, SUZ12, TAF1, TBP, TCF12, TCF7L2, TEAD4, TRIM28, USF2, YY1, ZNF143, ZNF263, ZNF384 |
| *IL15RA* | TP53, AR, TCF4, CREB1, FLI1, SPI1, SOX2, EGR1, FOXA2, HNF4A, GATA1, GATA2, SOX9, SRY, SRF, EED, PHC1, EZH2, RNF2, JARID2, MTF2, SUZ12, MYB, RELA, STAT5A, CEBPB, ARID3A, ATF3, BACH1, BCLAF1, BHLHE40, BRCA1, CBX3, CBX8, CCNT2, CHD1, CHD2, CHD4, CHD7, CTBP2, CTCF, CUX1, E2F4, E2F6, EBF1, ELF1, ELK1, EP300, ETS1, FOS, FOSL1, FOSL2, GABPA, GTF2B, GTF2F1, H2AFZ, HCFC1, HDAC1, HDAC2, HDAC6, HMGN3, IKZF1, IRF1, JUN, JUND, KAT2B, KDM4A, KDM5B, MAFF, MAFK, MAX, MAZ, MTA3, MXI1, MYC, NCOR1, NELFE, NR2F2, NRF1, PAX5, PHF8, PML, POLR2A, POU2F2, RAD21, RBBP5, RCOR1, REST, RFX5, RUNX3, SAP30, SETDB1, SIN3A, SMARCB1, SMC3, STAT1, TAF1, TAL1, TBL1XR1, TBP, TCF12, TCF3, TCF7L2, TEAD4, TFAP2C, TRIM28, UBTF, USF1, USF2, WRNIP1, YY1, ZBTB7A, ZC3H11A, ZEB1, ZKSCAN1, ZMIZ1, ZNF143, ZNF263, ZNF384 |
| *EPHA3* | ZNF217, CEBPD, AR, TCF4, TP63, NANOG, POU5F1, SOX2, TCF3, MITF, TP53, SIN3B, ASH2L, FOXP1, PPARD, EP300, OLIG2, SMARCA4, CUX1, SCLY, GATA4, SRF, BMI1, EED, MTF2, SUZ12, POU3F2, BACH1, CDKN2AIP, CEBPB, CHD1, CHD2, CHD7, CREB1, CTCF, E2F4, E2F6, EBF1, EGR1, ELF1, EZH2, FOSL1, FOXP2, GATA2, GATA3, GTF2F1, H2AFZ, HDAC1, HDAC2, KDM4A, KDM5A, MAX, MXI1, MYC, PAX5, PHF8, POLR2A, RAD21, RBBP5, RCOR1, REST, RFX5, SAP30, SETDB1, SIN3A, SIRT6, SP1, STAT1, TAF1, TBP, TCF12, TCF7L2, TEAD4, TRIM28, USF2, WRNIP1, YY1, ZNF143, ZNF263 |
| *SCN5A* | AR, STAT3, TP63, SPI1, NANOG, POU5F1, SOX2, TCF3, EGR1, MITF, KDM5B, TET1, SOX9, SRY, EP300, EZH2, RNF2, JARID2, MTF2, SUZ12, ESRRB, CTNNB1, TBX3, SMAD1, ATF2, ATF3, BACH1, BCL3, BHLHE40, CBX3, CCNT2, CHD1, CHD2, CHD7, CREB1, CTBP2, CTCF, E2F6, ELF1, ELK1, ETS1, FOXM1, GTF2F1, H2AFZ, HCFC1, HDAC1, HDAC2, HDAC6, HMGN3, IRF1, JUND, KDM4A, KDM5A, MAX, MAZ, MXI1, MYC, MYOD1, MYOG, NCOR1, NELFE, NFATC1, PBX3, PHF8, PML, POLR2A, RAD21, RBBP5, RCOR1, REST, RFX5, RUNX3, SAP30, SIN3A, SIRT6, SMC3, SP1, SP4, STAT1, STAT5A, TAF1, TAL1, TBL1XR1, TBP, TCF12, TEAD4, UBTF, USF1, USF2, WRNIP1, YY1, ZBTB7A, ZC3H11A, ZNF143, ZNF263 |
| *ATP6V1B1* | AR, NANOG, SOX2, EGR1, HNF4A, KDM5B, RCOR3, TET1, ZNF263, CEBPB, CHD2, CTCF, EP300, EZH2, FOXA1, FOXA2, H2AFZ, HDAC1, HDAC2, KDM5A, MAZ, MYC, NR2F2, POLR2A, RBBP5, REST, SAP30, SIN3A, TEAD4, TFAP2A, TFAP2C |

| | |
|---|---|
| *CSF1R* | AR, TCF4, FLI1, SPI1, TP53, HNF4A, EOMES, TFAP2C, TAL1, SMARCA4, ATF3, CUX1, SCLY, MECOM, LMO2, CBX2, CEBPB, CHD1, CTCF, E2F4, E2F6, EGR1, EP300, EZH2, FOS, FOSL2, GATA2, GATA3, GTF2F1, H2AFZ, HDAC1, JUND, KDM5B, MAFK, MAX, MAZ, MXI1, MYC, POLR2A, RAD21, RCOR1, SAP30, SIN3A, SMARCC1, STAT3, TCF7L2, TFAP2A, WHSC1, ZKSCAN1, ZNF143, ZNF384 |
| *KCNN4* | VDR, E2F1, EGR1, TP53, GATA1, GATA2, REST, TAL1, SRY, ATF3, SMAD2, SMAD3, STAT4, TCF7, SOX11, DNAJC2, ARID3A, ATF2, BATF, BCL3, BCLAF1, BHLHE40, BRCA1, CBX3, CCNT2, CEBPB, CHD1, CHD2, CHD4, CTCF, CTCFL, CUX1, E2F4, E2F6, EBF1, ELF1, ELK1, EP300, ETS1, EZH2, FOS, FOSL1, FOSL2, FOXA2, FOXM1, GATA3, GTF2F1, H2AFZ, HCFC1, HDAC1, HDAC2, HDAC6, HMGN3, IRF4, JUND, KAT2B, KDM5A, MAFK, MAX, MAZ, MTA3, MXI1, MYC, MYOD1, MYOG, NELFE, NFATC1, NFIC, NR2C2, NR2F2, PHF8, PML, POLR2A, POU2F2, RAD21, RBBP5, RCOR1, RELA, RFX5, RUNX3, SAP30, SIN3A, SMC3, SP1, STAT1, STAT3, STAT5A, TAF1, TBL1XR1, TBP, TCF12, TCF3, TCF7L2, TEAD4, TFAP2A, TFAP2C, UBTF, USF1, USF2, WHSC1, WRNIP1, YY1, ZBTB7A, ZC3H11A, ZKSCAN1, ZMIZ1, ZNF143, ZNF263, ZNF384 |
| *GPR17* | AR, MYC, HNF4A, SREBF2, STAT6, ARID3A, ATF3, CCNT2, CEBPB, CHD1, CHD7, CTCF, CTCFL, EP300, EZH2, FOS, H2AFZ, HDAC1, HDAC2, KDM5A, KDM5B, MAX, MAZ, POLR2A, RAD21, RBBP5, RCOR1, SUZ12, UBTF, USF1, USF2, ZNF143 |
| *FAM181B* | ZNF217, STAT3, TP63, POU5F1, SOX2, TP53, PPARG, GATA2, SIN3B, TFAP2C, PPARD, TET1, ERG, GFI1B, TAL1, TRIM28, SMARCA4, YAP1, KLF1, NR3C1, RUNX2, EZH2, RNF2, JARID2, MTF2, SUZ12, NR0B1, STAT1, BRCA1, CCNT2, CHD1, CHD2, CTBP2, CTCF, E2F6, EGR1, ELF1, EP300, FOXP2, GTF2F1, H2AFZ, HCFC1, HDAC1, HDAC2, HDAC6, HMGN3, JUND, KDM4A, KDM5B, MAX, MAZ, MYC, MYOG, PHF8, POLR2A, RAD21, RBBP5, RCOR1, REST, SAP30, SIN3A, TAF1, TBP, UBTF, YY1, ZBTB7A, ZMIZ1, ZNF143 |
| *MMP11* | CEBPA, E2F1, RUNX1, SPI1, KLF4, NANOG, POU5F1, SOX2, TCF3, EGR1, HNF4A, ZFX, TFAP2C, PPARD, SOX9, SRY, ZNF281, TFCP2L1, CUX1, TBX5, NR0B1, TAF7L, PAX6, DNAJC2, ARID3A, BACH1, BHLHE40, BRCA1, CCNT2, CEBPB, CEBPD, CEBPZ, CHD1, CHD2, CHD7, CTBP2, CTCF, CTCFL, E2F4, E2F6, EBF1, ELF1, ELK1, EP300, EZH2, FOS, FOSL2, FOXA1, FOXA2, FOXP2, GATA2, GTF2F1, H2AFZ, HCFC1, HDAC1, HDAC2, HDAC6, HMGN3, HNF4G, JUND, KDM4A, KDM5A, KDM5B, MAFF, MAFK, MAX, MAZ, MBD4, MEF2A, MXI1, MYBL2, MYC, MYOG, NELFE, NFIC, NR2F2, PHF8, POLR2A, RAD21, RBBP5, RCOR1, REST, RFX5, RUNX3, RXRA, SAP30, SIN3A, SMC3, SP1, SRF, STAT3, SUZ12, TAF1, TBL1XR1, TBP, TCF12, TCF7L2, TEAD4, TRIM28, UBTF, USF1, USF2, WRNIP1, YY1, ZBTB7A, ZEB1, ZNF143, ZNF384 |
| *MB* | AHR, ARNT, E2F1, SPI1, NANOG, POU5F1, GATA1, SIN3B, PPARD, EP300, ESR1, YAP1, CUX1, STAT4, BMI1, TFAP2A, NACC1, NR0B1, SMAD1, ESR2, HTT, BRCA1, CEBPB, CHD1, CHD2, CHD7, CTCF, EZH2, FOXA1, FOXA2, GABPA, GATA2, GATA3, GTF2F1, H2AFZ, HDAC2, HDAC6, MAFK, MAX, MAZ, MXI1, MYC, NFIC, NFYB, NR2F2, NR3C1, POLR2A, RAD21, RBBP5, RCOR1, REST, RFX5, SAP30, SIN3A, SIRT6, SMC3, SP1, SP4, STAT5A, SUZ12, TBP, TCF12, TCF7L2, TEAD4, TFAP2C, TRIM28, UBTF, WRNIP1, YY1, ZKSCAN1, ZNF143, ZNF217 |
| *PILRB* | HOXC9, XRN2, RUNX1, MYC, SPI1, EGR1, MITF, GATA1, ATF3, RUNX2, CUX1, TCF7, LMO2, STAT5A, ATF1, ATF2, BACH1, BATF, BCL11A, BCL3, BCLAF1, BHLHE40, BRCA1, CBX3, CCNT2, CEBPB, CEBPD, CHD1, CHD2, CREB1, CTCF, E2F4, E2F6, EBF1, ELF1, ELK1, ELK4, EP300, ETS1, FOS, FOSL1, FOSL2, FOXA2, FOXM1, FOXP2, GABPA, GATA2, GTF2B, GTF2F1, H2AFZ, HCFC1, HDAC1, HDAC2, HDAC6, HMGN3, IRF1, IRF3, IRF4, JUN, JUND, KDM4A, KDM5B, MAFK, MAX, MAZ, MEF2A, MTA3, MXI1, MYBL2, NANOG, NFATC1, NFIC, NFYA, NFYB, NR2C2, NR2F2, NR3C1, NRF1, PAX5, PBX3, PHF8, PML, POLR2A, POU2F2, RBBP5, RCOR1, RELA, REST, RFX5, RUNX3, RXRA, SAP30, SETDB1, SIN3A, SIX5, SMC3, SP1, SP2, STAT1, STAT3, TAF1, TAF7, TAL1, TBL1XR1, TBP, TCF12, TCF3, TCF7L2, TEAD4, TRIM28, UBTF, USF1, WRNIP1, YY1, ZBTB33, ZBTB7A, ZEB1, ZKSCAN1, ZMIZ1, ZNF143, ZNF263, ZNF384 |
| *CDH5* | E2F1, ERG, SP3, PAX3, CEBPD, NFE2L2, AR, TCF4, RUNX1, SPI1, KLF4, NANOG, POU5F1, SOX2, EGR1, MITF, PPARG, GATA2, E2F4, REST, TAL1, ESR1, YAP1, KLF1, RUNX2, TFAP2A, LMO2, MEIS1, ELF1, RBPJ, CDKN2AIP, BRCA1, CEBPB, CHD1, CHD2, CTCF, EBF1, EP300, EZH2, FOS, FOXA1, FOXA2, FOXP2, H2AFZ, HDAC2, HDAC6, JUN, JUND, KDM5B, MAFK, MAX, MYC, NR3C1, PBX3, POLR2A, RBBP5, RCOR1, RFX5, SRF, ZBTB7A, ZNF143, ZNF384 |
| *SYTL3* | VDR, SMAD4, TP63, RUNX1, SPI1, POU5F1, SOX2, FOXA2, HNF4A, GATA1, SETDB1, SRY, OLIG2, YY1, CTNNB1, PBX1, SOX11, NFIB, NR1H3, STAT6, ATF2, BATF, BCL11A, BCL3, BCLAF1, BHLHE40, CBX3, CEBPB, CHD1, CHD2, CREB1, CTCF, CUX1, E2F4, EBF1, ELF1, ELK1, EP300, EZH2, FOS, FOSL1, FOSL2, FOXM1, GATA3, H2AFZ, HDAC1, HDAC2, IKZF1, IRF3, IRF4, JUND, KDM5B, MAFK, MAX, MAZ, MEF2A, MEF2C, MTA3, MXI1, MYC, NFATC1, NFIC, NRF1, PAX5, PBX3, PML, POLR2A, POU2F2, RBBP5, RCOR1, RELA, REST, RNF2, RUNX3, RXRA, SIN3A, SMC3, SP1, SRF, STAT1, STAT3, STAT5A, TAF1, TBL1XR1, TBP, TCF12, TCF3, TEAD4, UBTF, USF1, USF2, WRNIP1, ZNF143, ZNF384 |
| *ZNF92* | HOXC9, STAT3, SMAD4, SPI1, SCLY, PBX1, PRDM5, ARID3A, ATF1, ATF2, ATF3, BCL3, BCLAF1, BHLHE40, BRCA1, CBX3, CCNT2, CEBPB, CEBPD, CHD1, CHD2, CREB1, CTCF, CUX1, E2F4, E2F6, EBF1, ELF1, ELK1, EP300, ETS1, EZH2, FOS, FOXM1, GABPA, GATA1, GATA2, GATA3, GTF2B, GTF2F1, H2AFZ, HCFC1, HDAC1, HDAC2, HDAC6, HMGN3, IRF1, IRF3, JUND, KDM4A, KDM5B, MAFF, MAFK, MAX, MAZ, MBD4, MTA3, MXI1, MYBL2, MYC, NANOG, NFATC1, NFIC, NFYA, NFYB, NR2F2, PAX5, PBX3, PHF8, PML, POLR2A, POU2F2, RAD21, RBBP5, RCOR1, RELA, REST, RFX5, RUNX3, SAP30, SETDB1, SIN3A, SMC3, SP1, SP2, SP4, STAT1, STAT5A, TAF1, TAF7, TAL1, TBL1XR1, TBP, TCF12, TCF3, TCF7L2, TEAD4, TRIM28, UBTF, USF1, USF2, WRNIP1, YY1, ZBTB33, ZBTB7A, ZC3H11A, ZEB1, ZKSCAN1, ZMIZ1, ZNF143, ZNF263, ZNF384 |

| | |
|---|---|
| *KYNU* | ZNF217, AR, STAT3, MYC, SPI1, NANOG, SOX2, TP53, HNF4A, RCOR3, SIN3B, REST, TFAP2C, PPARD, ESR1, ATF3, SMAD2, SMAD3, NR3C1, RUNX2, CUX1, ESRRB, ELF5, CEBPA, IRF8, NR1H3, ESR2, ARID3A, BATF, BCL3, BHLHE40, CEBPB, CHD1, CHD2, CTCF, E2F4, E2F6, EBF1, EP300, ETS1, EZH2, FOS, FOXA1, FOXA2, GATA3, GTF2F1, H2AFZ, HCFC1, HDAC1, HDAC2, HNF4G, MAFK, MAX, MAZ, MBD4, MEF2A, MXI1, MYBL2, NELFE, NFE2, NFIC, NFYB, POLR2A, RAD21, RCOR1, RUNX3, RXRA, SIN3A, SMC3, SP1, STAT1, TAF1, TBL1XR1, TBP, TCF12, TCF7L2, TEAD4, USF1, USF2, WRNIP1, YY1, ZC3H11A, ZNF143, ZNF384 |
| *TRIM55* | AHR, ARNT, TTF2, RUNX1, TP53, SETDB1, PPARD, TAL1, EP300, ESR1, KLF1, ATF3, SMAD2, SMAD3, CUX1, SCLY, GATA4, MEF2A, SRF, FOXP2, PBX1, POU3F2, RELA, ELF5, EWSR1, STAT5A, BHLHE40, CEBPB, CHD1, CTCF, E2F6, EGR1, EZH2, FOSL2, H2AFZ, HDAC6, MAFF, MAFK, MAX, MYC, MYOD1, MYOG, PAX5, PBX3, POLR2A, POU2F2, RCOR1, RFX5, SIN3A, TAF1, TCF12, TCF3, USF1, USF2, WRNIP1, ZNF384 |
| *DRD1* | AR, CREB1, SP1, TFAP2A, TFAP2B, AHR, ARNT, ZNF217, PAX3, RUNX1, MYC, SOX2, RCOR3, SIN3B, REST, PPARD, SRY, SOX17, RUNX2, TBX5, CTCF, MTF2, SUZ12, JUN, MEIS1, BACH1, BHLHE40, BRCA1, CCNT2, CEBPB, CHD1, CHD2, CTBP2, CTCFL, E2F6, EBF1, EP300, EZH2, FOSL1, FOXA2, FOXP2, GABPA, GATA2, GTF2F1, H2AFZ, HCFC1, HDAC1, HDAC2, HDAC6, HMGN3, JUND, KDM4A, KDM5A, KDM5B, MAFK, MAX, MAZ, MXI1, NR2F2, PAX5, PHF8, POLR2A, RAD21, RBBP5, RCOR1, SAP30, SIN3A, SMC3, TAF7, TAL1, TBP, TCF12, TEAD4, USF1, USF2, YY1, ZC3H11A, ZNF143, ZNF263, ZNF384 |
| *HAUS7* | EGR1, TET1, SRY, SALL4, SOX17, RUNX2, YY1, ELF1, ATF1, ATF3, BACH1, BHLHE40, BRCA1, CBX3, CCNT2, CHD1, CHD2, CREB1, CTCF, CTCFL, E2F4, E2F6, ELK1, EP300, ETS1, EZH2, FOS, FOXP2, GABPA, GATA1, GTF2F1, H2AFZ, HCFC1, HDAC1, HDAC2, HDAC6, HMGN3, IRF1, JUN, JUND, KDM4A, KDM5A, KDM5B, MAFK, MAX, MAZ, MXI1, MYC, NELFE, NFYA, NFYB, NR2F2, NRF1, PAX5, PBX3, PHF8, POLR2A, RAD21, RBBP5, RCOR1, REST, RFX5, RUNX3, SAP30, SIN3A, SIRT6, SMC3, SP1, SP2, SPI1, SRF, STAT1, STAT3, TAF1, TAF7, TBL1XR1, TBP, TCF12, TEAD4, UBTF, USF2, WHSC1, ZBTB7A, ZC3H11A, ZKSCAN1, ZMIZ1, ZNF143, ZNF263, ZNF384 |
| *ELOVL2* | ZNF217, SMAD4, CREM, SOX2, EGR1, TP53, FOXA2, HNF4A, REST, FOXP1, ERG, OLIG2, YAP1, SMAD3, CTCF, BACH1, SREBF2, NR1I2, HTT, ARID3A, ATF3, BHLHE40, BRCA1, CBX3, CBX8, CCNT2, CEBPB, CEBPD, CHD1, CHD2, CHD7, CREB1, CTBP2, CTCFL, E2F6, ELF1, EP300, ESR1, ETS1, EZH2, FOSL2, FOXA1, FOXP2, GABPA, GATA3, GTF2F1, H2AFZ, HCFC1, HDAC1, HDAC2, HMGN3, HNF4G, JUN, JUND, KDM4A, KDM5B, MAFK, MAX, MAZ, MXI1, MYBL2, MYC, NANOG, NFIC, NR3C1, NRF1, PAX5, PHF8, POLR2A, RAD21, RBBP5, RCOR1, RFX5, RUNX3, RXRA, SAP30, SETDB1, SIN3A, SMC3, SP4, SUZ12, TBL1XR1, TBP, TCF12, TCF7L2, TEAD4, TRIM28, UBTF, USF2, WRNIP1, YY1, ZBTB7A, ZC3H11A, ZMIZ1, ZNF143, ZNF263, ZNF384 |
| *TCTN2* | KDM5A, AR, FLI1, PPARG, E2F4, RCOR3, SIN3B, YY1, SRF, PDX1, BHLHE40, CEBPB, CHD1, CHD2, CREB1, CTCF, CUX1, E2F6, ELF1, ELK1, EP300, ETS1, EZH2, FOS, GTF2F1, H2AFZ, HCFC1, HDAC1, HDAC2, HMGN3, IRF1, IRF3, JUND, KAT2B, KDM4A, KDM5B, MAFF, MAFK, MAX, MAZ, MXI1, MYC, NFIC, NFYA, NFYB, PAX5, PHF8, POLR2A, RBBP5, RCOR1, REST, RFX5, RUNX3, SAP30, SIN3A, SMC3, SP1, SP2, SP4, TAF1, TBL1XR1, TBP, UBTF, WHSC1, WRNIP1, ZC3H11A, ZKSCAN1, ZMIZ1, ZNF143, ZNF384 |
| *LRP1B* | AHR, ARNT, ZNF217, PAX3, AR, STAT3, TCF4, SMAD4, TP63, NANOG, SOX2, TCF3, HNF4A, REST, TFAP2C, OLIG2, SMARCA4, TEAD4, PRDM14, KLF1, NR3C1, RUNX2, MTF2, SUZ12, POU3F2, SREBF1, ZNF322, BACH1, BRCA1, CEBPB, CHD1, CHD2, CHD7, CTBP2, CTCF, E2F4, E2F6, EP300, EZH2, FOXA1, FOXA2, GABPA, GTF2F1, H2AFZ, HDAC2, HNF4G, JUND, KDM4A, KDM5A, KDM5B, MAFF, MAFK, MAX, MAZ, MXI1, MYC, PHF8, POLR2A, PRDM1, RAD21, RBBP5, RCOR1, RFX5, SAP30, SIN3A, SMARCB1, SMARCC1, SMARCC2, SMC3, TAF1, TBP, TCF12, TCF7L2, TRIM28, USF2, WRNIP1, YY1, ZKSCAN1, ZNF143, ZNF263 |
| *B4GALT5* | TFAP2A, USF1, USF2, DACH1, AR, STAT3, TCF4, CREM, FLI1, KLF4, NANOG, POU5F1, SOX2, TCF3, EGR1, MITF, HNF4A, GATA1, EOMES, TFAP2C, FOXP1, PPARD, TAL1, EP300, ESR1, OLIG2, SALL4, ZNF281, PRDM14, KLF1, TFCP2L1, NR3C1, RUNX2, STAT4, ESRRB, CTNNB1, RELA, GATA3, ARID3A, ATF2, ATF3, BACH1, BATF, BCL3, BCLAF1, BHLHE40, BRCA1, CBX3, CCNT2, CEBPB, CEBPD, CHD1, CHD2, CHD4, CHD7, CREB1, CTCF, CUX1, E2F1, E2F4, E2F6, EBF1, ELF1, ELK1, ESRRA, ETS1, EZH2, FOS, FOSL1, FOSL2, FOXA1, FOXP2, GATA2, GTF2F1, H2AFZ, HCFC1, HDAC1, HDAC2, HDAC6, HMGN3, HNF4G, IKZF1, IRF1, IRF3, JUN, JUND, KDM1A, KDM4A, KDM5A, KDM5B, MAFF, MAFK, MAX, MAZ, MBD4, MTA3, MXI1, MYBL2, MYC, MYOD1, MYOG, NELFE, NFIC, NR2F2, NRF1, PAX5, PHF8, PML, POLR2A, POU2F2, RAD21, RBBP5, RCOR1, REST, RFX5, RUNX3, SAP30, SIN3A, SIRT6, SMARCB1, SMC3, SP1, SP4, SPI1, STAT1, STAT5A, SUPT20H, SUZ12, TAF1, TBL1XR1, TBP, TCF12, TCF7L2, TEAD4, UBTF, WRNIP1, YY1, ZBTB33, ZBTB7A, ZC3H11A, ZKSCAN1, ZMIZ1, ZNF143, ZNF384 |
| *PSMB9* | STAT1, ELK1, SOX2, SETDB1, TET1, MYCN, CNOT3, CUX1, STAT4, EZH2, MTF2, SUZ12, RELA, IRF1, IRF8, BHLHE40, CHD1, CHD2, CTCF, E2F4, EP300, ETS1, GATA1, HCFC1, KAT2A, MAFK, MAX, MAZ, MXI1, MYC, MYOG, NELFE, POLR2A, RCOR1, REST, SIN3A, SMC3, TAL1, TBP, UBTF, USF2, ZMIZ1 |
| *MDK* | HIF1A, WT1, STAT3, E2F1, MYC, KLF4, NANOG, POU5F1, SOX2, TCF3, EGR1, TP53, ZFX, SETDB1, EOMES, FOXP1, TET1, SOX9, SRY, ZNF281, CCND1, SIN3A, RELA, BACH1, CEBPB, IRF1, ARID3A, ATF2, ATF3, BCL3, BCLAF1, BHLHE40, CBX3, CCNT2, CEBPD, CHD1, CHD2, CHD4, CHD7, CREB1, CTBP2, CTCF, E2F4, E2F6, EBF1, ELF1, EP300, ETS1, EZH2, FOSL1, FOSL2, FOXA1, FOXA2, FOXM1, FOXP2, GTF2F1, H2AFZ, HCFC1, HDAC1, HDAC2, HDAC6, HMGN3, HNF4A, HNF4G, JUN, JUND, KAT2A, KDM1A, KDM4A, KDM5A, KDM5B, MAFF, MAFK, MAX, MAZ, MBD4, MTA3, MXI1, MYBL2, NCOR1, NFATC1, NFIC, NR2F2, PBX3, PHF8, PML, POLR2A, PRDM1, RAD21, RBBP5, RCOR1, REST, RFX5, RNF2, RUNX3, RXRA, SAP30, SMARCB1, SMC3, SP1, SP4, STAT5A, SUZ12, TAF1, TAF7, TAL1, TBL1XR1, TBP, TCF12, TCF7L2, TEAD4, TFAP2C, TRIM28, UBTF, USF1, USF2, YY1, ZBTB7A, ZKSCAN1, ZMIZ1, ZNF143, ZNF263, ZNF384 |

| | |
|---|---|
| *CD74* | E2F1, TP63, CREB1, FLI1, MITF, HNF4A, REST, SRY, JUN, RELA, TBP, IRF8, NR1H3, ATF1, ATF2, BATF, BCL11A, BCL3, BCLAF1, BHLHE40, BRCA1, CEBPB, CEBPD, CHD1, CHD2, CTBP2, CTCF, CUX1, E2F4, EBF1, EGR1, ELF1, ELK1, EP300, ETS1, EZH2, FOXM1, GATA1, GTF2F1, H2AFZ, HCFC1, HDAC2, HDAC6, IKZF1, IRF3, IRF4, JUND, KAT2A, KDM4A, KDM5A, MAFK, MAX, MAZ, MBD4, MEF2A, MEF2C, MTA3, MXI1, MYBL2, MYC, NELFE, NFATC1, NFE2, NFIC, NFYA, NFYB, NRF1, PAX5, PBX3, PML, POLR2A, POU2F2, RAD21, RBBP5, RCOR1, RFX5, RUNX3, SIN3A, SMC3, SP1, SPI1, SRF, STAT1, STAT3, STAT5A, TAF1, TBL1XR1, TCF12, TCF3, TCF7L2, TEAD4, UBTF, USF2, WRNIP1, YY1, ZBTB33, ZBTB7A, ZEB1, ZKSCAN1, ZNF143, ZNF384 |
| *GPR146* | STAT3, TP63, CREB1, MITF, TP53, PPARG, GATA1, SETDB1, TET1, TAL1, SOX9, SRY, EP300, SALL4, WT1, KLF1, TFCP2L1, STAT4, GATA4, FOXP2, MTF2, BCL3, RBPJ, RARG, ATF1, BACH1, BHLHE40, CCNT2, CEBPB, CHD1, CHD2, CHD7, CTCF, CUX1, EBF1, ELF1, ETS1, EZH2, GATA2, H2AFZ, HDAC1, HDAC2, HMGN3, JUND, KDM1A, KDM5B, MAX, MAZ, MXI1, MYC, MYOD1, MYOG, PAX5, POLR2A, RBBP5, RCOR1, RFX5, SIN3A, SIRT6, TBL1XR1, TCF12, TCF3, UBTF, USF1, USF2, YY1, ZMIZ1, ZNF384 |
| *CD200R1L* | AR, STAT3, SMAD4, GATA1, REST, RCOR1, PAX6, NR4A2, CEBPB, EBF1, EP300, EZH2, GATA3, H2AFZ, IRF1, JUND, KDM1A, TAL1, TCF12, TCF3 |
| *SLC46A1* | AR, TP63, CREB1, RUNX1, MYC, MITF, HNF4A, GATA1, E2F4, SIN3B, SETDB1, EOMES, TFAP2C, ERG, SMARCA4, MYB, SREBF2, ARID3A, ATF2, BACH1, BCL3, BCLAF1, BHLHE40, BRCA1, CBX3, CCNT2, CEBPB, CEBPD, CHD1, CHD2, CHD7, CTCF, E2F6, EBF1, EGR1, ELF1, ELK1, EP300, ETS1, EZH2, FOS, FOXA1, FOXM1, GTF2F1, H2AFZ, HCFC1, HDAC1, HDAC2, HDAC6, HMGN3, HNF4G, JUN, JUND, KAT2A, KDM4A, KDM5A, KDM5B, MAFF, MAFK, MAX, MAZ, MTA3, MXI1, MYBL2, MYOG, NELFE, NFIC, NR2F2, NRF1, PAX5, PHF8, PML, POLR2A, RAD21, RBBP5, RCOR1, REST, RFX5, RUNX3, SAP30, SIN3A, SMARCB1, SMC3, SP1, STAT1, STAT3, SUZ12, TAF1, TBL1XR1, TBP, TCF12, TCF3, TEAD4, TRIM28, UBTF, USF1, USF2, WHSC1, YY1, ZBTB7A, ZEB1, ZKSCAN1, ZMIZ1, ZNF143, ZNF263 |
| *CARNS1* | XRN2, SPI1, PPARG, SIN3B, ERG, EP300, YY1, MEIS1, CEBPB, CHD1, CTCF, EBF1, EGR1, EZH2, FOXA1, FOXA2, FOXP2, GATA1, H2AFZ, HDAC1, HDAC2, HDAC6, KDM4A, MAX, MYC, MYOD1, MYOG, NFIC, PHF8, POLR2A, RBBP5, RCOR1, REST, SAP30, SIN3A, SP1, TAL1, TBP, ZBTB7A, ZKSCAN1, ZNF384 |
| *CRISPLD1* | AR, SMAD4, MYC, NANOG, POU5F1, TCF3, TFCP2L1, RUNX2, FOXP2, RCOR1, EWSR1, GBX2, ATF3, BACH1, BRCA1, CHD1, CHD2, CHD7, CREB1, CTCF, E2F6, EP300, EZH2, GATA3, GTF2F1, H2AFZ, HDAC2, HDAC6, JUND, KDM4A, MAFF, MAFK, MAX, NR3C1, NRF1, PHF8, POLR2A, RAD21, RBBP5, REST, SAP30, SIN3A, SP1, TAF1, TAF7, TBP, TCF12, TEAD4, TRIM28, USF1, USF2, YY1, ZBTB7A, ZNF143, ZNF263 |
| *ERICH3* | SMAD4, SOX2, EGR1, HNF4A, KDM5B, REST, DMRT1, RUNX2, RCOR1, NR0B1, ZNF652, CHD1, CHD2, CTCF, EP300, EZH2, H2AFZ, HDAC2, KDM4A, KDM5A, MAX, NANOG, POLR2A, RAD21, RBBP5, RELA, RUNX3, SIN3A, SPI1, STAT1, TBP, ZNF143 |
| *CLEC12A* | STAT3, RUNX1, SPI1, NANOG, POU5F1, SOX2, TCF3, SETDB1, MECOM, LMO2, CEBPB, IRF8, BHLHE40, CHD2, CTCF, EBF1, EP300, ETS1, EZH2, GATA1, H2AFZ, JUN, MAFK, MAX, MYC, NFYB, RBBP5, RCOR1, RUNX3, TAL1, TBP |
| *LRRC61* | PAX3, TTF2, CREM, E2F1, FLI1, RUNX1, KLF4, POU5F1, EGR1, HNF4A, RCOR3, SIN3B, REST, SETDB1, TFAP2C, PPARD, GFI1B, ESR1, SALL4, TFCP2L1, SMAD2, SMAD3, GATA4, MEF2A, SIN3A, HSF1, BCL3, CEBPA, CEBPB, MYBL1, ARID3A, ATF1, ATF2, ATF3, BACH1, BCLAF1, BHLHE40, BRCA1, CBX2, CBX3, CCNT2, CEBPD, CHD1, CHD2, CHD7, CTCF, CTCFL, CUX1, E2F4, E2F6, EBF1, ELF1, ELK1, ELK4, EP300, ETS1, EZH2, FOS, FOXA1, FOXA2, FOXM1, FOXP2, GABPA, GATA2, GATA3, GTF2B, GTF2F1, H2AFZ, HCFC1, HDAC2, HMGN3, HNF4G, IRF1, JUN, JUND, KDM1A, KDM4A, KDM5A, KDM5B, MAFF, MAFK, MAX, MAZ, MBD4, MTA3, MXI1, MYBL2, MYC, NCOR1, NELFE, NFATC1, NFIC, NR2F2, NRF1, PAX5, PHF8, PML, POLR2A, POU2F2, RAD21, RBBP5, RCOR1, RELA, RFX5, RNF2, RUNX3, RXRA, SAP30, SIX5, SMARCA4, SMARCB1, SMARCC1, SMARCC2, SMC3, SP1, SPI1, SREBF1, SRF, STAT1, STAT3, STAT5A, TAF1, TAL1, TBL1XR1, TBP, TCF12, TCF3, TCF7L2, TEAD4, THAP1, TRIM28, UBTF, USF1, USF2, WRNIP1, YY1, ZBTB7A, ZC3H11A, ZEB1, ZKSCAN1, ZMIZ1, ZNF143, ZNF263, ZNF274, ZNF384 |
| *PPT1* | HIF1A, KDM5A, TTF2, AR, STAT3, E2F1, RUNX1, MYC, SPI1, NANOG, POU5F1, SOX2, TCF3, MITF, E2F4, SIN3B, ERG, FOXO3, TEAD4, CUX1, HOXB4, YY1, GATA4, SRF, BCL3, CEBPB, GATA3, TFEB, DCP1A, KDM6A, IRF8, ATF2, ATF3, BACH1, BATF, BCLAF1, BHLHE40, BRCA1, CBX3, CCNT2, CEBPD, CHD1, CHD2, CHD7, CREB1, CREBBP, CTBP2, CTCF, E2F6, ELF1, ELK1, EP300, ESR1, ETS1, EZH2, FOS, FOXM1, FOXP2, GABPA, GATA2, GTF2B, GTF2F1, H2AFZ, HCFC1, HDAC1, HDAC2, HDAC6, HMGN3, IRF1, JUN, JUND, KAT2A, KDM1A, KDM4A, KDM5B, MAFK, MAX, MAZ, MTA3, MXI1, MYBL2, MYOD1, MYOG, NELFE, NFATC1, NFIC, NFYB, NRF1, PAX5, PHF8, PML, POLR2A, POU2F2, RAD21, RBBP5, RCOR1, REST, RFX5, RUNX3, SAP30, SETDB1, SIN3A, SIRT6, SIX5, SMARCB1, SMARCC1, SMC3, SP1, SP4, STAT1, STAT5A, TAF1, TAF7, TAL1, TBL1XR1, TBP, TCF12, TCF7L2, THAP1, TRIM28, UBTF, USF1, USF2, WRNIP1, ZBTB33, ZBTB7A, ZC3H11A, ZEB1, ZKSCAN1, ZMIZ1, ZNF143, ZNF384 |
| *ZSWIM5* | STAT3, TCF4, SMAD4, CREM, MYC, TCF3, MITF, FOXA2, HNF4A, GATA2, REST, SETDB1, MYCN, TRIM28, SALL4, SMAD3, SCLY, MTF2, SUZ12, ARID3A, BACH1, BHLHE40, BRCA1, CBX2, CCNT2, CEBPB, CEBPD, CHD1, CHD2, CHD7, CREB1, CTBP2, CTCF, E2F4, E2F6, EGR1, ELF1, ELK1, EP300, EZH2, FOXA1, FOXP2, GATA3, GTF2F1, H2AFZ, HCFC1, HDAC1, HDAC2, HMGN3, JUN, JUND, KDM4A, KDM5A, KDM5B, MAFK, MAX, MAZ, MTA3, MXI1, NELFE, NRF1, PAX5, PHF8, PML, POLR2A, RAD21, RBBP5, RCOR1, RFX5, RUNX3, SAP30, SIN3A, SMARCB1, SMC3, TAF1, TAF7, TBL1XR1, TBP, TCF12, TCF7L2, TEAD4, UBTF, WHSC1, YY1, ZBTB7A, ZKSCAN1, ZNF143, ZNF263, ZNF384 |

| | |
|---|---|
| *ID2* | ATF1, CREB1, FLI1, HIF1A, MYC, SMAD1, ZNF217, TTF2, XRN2, PHF8, AR, STAT3, TCF4, SMAD4, TP63, CREM, E2F1, RUNX1, SPI1, KLF4, NANOG, POU5F1, SOX2, EGR1, MITF, TP53, PPARG, HNF4A, GATA1, GATA2, RCOR3, SIN3B, SETDB1, EOMES, TFAP2C, FOXP1, PPARD, TRIM28, SOX9, SRY, EP300, ESR1, OLIG2, SMARCA4, YAP1, SALL4, ZNF281, TEAD4, NR3C1, RUNX2, CCND1, HOXB4, GATA4, RAD21, MTF2, SUZ12, CTNNB1, TBX3, JUN, NUCKS1, STAT5A, NR0B1, NR1I2, DCP1A, PDX1, PAX6, ARID3A, ATF2, ATF3, BACH1, BATF, BCL11A, BCL3, BCLAF1, BHLHE40, BRCA1, CBX3, CCNT2, CEBPB, CEBPD, CHD1, CHD2, CHD4, CHD7, CTCF, CUX1, E2F4, E2F6, EBF1, ELF1, ELK1, ELK4, ESRRA, ETS1, EZH2, FOS, FOSL2, FOXA1, FOXA2, FOXM1, FOXP2, GABPA, GATA3, GTF2B, GTF2F1, H2AFZ, HCFC1, HDAC1, HDAC2, HMGN3, HNF4G, IKZF1, IRF1, IRF3, IRF4, JUND, KAT2A, KAT2B, KDM1A, KDM4A, KDM5B, MAFF, MAFK, MAX, MAZ, MBD4, MTA3, MXI1, MYBL2, NELFE, NFATC1, NFE2, NFIC, NFYA, NFYB, NRF1, PAX5, PBX3, PML, POLR2A, POU2F2, PRDM1, RCOR1, RELA, REST, RFX5, RNF2, RUNX3, SAP30, SIN3A, SIRT6, SIX5, SMC3, SP1, SP2, SP4, SRF, STAT1, TAF1, TAF7, TAL1, TBL1XR1, TBP, TCF12, TCF3, TCF7L2, THAP1, UBTF, USF1, USF2, WHSC1, WRNIP1, YY1, ZBTB33, ZBTB7A, ZC3H11A, ZEB1, ZKSCAN1, ZMIZ1, ZNF143, ZNF263, ZNF384 |
| *FSTL5* | ZNF217, AR, STAT3, TCF4, SMAD4, TP63, RUNX1, NANOG, POU5F1, SOX2, RCOR3, SIN3B, REST, FOXP1, PPARD, NR3C1, RUNX2, HOXB4, TBX3, JUN, POU3F2, BACH1, CEBPB, CHD1, CHD2, CHD7, CREB1, CTCF, E2F4, EP300, EZH2, FOS, FOXP2, GATA3, H2AFZ, HDAC2, JUND, KDM4A, MAX, MXI1, MYC, PAX5, PBX3, PHF8, POLR2A, POU2F2, PRDM1, RAD21, RBBP5, RCOR1, SAP30, SETDB1, SIN3A, SIRT6, SP1, SP4, TAF1, TBP, TCF12, TCF7L2, TEAD4, TRIM28, USF2, ZNF143, ZNF263 |
| *C9orf3* | NFE2L2, STAT3, CREB1, NANOG, TP53, SIN3B, REST, ASH2L, SOX9, EP300, TEAD4, MYBL2, GATA4, MEF2A, TBX5, TCF7, SIN3A, CEBPB, NR1I2 |
| *SELL* | FOXP3, FLI1, SALL4, CUX1, HOXB4, MECOM, CTCF, MYB, ELF5, CEBPB, ARID3A, BCLAF1, BHLHE40, CHD1, CHD2, EBF1, ELF1, ELK1, EP300, EZH2, FOXA1, FOXP2, GATA1, GATA3, GTF2F1, H2AFZ, HDAC2, IKZF1, IRF4, KDM5A, MAFF, MAFK, MAX, MAZ, MTA3, MXI1, MYC, PAX5, POLR2A, POU2F2, RAD21, RBBP5, RCOR1, RELA, SAP30, SMC3, SPI1, STAT1, TBL1XR1, TBP, TRIM28, UBTF, USF2, WRNIP1, YY1, ZC3H11A, ZNF143, ZNF263 |
| *SEMA3B* | AHR, ARNT, TP63, CREB1, CREM, NANOG, POU5F1, EGR1, MITF, HNF4A, TFAP2C, FOXP1, ERG, SOX9, SRY, EP300, ATF3, YY1, STAT4, TBX5, TCF7, RAD21, SUZ12, CEBPB, SREBF2, RBPJ, CRX, ESR2, BACH1, BCL3, BHLHE40, BRCA1, CEBPD, CHD2, CTCF, E2F4, ELF1, ELK1, ELK4, ESR1, ETS1, EZH2, FOS, FOSL2, FOXA2, FOXM1, GABPA, GATA1, GATA3, GTF2F1, H2AFZ, HDAC1, HDAC2, HMGN3, HNF4G, JUND, KDM4A, KDM5A, MAFK, MAX, MAZ, MXI1, MYBL2, MYC, NFATC1, NFIC, NFYA, NFYB, NR2F2, PBX3, PHF8, POLR2A, PRDM1, RBBP5, RCOR1, RFX5, SAP30, SIN3A, SIX5, SMARCB1, SMC3, SP1, STAT3, STAT5A, TAF1, TAL1, TBP, TCF12, TCF3, TCF7L2, TEAD4, TFAP2A, USF2, ZBTB33, ZBTB7A, ZKSCAN1, ZNF143, ZNF217 |
| *APIP* | NFE2L2, ELK1, AR, SMAD4, E2F1, SOX2, MITF, GATA2, E2F4, EOMES, MYCN, TAL1, TRIM28, FOXO3, CUX1, ZFP42, CTNNB1, PBX1, BCL3, DCP1A, KDM6A, PDX1, ARID3A, ATF1, ATF2, ATF3, BCLAF1, BHLHE40, BRCA1, CBX3, CCNT2, CEBPB, CEBPD, CHD1, CHD2, CHD7, CREB1, CTCF, E2F6, EBF1, EGR1, ELF1, ELK4, EP300, ETS1, EZH2, FOS, FOSL1, FOXA1, FOXM1, FOXP2, GABPA, GATA1, GATA3, GTF2F1, H2AFZ, HCFC1, HDAC1, HDAC2, HDAC6, HMGN3, IRF1, JUND, KAT2B, KDM1A, KDM4A, KDM5A, KDM5B, MAFK, MAX, MAZ, MTA3, MXI1, MYBL2, MYC, NANOG, NCOR1, NELFE, NFIC, NFYB, NR2F2, NR3C1, NRF1, PAX5, PBX3, PHF8, PML, POLR2A, POU2F2, RAD21, RBBP5, RCOR1, RELA, REST, RFX5, RUNX3, SAP30, SETDB1, SIN3A, SMARCB1, SMC3, SP1, SP4, SPI1, SRF, STAT1, STAT3, STAT5A, TAF1, TAF7, TBL1XR1, TBP, TCF12, TCF3, TCF7L2, TEAD4, UBTF, USF1, USF2, WRNIP1, YY1, ZBTB7A, ZC3H11A, ZEB1, ZKSCAN1, ZMIZ1, ZNF143, ZNF384 |
| *SLCO1B3* | STAT3, EGR1, POU3F2, BACH1, ARID3A, CEBPB, EP300, EZH2, FOXA1, FOXA2, H2AFZ, HDAC2, HNF4A, JUND, KDM5A, MYBL2, MYC, NFIC, NR2F2, POLR2A, RCOR1, RFX5, SP1, SPI1, TCF7L2, TEAD4 |
| *NMNAT2* | NFE2L2, VDR, STAT3, TCF4, SMAD4, TP63, NANOG, POU5F1, EGR1, MITF, TP53, PPARG, GATA1, GATA2, KDM5B, RCOR3, SIN3B, REST, SETDB1, FOXP1, TET1, ESR1, YAP1, ZNF281, PRDM14, NR3C1, CUX1, EED, EZH2, RNF2, SUZ12, PBX1, LMO2, LYL1, CDX2, ATF2, BACH1, BCL3, BHLHE40, CEBPB, CHD1, CHD2, CHD7, CREB1, CTCF, E2F6, EP300, GABPA, GATA3, GTF2F1, H2AFZ, HDAC2, HDAC6, JUN, JUND, KDM4A, KDM5A, MAX, MAZ, MXI1, MYC, MYOG, PHF8, POLR2A, RAD21, RBBP5, RCOR1, SAP30, SIN3A, SMC3, SP4, TAF1, TAL1, TBP, TCF12, TCF7L2, TEAD4, USF1, USF2, YY1, ZNF143, ZNF263 |
| *NR6A1* | ZNF217, CEBPD, ETS1, FOXP3, TTF2, AR, STAT3, TCF4, SMAD4, TP63, CREB1, CREM, E2F1, MYC, SPI1, NANOG, POU5F1, SOX2, TCF3, EGR1, MITF, TP53, PPARG, FOXA2, HNF4A, SIN3B, EOMES, TFAP2C, FOXP1, MYCN, ERG, FOXO3, EP300, DMRT1, CCND1, STAT4, PHC1, RNF2, CTNNB1, CDX2, THAP11, ARID3A, ATF1, ATF2, ATF3, BACH1, BCL3, BCLAF1, BHLHE40, BRCA1, CBX3, CCNT2, CEBPB, CHD1, CHD2, CHD7, CREBBP, CTBP2, CTCF, CTCFL, CUX1, E2F4, E2F6, EBF1, ELF1, ELK1, ELK4, EZH2, FOS, FOSL2, FOXA1, FOXM1, FOXP2, GABPA, GATA1, GATA2, GATA3, GTF2F1, H2AFZ, HCFC1, HDAC1, HDAC2, HDAC6, HMGN3, IRF1, IRF3, JUN, JUND, KAT2A, KAT2B, KDM1A, KDM4A, KDM5A, KDM5B, MAFF, MAFK, MAX, MAZ, MBD4, MTA3, MXI1, MYBL2, MYOG, NFATC1, NFE2, NFIC, NFYA, NFYB, NR2F2, NR3C1, NRF1, PAX5, PHF8, PML, POLR2A, POU2F2, RAD21, RBBP5, RCOR1, RELA, REST, RFX5, RUNX3, RXRA, SAP30, SETDB1, SIN3A, SIRT6, SIX5, SMARCB1, SMARCC2, SMC3, SP1, SP4, SREBF1, SRF, STAT1, STAT5A, SUPT20H, SUZ12, TAF1, TAF7, TBL1XR1, TBP, TCF12, TCF7L2, TEAD4, TFAP2A, TRIM28, UBTF, USF1, USF2, WHSC1, WRNIP1, YY1, ZBTB33, ZBTB7A, ZC3H11A, ZEB1, ZKSCAN1, ZMIZ1, ZNF143, ZNF263, ZNF384 |

| | |
|---|---|
| *STMN3* | TTF2, FLI1, RUNX1, MYC, KLF4, NANOG, POU5F1, SOX2, TCF3, EGR1, ZFX, RCOR3, SIN3B, REST, FOXP1, PPARD, TET1, ZNF281, SUZ12, SIN3A, MEIS1, ELF5, NR0B1, RCOR2, ARID3A, BACH1, BHLHE40, BRCA1, CBX3, CCNT2, CEBPB, CEBPD, CHD1, CHD2, CTBP2, CTCF, CTCFL, E2F4, E2F6, ELF1, ELK1, ELK4, EP300, ETS1, EZH2, FOS, FOSL1, FOXA1, FOXP2, GABPA, GTF2F1, H2AFZ, HCFC1, HDAC2, HDAC6, HMGN3, HNF4A, HNF4G, IRF1, JUN, JUND, KDM4A, KDM5A, KDM5B, MAFK, MAX, MAZ, MXI1, MYBL2, MYOG, NCOR1, NFIC, NR2F2, PAX5, PBX3, PHF8, PML, POLR2A, RAD21, RBBP5, RCOR1, RFX5, RUNX3, SAP30, SETDB1, SMC3, SP1, SP4, SPI1, STAT3, STAT5A, TAF1, TAF7, TAL1, TBL1XR1, TBP, TCF12, TCF7L2, TEAD4, THAP1, TRIM28, UBTF, USF1, YY1, ZBTB7A, ZC3H11A, ZKSCAN1, ZMIZ1, ZNF143, ZNF263, ZNF384 |
| *FAM50B* | VDR, STAT3, SMAD4, TP63, FLI1, MYC, SPI1, SOX2, EGR1, PPARG, FOXA2, HNF4A, ASH2L, PPARD, EP300, NR3C1, GATA4, RAD21, JUN, EWSR1, SREBF2, ZNF274, ARID3A, ATF2, BACH1, BATF, BCLAF1, BHLHE40, BRCA1, CBX3, CCNT2, CEBPB, CHD1, CHD2, CHD7, CREB1, CTCF, CTCFL, E2F4, E2F6, EBF1, ELF1, ELK1, ELK4, ETS1, EZH2, FOS, FOXA1, FOXM1, FOXP2, GABPA, GATA3, GTF2F1, H2AFZ, HCFC1, HDAC1, HDAC2, HMGN3, JUND, KDM4A, MAFK, MAX, MAZ, MTA3, MXI1, NANOG, NFATC1, NFIC, NFYA, NFYB, NRF1, PAX5, PHF8, PML, POLR2A, POU2F2, RBBP5, RCOR1, RFX5, RUNX3, SAP30, SETDB1, SIN3A, SIRT6, SMARCB1, SMARCC1, SMC3, SP1, SP4, STAT1, SUZ12, TAF1, TBL1XR1, TBP, TCF12, TCF3, TCF7L2, TEAD4, THAP1, TRIM28, YY1, ZBTB33, ZC3H11A, ZEB1, ZKSCAN1, ZMIZ1, ZNF143 |
| *C9orf116* | KDM5A, CREM, SPI1, NANOG, POU5F1, TP53, TET1, HSF1, TBP, PAX6, CTCF, E2F4, EP300, GABPA, HCFC1, KAT2A, MAX, MXI1, MYC, MYOG, NELFE, POLR2A, RAD21, REST, SIN3A, USF1, USF2, ZMIZ1 |
| *SUCNR1* | ZNF217, CEBPD, STAT3, FLI1, RUNX1, SOX2, PPARG, SIN3B, REST, TAL1, TRIM28, SMARCA4, SCLY, LMO2, POU3F2, RCOR1, CDX2, ATF1, ATF2, BATF, BHLHE40, CBX3, CHD1, CUX1, E2F6, ELF1, EP300, ETS1, EZH2, FOS, GABPA, GATA1, H2AFZ, HDAC2, IKZF1, JUN, JUND, KDM5B, MAX, MAZ, MTA3, MXI1, MYC, NFIC, NRF1, PHF8, POLR2A, RFX5, RUNX3, SAP30, SETDB1, SPI1, TAF1, TBP, TEAD4, UBTF, USF2, WHSC1, WRNIP1, ZNF143 |
| *PSMB10* | VDR, TP63, CREB1, FLI1, MYC, EGR1, PPARG, HNF4A, SIN3B, SETDB1, SRY, ESR1, DMRT1, CCND1, TBX5, SIN3A, STAT5A, SMAD1, IRF8, ARID3A, ATF2, ATF3, BACH1, BCL3, BCLAF1, BHLHE40, BRCA1, CBX3, CCNT2, CEBPB, CEBPD, CHD1, CHD2, CHD4, CHD7, CTCF, CUX1, E2F4, E2F6, EBF1, ELF1, ELK1, ELK4, EP300, ETS1, EZH2, FOS, FOSL2, FOXM1, FOXP2, GABPA, GATA1, GATA2, GATA3, GTF2B, GTF2F1, H2AFZ, HCFC1, HDAC1, HDAC2, HDAC6, HMGN3, IRF1, JUN, JUND, KAT2A, KDM4A, KDM5A, KDM5B, MAFF, MAFK, MAX, MAZ, MBD4, MTA3, MXI1, MYBL2, MYOG, NANOG, NELFE, NFATC1, NFE2, NFIC, NFYA, NFYB, NR2F2, NRF1, PAX5, PBX3, PHF8, PML, POLR2A, POU2F2, PRDM1, RAD21, RBBP5, RCOR1, RELA, REST, RFX5, RUNX3, SAP30, SIX5, SMARCB1, SMC3, SP1, SP2, SP4, SPI1, STAT1, STAT3, TAF1, TBL1XR1, TBP, TCF12, TCF3, TCF7L2, TEAD4, UBTF, USF1, USF2, WHSC1, WRNIP1, YY1, ZBTB33, ZBTB7A, ZC3H11A, ZEB1, ZKSCAN1, ZMIZ1, ZNF143, ZNF263, ZNF384 |
| *EPCAM* | MYB, TP53, TP63, CREM, E2F1, RUNX1, SPI1, NANOG, POU5F1, SOX2, PPARG, HNF4A, GATA2, KDM5B, REST, SETDB1, ASH2L, EOMES, TFAP2C, SOX9, MYBL2, SUZ12, CDX2, GATA3, ATF2, ATF3, BACH1, BHLHE40, CBX3, CCNT2, CEBPD, CHD1, CHD2, CHD7, CREB1, CTBP2, CTCF, E2F4, E2F6, EGR1, ELF1, EP300, ESR1, ETS1, EZH2, FOS, FOSL1, FOSL2, FOXA1, GABPA, GTF2F1, H2AFZ, HCFC1, HDAC1, HDAC2, HDAC6, HMGN3, JUN, JUND, KDM4A, KDM5A, MAFK, MAX, MAZ, MXI1, MYC, MYOG, NFIC, NR2F2, NRF1, PHF8, PML, POLR2A, RAD21, RBBP5, RCOR1, RFX5, RXRA, SAP30, SIN3A, SIRT6, SMARCB1, SMARCC1, SP1, SP4, SRF, TAF1, TAF7, TBP, TCF12, TCF7L2, TEAD4, UBTF, USF1, YY1, ZBTB7A, ZEB1, ZNF143, ZNF217, ZNF263, ZNF384 |
| *RPRM* | AHR, ARNT, ZNF217, PAX3, CEBPD, NFE2L2, TTF2, AR, SMAD4, TP63, RUNX1, MYC, KLF4, NANOG, POU5F1, SOX2, TCF3, TP53, ZFX, REST, SETDB1, FOXP1, PPARD, SOX9, SRY, ESR1, OLIG2, ZNF281, SMAD3, RUNX2, CTCF, BMI1, RNF2, MTF2, SUZ12, JUN, PBX1, POU3F2, RCOR1, ARID3A, BACH1, BCLAF1, BHLHE40, BRCA1, CBX3, CCNT2, CEBPB, CHD1, CHD2, CHD7, CREB1, CTBP2, E2F4, E2F6, EBF1, EGR1, ELF1, ELK1, EP300, EZH2, FOS, FOXA2, FOXP2, GABPA, GTF2F1, H2AFZ, HDAC2, HMGN3, JUND, KDM4A, KDM5A, KDM5B, MAFF, MAFK, MAX, MAZ, MXI1, NFIC, NR2F2, PAX5, PHF8, POLR2A, RAD21, RBBP5, RFX5, SAP30, SIN3A, SIRT6, SMARCB1, SMC3, SP4, SPI1, STAT1, TAF1, TBL1XR1, TBP, TCF12, TCF7L2, TEAD4, TFAP2A, TFAP2C, TRIM28, UBTF, USF1, USF2, YY1, ZBTB7A, ZC3H11A, ZKSCAN1, ZMIZ1, ZNF143, ZNF263, ZNF384 |
| *ERP27* | ELK1, TP63, SOX2, MITF, TP53, ASH2L, TAL1, SOX17, PAX6, CHD2, CTCF, EP300, FOS, H2AFZ, KDM5A, NELFE, NFATC1, PML, POLR2A, POU2F2, RAD21, REST, RUNX3, SMC3, SPI1, YY1, ZMIZ1, ZNF143, ZNF384 |
| *SLC29A3* | STAT3, TP63, FLI1, KLF4, POU5F1, SOX2, FOXA2, HNF4A, ZFX, SIN3B, EOMES, TFAP2C, PPARD, TET1, FOXO3, SMARCA4, SALL4, NR3C1, SRF, CTCF, EWSR1, CEBPB, NR1H3, ARID3A, ATF3, BACH1, BCL3, BCLAF1, BHLHE40, BRCA1, CBX3, CCNT2, CHD1, CHD2, CHD4, CHD7, CREB1, CTCFL, CUX1, E2F4, E2F6, EGR1, ELF1, ELK1, EP300, ETS1, EZH2, FOS, FOSL2, FOXA1, FOXM1, FOXP2, GABPA, GTF2B, GTF2F1, H2AFZ, HCFC1, HDAC1, HDAC2, HMGN3, HNF4G, IRF1, JUN, JUND, KDM4A, KDM5A, KDM5B, MAFF, MAFK, MAX, MAZ, MTA3, MXI1, MYBL2, MYC, MYOG, NCOR1, NELFE, NFIC, NFYA, NR2F2, NRF1, PAX5, PHF8, PML, POLR2A, RAD21, RBBP5, RCOR1, RELA, REST, RFX5, RNF2, RUNX3, SAP30, SIN3A, SMC3, SP1, SPI1, STAT1, STAT5A, TAF1, TAL1, TBL1XR1, TBP, TCF12, TCF3, TCF7L2, TEAD4, TRIM28, UBTF, USF1, USF2, WRNIP1, YY1, ZBTB7A, ZC3H11A, ZKSCAN1, ZMIZ1, ZNF143, ZNF263, ZNF384 |
| *SPATA13* | ZNF217, VDR, AR, STAT3, E2F1, RUNX1, EGR1, MITF, FOXA2, HNF4A, SIN3B, TET1, FOXO3, OLIG2, SMARCA4, TEAD4, KLF1, RUNX2, EZH2, TFAP2A, MYB, NR1I2, ARID3A, BACH1, BHLHE40, BRCA1, CCNT2, CEBPB, CHD1, CHD2, CTCF, E2F4, E2F6, EBF1, EP300, ESR1, FOS, FOXA1, FOXP2, GATA3, GTF2F1, H2AFZ, HCFC1, HDAC1, HDAC2, HDAC6, HMGN3, IRF1, JUND, KDM4A, KDM5B, MAFF, MAFK, MAX, MAZ, MXI1, MYC, MYOG, NELFE, NFIC, PBX3, PHF8, POLR2A, RAD21, RBBP5, RCOR1, REST, RFX5, RUNX3, SAP30, SIN3A, SIRT6, SMC3, SP1, SP4, SRF, STAT1, SUZ12, TAF1, TBP, TCF12, TCF7L2, UBTF, USF1, YY1, ZBTB7A, ZKSCAN1, ZMIZ1, ZNF143, ZNF263, ZNF384 |

| | |
|---|---|
| *DOK7* | TP63, CREB1, FLI1, SPI1, EGR1, MITF, TFAP2C, SOX9, ESR1, TEAD4, NR3C1, SCLY, CTCF, MTF2, SUZ12, CTNNB1, ESR2, CCNT2, CHD2, E2F4, EBF1, ELF1, EP300, EZH2, FOS, FOXA1, FOXM1, GABPA, GATA3, GTF2F1, H2AFZ, HDAC2, HDAC6, HNF4A, HNF4G, KDM4A, MAFF, MAFK, MAX, MAZ, MXI1, MYC, MYOD1, MYOG, NFIC, NR2F2, PAX5, POLR2A, RAD21, RBBP5, RCOR1, REST, RFX5, RUNX3, SIN3A, SMC3, SRF, STAT3, TAF1, TBP, TCF12, TCF3, TCF7L2, TFAP2A, UBTF, YY1, ZBTB7A, ZNF217 |
| *KCNT2* | AR, TCF4, SMAD4, POU5F1, SOX2, FOXA2, HNF4A, PPARD, TET1, ESR1, SMARCA4, PRDM14, TFCP2L1, YY1, FOXP2, RELA, NR1I2, PADI4, IRF1, ATF2, ATF3, BACH1, CBX3, CCNT2, CEBPB, CHD1, CHD2, CHD7, CREB1, CTBP2, CTCF, E2F6, EGR1, EP300, EZH2, FOXA1, GATA1, GATA2, GATA3, GTF2F1, H2AFZ, HCFC1, HDAC2, JUN, JUND, KDM4A, KDM5A, MAFK, MAX, MAZ, MXI1, MYC, NANOG, PHF8, POLR2A, RAD21, RBBP5, RCOR1, REST, RFX5, SAP30, SETDB1, SIN3A, SIRT6, SP1, SP4, SUZ12, TAF1, TAF7, TBP, TCF12, TEAD4, TRIM28, USF2, ZBTB7A, ZNF143, ZNF384 |
| *MYPN* | AR, CREB1, NANOG, TP53, FOXA2, SIN3B, TET1, NR1I2, ESR2, BRCA1, CBX2, CBX3, CEBPB, CHD1, CHD2, CTCF, EGR1, EP300, EZH2, FOS, FOSL2, FOXA1, FOXP2, GATA1, GATA3, GTF2F1, H2AFZ, HMGN3, IKZF1, JUND, KDM5A, MAFK, MAX, MXI1, MYC, MYOD1, MYOG, NFIC, NR2F2, NR3C1, PBX3, POLR2A, RAD21, RCOR1, RFX5, RNF2, SAP30, SMC3, SP1, STAT3, TAF1, TBP, TCF12, TCF3, TCF7L2, TFAP2A, TFAP2C, TRIM28, USF1, USF2, YY1, ZNF384 |
| *PPM1H* | ZNF217, PAX3, NFE2L2, AR, STAT3, TCF4, SMAD4, FLI1, RUNX1, SPI1, KLF4, POU5F1, MITF, TP53, PPARG, FOXA2, HNF4A, GATA1, GATA2, E2F4, SIN3B, REST, SETDB1, ASH2L, TET1, MYCN, SOX9, ESR1, YAP1, DMRT1, TFCP2L1, SMAD3, RUNX2, CUX1, SCLY, TBX5, CTCF, EZH2, RNF2, JARID2, MTF2, SUZ12, CTNNB1, MYB, NR1I2, ASXL1, TFEB, CRX, NR1H3, ARID3A, ATF2, ATF3, BACH1, BCL3, BHLHE40, BRCA1, CBX2, CBX3, CBX8, CCNT2, CEBPB, CEBPD, CHD1, CHD2, CHD7, CREB1, CTBP2, CTCFL, E2F6, EGR1, ELF1, ELK1, ELK4, EP300, ETS1, FOSL2, FOXA1, FOXP2, GATA3, GTF2F1, H2AFZ, HCFC1, HDAC1, HDAC2, HDAC6, HMGN3, IRF1, JUN, JUND, KDM1A, KDM4A, KDM5A, KDM5B, MAFF, MAFK, MAX, MAZ, MBD4, MXI1, MYBL2, MYC, NANOG, NELFE, NFIC, NFYA, NFYB, NR2F2, NR3C1, PAX5, PHF8, PML, POLR2A, RAD21, RBBP5, RCOR1, RFX5, RUNX3, RXRA, SAP30, SIN3A, SIRT6, SMARCB1, SMC3, SP1, SP4, SRF, TAF1, TAF7, TAL1, TBL1XR1, TBP, TCF12, TCF3, TCF7L2, TEAD4, TRIM28, UBTF, USF1, USF2, YY1, ZBTB7A, ZC3H11A, ZEB1, ZKSCAN1, ZMIZ1, ZNF143, ZNF263, ZNF384 |
| *LCP1* | AR, ETS1, JUN, NFE2L2, AP1S2, TCF4, E2F1, FLI1, RUNX1, MYC, SPI1, SOX2, PPARG, FOXA2, HNF4A, GATA1, TFAP2C, SALL4, DMRT1, KLF1, YY1, MECOM, STAT4, BMI1, PBX1, MYB, POU3F2, STAT5A, CEBPA, CEBPB, PADI4, DNAJC2, ESR2, ARID3A, ATF2, BATF, BCL11A, BCL3, BCLAF1, BHLHE40, BRCA1, CBX3, CCNT2, CHD1, CHD2, CREB1, CTCF, CUX1, E2F4, E2F6, EBF1, EGR1, ELF1, ELK1, EP300, EZH2, FOXM1, GATA2, GATA3, GTF2B, GTF2F1, H2AFZ, HCFC1, HDAC1, HDAC2, HDAC6, HMGN3, IKZF1, IRF1, IRF3, IRF4, JUND, KAT2A, KDM5A, KDM5B, MAFF, MAFK, MAX, MAZ, MEF2A, MTA3, MXI1, NFATC1, NFE2, NFIC, NFYB, NRF1, PAX5, PBX3, PHF8, PML, POLR2A, POU2F2, RBBP5, RCOR1, RELA, REST, RFX5, RUNX3, RXRA, SAP30, SETDB1, SIN3A, SMC3, SP1, SRF, STAT1, STAT3, SUPT20H, TAF1, TAL1, TBL1XR1, TBP, TCF12, TCF3, TEAD4, TRIM28, UBTF, USF1, USF2, WRNIP1, ZC3H11A, ZEB1, ZMIZ1, ZNF143, ZNF263, ZNF384 |
| *FAM151A* | TP63, PPARG, RCOR3, REST, TET1, TEAD4, SREBF2, DNAJC2, ATF2, BATF, BCL11A, BHLHE40, CHD2, CTCF, EBF1, ELF1, EP300, EZH2, FOXM1, GATA1, GATA2, GATA3, H2AFZ, HDAC1, HDAC6, HNF4G, MTA3, NFIC, NR3C1, PAX5, POLR2A, RBBP5, RELA, RUNX3, SP1, SPI1, STAT5A, TCF12, ZNF143, ZNF384 |
| *TMEM191C* | CREM, SPI1, RCOR3, SUZ12, ATF1, BATF, BHLHE40, CBX3, CEBPD, CTCF, E2F4, E2F6, EBF1, EGR1, ELF1, ELK1, EP300, FOSL2, FOXA1, FOXP2, GABPA, GATA1, GATA2, H2AFZ, HCFC1, HDAC2, IRF4, JUND, KDM5B, MAX, MAZ, MXI1, MYC, MYOD1, MYOG, NELFE, NR2F2, PAX5, PBX3, PHF8, POLR2A, POU2F2, RAD21, RBBP5, RCOR1, REST, RXRA, SAP30, SIN3A, SIX5, SMC3, SP1, SRF, STAT1, STAT3, STAT5A, TAF1, TBP, TCF12, TCF3, TRIM28, UBTF, USF1, YY1, ZBTB33, ZBTB7A, ZC3H11A, ZEB1, ZNF143, ZNF384 |
| *PRTG* | AR, TCF4, E2F1, MYC, NANOG, POU5F1, SOX2, TCF3, MITF, GATA1, GATA2, REST, ASH2L, EOMES, ERG, DMRT1, TEAD4, PRDM14, TFCP2L1, TCF7, JARID2, MTF2, SUZ12, LMO2, RCOR1, ARID3A, BACH1, BCL3, BHLHE40, BRCA1, CBX8, CEBPB, CEBPD, CHD1, CHD2, CHD7, CREB1, CTBP2, CTCF, E2F4, E2F6, EGR1, ELF1, ELK1, EP300, ETS1, EZH2, FOSL2, FOXA1, FOXA2, FOXP2, GATA3, H2AFZ, HCFC1, HDAC1, HDAC2, HDAC6, HNF4A, HNF4G, JUND, KDM4A, KDM5B, MAFF, MAFK, MAX, MAZ, MEF2A, MXI1, MYOD1, MYOG, NFIC, NR2F2, PBX3, PHF8, POLR2A, RAD21, RBBP5, RFX5, RNF2, SAP30, SIN3A, SIX5, SMARCB1, SMC3, SP1, TAF1, TBP, TCF12, TCF7L2, TRIM28, UBTF, USF1, YY1, ZBTB33, ZBTB7A, ZKSCAN1, ZNF143, ZNF384 |
| *FUT9* | AR, STAT3, TCF4, E2F1, RUNX1, MYC, NANOG, POU5F1, SOX2, TCF3, TP53, HNF4A, RCOR3, REST, ASH2L, PPARD, SRY, OLIG2, SMARCA4, ZNF281, TFCP2L1, ESRRB, PBX1, POU3F2, RCOR1, NACC1, ATF2, BACH1, BRCA1, CEBPB, CHD1, CHD2, CHD7, CTBP2, CTCF, E2F6, EGR1, EP300, EZH2, FOSL1, GATA1, GATA2, GATA3, GTF2F1, H2AFZ, HCFC1, HDAC2, HDAC6, JUND, KDM4A, KDM5A, MAX, MXI1, MYOG, PHF8, POLR2A, RAD21, RBBP5, SAP30, SIN3A, SP4, SUZ12, TAF1, TAF7, TBP, TCF12, TEAD4, USF2, YY1, ZBTB7A, ZNF143 |
| *CDC42EP3* | ZNF217, NFE2L2, ELK1, AR, TCF4, SMAD4, TP63, FLI1, RUNX1, KLF4, NANOG, POU5F1, SOX2, TCF3, MITF, TP53, ASH2L, EOMES, PPARD, MYCN, ERG, TAL1, SRY, EP300, SALL4, SOX17, PRDM14, ATF3, SMAD2, SMAD3, NR3C1, RUNX2, SCLY, MTF2, SUZ12, JUN, MYB, RCOR1, NR0B1, SMAD1, STAT1, ETS2, HOXD13, ARID3A, ATF2, BACH1, BCL11A, BCL3, BCLAF1, BHLHE40, BRCA1, CBX2, CBX8, CCNT2, CEBPB, CHD1, CHD2, CHD7, CREB1, CTBP2, CTCF, CUX1, E2F4, E2F6, EBF1, EGR1, ELF1, ETS1, EZH2, FOS, FOSL2, FOXA2, FOXM1, FOXP2, GABPA, GATA1, GATA2, GATA3, GTF2F1, H2AFZ, HCFC1, HDAC1, HDAC2, HDAC6, HMGN3, IRF1, IRF4, JUND, KDM4A, KDM5B, MAFK, MAX, MAZ, MTA3, MXI1, MYC, MYOD1, MYOG, NELFE, NFATC1, NFE2, NFIC, NFYB, NR2F2, NRF1, PAX5, PBX3, PHF8, PML, POLR2A, POU2F2, RAD21, RBBP5, RELA, |

| | |
|---|---|
| | REST, RFX5, RNF2, RUNX3, RXRA, SAP30, SIN3A, SMC3, SP1, SP4, STAT3, STAT5A, TAF1, TAF7, TBL1XR1, TBP, TCF12, TCF7L2, TEAD4, TRIM28, UBTF, USF1, USF2, WRNIP1, YY1, ZBTB7A, ZC3H11A, ZEB1, ZMIZ1, ZNF143, ZNF263, ZNF384 |
| *ACOT11* | ELK1, STAT3, TCF4, TP63, FLI1, RUNX1, KLF4, EGR1, MITF, GATA1, REST, MYCN, SOX9, YAP1, DMRT1, KLF1, TCF7, CTCF, RAD21, BACH1, ATF1, BHLHE40, CBX3, CEBPB, CHD2, CTCFL, EP300, EZH2, FOSL2, GATA2, GATA3, H2AFZ, HDAC1, HDAC2, HDAC6, JUND, KDM5A, KDM5B, MAFK, MAX, MAZ, MYC, MYOD1, MYOG, NFIC, PBX3, PHF8, POLR2A, RBBP5, RCOR1, RXRA, SIN3A, SMC3, SPI1, SUPT20H, SUZ12, TAF1, TBP, TCF12, TEAD4, USF1, USF2, WHSC1, YY1, ZBTB7A, ZNF143 |
| *RTP4* | E2F2, PAX3, NFE2L2, AR, STAT3, NANOG, POU5F1, SOX2, TCF3, SOX9, ZFP42, NACC1, NR0B1, IRF8, ATF1, ATF2, BCLAF1, BHLHE40, CEBPB, CHD1, CHD2, CTCF, CUX1, EBF1, ELK1, EP300, EZH2, H2AFZ, HDAC2, IKZF1, IRF1, IRF3, IRF4, MAFF, MAFK, MAX, MTA3, MXI1, MYC, NFIC, NFYB, NRF1, PML, POLR2A, POU2F2, RBBP5, RCOR1, RFX5, RUNX3, SIN3A, SMC3, SPI1, SRF, STAT1, STAT2, TAF1, TBP, USF2, WRNIP1, YY1, ZC3H11A, ZKSCAN1, ZNF143, ZNF384 |
| *MERTK* | ZNF217, AR, TCF4, TP63, SPI1, SOX2, MITF, TP53, FOXA2, HNF4A, GATA2, TET1, ERG, TRIM28, SOX9, WT1, STAT4, CTCF, RAD21, PHC1, CEBPB, NR1H3, ARID3A, BACH1, BHLHE40, BRCA1, CCNT2, CEBPD, CHD1, CHD2, CHD7, CREB1, CTBP2, CTCFL, CUX1, E2F6, EBF1, EGR1, ELF1, ELK1, EP300, ETS1, EZH2, FOS, FOXP2, GABPA, GTF2F1, H2AFZ, HCFC1, HDAC1, HDAC2, HDAC6, HMGN3, HNF4G, IRF1, JUND, KDM1A, KDM4A, KDM5A, KDM5B, MAFF, MAFK, MAX, MAZ, MBD4, MXI1, MYBL2, MYC, MYOD1, MYOG, NANOG, NFIC, NRF1, PHF8, POLR2A, RBBP5, RCOR1, REST, RFX5, RNF2, SAP30, SIN3A, SMC3, SP1, SP4, SRF, STAT1, STAT3, SUZ12, TAF1, TAF7, TBL1XR1, TBP, TCF12, TCF3, TEAD4, UBTF, USF2, WRNIP1, YY1, ZBTB7A, ZC3H11A, ZMIZ1, ZNF143, ZNF263, ZNF384 |
| *STYK1* | NFE2L2, STAT3, RUNX1, MYC, SOX2, TP53, PPARG, SIN3B, EOMES, SALL4, SOX17, MYBL2, STAT4, MEIS1, STAT1, ATF2, BACH1, BATF, BCLAF1, BHLHE40, BRCA1, CBX3, CEBPB, CHD1, CHD2, CHD7, CREB1, CTBP2, CTCF, CUX1, E2F4, E2F6, ELF1, ELK1, EP300, EZH2, FOS, FOXM1, GATA1, GATA2, GTF2F1, H2AFZ, HDAC1, HDAC2, IRF1, IRF4, JUND, KDM1A, KDM4A, KDM5A, KDM5B, MAFF, MAFK, MAX, MAZ, MTA3, MXI1, NFATC1, NFIC, PHF8, PML, POLR2A, RAD21, RBBP5, RCOR1, REST, RFX5, RUNX3, SAP30, SIN3A, SMC3, SP1, SPI1, SRF, STAT5A, TAF1, TAL1, TBL1XR1, TBP, TCF12, TCF7L2, TEAD4, TRIM28, UBTF, USF1, USF2, WRNIP1, YY1, ZBTB7A, ZEB1, ZKSCAN1, ZNF143, ZNF263, ZNF384 |
| *HHIP* | STAT3, TCF4, TP63, CREM, RUNX1, MYC, NANOG, POU5F1, SOX2, TCF3, EGR1, GATA2, E2F4, TRIM28, EP300, SMARCA4, YAP1, SALL4, RUNX2, PHC1, EZH2, RNF2, MTF2, SUZ12, TBX3, NACC1, GLI1, ATF1, BACH1, BHLHE40, CHD1, CHD2, CREB1, CTBP2, CTCF, CUX1, E2F6, EBF1, ESR1, FOS, GATA3, H2AFZ, HDAC2, HDAC6, JUND, KDM4A, KDM5B, MAX, MAZ, MXI1, NRF1, PAX5, PHF8, PML, POLR2A, RAD21, RBBP5, RCOR1, SAP30, SIN3A, SPI1, STAT5A, TAL1, TCF12, TEAD4, UBTF, WRNIP1, ZBTB7A, ZMIZ1, ZNF143, ZNF384 |
| *CPLX1* | TCF4, CREM, RUNX1, NANOG, POU5F1, SOX2, EGR1, HNF4A, REST, TRIM28, SRY, KLF1, RUNX2, SUZ12, JUN, NR0B1, BACH1, BHLHE40, CCNT2, CHD1, CHD2, CREB1, CREBBP, CTBP2, CTCF, E2F6, ELF1, EP300, ETS1, EZH2, FOXP2, GABPA, H2AFZ, HDAC2, HMGN3, IRF1, JUND, KDM4A, KDM5A, KDM5B, MAFF, MAFK, MAX, MAZ, MXI1, MYC, PAX5, PHF8, POLR2A, RAD21, RBBP5, RCOR1, RNF2, RXRA, SAP30, SIN3A, SP1, TBP, TEAD4, UBTF, YY1, ZBTB7A, ZNF143 |
| *PI16* | TP63, FLI1, RUNX1, SPI1, SOX2, EGR1, MITF, GATA1, GATA2, PPARD, ERG, GFI1B, TAL1, SOX9, TEAD4, CUX1, YY1, MECOM, FOXP2, JARID2, SUZ12, MYB, LMO2, LYL1, MEIS1, BCL3, BHLHE40, CBX2, CBX8, CHD1, CTCF, EBF1, EP300, ESR1, ETS1, EZH2, FOS, FOSL1, FOSL2, GATA3, H2AFZ, HDAC2, HDAC6, HMGN3, JUN, JUND, KDM5A, KDM5B, MAFK, MAX, MAZ, MXI1, MYC, MYOG, PBX3, PHF8, POLR2A, RAD21, RBBP5, RCOR1, RNF2, SAP30, SIN3A, SMC3, STAT3, TCF12, ZNF263, ZNF384 |
| *ULBP3* | HOXC9, AR, SMAD2, SMAD3, BACH1, BRCA1, CBX3, CCNT2, CEBPB, CHD1, CHD2, CREB1, CTCF, E2F4, E2F6, EBF1, EGR1, ELK1, EP300, EZH2, FOS, FOSL2, FOXP2, GABPA, GTF2F1, H2AFZ, HCFC1, HDAC2, HMGN3, IRF1, JUND, KDM4A, KDM5A, KDM5B, MAX, MAZ, MXI1, MYC, NFYA, NFYB, PAX5, PBX3, PHF8, POLR2A, RAD21, RBBP5, RCOR1, REST, RFX5, SAP30, SIN3A, SMC3, SP1, STAT3, SUZ12, TAF1, TAF7, TBP, TCF12, TCF7L2, TEAD4, UBTF, USF1, YY1, ZBTB33, ZBTB7A, ZKSCAN1, ZNF143, ZNF263 |
| *PTCH1* | FOXP3, AR, SMAD4, TP63, E2F1, FLI1, MYC, SPI1, NANOG, POU5F1, SOX2, TCF3, EGR1, MITF, FOXA2, GATA2, RCOR3, SIN3B, REST, SETDB1, ASH2L, EOMES, FOXP1, PPARD, TRIM28, EP300, OLIG2, YAP1, SALL4, WT1, ZNF281, TEAD4, PRDM14, KLF1, ATF3, RUNX2, CCND1, HOXB4, MYBL2, ZFP42, STAT4, FOXP2, TCF7, MTF2, SUZ12, CTNNB1, TBX3, SIN3A, MYB, HSF1, TBP, NACC1, NR0B1, ZIC3, SMAD1, BCL3, PADI4, RBPJ, IRF8, BACH1, BHLHE40, CCNT2, CHD1, CHD2, CHD4, CHD7, CTBP2, CTCF, E2F4, E2F6, ELF1, ETS1, EZH2, FOXA1, GATA1, GTF2F1, H2AFZ, HCFC1, HDAC1, HDAC2, HMGN3, JUN, JUND, KAT2A, KDM4A, KDM5A, MAFK, MAX, MAZ, MXI1, MYOG, NELFE, PBX3, PHF8, POLR2A, RAD21, RBBP5, RCOR1, SAP30, SMC3, STAT3, TAF1, TAL1, TBL1XR1, TCF12, UBTF, USF1, USF2, YY1, ZBTB33, ZBTB7A, ZC3H11A, ZKSCAN1, ZMIZ1, ZNF143, ZNF263, ZNF384 |

| | |
|---|---|
| *DISC1* | ATF4, ATF5, AHR, ARNT, ZNF217, TTF2, AR, STAT3, TCF4, SMAD4, TP63, RUNX1, SPI1, NANOG, SOX2, TCF3, MITF, TP53, FOXA2, HNF4A, GATA1, MYCN, TAL1, TRIM28, SRY, EP300, SCLY, MEF2A, FOXP2, TCF7, BMI1, PHC1, EZH2, RNF2, JARID2, SUZ12, NR0B1, GATA3, DROSHA, ARID3A, ATF1, ATF2, BACH1, BATF, BCLAF1, BHLHE40, CBX2, CCNT2, CEBPB, CEBPD, CHD1, CHD2, CHD7, CREB1, CTBP2, CTCF, E2F4, E2F6, EBF1, EGR1, ELF1, FOS, FOSL2, FOXA1, GABPA, GATA2, GTF2F1, H2AFZ, HDAC1, HDAC2, HDAC6, HMGN3, HNF4G, IRF4, JUN, JUND, KDM4A, KDM5B, MAFF, MAX, MAZ, MBD4, MTA3, MXI1, MYBL2, MYC, NFIC, NR3C1, NRF1, PAX5, PHF8, POLR2A, RAD21, RBBP5, RCOR1, REST, RFX5, RUNX3, RXRA, SAP30, SIN3A, SMC3, SP1, SP4, TAF1, TBL1XR1, TBP, TCF12, TCF7L2, TEAD4, USF1, YY1, ZBTB7A, ZEB1, ZNF143, ZNF263 |
| *CPNE1* | AR, STAT3, CREB1, E2F1, FLI1, SPI1, FOXA2, ZFX, FOXP1, CNOT3, TRIM28, TFAP2A, CTNNB1, PBX1, PADI4, ARID3A, ATF1, ATF2, ATF3, BACH1, BCL3, BCLAF1, BHLHE40, BRCA1, CBX3, CCNT2, CEBPB, CEBPD, CHD1, CHD2, CHD7, CTBP2, CTCF, CUX1, E2F4, E2F6, EBF1, EGR1, ELF1, ELK1, ELK4, EP300, ETS1, EZH2, FOS, FOSL2, FOXA1, FOXM1, FOXP2, GABPA, GATA1, GATA3, GTF2B, GTF2F1, H2AFZ, HCFC1, HDAC1, HDAC2, HDAC6, HMGN3, IRF1, JUN, JUND, KAT2A, KAT2B, KDM1A, KDM4A, KDM5A, KDM5B, MAFF, MAFK, MAX, MAZ, MBD4, MTA3, MXI1, MYBL2, MYC, NELFE, NFATC1, NFIC, NR2F2, NR3C1, NRF1, PAX5, PHF8, PML, POLR2A, POU2F2, RAD21, RBBP5, RCOR1, RELA, REST, RFX5, RUNX3, SAP30, SETDB1, SIN3A, SIRT6, SIX5, SMARCB1, SMC3, SP1, SP4, SRF, STAT1, STAT5A, SUZ12, TAF1, TAF7, TBL1XR1, TBP, TCF12, TCF3, TCF7L2, TEAD4, THAP1, UBTF, USF1, USF2, WHSC1, WRNIP1, YY1, ZBTB33, ZBTB7A, ZC3H11A, ZEB1, ZKSCAN1, ZMIZ1, ZNF143, ZNF263, ZNF384 |
| *ARRDC2* | VDR, DACH1, STAT3, CREM, E2F1, FLI1, RUNX1, MYC, SPI1, KLF4, POU5F1, SOX2, EGR1, MITF, FOXA2, HNF4A, GATA1, GATA2, ZFX, SETDB1, FOXP1, TAL1, CNOT3, TRIM28, FOXO3, ESR1, KLF1, NR3C1, STAT4, FOXP2, RCOR1, SMAD1, TAF7L, ELF1, STAT6, ESR2, ATF1, ATF2, ATF3, BACH1, BCL3, BCLAF1, BHLHE40, BRCA1, CBX3, CCNT2, CEBPB, CEBPD, CHD1, CHD2, CHD4, CHD7, CREB1, CTCF, CUX1, E2F4, E2F6, EBF1, ELK1, ELK4, EP300, ETS1, EZH2, FOS, FOSL2, FOXA1, FOXM1, GABPA, GATA3, GTF2B, GTF2F1, GTF3C2, H2AFZ, HCFC1, HDAC1, HDAC2, HDAC6, HMGN3, IRF1, JUN, JUND, KAT2A, KDM1A, KDM4A, KDM5A, KDM5B, MAFK, MAX, MAZ, MTA3, MXI1, MYOG, NCOR1, NELFE, NFIC, NR2F2, NRF1, PAX5, PBX3, PHF8, PML, POLR2A, POU2F2, RAD21, RBBP5, RELA, REST, RFX5, RUNX3, SAP30, SIN3A, SIRT6, SMARCB1, SMARCC1, SMC3, SP1, SRF, STAT1, STAT5A, SUZ12, TAF1, TBL1XR1, TBP, TCF12, TCF3, TCF7L2, TEAD4, THAP1, UBTF, USF1, USF2, WHSC1, YY1, ZBTB7A, ZC3H11A, ZEB1, ZKSCAN1, ZMIZ1, ZNF143, ZNF274, ZNF384 |
| *CTSS* | VDR, STAT3, TP63, SPI1, POU5F1, SOX2, GATA1, GATA2, KLF1, MYBL2, RELA, ELF1, CRX, IRF8, ARID3A, ATF2, ATF3, BATF, BCL3, BCLAF1, BHLHE40, CCNT2, CEBPB, CHD1, CREB1, CTCF, CUX1, EBF1, EP300, ETS1, FOS, FOSL2, GATA3, GTF3C2, H2AFZ, HCFC1, HDAC6, IKZF1, IRF1, IRF4, JUN, KAT2B, KDM1A, KDM5A, KDM5B, MAFK, MAX, MEF2A, MXI1, MYC, NELFE, NFE2, NFIC, PAX5, PHF8, PML, POLR2A, POLR3A, RAD21, RBBP5, RCOR1, REST, RUNX3, SAP30, SIN3A, SP1, STAT1, TAF1, TBL1XR1, TBP, TCF12, TCF7L2, TEAD4, USF1, USF2, WRNIP1, ZMIZ1, ZNF143, ZNF384 |
| *ANGPT4* | AR, TP63, MYC, GATA1, GATA2, RCOR3, SIN3B, REST, TFAP2C, TET1, SOX9, ESR1, BMI1, HOXD13, CEBPB, CTCF, EZH2, H2AFZ, HDAC1, HDAC2, HDAC6, JUND, KDM5B, POLR2A, RBBP5, USF1 |
| *TFEC* | ETS2, SPI1, AR, STAT3, RUNX1, PPARD, OLIG2, SMARCA4, CUX1, MYB, POU3F2, RELA, BACH1, CEBPB, PRDM5, CTCF, EP300, EZH2, FOS, GATA2, GATA3, H2AFZ, MAFK, MEF2A, NFIC, POLR2A, RFX5, SAP30, TBL1XR1 |
| *COLEC11* | HOXC9, AR, TCF4, SMAD4, TP63, FLI1, RUNX1, EGR1, MITF, FOXA2, HNF4A, GATA2, PPARD, TEAD4, BMI1, ARID3A, ATF2, BACH1, BCL11A, BCL3, BCLAF1, BHLHE40, BRCA1, CBX3, CCNT2, CEBPB, CHD1, CTCF, CUX1, E2F6, EBF1, ELF1, EP300, ETS1, EZH2, FOXM1, FOXP2, GABPA, GATA1, H2AFZ, HCFC1, HDAC1, HDAC2, HMGN3, IRF1, JUND, KDM1A, KDM4A, KDM5A, KDM5B, MAX, MAZ, MTA3, MXI1, MYC, NFATC1, NFIC, NFYB, NR2F2, PAX5, PHF8, PML, POLR2A, RAD21, RBBP5, RCOR1, RELA, REST, RNF2, RUNX3, SMC3, SP1, SPI1, STAT5A, TAF1, TAL1, TBL1XR1, TBP, TCF12, THAP1, UBTF, USF1, YY1, ZBTB7A, ZC3H11A, ZMIZ1, ZNF143, ZNF384 |
| *MAST1* | TCF4, RUNX1, MYC, KLF4, NANOG, POU5F1, SOX2, EGR1, MITF, PPARG, HNF4A, KDM5B, SIN3B, REST, FOXP1, MYCN, ZNF281, TBX5, FOXP2, CTCF, EZH2, SUZ12, MEIS1, NR4A2, ARID3A, BACH1, BCL3, BHLHE40, CBX3, CCNT2, CEBPB, CEBPD, CHD1, CHD2, CHD7, CREB1, CREBBP, CTCFL, CUX1, E2F4, E2F6, ELF1, EP300, ETS1, FOS, FOSL1, GATA1, GATA2, GATA3, GTF2F1, H2AFZ, HCFC1, HDAC1, HDAC2, HDAC6, HMGN3, IRF1, JUN, JUND, KAT2B, KDM4A, MAFK, MAX, MAZ, MXI1, NFIC, NR2F2, PAX5, PBX3, PHF8, POLR2A, RAD21, RBBP5, RCOR1, RFX5, RNF2, RUNX3, SAP30, SETDB1, SIN3A, SIRT6, SMC3, SP1, SP4, STAT3, TAF1, TAL1, TBL1XR1, TBP, TCF12, TCF3, TEAD4, UBTF, YY1, ZBTB7A, ZC3H11A, ZKSCAN1, ZMIZ1, ZNF143, ZNF263, ZNF384 |
| *CD47* | ZNF217, VDR, DACH1, STAT3, TP63, CREM, E2F1, FLI1, RUNX1, SPI1, KLF4, NANOG, POU5F1, SOX2, TCF3, MITF, TP53, PPARG, GATA1, REST, SETDB1, ASH2L, EOMES, TFAP2C, TET1, TAL1, TRIM28, SRY, FOXO3, SOX17, DMRT1, PRDM14, KLF1, NR3C1, SCLY, STAT4, MEF2A, SRF, POU3F2, BACH1, ZIC3, BCL3, TAF7L, CEBPB, DNAJC2, BCLAF1, BHLHE40, BRCA1, CBX3, CCNT2, CHD1, CHD2, CHD4, CHD7, CREB1, CTBP2, CTCF, CTCFL, CUX1, E2F4, E2F6, EBF1, EGR1, ELF1, ELK1, EP300, ETS1, EZH2, FOXA1, FOXA2, FOXM1, FOXP2, GABPA, GTF2F1, H2AFZ, HCFC1, HDAC1, HDAC2, HDAC6, HMGN3, IKZF1, IRF1, JUND, KAT2A, KDM1A, KDM4A, KDM5A, KDM5B, MAFK, MAX, MAZ, MTA3, MXI1, MYC, MYOD1, MYOG, NELFE, NFATC1, NFYB, NRF1, PAX5, PHF8, PML, POLR2A, POU2F2, RAD21, RBBP5, RCOR1, RELA, RFX5, RUNX3, SAP30, SIN3A, SMARCB1, SMC3, SP1, STAT1, STAT5A, TAF1, TBL1XR1, TBP, TCF12, TCF7L2, UBTF, USF1, USF2, WRNIP1, YY1, ZBTB7A, ZEB1, ZKSCAN1, ZMIZ1, ZNF143, ZNF384 |

| | |
|---|---|
| *MEGF6* | AR, TP63, FLI1, MYC, SPI1, EGR1, MITF, HNF4A, GATA2, FOXP1, SOX9, SUZ12, BHLHE40, CHD1, CTCF, E2F6, EZH2, FOS, GABPA, H2AFZ, HDAC2, HDAC6, HMGN3, IRF1, JUND, KDM4A, KDM5A, KDM5B, MAX, MAZ, MXI1, NFIC, PHF8, POLR2A, REST, RNF2, SAP30, SIN3A, TAF1, TBP, TCF12, UBTF, USF1, ZBTB7A, ZNF263 |
| *HECW2* | AHR, ARNT, ZNF217, STAT3, TCF4, SMAD4, TP63, FLI1, RUNX1, SPI1, TP53, GATA1, CNOT3, TFCP2L1, RUNX2, FOXP2, BMI1, JARID2, MTF2, SUZ12, CTNNB1, JUN, CDX2, STAT6, BACH1, BCLAF1, BRCA1, CCNT2, CEBPB, CHD1, CHD2, CHD7, CREB1, CTBP2, CTCF, E2F6, EGR1, ELF1, ELK1, EP300, EZH2, GABPA, GATA2, GATA3, GTF2F1, H2AFZ, HCFC1, HDAC1, HDAC2, HDAC6, HMGN3, JUND, KDM4A, KDM5A, KDM5B, MAFF, MAFK, MAX, MAZ, MXI1, MYC, MYOD1, MYOG, PAX5, PHF8, POLR2A, RAD21, RBBP5, RCOR1, REST, RFX5, RUNX3, SIN3A, SMARCB1, SMC3, SP4, TAF1, TAF7, TBP, TCF12, TCF7L2, TEAD4, TRIM28, UBTF, YY1, ZBTB7A, ZEB1, ZKSCAN1, ZNF143, ZNF263 |
| *TMEM185B* | TP63, CREM, E2F1, FLI1, MYC, SPI1, POU5F1, ZFX, SIN3B, ASH2L, EOMES, TFAP2C, ERG, GFI1B, TAL1, EP300, SOX17, DMRT1, SCLY, RNF2, SUZ12, ATF2, ATF3, BACH1, BCL3, BCLAF1, BHLHE40, BRCA1, CBX3, CCNT2, CEBPB, CEBPD, CHD1, CHD2, CHD7, CREB1, CTBP2, CTCF, CTCFL, E2F4, E2F6, EBF1, EGR1, ELF1, ELK1, ELK4, ESRRA, ETS1, EZH2, FOSL1, FOSL2, FOXM1, FOXP2, GABPA, GATA1, GATA2, GATA3, GTF2B, GTF2F1, H2AFZ, HCFC1, HDAC1, HDAC2, HDAC6, HMGN3, HNF4A, IRF1, JUN, JUND, KAT2A, KAT2B, KDM1A, KDM4A, KDM5A, KDM5B, MAFF, MAFK, MAX, MAZ, MTA3, MXI1, MYBL2, MYOD1, MYOG, NANOG, NELFE, NFATC1, NFE2, NFIC, NFYA, NFYB, NR2F2, NR3C1, NRF1, PAX5, PHF8, PML, POLR2A, POU2F2, RAD21, RBBP5, RCOR1, REST, RFX5, RUNX3, SAP30, SETDB1, SIN3A, SMARCB1, SMC3, SP1, SP4, STAT1, STAT3, STAT5A, TAF1, TAF7, TBL1XR1, TBP, TCF12, TCF3, TCF7L2, TEAD4, TRIM28, UBTF, USF1, USF2, WHSC1, WRNIP1, YY1, ZC3H11A, ZEB1, ZKSCAN1, ZMIZ1, ZNF143, ZNF263, ZNF384 |
| *SCGB1A1* | JUN, PGR, POU2F1, SMAD4, SPI1, NANOG, SOX2, TP53, PPARG, GATA1, GATA2, ASH2L, PPARD, ERG, TAL1, EP300, ESR1, SMAD3, CUX1, YY1, RAD21, TFAP2A, CEBPB, BCL3, ELF1, EZH2, H2AFZ, HDAC1, HDAC6, JUND, KDM5B, NR2F2, PHF8, PML, POLR2A, RBBP5, RCOR1, SAP30, SETDB1, TEAD4, TRIM28, ZNF263 |
| *IL12A* | NFE2L2, ELK1, STAT3, FLI1, RUNX1, MYC, SOX2, GATA2, ASH2L, PPARD, ERG, SMARCA4, MECOM, CTNNB1, MYB, LMO2, RELA, BACH1, ARID3A, ATF2, BATF, BCL3, BCLAF1, BHLHE40, BRCA1, CBX3, CCNT2, CEBPB, CHD1, CHD2, CHD7, CREB1, CTBP2, CTCF, CTCFL, E2F4, E2F6, EBF1, EGR1, ELF1, EP300, EZH2, FOS, FOXA1, FOXM1, GABPA, GATA1, GATA3, GTF2F1, H2AFZ, HCFC1, HDAC1, HDAC2, HDAC6, HMGN3, IKZF1, IRF1, IRF4, JUN, JUND, KDM4A, KDM5B, MAFF, MAFK, MAX, MAZ, MEF2A, MEF2C, MTA3, MXI1, MYBL2, NFIC, NRF1, PAX5, PHF8, POLR2A, POU2F2, RAD21, RBBP5, RCOR1, RFX5, RUNX3, SAP30, SETDB1, SIN3A, SMC3, SP1, SP4, SRF, STAT1, STAT5A, SUZ12, TAF1, TBL1XR1, TBP, TCF12, TCF3, TCF7L2, TEAD4, USF1, USF2, WRNIP1, YY1, ZBTB7A, ZEB1, ZNF143, ZNF263 |
| *HIST1H3H* | HOXC9, VDR, CREB1, MYC, SPI1, KLF4, SOX2, EGR1, GATA1, E2F4, EOMES, TAL1, CNOT3, SALL4, CUX1, CCND1, HOXB4, MYBL2, CHD1, ZFP42, FOXP2, RNF2, POU3F2, HSF1, TBP, ZIC3, ARID3A, ATF1, ATF2, ATF3, BACH1, BATF, BCL11A, BCL3, BCLAF1, BHLHE40, BRCA1, CBX3, CCNT2, CEBPB, CEBPD, CHD2, CHD7, CREBBP, CTCF, E2F6, EBF1, ELF1, ELK1, ELK4, EP300, ETS1, EZH2, FLI1, FOS, FOSL1, FOXA1, FOXA2, FOXM1, GABPA, GATA2, GATA3, GTF2B, GTF2F1, H2AFZ, HCFC1, HDAC1, HDAC2, HDAC6, HMGN3, IKZF1, IRF1, IRF3, IRF4, JUN, JUND, KAT2A, KAT2B, KDM4A, KDM5A, KDM5B, MAFF, MAFK, MAX, MAZ, MBD4, MEF2A, MTA3, MXI1, NANOG, NCOR1, NELFE, NFATC1, NFIC, NFYA, NFYB, NR2F2, NR3C1, NRF1, PAX5, PBX3, PHF8, PML, POLR2A, POU2F2, RAD21, RBBP5, RCOR1, RELA, REST, RFX5, RUNX3, RXRA, SAP30, SETDB1, SIN3A, SIRT6, SIX5, SMC3, SP1, SP2, SP4, STAT1, STAT3, STAT5A, SUZ12, TAF1, TAF7, TBL1XR1, TCF12, TCF3, TCF7L2, TEAD4, THAP1, TRIM28, UBTF, USF1, USF2, WHSC1, WRNIP1, YY1, ZBTB33, ZC3H11A, ZEB1, ZKSCAN1, ZMIZ1, ZNF143, ZNF384 |
| *BTN3A2* | ZNF217, FLI1, MITF, FOXA2, HNF4A, GATA1, ARID3A, ATF1, ATF2, BACH1, BCLAF1, BHLHE40, BRCA1, CBX3, CEBPB, CHD1, CHD2, CREB1, CTCF, CUX1, EBF1, ELF1, ELK1, EP300, EZH2, FOS, FOXA1, FOXM1, FOXP2, GABPA, GTF2F1, H2AFZ, HCFC1, HDAC2, HNF4G, IRF1, IRF4, JUND, KDM1A, KDM4A, KDM5B, MAFF, MAFK, MAX, MAZ, MTA3, MXI1, MYBL2, MYC, NFE2, NFIC, NFYA, NFYB, NRF1, PAX5, PHF8, PML, POLR2A, POU2F2, PRDM1, RAD21, RBBP5, RCOR1, REST, RFX5, RUNX3, SAP30, SIN3A, SMC3, SP1, SPI1, STAT1, STAT3, TAF1, TBL1XR1, TBP, TCF12, TCF7L2, TEAD4, UBTF, USF1, USF2, WHSC1, WRNIP1, YY1, ZBTB33, ZC3H11A, ZMIZ1, ZNF143, ZNF263, ZNF384 |
| *SORBS1* | AR, TCF4, SMAD4, TP63, E2F1, MYC, SPI1, NANOG, POU5F1, SOX2, TCF3, MITF, TP53, PPARG, FOXA2, HNF4A, GATA1, GATA2, RCOR3, SIN3B, REST, EOMES, TFAP2C, TRIM28, EP300, SALL4, PRDM14, KLF1, TFCP2L1, RUNX2, CUX1, SCLY, YY1, GATA4, EZH2, RNF2, CTNNB1, PBX1, NUCKS1, TBP, NACC1, NR0B1, CDX2, ARID3A, BACH1, BHLHE40, BRCA1, CBX3, CCNT2, CEBPB, CEBPD, CHD1, CHD2, CREB1, CTBP2, CTCF, E2F4, E2F6, EBF1, EGR1, ELF1, ELK1, ELK4, ETS1, FOS, FOXA1, FOXP2, GABPA, GTF2F1, H2AFZ, HCFC1, HDAC1, HDAC2, HDAC6, HMGN3, HNF4G, IRF1, JUN, JUND, KDM1A, KDM4A, KDM5A, KDM5B, MAFF, MAFK, MAX, MAZ, MXI1, MYOG, NELFE, NFIC, NFYA, NFYB, NR2F2, NRF1, PAX5, PHF8, PML, POLR2A, POU2F2, RAD21, RBBP5, RCOR1, RFX5, SAP30, SETDB1, SIN3A, SIRT6, SMARCB1, SMC3, SP1, SP4, SRF, STAT1, STAT3, TAF1, TAF7, TAL1, TBL1XR1, TCF12, TCF7L2, THAP1, UBTF, USF1, USF2, ZBTB7A, ZEB1, ZKSCAN1, ZMIZ1, ZNF143, ZNF384 |
| *SNAP91* | TFAP2A, TCF4, CREM, RUNX1, POU5F1, PPARG, KDM5B, TFAP2C, EED, EZH2, RNF2, JARID2, MTF2, SUZ12, BACH1, BHLHE40, CEBPB, CHD1, CHD2, CHD7, CREB1, CTBP2, CTCF, E2F6, EBF1, EGR1, ELF1, EP300, GTF2F1, H2AFZ, HCFC1, HDAC2, JUND, KDM4A, KDM5A, MAX, MAZ, MXI1, MYC, MYOG, NANOG, NRF1, PHF8, POLR2A, RAD21, RBBP5, REST, SAP30, SIN3A, TAF1, TBP, TCF12, TCF7L2, TRIM28, USF1, USF2, YY1, ZBTB7A, ZNF143, ZNF263 |
| *LINC00242* | TTF2, TCF4, E2F1, RUNX1, RUNX2, SCLY, HIF1A, ARID3A, BACH1, BCL3, BHLHE40, BRCA1, CBX2, CBX8, CEBPB, CEBPD, CHD1, CTCF, CUX1, ELF1, EP300, EZH2, FOS, FOSL1, FOSL2, FOXA1, FOXA2, GATA3, H2AFZ, HDAC2, HNF4A, HNF4G, |

| | |
|---|---|
| | IRF3, JUN, JUND, KDM5B, MAFF, MAFK, MAX, MBD4, MXI1, MYBL2, MYC, NFIC, NR3C1, POLR2A, RAD21, RCOR1, RFX5, RNF2, SMC3, SP1, SRF, STAT3, TBP, TCF12, TCF7L2, TEAD4, TRIM28, UBTF |
| *LINC00574* | TP63, RUNX1, HNF4A, MYCN, RUNX2, SCLY, HIF1A, ARID3A, ATF3, BACH1, BCLAF1, BHLHE40, BRCA1, CBX2, CBX3, CBX8, CCNT2, CEBPB, CEBPD, CHD1, CHD2, CHD7, CREB1, CTCF, CTCFL, CUX1, E2F4, E2F6, EBF1, ELF1, EP300, ESR1, EZH2, FOS, FOXM1, FOXP2, GABPA, GTF2F1, H2AFZ, HCFC1, HDAC1, HDAC2, HDAC6, HMGN3, HNF4G, JUN, JUND, KDM4A, KDM5A, KDM5B, MAFK, MAX, MAZ, MBD4, MTA3, MXI1, MYBL2, MYC, NFIC, NR2F2, NRF1, PAX5, PHF8, POLR2A, RAD21, RBBP5, RCOR1, REST, RFX5, RNF2, RUNX3, RXRA, SAP30, SETDB1, SIN3A, SMC3, SP1, SP4, SPI1, STAT1, STAT3, STAT5A, SUZ12, TAF1, TBL1XR1, TBP, TCF12, TCF7L2, TEAD4, TRIM28, UBTF, USF2, WRNIP1, YY1, ZBTB7A, ZC3H11A, ZEB1, ZKSCAN1, ZMIZ1, ZNF143, ZNF263, ZNF384 |
| *HOTTIP* | SPI1, BACH1, CBX8, CHD2, CREB1, CTBP2, CTCF, E2F6, EGR1, ELK1, EZH2, FOXA1, FOXA2, GABPA, GTF2F1, H2AFZ, HDAC2, HMGN3, KDM4A, KDM5B, MAFK, MAX, MAZ, MYC, NFYA, POLR2A, RAD21, RCOR1, REST, RFX5, SAP30, SIN3A, SMARCB1, SMC3, SUZ12, TAF1, TBP, TCF12, TCF7L2, TRIM28, WRNIP1, YY1, ZBTB7A, ZKSCAN1, ZNF143, ZNF263 |
| *HCG27* | EGR1, MITF, FOXA2, HNF4A, TFAP2C, ATF3 |
| *AFAP1-AS1* | ATF1, ATF3, BHLHE40, CBX3, CEBPB, CHD1, CHD2, CTCF, E2F6, ELF1, EP300, ETS1, EZH2, FOS, H2AFZ, HCFC1, HDAC1, HDAC2, HDAC6, JUND, KDM5B, MAX, MAZ, MYC, PHF8, PML, POLR2A, RCOR1, SAP30, SIN3A, SPI1, STAT3, TBL1XR1, TBP, USF1, USF2, YY1, ZBTB33, ZNF143, ZNF263 |
| *MYLK-AS1* | ARID3A, ATF2, BACH1, BATF, BCL3, BCLAF1, BHLHE40, BRCA1, CBX3, CCNT2, CEBPB, CEBPD, CHD1, CHD2, CHD7, CREB1, CTBP2, CTCF, CUX1, E2F1, E2F4, E2F6, EBF1, EGR1, ELF1, ELK1, EP300, ETS1, EZH2, FOS, FOSL2, FOXA1, FOXA2, FOXM1, FOXP2, GABPA, GATA1, GATA3, GTF2F1, H2AFZ, HCFC1, HDAC1, HDAC2, HDAC6, HMGN3, HNF4A, IKZF1, IRF1, IRF4, JUN, JUND, KDM1A, KDM4A, KDM5A, KDM5B, MAFF, MAFK, MAX, MAZ, MBD4, MTA3, MXI1, MYBL2, MYC, NANOG, NFATC1, NFIC, NFYB, NR3C1, NRF1, PAX5, PBX3, PHF8, PML, POLR2A, POU2F2, RAD21, RBBP5, RCOR1, RELA, REST, RFX5, RUNX3, SAP30, SIN3A, SIRT6, SIX5, SMARCB1, SMARCC1, SMC3, SP1, SP4, SPI1, STAT1, STAT3, STAT5A, TAF1, TAF7, TAL1, TBL1XR1, TBP, TCF12, TCF3, TCF7L2, TEAD4, TFAP2A, TFAP2C, THAP1, TRIM28, UBTF, USF1, USF2, WHSC1, WRNIP1, YY1, ZBTB7A, ZC3H11A, ZEB1, ZKSCAN1, ZMIZ1, ZNF143, ZNF263, ZNF384 |
| *DLX6-AS1* | BMI1, JARID2, SUZ12, BACH1, CBX8, CTBP2, CTCF, EZH2, POLR2A, REST, SP1, STAT3, TAF1, TCF12 |
| *RHPN1-AS1* | TP63, MITF, GATA1, PPARD, ERG, RUNX2, ARID3A, ATF3, BACH1, BCL3, BHLHE40, BRCA1, CBX3, CCNT2, CEBPB, CEBPD, CHD1, CHD2, CHD4, CHD7, CREB1, CTCF, CTCFL, CUX1, E2F4, E2F6, EBF1, EGR1, ELF1, ELK1, ELK4, EP300, ETS1, EZH2, FOS, FOSL2, FOXA1, FOXM1, FOXP2, GABPA, GTF2F1, H2AFZ, HCFC1, HDAC2, HMGN3, HNF4A, IRF1, IRF3, JUN, JUND, KAT2B, KDM1A, KDM4A, KDM5B, MAFK, MAX, MAZ, MTA3, MXI1, MYBL2, MYC, NFYA, NFYB, NR2F2, NR3C1, NRF1, PAX5, PBX3, PHF8, PML, POLR2A, RBBP5, RCOR1, REST, RFX5, RNF2, RUNX3, RXRA, SAP30, SETDB1, SIN3A, SIRT6, SMARCB1, SMARCC1, SMC3, SP1, SP2, SP4, SREBF1, SRF, STAT1, STAT3, STAT5A, SUZ12, TAF1, TAL1, TBL1XR1, TBP, TCF12, TCF3, TEAD4, TFAP2A, TFAP2C, THAP1, TRIM28, UBTF, USF1, USF2, WRNIP1, YY1, ZBTB7A, ZC3H11A, ZEB1, ZKSCAN1, ZMIZ1, ZNF143, ZNF263, ZNF274, ZNF384 |
